# The connectome of the adult *Drosophila* mushroom body: implications for function

**DOI:** 10.1101/2020.08.29.273276

**Authors:** Feng Li, Jack Lindsey, Elizabeth C. Marin, Nils Otto, Marisa Dreher, Georgia Dempsey, Ildiko Stark, Alexander Shakeel Bates, Markus William Pleijzier, Philipp Schlegel, Aljoscha Nern, Shinya Takemura, Tansy Yang, Audrey Francis, Amalia Braun, Ruchi Parekh, Marta Costa, Louis Scheffer, Yoshinori Aso, Gregory S. X. E. Jefferis, L.F. Abbott, Ashok Litwin-Kumar, Scott Waddell, Gerald M. Rubin

## Abstract

Making inferences about the computations performed by neuronal circuits from synapse-level connectivity maps is an emerging opportunity in neuroscience. The mushroom body (MB) is well positioned for developing and testing such an approach due to its conserved neuronal architecture, recently completed dense connectome, and extensive prior experimental studies of its roles in learning, memory and activity regulation. Here we identify new components of the MB circuit in *Drosophila*, including extensive visual input and MB output neurons (MBONs) with direct connections to descending neurons. We find unexpected structure in sensory inputs, in the transfer of information about different sensory modalities to MBONs, and in the modulation of that transfer by dopaminergic neurons (DANs). We provide insights into the circuitry used to integrate MB outputs, connectivity between the MB and the central complex and inputs to DANs, including feedback from MBONs. Our results provide a foundation for further theoretical and experimental work.

## Introduction

Dramatic increases in the speed and quality of imaging, segmentation and reconstruction in electron microscopy now allow large-scale, dense connectomic studies of nervous systems. Such studies can reveal the chemical synapses between all neurons, generating a complete connectivity map. Connectomics is particularly useful in generating biological insights when applied to an ensemble of neurons with interesting behavioral functions that have already been extensively studied experimentally. Knowing the effects on behavior and physiology of perturbing individual cell types that can also be unambiguously identified in the connectome is of considerable value. Here we present a connectomic analysis of one such neuronal ensemble, the mushroom body (MB) of adult *Drosophila melanogaster*.

Understanding how memories of past events are formed and then used to influence ongoing behavior are key challenges in neuroscience. It is generally accepted that parallel changes in connection strength across multiple circuits underlie the formation of a memory and that these changes are integrated to produce net changes in behavior. Animals learn to predict the value of sensory cues based on temporal correlations with reward or punishment (Pavlov and Thompson, 1902). Such associative learning entails lasting changes in connections between neurons (reviewed in Abraham et al., 2019; Martin et al., 2000). It is now clear that different parts of the brain process and store different aspects of the information learned in a single event (reviewed in Josselyn and Frankland, 2018). In both flies and mammals, dopaminergic neurons play a key role in conveying information about whether an event has a positive or negative valence and there are compelling parallels between the molecular diversity of dopaminergic cell types across these evolutionarily distant animals (Watabe-Uchida and Uchida, 2018). However, we have limited understanding of how information about the outside world or internal brain state reaches different dopaminergic populations. Nor do we understand the nature of the information that is stored in each parallel memory system or how these parallel memories interact to guide coherent behavior. We believe such processes are governed by general and evolutionarily conserved principles. In particular, we believe the circuit logic that allows a brain to balance the competing demands of rapid learning with long-term stability of memory are likely to be the same in flies and mammals. Developing a comprehensive understanding of these circuits at the resolution of individual neurons and synapses will require the synergistic application of a variety of experimental methods together with theory and modeling. Many of the required methods are well developed in *Drosophila,* where the circuits underlying learning and memory are less complex than in mammals, and where detailed anatomical knowledge of the relevant circuits, which we believe will be essential, has just now become available. Here we provide analysis of the complete connectome of a circuit involved in parallel processing of associative memories in adult fruit flies. The core architecture of this circuit is strikingly similar to that of the vertebrate cerebellum (Figure 1; Laurent, 2002; Farris, 2011; Litwin-Kumar et al., 2017).

**Figure 1.**
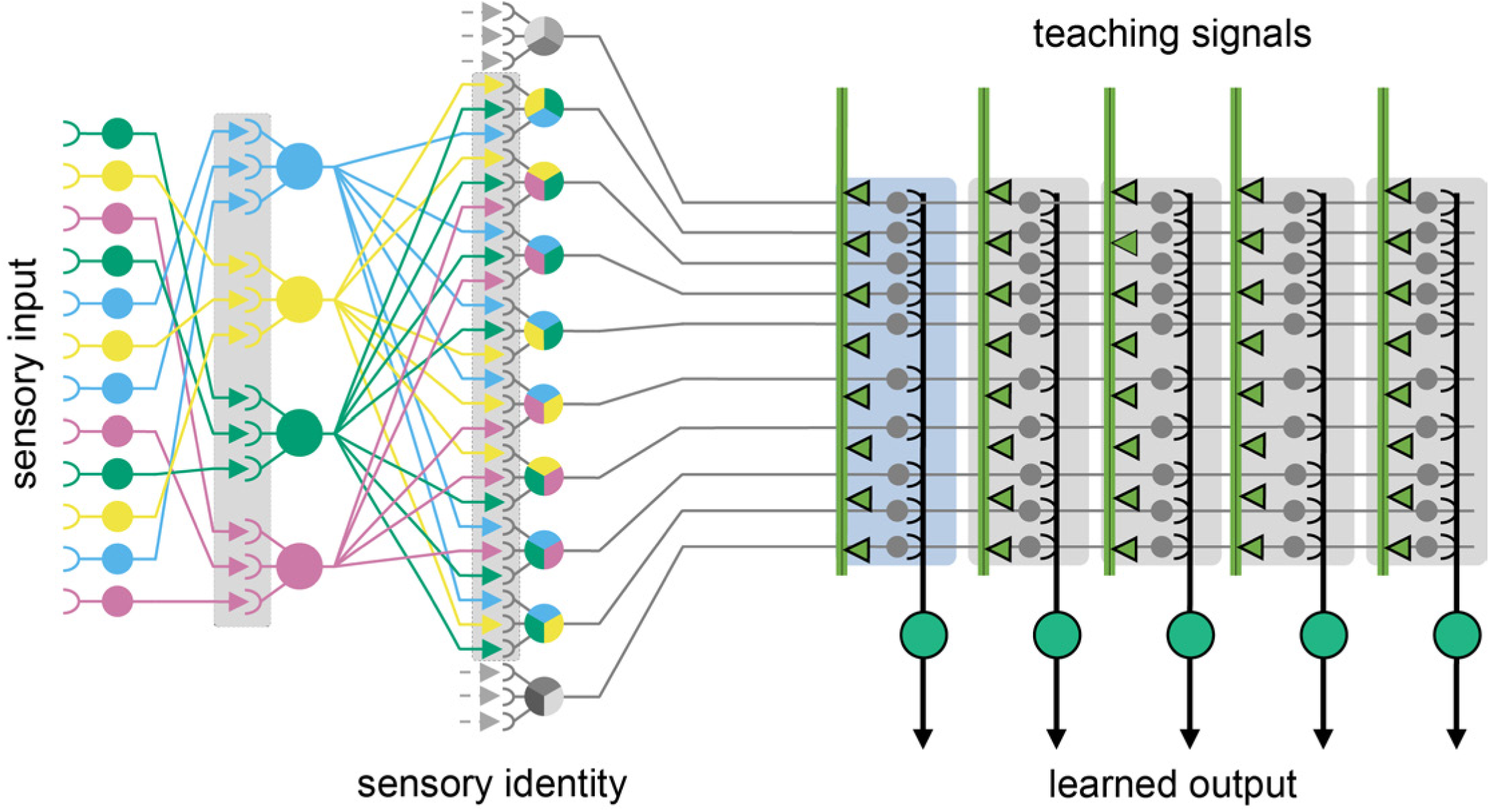
The shared circuit architecture of the mushroom body and the cerebellum. In both the insect MB and vertebrate cerebellum sensory information is represented by sparse activity in parallel axonal fibers — Kenyon cells (KCs) in the MB and granule cells (GC) in the cerebellum (reviewed in Modi et al., 2020). In general, each KC or GC has claw-like dendrites that integrate sensory input from a small number of neurons, called projection neurons in insects and mossy fibers in vertebrates. In the MB, teaching signals are provided by dopaminergic neurons (DANs) and in the cerebellum by climbing fibers. Learned output is conveyed to the rest of the brain from the MB lobes by MB output neurons (MBONs) or, from the cerebellar cortex, by Purkinje cells. The arbors of the DANs and MBONs overlap in the MB lobes and define a series of 15 compartments (Aso et al., 2014a; Gao et al., 2019); Figure 2; Video 2); similarly, overlap between the arbors of climbing fibers and Purkinje cells define zones along the GC parallel fibers.

The MB is the major site of associative learning in insects (reviewed in Heisenberg, 2003; Modi et al., 2020), and species that perform more complex behavioral tasks tend to have larger MBs (O’Donnell et al., 2004; Sivinski, 1989). In the MB of each brain hemisphere, sensory stimuli are represented by the sparse activity of ∼2000 Kenyon cells (KCs) whose dendrites form a structure called the MB calyx and whose parallel axonal fibers form the lobes of the MB (Figure 2; Video 1).

**Figure 2.**
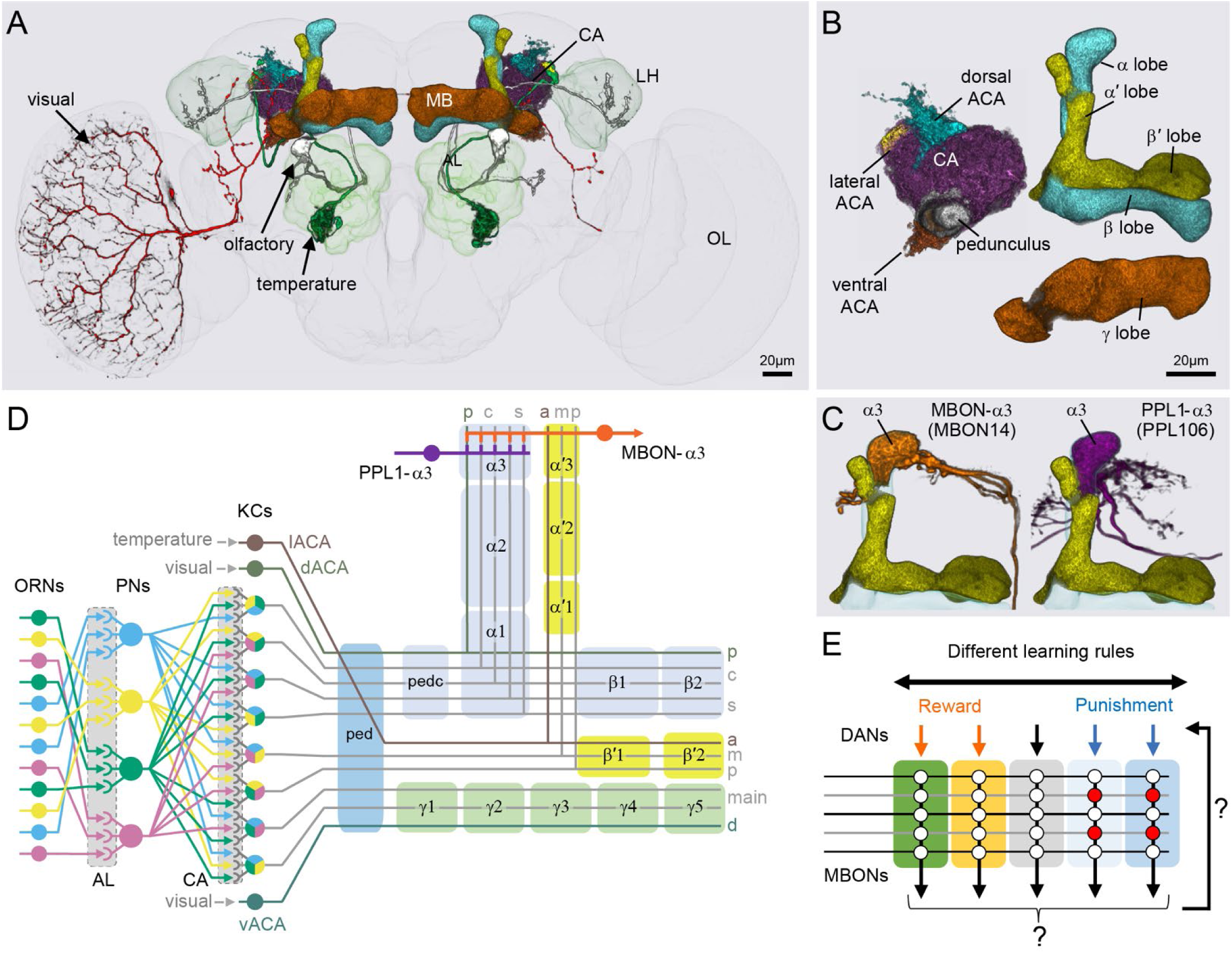
Anatomy of the adult Drosophila MB. Diagram of structure and information flow in the MB. (A) An image of the brain showing subregions of the MB (see panel B for more detail) and examples of the sensory pathways that provide information to the KCs. Projection neurons (PNs) from the 51 olfactory glomeruli of the antennal lobe (AL) extend projection neurons (PN) to the calyx (CA) of the MB and the lateral horn (LH). A total of 126 PNs, using a threshold of 25 synapses, innervate the CA and 3 innervate the lACA. Six olfactory PNs from the DL3 glomerulus are shown (white). Also shown is a visual projection neuron, aMe12 (red) that conveys information from the optic lobe (OL) to the ventral accessory calyx (vACA) and a thermosensory projection neuron (green) that conveys cold temperature information from arista sensory neurons in glomerulus VP3 to the lACA; the positions of the accessory calyces are shown in (B). See Video 1 for additional details. (B) Subregions within the MB. The γ lobe, CA, and pedunculus are displayed separately from other lobes; their normal positions are as shown in panel A. Color-coding is as in panel A. (C) The MB output neuron (MBON14) whose dendrites fill the α3 compartment at the tip of the vertical lobe is shown along with the dopaminergic neuron (PPL106), whose axonal terminals lie in the same compartment. See Video 2 for more detailed examples of the structure of a compartment. (D) A schematic representation of the key cellular components and information flow during processing of sensory inputs to the MB. Olfactory receptor neurons (ORNs) expressing the same odorant receptor converge onto a single glomerulus in the AL. A small number (generally 3 - 4) of PNs from each of the 51 olfactory glomeruli innervate the MB, where they synapse on the dendrites of the ∼2000 Kenyon cells (KCs) in a globular structure, the CA. Each KC exhibits, on average, 6 dendritic ‘claws’, and each claw is innervated by a single PN. The axons of the KCs project in parallel anteriorly through the pedunculus (ped) to the lobes, where KCs synapse onto the dendrites of MB output neurons (MBONs). KCs can be categorized into three major classes α/β, α*′*/β*′*, and γ, based on their projection patterns in the lobes (Crittenden et al., 1998). The β, β*′*, and γ lobes constitute the medial lobes (also known as horizontal lobes), while the α and α*′* lobes constitute the vertical lobes. These lobes are separately wrapped by ensheathing glia (Awasaki et al., 2008). The α/β and α*′*/β*′* neurons bifurcate at the anterior end of the ped (pedc) and project to both the medial and vertical lobes (Lee et al., 1999). The γ neurons project only to the medial lobe. Dendrites of MBONs and terminals of modulatory dopaminergic neurons (DANs) intersect the longitudinal axis of the KC axon bundle, forming 15 subdomains or compartments, five each in the α/β, α*′*/β*′*, and γ lobes (numbered α1, α2, and α3 for the compartments in the α lobe from proximal to distal and similarly for the other lobes; Aso et al., 2014a; Tanaka et al., 2008). Additionally, one MBON and one DAN innervate the core of the distal pedunculus (pedc) intersecting the α/β KCs. In the current work, we further classified KCs into 14 types, 10 main types and 4 unusual embryonic born KCs, named KCγs1-s4 (see Figure 3); the main KC types have their dendrites in the main calyx, with the following exceptions: The dendrites of γd KCs form the ventral accessory calyx (vACA; Aso et al., 2009; Butcher et al., 2012); those of the α/βp KCs form the dorsal accessory calyx (dACA; Lin et al., 2007; Tanaka et al., 2008); and the dendrites of a subset of α*′*/β*′* cells form the lateral accessory calyx (lACA) (Marin et al., 2020; Yagi et al., 2016). These accessory calyces receive non-olfactory input (Tanaka et al., 2008). Different KCs occupy distinct layers in the lobes as indicated (p: posterior; c: core; s: surface; a: anterior; m: middle; main and d: dorsal). Some MB extrinsic neurons extend processes only to a specific layer within a compartment. (E) Individual compartments serve as parallel units of memory formation (see Aso and Rubin, 2016). Reward or punishment is conveyed by dopaminergic neurons, and the coincidence of dopamine release with activity of a KC modifies the strength of that KC’s synapses onto the MBONs in that compartment. The circuit structure by which those MBONs combine their outputs to influence behavior and provide feedback to dopaminergic neurons are investigated in this paper.

The major sensory inputs to the *Drosophila* MB are olfactory, delivered by ∼150 projection neurons (PNs) from the antennal lobe to the dendrites of the KCs in the MB calyx (Bates et al., 2020b). KCs each receive input from an average of six PNs. For a KC to fire a spike, multiple of its PN inputs need to be simultaneously activated (Gruntman and Turner, 2013). This requirement, together with global feedback inhibition (Lin et al., 2014a; Papadopoulou et al., 2011), ensures a sparse representation where only a small percentage of KCs are activated by an odor (Honegger et al., 2011; Perez-Orive et al., 2002). The MB has a three layered divergent-convergent architecture (Huerta et al., 2004; Jortner et al., 2007; Laurent, 2002; Litwin-Kumar et al., 2017; Shomrat et al., 2011; Stevens, 2015) in which the coherent information represented by olfactory PNs is expanded and decorrelated when delivered to the KCs (Caron et al., 2013; Zheng et al., 2020). But the degree to which the structure of the sensory input representation is maintained by the KCs has been debated.

While best studied for its role in olfactory associative learning, the MB also receives inputs from several other sensory modalities. A subset of projection neurons from the antennal lobe delivers information about temperature and humidity in both the larva (Eichler et al., 2017) and the adult (Marin et al., 2020). Taste conditioning also requires the MB and is believed to depend on specific KC populations, although the relevant inputs to these KCs have not yet been reported (Kirkhart and Scott, 2015; Masek and Scott, 2010). We identified one likely path for gustatory input to the MB.

*Drosophila* MBs are also known to be able to form memories based on visual cues (Aso et al., 2014b; Brembs, 2009; Vogt et al., 2016, 2014; Liu et al., 1999; Zhang et al., 2007). Until a few years ago, it was thought that visual input reached the *Drosophila* MBs using only indirect, multisynaptic pathways (Farris and Van Dyke, 2015; Tanaka et al., 2008) as direct visual input from the optic lobes to the MBs, well known in Hymenoptera (Ehmer and Gronenberg, 2002), had not been observed in any dipteran insect (Mu et al., 2012; Otsuna and Ito, 2006). In 2016, Vogt et al. (2016) identified two types of visual projection neurons (VPNs) connecting the optic lobes and the MB and additional connections have been observed recently by light microscopy (Li et al., 2020). We found that visual input was much more extensive than previously appreciated, with about eight percent of KCs receiving predominantly visual input, and present here a detailed description of neuronal pathways connecting the optic lobe and the MB. Visual sensory input appears to be segregated into distinct KC populations in both the larva (Eichler et al., 2017) and the adult (Vogt et al., 2016; Li et al., 2020), as is the case in honeybees (Ehmer and Gronenberg, 2002). We found two classes of KCs that receive predominantly visual sensory input, as well as MBONs that get the majority of their input from these segregated KC populations.

MBONs provide the convergence element of the MB’s three layer divergent-convergent circuit architecture. Previous work has identified 22 types of MBONs whose dendrites receive input from specific axonal segments of the KCs. The outputs of the MBONs drive learned behaviors. Approximately 20 types of dopaminergic neurons (DANs) innervate corresponding regions along the KC axons and are required for associative olfactory conditioning. Specifically, the presynaptic arbors of the DANs and postsynaptic dendrites of the MBONs overlap in distinct zones along the KC axons, defining the 15 compartmental units of the MB lobes (Aso et al., 2014a; Mao and Davis, 2009; Takemura et al., 2017; Tanaka et al., 2008); Figure 2; Video 2). A large body of evidence indicates that these anatomically defined compartments of the MB are the units of associative learning (Aso et al., 2012, 2010, 2019, 2014a, 2014b; Aso and Rubin, 2016; Berry et al., 2018; Blum et al., 2009; Bouzaiane et al., 2015; Burke et al., 2012; Claridge-Chang et al., 2009; Isabel et al., 2004; Jacob and Waddell, 2020; Krashes et al., 2009; S. Lin et al., 2014; Liu et al., 2012; Owald et al., 2015; Pai et al., 2013; Plaçais et al., 2013; Schwaerzel et al., 2003; Séjourné et al., 2011; Trannoy et al., 2011; Yamagata et al., 2015; Zars et al., 2000).

The DANs innervating different MBON compartments appear to play distinct roles in signalling reward versus punishment, novelty versus familiarity, the presence of olfactory cues and the activity state of the fly (Aso et al., 2010; Aso and Rubin, 2016; Burke et al., 2012; Cohn et al., 2015; Hattori et al., 2017; Liu et al., 2012; Sitaraman et al., 2015b; Tsao et al., 2018). These differences between DAN cell types presumably reflect in large part the nature of the inputs that each DAN receives, but our knowledge of these inputs is just emerging (Otto et al., 2020) and is far from comprehensive. DANs adjust synaptic weights between KCs and MBONs with cell type-specific rules and, in at least some cases, these differences arise from the effects of co-transmitters (Aso et al., 2019). In general, a causal association of KC responses with the activation of a DAN in a compartment results in depression of the synapses from the active KCs onto MBONs innervating that compartment (Hige et al., 2015; Handler et al, 2019). Different MB compartments are known to store and update non-redundant information as an animal experiences a series of learning events (Berry et al., 2018; Felsenberg et al., 2017, 2018). In rodent and primate brains, recent studies have revealed that dopaminergic neurons are also molecularly diverse and encode prediction errors and other information based on cell type-specific rules (Hu, 2016; Menegas et al., 2018; Poulin et al., 2020; Watabe-Uchida et al., 2017).

MBONs convey information about learned associations to the rest of the brain. Activation of individual MBONs can cause behavioral attraction or repulsion, according to the compartment in which their dendrites arborize (Aso et al., 2014b; Owald et al., 2015; Perisse et al., 2016). The combined output of multiple MBONs is likely to be integrated in downstream networks, but we do not understand how memories stored in multiple MB compartments alter these integrated signals to guide coherent and appropriate behaviors. Prior anatomical studies implied the existence of multiple layers of interneurons between MBONs and descending motor pathways (Aso et al., 2014a). What is the nature of information processing in those layers? Anatomical studies using light microscopy provided the first hints. MBONs from different compartments send their outputs to the same brain regions, suggesting that they might converge on shared downstream targets. DANs often project to these same brain areas, raising the possibility of direct interaction between MBONs and DANs. The functional significance of such interactions has just begun to be investigated (Felsenberg et al., 2017, 2018; Ichinose et al., 2015; Jacob and Waddell, 2020; Perisse et al., 2016; Zhao et al., 2018b), and studies of the *Drosophila* larva, where a connectome of a numerically less complex MB is available (Eichler et al., 2017), are providing valuable insights (Eschbach et al., 2020a, 2020b; Saumweber et al., 2018).

The recently determined connectome of a portion of an adult female fly brain (hemibrain; see Figure 2-figure supplement 1) provides connectivity data for ∼22,500 neurons (Scheffer et al., 2020). Among them, ∼2,600 neurons have axons or dendrites in the MB, while ∼1,500 neurons are directly downstream of MBONs (using a threshold of 10 synapses from each MBON to each downstream target) and ∼3200 are upstream of MB dopaminergic neurons (using a threshold of 5 synapses from each upstream neuron to each DAN). Thus we will consider approximately one-third of the neurons in the central brain in our analysis of the MB ensemble. There were some limitations resulting from not having a wiring diagram of the full central nervous system, as we lacked complete connectivity information for neurons with processes that extended outside the hemibrain volume (Figure 2-figure supplement 1). We were generally able to mitigate these limitations by identifying the corresponding neurons in other EM or light microscopic data sets when the missing information was important for our analyses. Thus the hemibrain dataset was able to support a nearly comprehensive examination of the full neural network underlying the MB ensemble. Studies of the larval MB are providing parallel information on the structure and function of an MB with most of the same cell types, albeit fewer copies of each (Eichler et al., 2017; Eschbach et al., 2020a, 2020b; Saumweber et al., 2018). The microcircuits inside three MB compartments in the adult were previously described (Takemura et al., 2017) and we report here that the overall organization of these three compartments is conserved in a second individual of a different gender. More importantly, we extend the analysis of microcircuits within the MB lobes to all 15 compartments, revealing additional aspects of spatial organization within individual compartments.

In the current study, we were able to discover new morphological subtypes of KCs and to determine the sensory inputs delivered to the dendrites of each of the ∼2,000 KC. We found considerable structure in the organization of those inputs and unexpectedly high levels of visual input, which was the majority sensory input for two classes of KCs. This segregation of distinct sensory representations into channels is maintained across the MB, such that MBONs, by sampling from different KCs, have access to different sensory modalities and representations.

We discovered a new class of “atypical” MBONs, consisting of 14 cell types, that have part of their dendritic arbors outside the MB lobes, allowing them to integrate input from KCs with other information; at least five of them make strong, direct synaptic contact onto descending neurons that drive motor action. We describe how MBONs from different compartments interact with each other to potentially integrate and transform the signals passed from the MB to the rest of the brain, revealing a number of circuit motifs including multi-layered MBON to MBON feedforward networks and extensive convergence both onto common targets and onto each other through axo-axonal connections. Finally, we analyzed the inputs to all 158 DANs that innervate the MB. We found extensive direct feedback from MBONs to the dendrites of DANs, providing a mechanism of communication within and between MB compartments. We also found groups of DANs that share common inputs, providing mechanistic insights into the distributed parallel processing of aversive and appetitive reinforcement and other experimental observations.

## Results

### An updated MB cell type catalog

The MB can be divided into three distinct parts: the calyx, the pedunculus and the lobes (Figure 2; Video 1). The calyx is the input region for sensory information; KCs have their dendrites in the calyx where they receive inputs from projection neurons. The calyx has subregions: the main calyx (CA) and three accessory calyces. The CA gets over 90 percent of its sensory information from olfactory projection neurons, whereas the smaller accessory calyces are sites of non-olfactory input. The lobes are the main output region of the MB; the axons of the KCs make synapses along their length, as they transverse the lobes, to the dendrites of the MBONs. The pedunculus consists of parallel KC axonal fibers that connect the CA and the lobes and is largely devoid of external innervation in the adult. There are five MB lobes: α, β, α′, β′ and γ. In *Drosophila*, the α and α′ lobes are often called the vertical lobes, and the β, β′ and γ lobes are collectively called the medial, or horizontal, lobes. Each lobe is further divided into compartments by the innervation patterns of DANs and MBONs (Figure 2; Video 2).

We compared the morphology of each EM reconstructed neuron to light microscopy images of genetic driver lines that had been used previously to define the cell types in the MB. Guided by these comparisons, we assigned cell type names that corresponded to established names to the extent possible. In some cases, the availability of full EM reconstructed morphologies allowed us to discern additional subtypes. We also discovered an entirely new class of MBONs, the atypical MBONs, that differed from previously described MBONs in having dendrites both inside and outside the MB lobes. The next few sections describe this updated catalog of MB cell types, including KCs (Figures 3-5; Videos 3 and 6), other MB intrinsic and modulatory neurons (Figure 3-figure supplement 1; Videos 4 and 5), DANs (Figure 6), MBONs (Figure 7) and atypical MBONs (Figure 8; Figure 8-figure supplements 1-14; Videos 7-20). Most MB cell types were named based on light microscopy analyses of their specific innervation areas inside the MB. For instance, MBON-α3 has its dendrites in the third compartment of the α lobe. MBONs and DANs also have synonymous names based on numbering (for example, MBON14 for MBON-α3), which are primarily used in this report; Figure 6-figure supplement 1 shows the neurons contained in each MB compartment and lists their alternative names.

**Figure 3.**
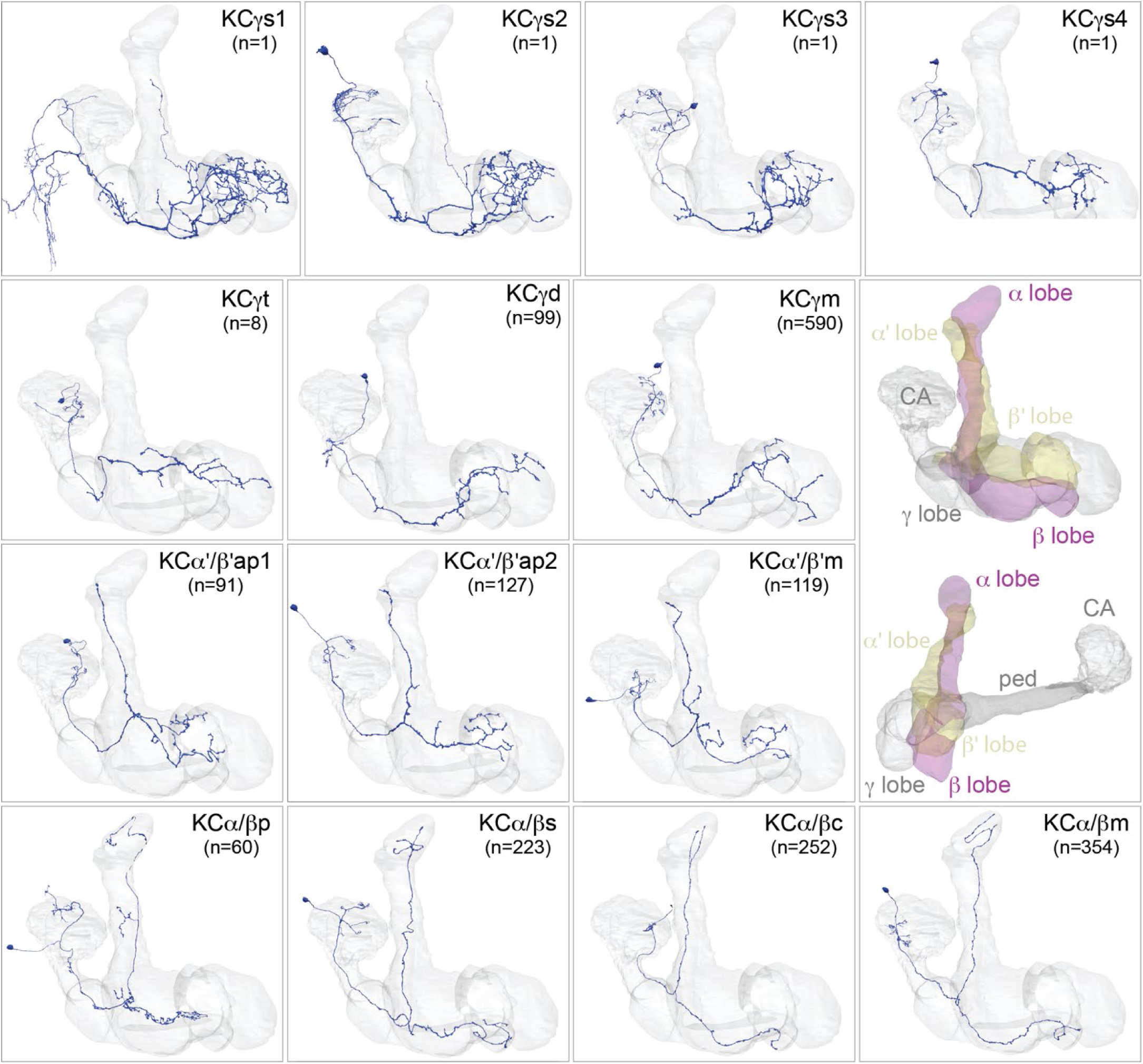
Kenyon cells Each panel shows a representative neuron of the indicated KC subtype together with the outline of the MB lobes and CA in grey, in a perspective view from an oblique angle to better display neuronal morphology. The insert on the right provides a key to the position of the individual lobes, the pedunculus (ped) and CA; the upper image presents the same view as KC subtype panels and the lower image shows a rotated view to better visualize the ped and CA. The numbers (n=) indicate the number of cells that comprise each KC subtype in this animal; the number of KCs is known to vary between animals (reviewed in Aso et al., 2009). Several of the KC subtypes are defined here for the first time, based on morphological clustering, as described in Figure 4. Although reconstruction of 78 KCα/β was incomplete in the CA as a consequence of a small area of reduced image quality, it did not affect morphological clustering, which was based on simplified axonal skeletons. More information about each of these cell types is shown in Video 3. Additional intrinsic and extrinsic neurons with processes in the MB are shown in Figure 3-figure supplement 1.

**Figure 4.**
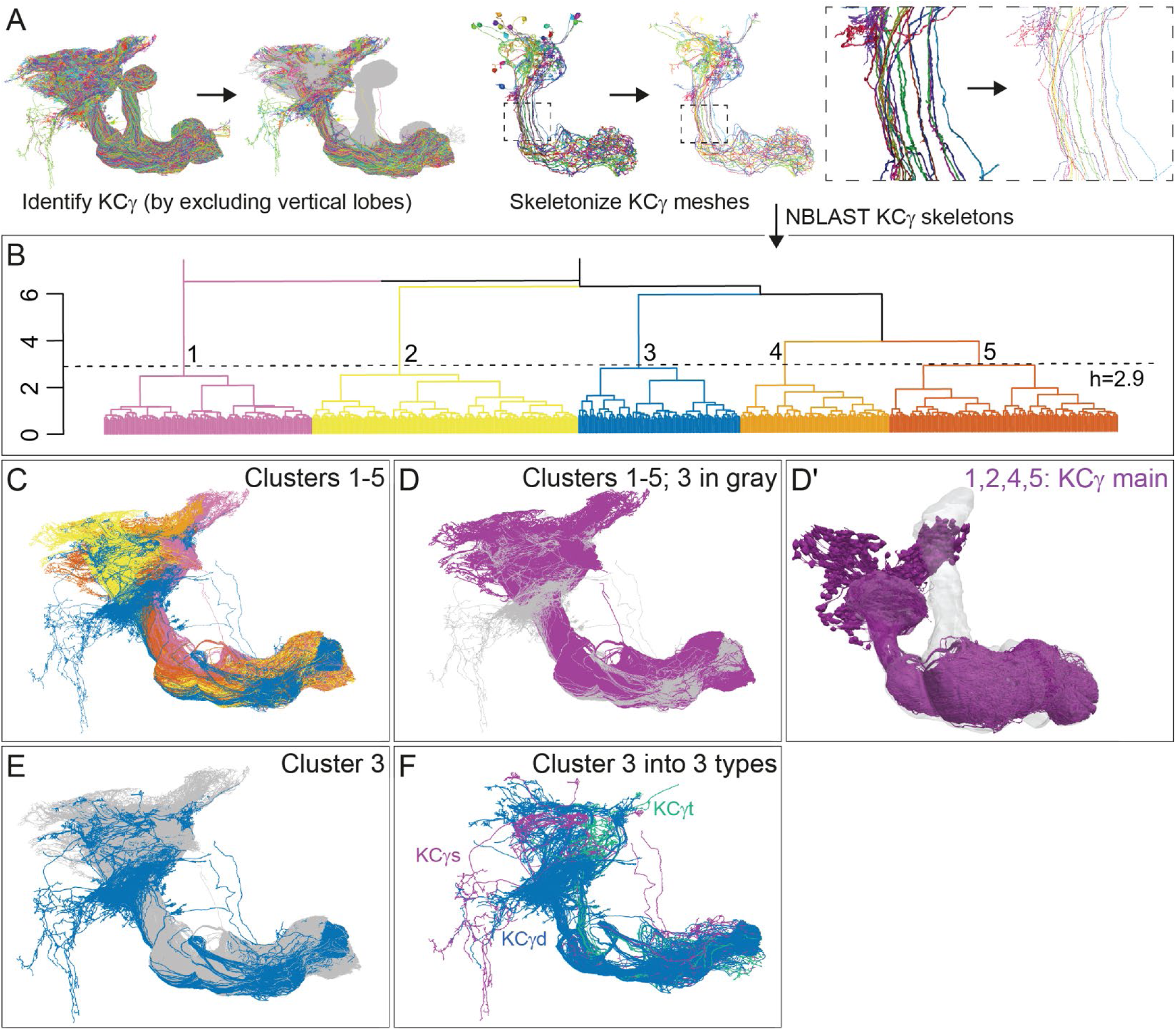
Morphological hierarchical clustering reveals previously unrecognized KC subtypes. (A) KC typing workflow, using KCγ as an example. All γ KCs in the population of annotated KCs were identified by excluding all KCs with axons in the vertical lobes. The space filling morphologies of γ KCs were converted to skeletons (enlarged in the dashed box for clarity) and then NBLAST all-by-all neuron clustering (Costa et al., 2016) was used to reveal morphological groups. (B) Morphological hierarchical clustering of KCγ based on NBLAST scores is shown, cut at height 2.9 (dashed line), which produces 5 clusters. (C) KCγ skeletons of those five clusters are shown, color-coded as in (B). (D) KCγ skeletons in clusters 1, 2, 4 and 5, which includes all 590 KCγm, are shown in magenta; cluster 3 is shown in grey. (D*′*) Space filling morphologies of KCγm (clusters 1, 2, 4 and 5) shown in magenta with the MB in grey. (E) KCγ skeletons in cluster 3 (blue), which includes all KCγd, and clusters 1-2 and 4-5 (grey), which correspond to the KCγm type are shown. (F) KCγ skeletons from cluster 3, cut at height 1.3, which produces 6 sub-clusters corresponding to three color-coded subtypes: green, 3.1 (eight KCγt); magenta, 3.2 (four KCγs); and blue, 3.3-3.6 (99 KCγd); see Figure 4-figure supplement 1 for details).

**Figure 5.**
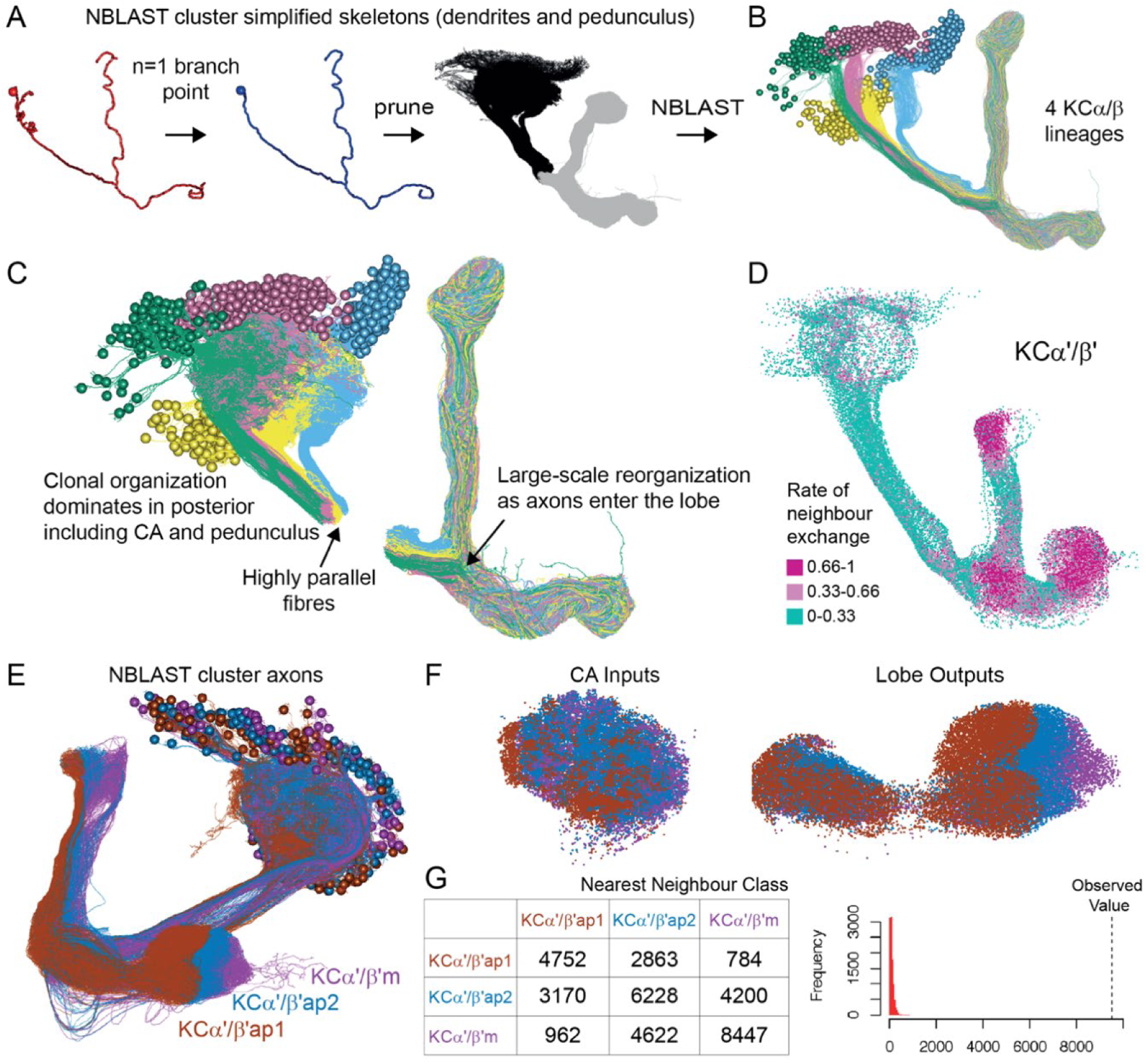
Organizational features of KC projections. (A) KCs were simplified to skeletons with one major branch point which were used as input for NBLAST all-by-all whole neuron clustering. (B) This clustering revealed the four clonal units that make up the mushroom body (shown here for KCα/β). (C) Full neuronal morphologies of the four clusters show that the positions of the neurons in the CA are strongly influenced by this fourfold clonal unit structure, which is also reflected in the arrangement of the highly parallel fibers in the pedunculus. On entering the lobes, the axons reorganize, and the neurons from the four clusters become intermingled. (D) Visualization of the rate of change in KC neighbors quantified as the fraction of 10 nearest neighbors that change compared with a position 5 µm closer to the soma. Large values imply a rapid change in the neighbors of individual KC fibers, which is observed at the entry and tips of the KCα*′*/β*′* lobe as illustrated here. Figure 5-figure supplement 1B shows similar visualizations for the KCγ and KCα/β lobes; rapid change in neighbors is seen throughout the KCγ lobe while KCα/β neurons show an intermediate rate of change. (E) NBLAST clustering of the axons of α*′*/β*′* KCs reveals three clear laminae in the vertical and horizontal lobes, which correlate with a layered organization in the CA and correspond to the three KC α*′*/β*′* subtypes. (F) A similar organization is seen for KCα*′*/β*′* dendrites in the CA and axon outputs in the lobes. (G) As a statistical test for the correlated lobe/CA organization into three subtypes, each synapse in the CA was matched with its closest neighbor from another neuron and the subtype of that neighbor recorded. The contingency table (left) shows that nearest neighbor synapses were most commonly from the same subtype. A permutation test (n=10000) confirmed that this statistic was far higher than expected by chance (right).

**Figure 6.**
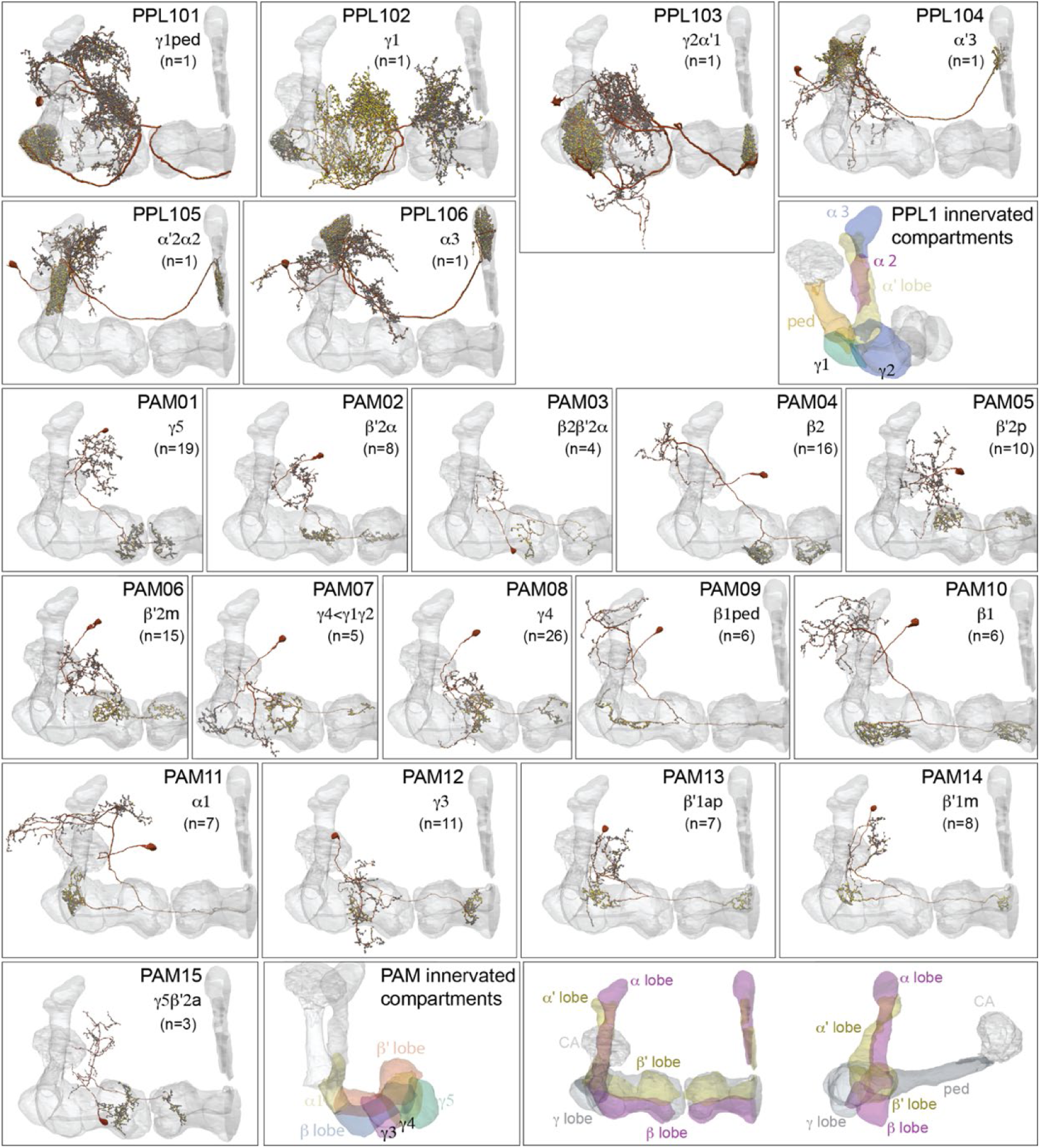
Dopaminergic neurons (DANs). Each panel shows a DAN cell type, with its name, the compartment(s) it innervates and the number of cells of that type per brain hemisphere indicated; the outline of the MB lobes and CA are shown in grey, in a perspective view from an oblique angle to better display neuronal morphology. Figure 6-figure supplement 1 shows which DANs, MBONs and KCs are found in each compartment. PPL1 dopaminergic neurons are divided into six cell types, <u>PPL101</u>, <u>PPL102</u>, <u>PPL103</u>, <u>PPL104</u>, <u>PPL105</u>, and <u>PPL106</u>. As a population, the PPL1 neurons innervate the α’ lobe, α lobe compartments 2 and 3, and γ lobe compartments 1 and 2, as illustrated. There is only one PPL1 DAN of each type per hemisphere, but they send their axons bilaterally to innervate the same MB compartments, although less densely, in the other brain hemisphere (see Aso et al. 2014a). All other compartments are innervated by PAM DANs, as illustrated: <u>PAM01</u>, <u>PAM02</u>, <u>PAM03</u>, <u>PAM04</u>, <u>PAM05</u>, <u>PAM06</u>, <u>PAM07</u>, <u>PAM08</u>, <u>PAM09</u>, <u>PAM10</u>, <u>PAM11</u>, <u>PAM12</u>, <u>PAM13</u>, <u>PAM14</u>, and <u>PAM15</u>. Unlike the PPL1 DANs, multiple PAM DANs of the same cell type innervate the same compartment, and in some cases the same compartment has different PAM DAN types innervating different subdomains of the compartment. MB lobes are shown in grey. A single representative neuron is shown for each cell type in magenta, with grey dots indicating postsynaptic sites and yellow dots indicating presynaptic sites. Images showing the identity of the MB lobes are shown in the lower right.

**Figure 7.**
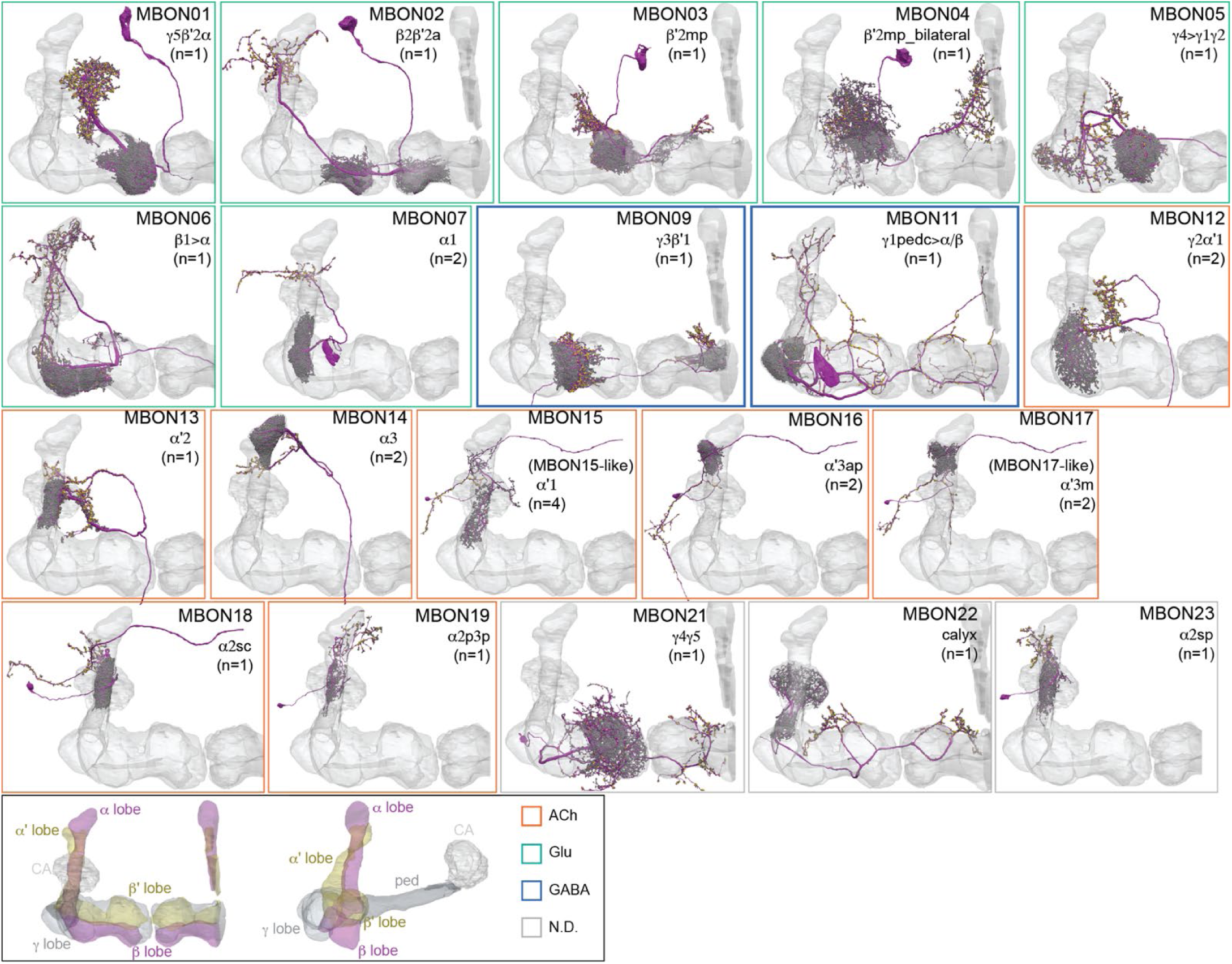
Mushroom Body Output Neurons (MBONs). Each panel shows one of the previously described 20 types of MBONs, with its name, the compartment(s) it innervates and the number of cells of that type per brain hemisphere indicated (Aso et al., 2014a; Takemura et al., 2017); the outline of the MB lobes and CA are shown in grey, in a perspective view from an oblique angle to better display neuronal morphology. MB lobes are shown in grey. A single representative neuron is shown for each cell type (magenta), with grey and yellow dots indicating pre synaptic and presynaptic sites repectively. The bounding box for each neuron is color coded by the neurotransmitter used by that MBON; color key shown in the last panel. These MBONs are considered to be typical in that their dendritic arbors are confined to the MB lobes. We reclassified MBON10 and MBON20 (Aso et al., 2014a) as atypical MBONs since their dendrites extend outside the MB lobes. MBON08, defined by split-GAL4 line MB083C (Aso et al., 2014a), was not found in the hemibrain volume. For the other 21 MBON types, we found only minor differences with previous studies (Aso et al., 2014a; Takemura et al., 2017). For example, MBON15 (α*′*1) and MBON17 (α*′*3m), which each were described as having two cells in (Aso et al., 2014a; Takemura et al., 2017), had additional cells in the hemibrain that were similar in morphology, but had some connectivity differences, that we refer to as MBON15-like and MBON17-like. However, since our observations are based on a single individual, we did not split them into separate cell types. Links to the neuPrint records of these MBON types are as follows: <u>MBON01</u>, <u>MBON02</u>, <u>MBON03</u>, <u>MBON04</u>, <u>MBON05</u>, <u>MBON06</u>, <u>MBON07</u>, <u>MBON09</u>, <u>MBON11</u>, <u>MBON12</u>, <u>MBON13</u>, <u>MBON14</u>, <u>MBON15 (including MBON15-like)</u>, <u>MBON16</u>, <u>MBON17 (including MBON17-like), MBON18, MBON19, MBON20, MBON21, MBON22, and MBON23.</u>

**Figure 8.**
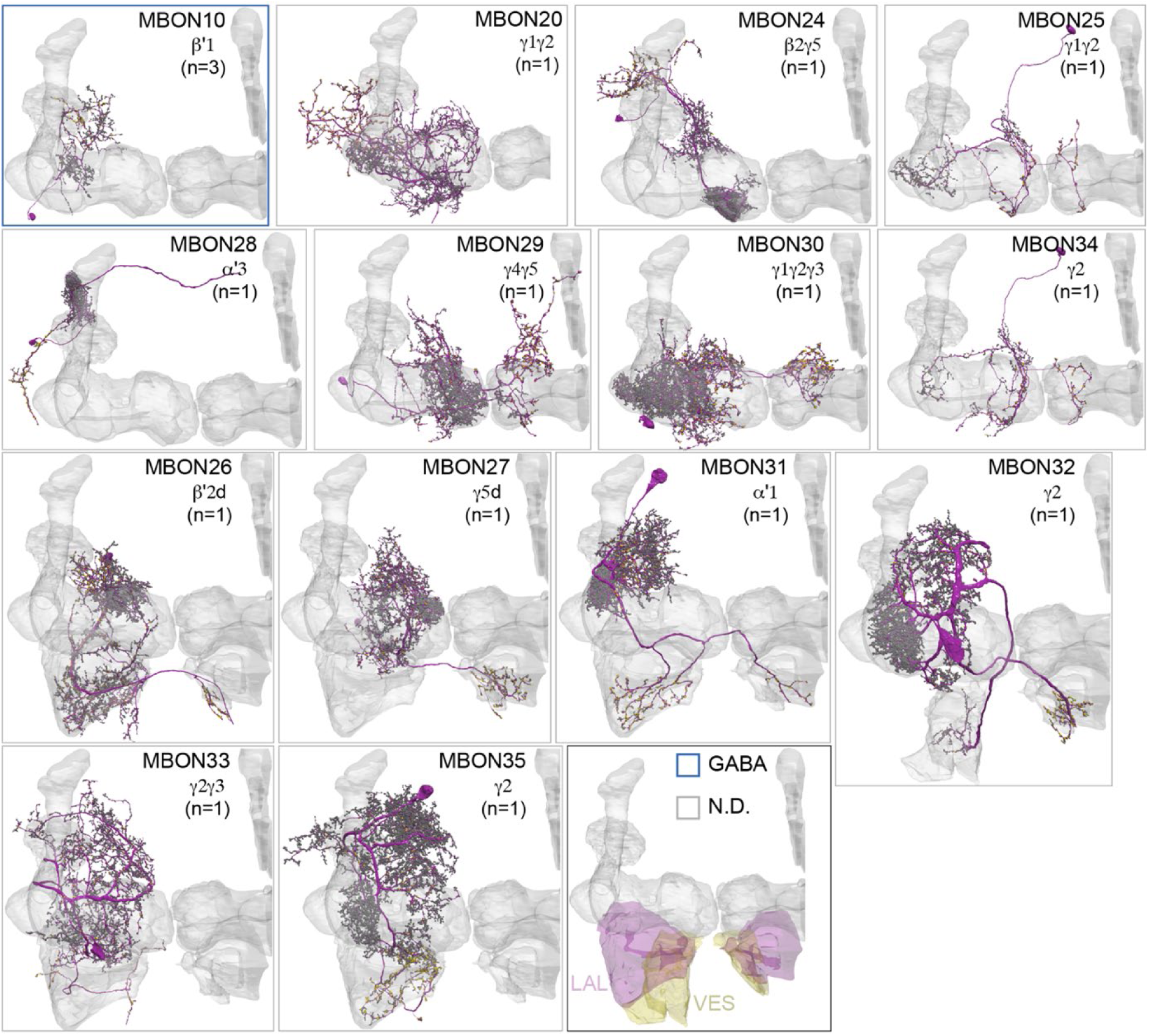
Atypical MBONs. Each panel shows one of the 14 types of atypical MBONs, with its name, the compartment(s) it innervates and the number of cells of that type per brain hemisphere indicated. Figure 6-figure supplement 1 shows which DANs, MBONs and KCs are found in each compartment. MB lobes are shown in grey and in the bottom right panel the lateral accessory lobe (LAL) and vest (VES) brain areas are highlighted. Neuronal morphologies are shown with dark grey dots indicating postsynaptic sites and yellow dots indicating presynaptic sites. The MBONs shown in this figure are considered to be atypical in that their dendritic arbors are only partially within the MB lobes. Twelve of these types were discovered in the course of the current study. The other two, the GABAergic MBON10 and MBON20, were described by Aso et al. (2014a), but we have reclassified them here as atypical MBONs because they have dendrites both inside and outside the MB lobes. The three MBON10s in the EM volume have 69%, 68% and 50% of their postsynaptic sites outside the MB lobes; one is shown here. All the other atypical MBONs occur once per hemisphere. Unlike most typical MBONs that innervate brain areas that are dorsal to the MB, six of the atypical MBONs innervate areas that are ventral to the MB. The LAL is a target of several atypical MBONs, and one also innervates the VES. More detailed information about each of the atypical MBONs can be found in Figure 8-figure supplements 1-14 and Videos 7-20. Figure 8-figure supplement 15 compares the non-MB inputs to these MBONs. Neurotransmitters of atypical MBONs are unknown except MBON10, as indicated by the color of the bounding box. Links to the neuPrint records of these MBON types are as follows: <u>MBON10</u>, <u>MBON24</u>, <u>MBON25</u>, <u>MBON26</u>, <u>MBON27</u>, <u>MBON28</u>, <u>MBON29</u>, <u>MBON30</u>, <u>MBON31</u>, <u>MBON32</u>, <u>MBON33</u>, <u>MBON34</u>, and <u>MBON35</u>.

### KCs: the major MB intrinsic neurons and conveyors of sensory identity

Associative memories in the MB are stored as altered synaptic weights between KCs, which represent sensory information, and their target MBONs (Bouzaiane et al., 2015; Cassenaer and Laurent, 2012; Hige et al., 2015; Owald et al., 2015; Pai et al., 2013; Perisse et al., 2016; Sejourne et al., 2011). Each of the 15 MB compartments is unique in cellular composition, and individual compartments can exhibit internal substructure, which we discuss later in the paper. In this next section we consider the different types of KCs that project to each compartment. In later sections, we examine how the various KC types receive distinct sensory information from projection neurons in the calyces and then connect differentially with MBONs to provide each MBON cell type with access to a different sensory space to use in forming memories. The number of KCs connected with each MBON also varies significantly from 122 (MBON10) to 1694 (MBON11), which can influence memory capacity and specificity.

We identified 1927 KCs in the right brain hemisphere. There are three major KC classes: α/β, α′/β′ and γ KCs. KCs are sequentially generated from four neuroblasts in the order of γ, α′/β′ and α/β (Video 6; Crittenden et al., 1998; Ito et al., 1997; Lee et al., 1999; Zhu et al., 2003). The axons of α/β and α′/β′ KCs bifurcate at the distal pedunculus to innervate the α and β lobes or the α′ and β′ lobes, respectively. The axons of γ KCs also branch but are confined to the γ lobe. Genetic driver lines, immunohistochemistry and single cell morphology has revealed further subtypes (Aso et al., 2009; Lin et al., 2007; Strausfeld et al., 2003; Tanaka et al., 2008). Here we grouped 1927 KCs into 14 subtypes (Figure 3; Video 3) by applying NBLAST morphological clustering (Bates et al., 2020a; Costa et al., 2016) to each major class of KCs (Figure 4). Despite the dominance of olfactory input to the MB, all three major classes of KCs were found to contain small subsets dedicated to non-olfactory information.

γ KCs: KCs that innervate the γ lobe (KCγ) have been traditionally divided into two groups, dorsal (KCγd) and main (KCγm). Axons of γd KCs innervate the dorsal layer of the γ lobe and γm KCs innervate the rest of the γ lobe. The dendrites of γd KCs arborize in the ventral accessory calyx (vACA), whereas those of γm KCs are found in the CA (Aso et al., 2014a, 2009; Vogt et al., 2016). We identified the 701 γ KCs by excluding any KCs that innervated the vertical lobes (Figure 4). We then converted their 3D morphologies (meshes) into skeletons and performed an all-by-all NBLAST, which allowed us to define new KCγ subtypes and enumerate the members of each type: 590 KCγm, 99 KCγd, eight KCγt neurons with dendrites in the anterior CA (targeted preferentially by thermo/hygrosensory neurons), and four unique KCγs neurons sampling from one or more accessory calyces (Figure 4-figure supplement 1). All NBLAST clusters were validated by an independent clustering based on connectivity, CBLAST (Scheffer et al., 2020), and the small number of discrepancies were resolved by manual inspection (see Materials and methods).

α′/β′ KCs: We identified 337 α′/β′ KCs, which could be divided into three subtypes using all-by-all NBLAST on their axons (Figure 4-figure supplement 2). The axons of each subtype formed a distinct layer in both the vertical and horizontal lobes. We named these subtypes to be consistent with prior nomenclature based on split-GAL4 driver lines (Figure 4-figure supplement 3). There are 91 α′/β′ap1 (Figure 4-figure supplement 2 B, B′), 127 α′/β′ap2 (Figure 4-figure supplement 2 D, D′), and 119 α′/β′m (Figure 4-figure supplement 2 C, C′) KCs. Moreover, we found that each subtype’s axon layer was correlated with the position of its dendrites in the CA (Figure 5E). The dendrites of the α′/β′ap1 KCs lie in the lateral accessory calyx and anterior CA, areas that are preferentially targeted by thermo/hygrosensory sensory projection neurons (Figure 12B).

α/β KCs: We identified 889 α/β KCs, which could be divided into four subtypes using all-by-all NBLAST on their axons (Figure 4-figure supplement 4). The first subtype corresponds to the 60 KCα/βp that form the posterior layer of the α and β lobes (Figure 4-figure supplement 4 C, C’); these are the first α/β KCs to be born and have been referred to as pioneer KCα/βs for this reason (Lin et al., 2007; Zhu et al., 2003). The remaining three groups form concentric layers (surface, middle, and core) in both the α and β lobes, yielding 223 KCα/βs (Figure 4-figure supplement 4 D, D′), 354 KCα/βm (Figure 4-figure supplement 4 F, F′), and 252 KCα/βc (Figure 4-figure supplement 4 E, E′). Dendrites of α/βp KCs form the dorsal accessory calyx (dACA), while the rest of α/β KCs have dendrites in the CA. Our classification of α/β KC subtypes is consistent with prior light level studies (Tanaka et al., 2008; Zhu et al., 2003).

Each of the four neuroblasts whose progeny form the MB lobes is thought to generate all classes of KCs, but their exact contributions have been difficult to assess. There is no labelling of neuroblast origin in EM images, but neurons derived from the same neuroblast tend to have cell bodies in close proximity and primary neurites bundled into the same fiber tract. Tight groupings of cell bodies are particularly evident for α/β KCs. To classify α/β KCs into four neuroblast groups, we applied NBLAST to simplified skeletons of α/β KCs whose axons in the lobes had been removed (Figure 5A). As expected, we found four equal-sized groups of α/β KCs that we believe are each the descendants of a single neuroblast (Figure 5B). These four groups form subregions in the CA and pedunculus, but their axons are scrambled in the lobes (Figure 5C) as previously demonstrated by genetic methods (Lin et al., 2007; Zhu et al., 2003).

Upon entering the MB lobes, the axons of each KC type project to spatially segregated layers in the lobes (Video 6), with the exception of γm KCs axons which meander along the length of the horizontal lobe (Figure 5-figure supplement 1B). This maintained segregation is most prominent for α′/β′ KCs but is also seen in α/β KCs (Figure 5E-G). The dendrites of each KC type also tend to be found in the same region of the CA (Video 6; Leiss et al., 2009; Lin et al., 2007; Zheng et al., 2020), which, in some cases, appears to support input specialization. These features of the spatial mapping from CA to lobes and the organisation of the parallel fiber system presumably evolved to facilitate associative learning. This spatial arrangement gives each MBON the possibility of receiving mixed or segregated sensory information depending on where that MBON extends its dendrite within the different KC layers, and, similarly, gives DANs the potential ability to modify strengths of synapses from KCs conveying specific sensory information. KC make synapses to neighboring KCs in the calyx, pedunculus and lobes. These were described for the α lobe in Takemura et al., (2017), and Figure 5-figure supplement 2 provides a summary for the entire MB.

### DANs: the providers of localized neuromodulation

For associative learning to occur, the neuronal pathways that convey punishment or reward must converge with those that convey sensory cues. In the fly brain, this anatomical convergence takes place in the compartments of MB lobes: sensory cues are represented by the sparse activity of KCs and reinforcement signals by the DANs that innervate the MB lobes. DANs have been traditionally grouped into two clusters, PPL1 and PAM, based on the position of their cell bodies (Figure 6). PPL1 DANs innervate the vertical lobes and generally convey punishment, whereas PAM DANs innervate horizontal lobes and generally convey reward (Aso et al., 2019, 2012, 2010; Aso and Rubin, 2016; Burke et al., 2012; Claridge-Chang et al., 2009; Felsenberg et al., 2018, 2017; Huetteroth et al., 2015; Jacob and Waddell, 2020; Lin et al., 2014b; Liu et al., 2012; Mao and Davis, 2009; Schwaerzel et al., 2003). There is also a DAN from the PPL2ab cluster, PPL201 (Figure 3-figure supplement 1G), that innervates the CA (Mao and Davis, 2009; Tanaka et al., 2008; Zheng et al., 2018) and has been reported to play a role in signalling saliency (Boto et al., 2019). We defined six PPL1 DAN cell types (PPL101-PPL106; see Video 2 for PPL106) and 15 PAM DAN cell types (PAM01-PAM15). These cell type classifications are consistent with previous studies (Aso et al., 2014a), except for the addition of one new type, PAM15 (γ5β′2a). There is only a single cell per PPL1-DAN cell type in a hemisphere, and axons of each cell broadly arborize in the compartment(s) they innervate. In contrast, there are between 3 and 26 cells per PAM DAN cell type, and the axonal terminals of an individual PAM DAN occupy only a portion of the compartment it innervates (see Video 32 for PAM11 and Video 33 and Figure 32 for PAM12). Thus, it is possible to further subdivide the members of PAM DAN cell types in Figure 6 into smaller groups based on morphology and connectivity as described in Otto et al. (2020). We present an extensive analysis of such subtypes later in the paper (Figures 28-37).

### MBONs: the MB’s conduit to the brain for learned associations

The representations of sensory cues and memory traces encoded in KC axon terminals have been reported to be read out by a network of 22 types of MBONs (Aso et al., 2014a; Takemura et al., 2017). We found all the previously described MBON types in the hemibrain volume (Figure 7), except for MBON08 which is not present in the imaged fly. MBONs can be categorized into three groups by their transmitters, which also correspond to anatomical and functional groups. Dendrites of glutamatergic MBONs arborize in the medial compartments of the horizontal lobes, which are also innervated by reward-representing PAM-DANs. Most cholinergic MBONs arborize in the vertical lobes, in compartments that are also innervated by punishment-representing PPL1-DANs. GABAergic MBONs also arborize in compartments innervated by punishment-representing DANs. As described above, axon fibers of distinct types of olfactory and non-olfactory KCs form layers in the MB lobes. Each MBON arborizes its dendrites in a subset of layers where they receive excitatory, cholinergic synapses from KCs (Barnstedt et al., 2016; Takemura et al., 2017). These KC synapses are known to be presynaptically modulated by dopamine (Davis, 2005; Hige et al., 2015; Kim et al., 2007). Within the MB lobes, MBONs also receive input from APL (Liu and Davis, 2009) and DPM (Waddell et al., 2000) as well as from DANs (Takemura et al., 2017).

MBONs generally project their axons outside the MB lobes, with the exception of three feedforward MBONs that project to other MB compartments (Aso et al., 2014a). As discussed in detail below (Figures 18-25, Videos 23-25), MBONs most heavily innervate dorsal brain areas such as the CRE, SIP and SMP (Figure 18), make direct connections to the fan-shaped-body of the central complex (Figures 19 and 20), tend to converge on common targets (Figure 21), form a multi-layer feedforward network employing axo-axonal synapses (Figure 24), and provide input to the dendrites of DANs (Figure 26).

### Atypical MBONs: integrators of information from inside and outside the MB lobes

We identified 14 additional types of MBONs that differ from MBONs previously described in the adult. We refer to these cell types as ‘atypical MBONs’ in that their dendritic arbors, in addition to having extensive KC input within the MB lobes, extend outside the MB lobes into adjacent brain areas (Figure 8, Figure 8-figure supplements 1-14, Videos 7-20). We reclassified MBON10 and MBON20 as atypical MBONs since these two cell types had significant dendritic arbors outside the MB lobes (Figure 8, Figure 8-figure supplements 1 and 2, Videos 7 and 8). Twelve of the 14 atypical MBON types innervate the horizontal lobes. Unlike typical MBONs, six of the atypical MBONs have significant innervation in ventral neuropils, in particular the LAL.

For each of the atypical MBONs we provide a figure supplement (Figure 8-figure supplements 1-14) that provides information on its top inputs and outputs as well as a video (Videos 7-20) that displays additional morphological and connectivity features.

As these MBONs are described here for the first time, no experimental data yet exists on their function(s) or physiology. Nevertheless, their connectivity provides clues. Each atypical MBON is poised to integrate information conveyed by KCs with additional inputs to the portion of its dendritic arbor that lies outside the MB lobes. Figure 8-figure supplement 15 shows which brain regions supply input to each of the atypical MBONs. Frequently, these inputs include the outputs of other MBONs; nine of the 14 atypical MBONs have at least two other typical or atypical MBONs among the top 10 inputs to their dendrites that lie outside the lobes. Some atypical MBONs receive sensory information directly. MBON28 (α′3) receives strong multiglomerular PN input, with three mPNs among its top 10 inputs outside the MB. MBON24 (β2γ5)’s top two inputs outside the MB are putative suboesophageal zone output neurons (SEZONs), likely to convey mechanosensory or gustatory information based on their arbors traced in the FAFB volume (Otto et al., 2020; Zheng et al., 2017) that contains the full SEZ. One atypical MBON, MBON30 (γ1γ2γ3), is the only MBON that receives significant input directly from the central complex; the nine cells of one fan-shaped body (FB) columnar cell type, FB2-5RUB, converge in a small brain area called the rubus where they make nearly 500 synapses onto MBON30 (Video 15).

Such features suggest that the atypical MBONs might be involved in information convergence. Six of the atypical MBONs project to the LAL, positioning them to connect more directly with the motor network than typical MBONs which send their outputs to dorsal brain areas, a feature we explore in detail later in the paper.

### Sensory inputs to the KCs: the calyces

Sensory information is conveyed to the dendrites of KCs in specialized MB structures called calyces. The main calyx (CA) contains the dendrites of 90% percent of the KCs; the dendrites of the remaining KCs are in one of three accessory calyces, each with a specialized function and distinct KC composition. Our work confirms and extends prior descriptions of the calyces (Aso et al., 2009; Bates et al., 2020b; Butcher et al., 2012; Marin et al., 2020; Tanaka et al., 2008).

Main calyx (CA): Olfactory sensory neurons (ORNs) that express the same odorant receptor project their axons to the same glomerulus in the antennal lobe. Uniglomerular PNs (uPNs) arborize their dendrites in a single glomerulus and thus receive direct sensory signals from one type of ORN. These olfactory uPNs are the major sensory inputs to the main calyx (Figure 9A). PNs branch when they enter the CA and their axons terminate in an average of ∼6 synaptic boutons, although the number varies between PN cell types (Figure 9-figure supplement 1) and across individuals (Gao et al., 2019). The number of boutons per KC also differs between KC cell types in the CA: KCα′/β′, 4.40 +/- 1.58; KCα/β, 4.67 +/- 1.78; KCγ, 8.49 +/- 2.17. The synaptic boutons from a single PN can be spread over a large fraction of the calyx. Thermo- and hygrosensory uPNs and multiglomerular PNs (mPNs) can also terminate in the CA, mainly in the anterior (Figure 9B). PNs that convey particular odor scenes (for example decaying fruit or pheromones) appear to target specific areas of the CA (Figure 9C; Figure 9-figure supplement 2). We identified a subset of γm KCs that, while receiving input from olfactory PNs in CA, also extended dendritic claws outside the main calyx to receive gustatory input from a single PN delivering both hygrosensory and gustatory information, providing the first description of a pathway for gustatory information to reach KCs (Figure 9-figure supplement 3).

**Figure 9.**
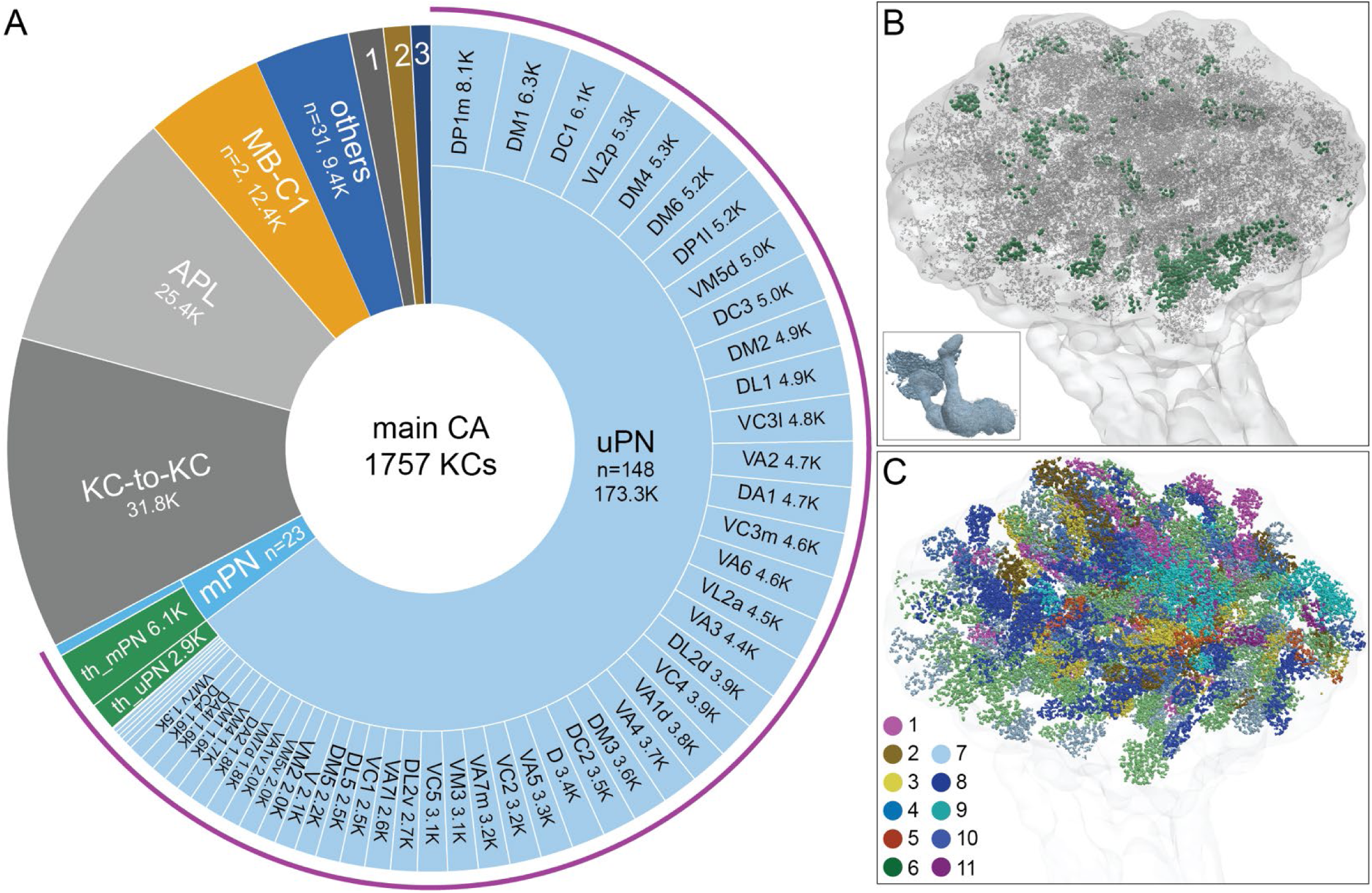
Main calyx (CA). The dendrites of 1757 KCs of the α/β, α′/β′ and γ cell types define the CA. (A) The pie chart shows a breakdown of the inputs to these KCs. The largest source of input is from 129 uniglomerular olfactory projection neurons as judged by synapse number (uPNs; 63.6% of total input to the KCs); the number of synapses is indicated for each uPN cell type. Additional olfactory sensory input is provided by 23 multiglomerular projection neurons (mPNs). Information about temperature is provided by both 35 mPNs (th_mPN) and 19 uPNs (th_uPN). The next most prominent inputs are KC-to-KC synapses within the CA (11.9%), from APL (9.5%; see Figure 3-figure supplement 1A) and from MB-C1 (4.6%; see Figure 3-figure supplement 1H). Smaller sources of input are indicated by the numbered sectors: 1, a group of 9 neurons previously described as “centrifugal” neurons (Bates et al., 2020) that innervate both CA and LH (1.3%). 2, <u>MB-CP2</u> (1.0%); 3, <u>PPL201</u> (see Figure 3-figure supplement 1G). The remaining 3.4% is provided by 31 other neurons (blue). (B) An image of the CA showing the locations of olfactory PN and thermo PN synapses onto KCs. The green dots representing thermo PN olfactory input synapses are of larger diameter to allow better visibility in the presence of the larger number of grey dots representing olfactory input synapses. Note the thermo PN inputs are located in the anterior and at the periphery of the CA, corresponding to the position of α′/β′ap1 and γt KC dendrites. The inset shows the orientation of the image. (C) Inputs from olfactory PNs are shown color-coded based on the type of olfactory information they are thought to convey (see Bates et al., 2020): 1, fruity; 2, plant matter; 3, animal matter; 4, wasp pheromone; 5, insect alarm pheromone; 6, yeasty; 7, alcoholic fermentation; 8, decaying fruit; 9, pheromonal; 10, egg-laying related. 11. geosmin.

Ventral accessory calyx: The dendrites of 99 γd KCs surround the base of the CA in a loose ring and form the ventral accessory calyx (vACA; Figure 10). The accessory calyces are thought to be specialized for non-olfactory information, and indeed we found that visual projection neurons (VPNs) from the medulla (ME) and the lobula (LO) are the predominant inputs to the γd KCs. While the LO is largely contained in the hemibrain, the ME is not. In many cases, we were able to confirm predicted ME VPNs by matching neuronal fragments with their complete counterparts in FAFB or light-microscopy images for conclusive annotation of ME VPNs (Figure 10-figure supplement 1).

**Figure 10.**
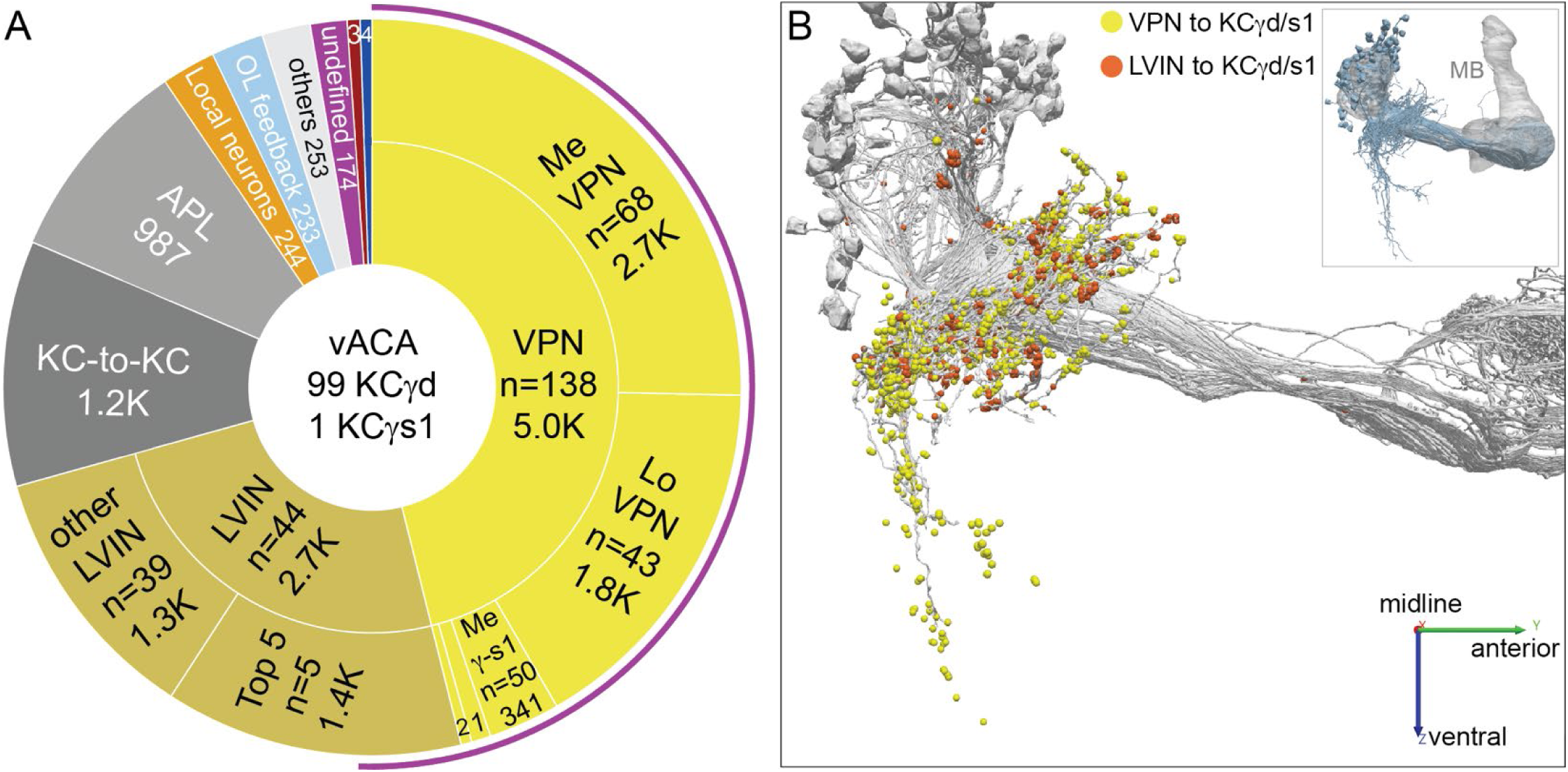
Ventral accessory calyx (vACA). The dendrites of the 99 γd and γs1 KCs define the vACA. (A) The pie chart shows a breakdown of the inputs to these KCs; the number of cell (n=) and the number of the total synapses contributed by the cells in that sector are shown without applying a threshold. The majority of inputs convey visual information, either directly from visual projection neurons (VPNs; 46.1%) or through intermediate local visual interneurons (LVIN; 24.6%) that themselves receive input from VPNs. The number of VPNs shown in pie chart counts VPNs that make as few as one synapse. When a threshold is applied that requires a VPN to make at least 5 synapses to a single KC, then we find 49 VPNs, including 26 ME VPNs and 21 LO VPNs. The synapses from the VPNs and the LVINs onto the KCγd dendrites do not show the claw-like structure seen in the CA (Video 22). A ranking of LVINs based on the amount of visual input conveyed is shown in Figure 10-figure supplement 1A. More than half of the indirect input is mediated by five LVINs (Top 5), which are shown in Figure 10-figure supplement 3B. VPNs can be subdivided based on the location of their dendrites in either the medulla (ME) or lobula (LO), as indicated in the outer circle. There are 68 VPNs that connect to the single KCγs1, with a total of 483 synapses: 50 from the Me, 34 of which are shared with other γd KCs, and 14 from the LO, eight of which are shared with other γd KCs (represented by the numbered sector 1). The next most prominent inputs to KCs in the vACA are synapses between the KCs themselves (10.8%), from APL (9.1%), from local interneurons that do not appear to convey significant visual information (2.3%), from interneurons that send feedback from the vACA to optic lobe neurons (OL feedback; 2.2%) and neurons that leave the volume with undefined identity (undefined; 1.6%). Other sources of input are indicated by the other numbered sectors: 2, other VPN input that we could not classify as from the Me or Lo, due to incomplete morphology (0.9%); 3, <u>putative mPNs</u> (0.6%) (5813063239, 1442819296, 5813040515); 4, a <u>putative SEZ cell type</u> (0.5%). The remaining 2.3% is provided by 253 interneurons that are weakly connected to these KCs, with each providing 1 synapse to each of less than 4 KCs (others). The fraction of input to the vACA KCs conveying visual information is indicated by the outer purple arc; it reflects the direct input from the VPNs plus the fraction of the LVIN input that represents visual input. (B) Synaptic connections from visual projection neurons (VPN) and local visual interneurons (LVIN) onto γd and ©s1 KCs (grey), color-coded. Note the different spatial distribution of synapses from VPNs and LVINs. VPNs make synapses onto KCγd dendrites in an area ventral to the CA, previously recognized as the vACA (Butcher et al., 2012), as well as in a diffuse ring surrounding the base of the CA; synapses from LVINs are restricted to the ring. Additional views are shown in Figure 10-figure supplement 4A and B.

Although the γd KCs respond to light and are required for learning the predictive value of color (Vogt et al., 2016, 2014), we have little definitive insight into the type of visual information conveyed by their VPN inputs. In bees, ME and LO VPNs convey specific chromatic, temporal, and motion features, including sensory information required for associations bees make during foraging tasks (Paulk and Gronenberg, 2008). The largest group of VPNs in *Drosophila* are ipsilateral, unilateral ME neurons. The ME VPNs have dendrites in the outer part of the ME (up to layer M8); the LO VPNs primarily arborize in the deeper layers of the LO (Lo4-Lo6). These layers patterns are consistent with a possible role in conveying information about color and intensity but notably exclude optic lobe regions that are strongly associated with motion vision such as the lobula plate, ME layer M10 and LO layer Lo1 (Borst, 2014). The LO VPNs are also clearly distinct from the well-studied lobula columnar cells with response to visual features such as visual looming or small moving objects (Wu et al., 2016). The optic lobe has a retinotopic organization. Several ME and LO VPNs have arbors that are restricted to parts of the ME and LO and thus are predicted to preferentially respond to stimuli in different parts of the visual field. However, we did not observe evidence for a high-resolution spatial map formed by KC inputs. Local visual interneurons (LVINs) that do not themselves arborize in the optic lobe but are downstream of VPNs convey additional visual information (Figure 10A). The connections from VPNs and LVINs onto KCγd dendrites are spatially segregated (Figure 10B). LVINs are discussed in more detail below.

Dorsal accessory calyx: The dendrites of 60 α/βp KCs define the dorsal accessory calyx (dACA) and receive predominantly visual input (Figure 11A). We found that the VPNs that directly connect to α/βp KCs come mostly from the ME (Figure 11A), with a much smaller contribution from the LO; VPNs projecting from the LO to the dACA have also been recently noted by Li et al., (2020). However, unlike in the vACA, indirect visual input conveyed by LVINs outweighs direct VPN input (Figure 11A), and VPN and LVIN inputs are less segregated (Figure 11B). Among these LVINs, a cluster of 13 morphologically similar neurons (SLP360, SLP362 and SLP371) contributes over 50% of input from all LVINs. One LVIN, MB-CP2, which has been suggested to integrate multi-sensory inputs (Zheng et al., 2018), is the single strongest dACA input neuron (Figure 11-figure supplement 2E) and also seems to relay input from the subesophageal zone (SEZ).

**Figure 11.**
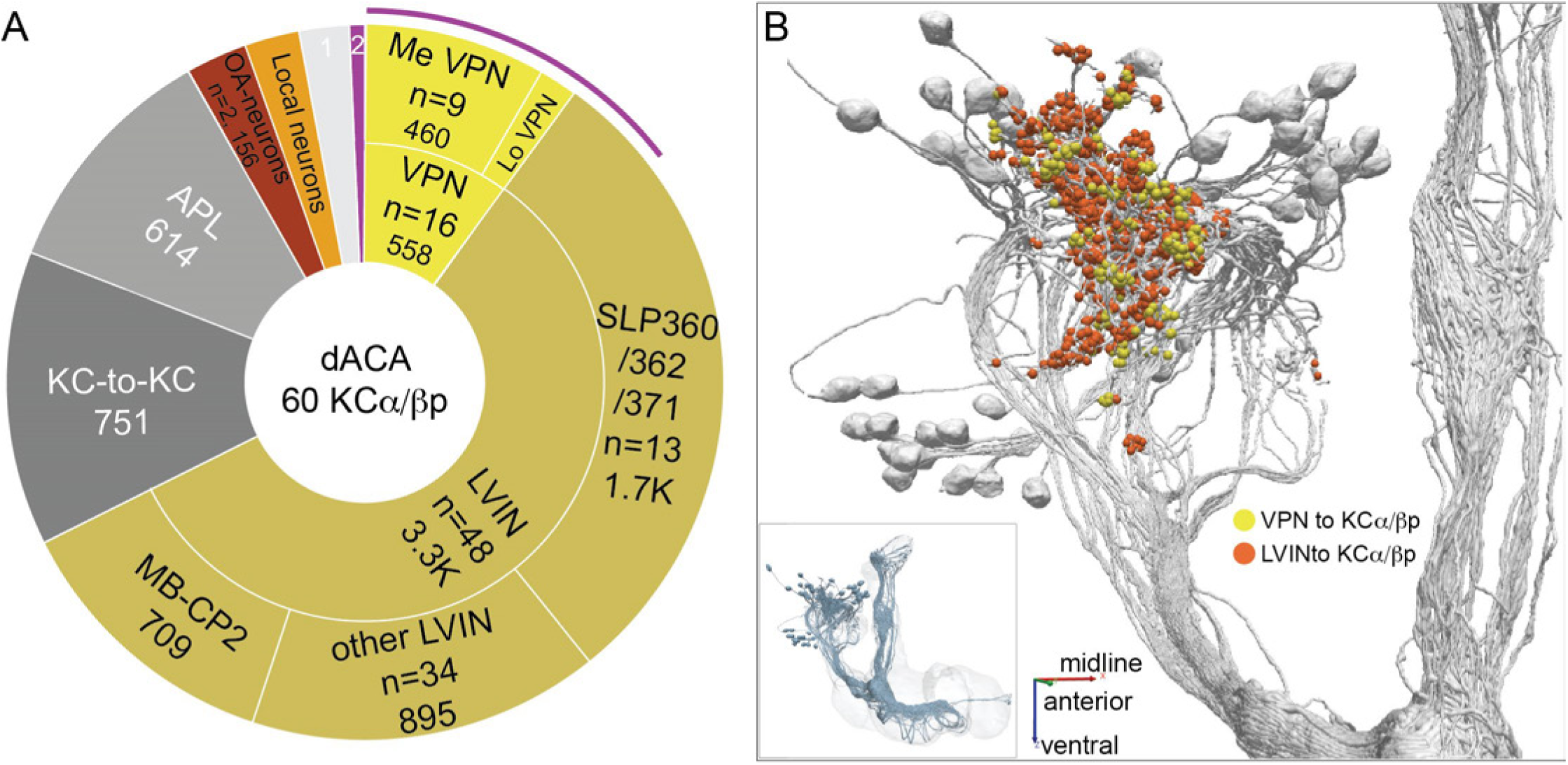
Dorsal accessory calyx (dACA). The dendrites of the 60 α/βp KCs define the dACA. The pie chart shows a breakdown of the inputs to these KCs. The majority convey visual information, either directly from visual projection neurons (VPN; 9.8%) or through intermediate local visual interneurons (LVIN; 57.8%) that receive input from VPNs (see Figure 11-figure supplement 1). VPNs can be subdivided based on the location of their dendrites in either ME or LO, as indicated in the outer circle. More than two-thirds of the indirect input is mediated by the LVIN cell types <u>SLP360</u>, <u>SLP362</u>, and <u>SLP371</u>, shown in Figure 11-figure supplement 1C; this SLP360/361/371 cluster of 13 neurons contributes about 30% of total input to the 60 α/βp KCs in the dACA. Neurons of similar morphology have also been observed to be presynaptic to KCα/βp in the dACA in a recent study (Li et al., 2020). Another LVIN, <u>MB-CP2</u> (LHPV3c1) (479935033), provides 12.6% of the input to KCs in the dACA; however, only a small percentage of its inputs are visual (see Figure 11-figure supplement 1A and E). The total visual information presented to KCs by VPNs and LVINs is indicated by the purple arc around the outer layer; it reflects the direct input from the VPNs plus the fraction of the LVIN input that represents visual input. The next most prominent inputs are KC-to-KC synapses in the dACA (13.3%), from APL (10.9%), from two octopaminergic neurons (2.8%; <u>OA-VPM3</u>, see Figure 3-figure supplement 1D, and <u>OA-VUMa2</u>, see Figure 3-figure supplement 1F); and <u>local interneurons</u> (n=22; 2.3%). Remaining input, “others”, are input from 102 different neurons that are all weakly connected; and numbered sector 1 are mPNs (0.7%). The dendrites of KCα/βp neurons in the dACA (Tanaka et al., 2008; Zhu et al., 2003) are reportedly activated by bitter or sweet tastants (Kirkhart and Scott, 2015). However, the KCα/βp are not required for taste conditioning, which instead appears to depend on γ KCs (Kirkhart and Scott, 2015) and we were unable to identify strong candidates for delivering gustatory sensory information to the dACA. The PN VP5+Z adPN (<u>5813063239</u>) connects to two α/βp KCs has dendrites in the SEZ (Figure 9-figure supplement 3). But this is the only gustatory PN we can associate with the dACA, and it primarily projects to KCγm neurons through which it might participate in conditioned taste aversion (Kirkhart and Scott, 2015). (B) Color-coded synaptic connections from visual projection neurons (VPN; yellow) and local visual interneurons (LVIN; orange) onto α/βp KCs (grey). Note that, unlike in the vACA, there are more connections from LVINs than VPNs in the dACA.

As described above, local visual interneurons (LVINs) make up a substantial portion of the inputs to γd KCs (Figure 10-figure supplement 2) and α/βp KCs (Figure 11-figure supplement 1) in the vACA and dACA, respectively. LVINs get input from multiple VPNs, as well as nonvisual inputs, and then convey this integrated information to KCs. The neuronal morphologies and connectivity patterns of the most strongly connected LVINs are shown in Figure 10-figure supplements 3-5 for the vACA and Figure 11-figure supplements 2 and 3 for the dACA. We observed that clusters of LVINs, or sometimes single LVINs, receive input from distinct subpopulations of VPNs. Moreover, some of the LVINs that receive inputs from similar VPN subpopulations tend also to receive similar nonvisual inputs (Figure 11-figure supplements 4 and 5). These observations suggest that, rather than simply relaying visual information, LVINs may perform more complex processing, including integration of visual and nonvisual signals.

Lateral accessory calyx: The lateral accessory calyx (lACA) is a small subcompartment of the CA innervated by 14 α′/β′ap1 KCs (Yagi et al., 2016) and one KCγs2 (Figure 12). The lACA, which has been recently described in detail in Marin et al. (2020), is thought to be a thermosensory center as >90% of its input comes from two PNs: the slow-adapting, cooling air-responsive PN, VP3+ vPN (Liu et al., 2015; Stocker et al., 1990), which solely targets the lACA (Jenett et al., 2012), and the predicted warming air-responsive VP2 adPN (Marin et al., 2020). Other inputs to the lACA are described in Figure 12 and the inputs to KCγs2 are separately detailed in Figure 12-figure supplement 2. Most KCs in the lACA also receive inputs in the CA from olfactory and thermo/hygrosensory PNs. There are direct connections to DN1a clock neurons within the lACA (Yagi et al., 2016; Marin et al., 2020), as well as from the DN1a neurons and the aMe23/DN1-like neuron to the 5^th^ s-LNv and one LNd (Figure 12-figure supplement 1), which could play a role in entrainment of the circadian clock by temperature (Figure 12-figure supplement 2) or the adjustment of sleep patterns to different temperatures (Yadlapalli et al., 2018; Alpert et al., 2020). These connections appear to reflect a function of the lACA that is distinct from its role as a site of thermosensory inputs to KCs.

**Figure 12.**
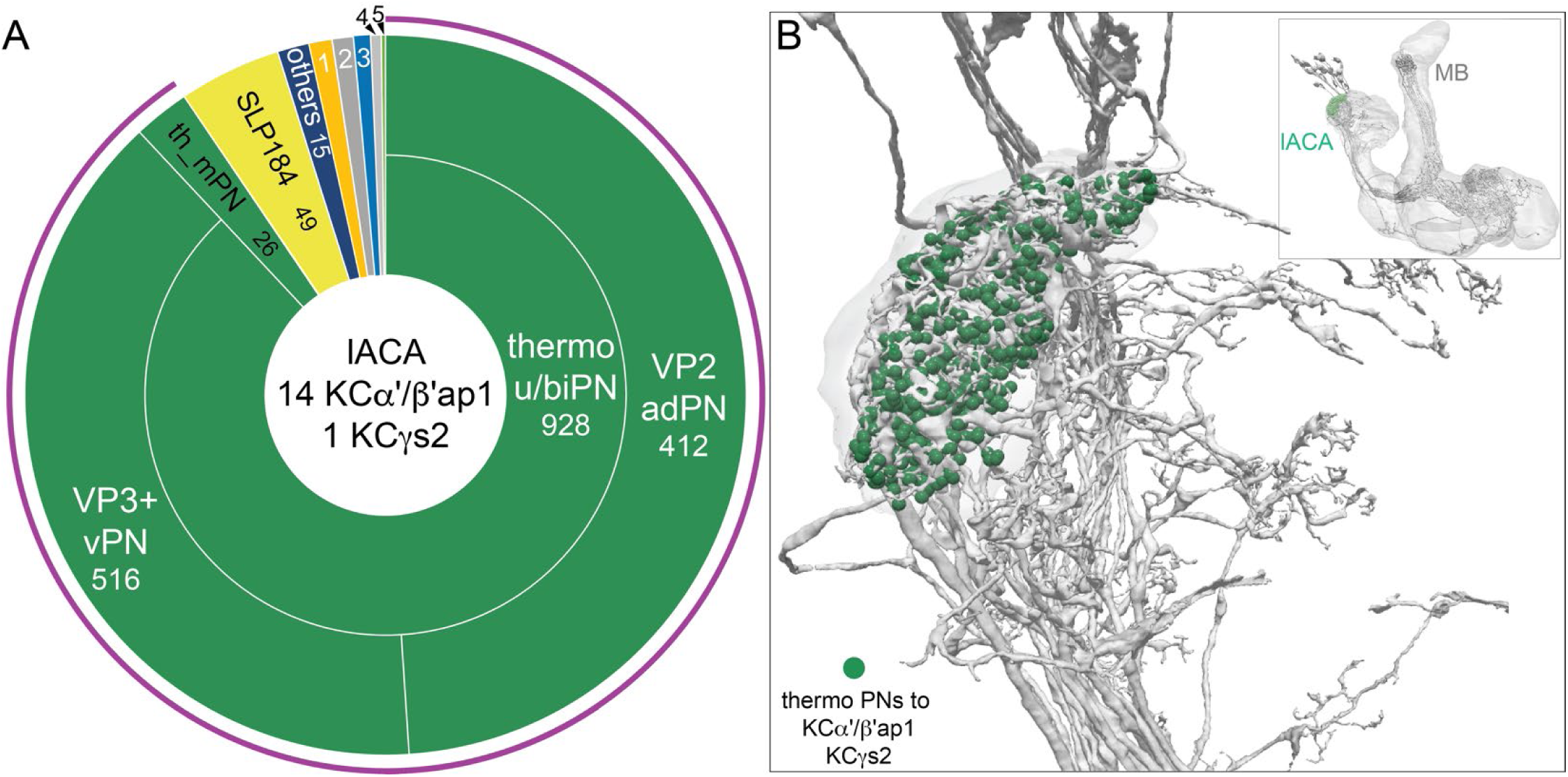
Lateral accessory calyx (lACA). The lACA is defined by the limits of the presynaptic boutons of <u>VP2 adPN</u> (1975878958) and <u>VP3+ vPN</u> (663432544); there also appear to be glia separating the lACA from main CA. Fourteen α′/β′ap1 KCs and the γs2 KC innervate the lACA. (A). A pie chart showing inputs to the 15 lACA-innervating KCs. The majority of input to these KCs is from the two thermosensory uPNs, VP2 adPN and VP3+ vPN, that contribute 928 synapses, of which the single γs2 KC receives 268. Other prominent inputs are one local interneuron (<u>SLP184</u>) and two thermo/hygrosensory mPNs. Eight interneurons (other) contribute 15 synapses. Sources of input indicated by the other numbered sectors are as follows: 1, circadian clock-associated neurons (1.0%); 2, APL (0.9%); 3, MB-C1 (0.7%); 4, KC-to-KC connections and 5, other PNs (0.2%). The total temperature information presented to KCs by PNs is indicated by the purple arc around the outer layer. Note that KCs in lACA have dramatically less KC-to-KC and APL input than those in the dACA and the vACA. (B) Synaptic connections from thermo uPNs (green) to KCs (grey). The inset shows the orientation of the MB.

### Randomness and structure in sensory inputs to KCs

Sensory input to the MB calyces shows clear structure across modalities, with visual VPN/LVIN and thermo/hygrosensory PN input targeted to specific KC types (as described above and discussed more fully below). This raises the question of whether olfactory inputs, in particular from uPNs, also exhibit structure in their inputs to the KCs. PN synapses onto KCs in the CA have a characteristic structure in which each bouton is surrounded by a claw-like KC process (Yasuyama et al., 2002; Figure 9-figure supplement 1F; Video 21), which are strikingly reminiscent of the mossy fiber-granule cell synapses found in the vertebrate cerebellum (Huang et al., 2013). Each KC has an average of 5.6 dendritic claws in the CA and requires simultaneous inputs from a combination of PNs to spike (Gruntman and Turner, 2013). The synapses between the VPNs and the KCγd dendrites have a more typical morphology, lacking the claw-like structure seen in PN-to-KC synapses in the CA (Video 22).

Previous work suggested that KCs sample olfactory uPNs without apparent structure in both the larva (Eichler et al., 2017) and the adult (Caron et al., 2013), but a recent analysis of the FAFB EM dataset identified convergence of specific PNs that was inconsistent with random sampling (Zheng et al., 2020; see also Gruntman and Turner, 2013). Developmental mechanisms have the potential to bias PN-KC connections. Both PN and KC cell types are generated in a highly stereotyped developmental order (Lee et al., 1999; Yu et al., 2010), and PNs flexibly adjust the number of their presynaptic boutons based on the availability of KC dendrites (Elkahlah et al., 2020).

To look for potential structure in olfactory uPN inputs to KCs, we computed a binary uPN-to- KC connection matrix using a cutoff of five synapses. We performed principal components analysis (PCA) on this connectivity, which can provide indications of structure (Caron et al., 2013; Eichler et al., 2017), and compared the results with PCA on synthetic connectivity matrices constructed by assuming KCs randomly sample their inputs in proportion to the total number of KC connections formed with each uPN (Figure 13A). Three principal components (PCs) are clearly larger than the corresponding values in the random model. Much of this deviation from the random model is due to differential sampling of olfactory glomeruli by γ, α′/β′ and α/β KCs. In particular, input to α′/β′ and α/β KCs is more strongly skewed toward specific highly-represented glomeruli, most notably DP1m and DM1, while the distribution for γ KCs is more uniform (Figure 13B). This suggests that the random model should be extended to allow for uPN connection probabilities that depend on both the uPN and KC types. However, some deviations from the extended random model are still present in the data, as can be seen when PCA is performed on lobe-specific connectivity matrices (Figure 13A). Note, for example, the first PC for the α/β KCs (also see Figure 13-figure supplements 1 and 2).

**Figure 13.**
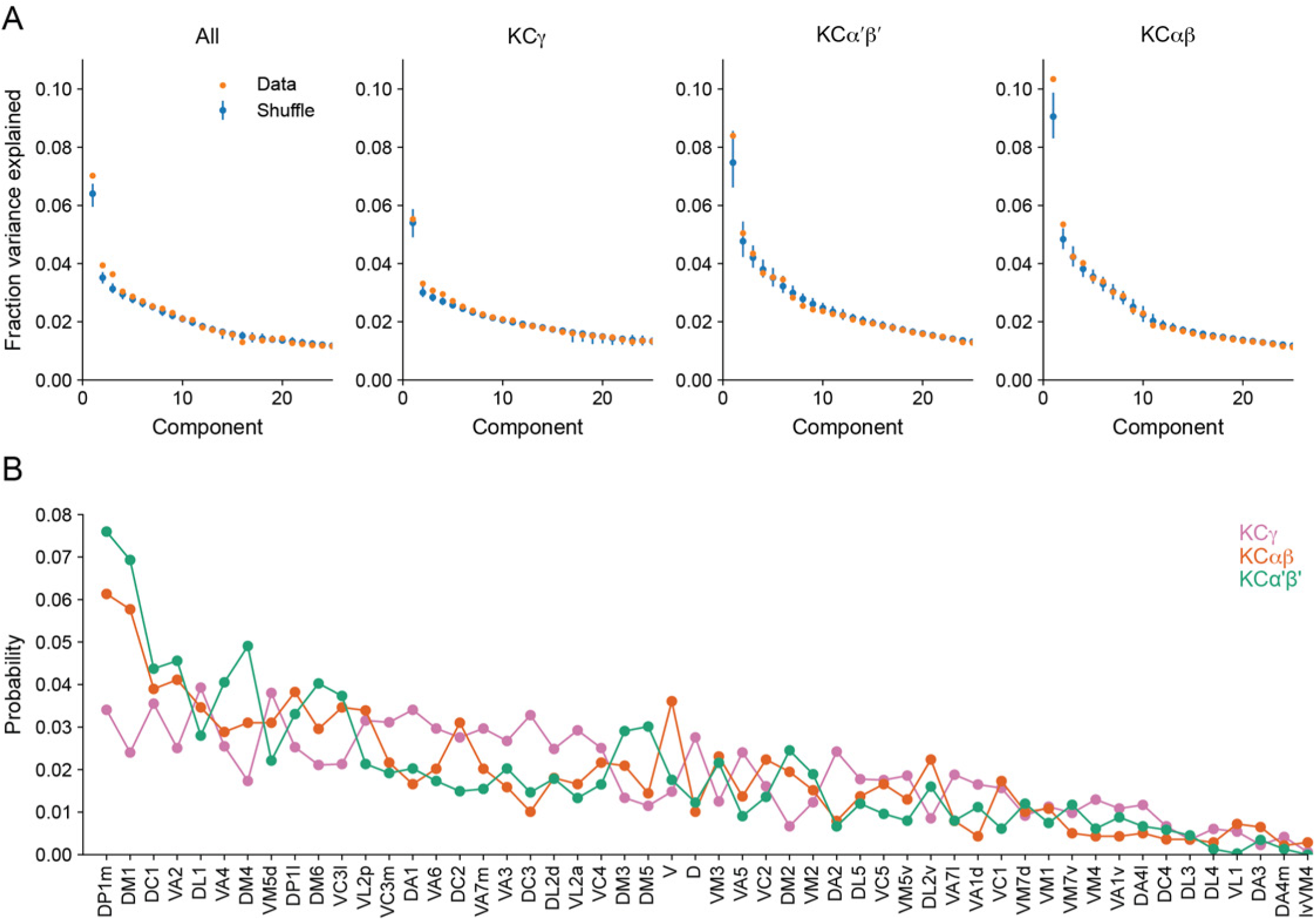
Comparison of KC input connectivity to random models. (A) Fraction of variance explained by components identified via principal components analysis of the olfactory uPN to KC input connectivity matrix—a binary matrix containing ones and zeros for present or absent connections between KCs and their inputs, at a threshold of 5 synapses. Results are shown for the reconstructed data (orange) and for a collection of shuffled models (blue) in which each KC retains the same total number of connections but samples among all uPNs randomly, with a probability proportional to the total number of connections each uPN makes. Such models therefore retain the degree distribution across KCs, (that is, the probability distribution of the number of claws formed by individual KCs) and the average connection probability for each uPN, but no other structure. Bars indicate 95% confidence intervals for shuffled models. Deviations in the first few components indicate structure inconsistent with a random model. Left: All KCs; Right: Analysis restricted to the indicated KC subtypes (visual α/βp and γd KCs, as well as γs KCs, were excluded). (B) Probability of uPN-to-KC connections from each olfactory glomerulus, sorted by most to least well-connected. Probabilities are plotted separately for the KC subtypes shown in (A).

We reasoned that this residual structure might arise from the spatial organization of inputs in the CA, so we analysed the spatial arrangement of uPN-to-KC connections and its impact on the KC odor representation. We determined the centroid locations of uPN axonal boutons within the CA by using spatial clustering of PN-to-KC synapses. Boutons belonging to uPNs from the same glomerulus were nearer, on average, than those from different glomeruli (Figure 14A). From these distributions, we computed the average number of boutons within a given radius of each centroid (Fig. 14B). These neighboring boutons were used to construct models with PN-to-KC connectivity randomly shuffled within a specified radius r (Figure 14C). This produces models in which large-scale organization (at spatial scales greater than r) is preserved, while local organization (at spatial scales less than r) is random (note that the model and the data are identical for r = 0). We computed statistics that quantified properties of the KC representation for our shuffled models, as a function of r. The first is the participation ratio of the PN-to-KC weight matrix, which quantifies how uniformly represented each glomerulus is across the inputs to all KCs. The second statistic is the dimension of the KC representation in a model in which KCs fire sparsely in response to odors that activate random patterns of PNs. Previous work has shown that this quantity determines the ability of a linear readout of KC activity to perform odor discrimination (Litwin-Kumar et al., 2017). Our analysis reveals that the participation ratio and dimension are lower for the true data than for the shuffled models, as expected from non-random structure, although the effect is modest (Figure 14D). Noticeable effects are present when the length scales for random shuffling is greater than approximately 10 μm. The effect is strongest for α/β and α′/β′ KCs, while the effect of spatial organization of the γ KC inputs appears to be minimal.

**Figure 14.**
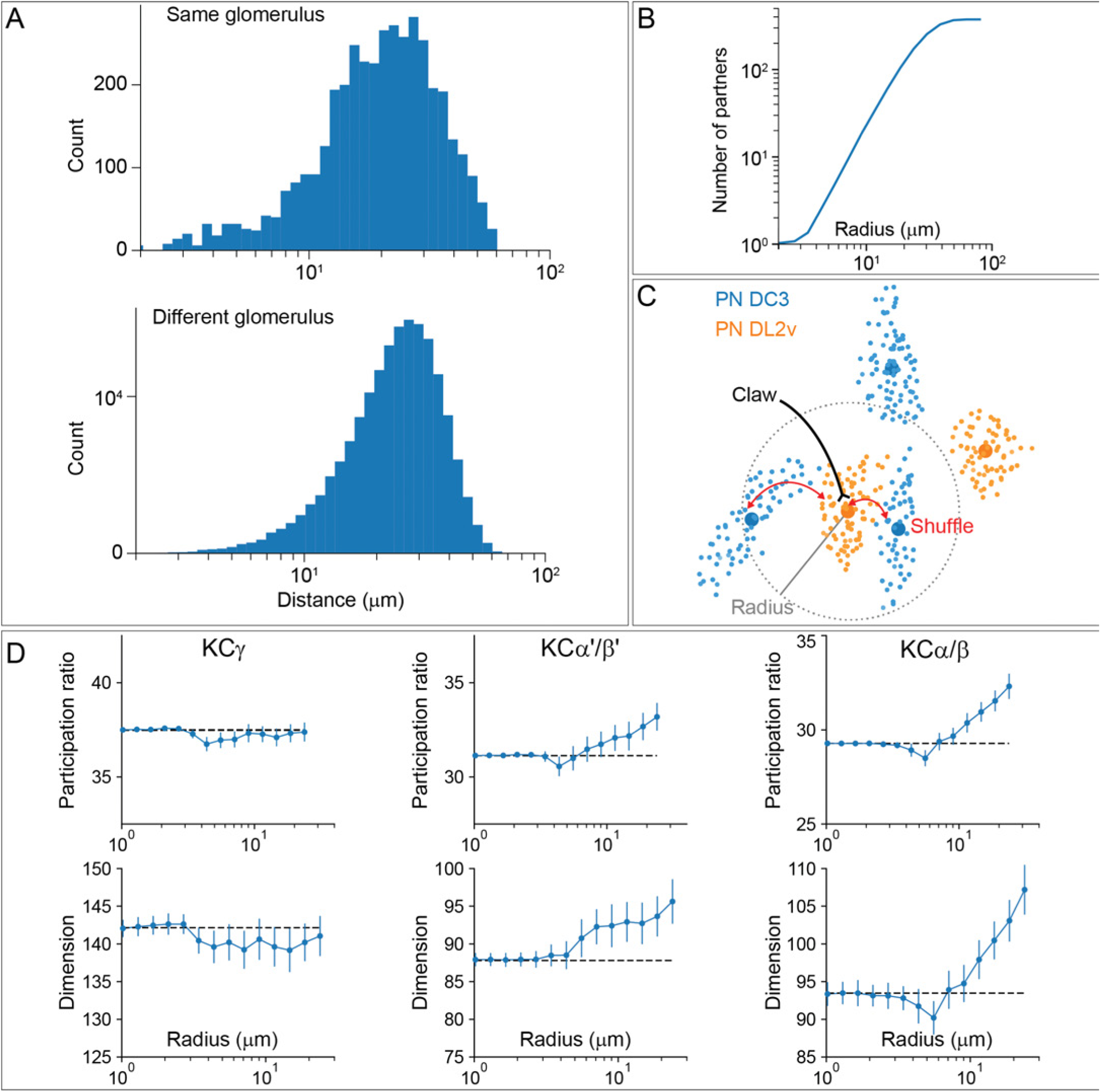
Effect of spatial organization on the KC representation. (A) Histogram of pairwise distances between boutons formed by PNs from the same glomerulus (top) or different glomeruli (bottom). (B) Average number of nearby boutons within a given radius of each PN bouton. (C) Illustration of shuffling procedure used to randomize PN-to-KC connectivity within a given radius. Small dots: synapses formed by each PN onto KCs. Large dots: Centroid location of identified boutons. Claws are randomly reassigned (red arrows) to boutons whose centroids lie within the given radius, indicated by the dotted grey line. (D) Participation ratio (top) and dimension (bottom) of models constructed by shuffling connectivity within a specific radius. The dependence of these two quantities on the shuffle radius shows a modest effect of the spatial organization of the CA on the KC representation in the model.

This analysis indicates that uPN inputs depend on KC subtype and on the spatial organization of connections within the main calyx due to locally restricted sampling. This spatial structure has only a modest effect on the dimensionality of the KC olfactory representation for simulated odors that activate random ensembles of PNs, and its effect on KC inputs does not noticeably persist at the level of KC outputs to MBONs (Figure 13-figure supplement 2). Nevertheless, it might be relevant for specific odor categories. More broadly, our analysis of sensory inputs to KCs reveals clear specialization of function across modalities supported by segregated modality-specific connectivity within the accessory calyces as well as non-olfactory input to the main calyx (see Figure 15-figure supplement 1A).

### Segregation of information flow through the MB

If the segregation of information seen in the KCs, especially across sensory modalities, is maintained at the level of MB output, this could have important implications for the specificity of learned associations. To explore this possibility, we first computed the fraction of inputs that KC types receive from PNs conveying different sensory modalities (Figure 15-figure supplement 2A) and then we computed the fraction of input to each MBON provided by KCs specialized for different sensory modalities (Figure 15-figure supplement 2B and C). The PN-to-KC input fractions (Figure 15-figure supplement 2A) quantify the specialization of KCs for particular sensory modalities as described above. For example, α/βp, γd and γs1 KCs receive more than 90% of their input from VPNs and LVINs, while γs2,3,4 KCs receive more than 50% of their input from thermo-hygrosensory PNs. From this information, we divided KCs into three groups: 1664 olfactory KCs, 102 olfactory + thermo/hygrosensory KCs, and 161 olfactory + visual KCs. We then computed the fraction of KC input to each of the MBONs that came from each of these three KC groups (Figure 15-figure supplement 2B and C; this connectivity is also shown more finely divided into KC types in Figure 15-figure supplement 3). Typical MBONs innervating the α′/β′ lobes show a gradation of input from KCs that receive thermo/hygrosensory inputs, with MBON16 (α′3ap) having 36% of its effective input coming from this source. Typical MBONs of the α/β lobes similarly show varying degrees of KC input of the olfactory + visual class. Typical MBONs of the γ lobe and CA are driven predominantly by olfactory KCs. Atypical MBONs are more strongly innervated by KCs with non-olfactory input.

By multiplying matrices of PN-KC and KC-MBON connectivity fractions, we computed an effective PN-to-MBON connectivity (Figure 15, Figure 15-figure supplement 1B and C). This effective connectivity shows a surprisingly high amount of non-olfactory sensory input to the MBONs, with some MBONs predominantly devoted to non-olfactory modalities. While the majority of the effective input to the typical MBONs is olfactory, 16% of the effective input to MBONs innervating the α/β lobes, and 9% for the γ lobe MBONs, is visual, with one of the two MBON19 (α2p3p)s having 62% visual effective input (Figures 15, 15-figure supplement 1). Thermo-hygrosensory PNs constitute 7% of the effective input to the α′/β′ MBONs. Compared to typical MBONs, atypical MBONs have a higher fraction of their input from non-olfactory sources: 24% of the effective input to atypical MBONs innervating the α′/β′ lobes is thermo-hygrosensory, and 26% of the effective input to γ-lobe atypical MBONs is visual. In particular, MBON26 (β′2d) has 30% thermo-hygrosensory effective input, and MBON27 (γ5d) has 80% visual effective input. Thus, these MBONs may preferentially participate in non-olfactory or multimodal associative memories.

**Figure 15.**
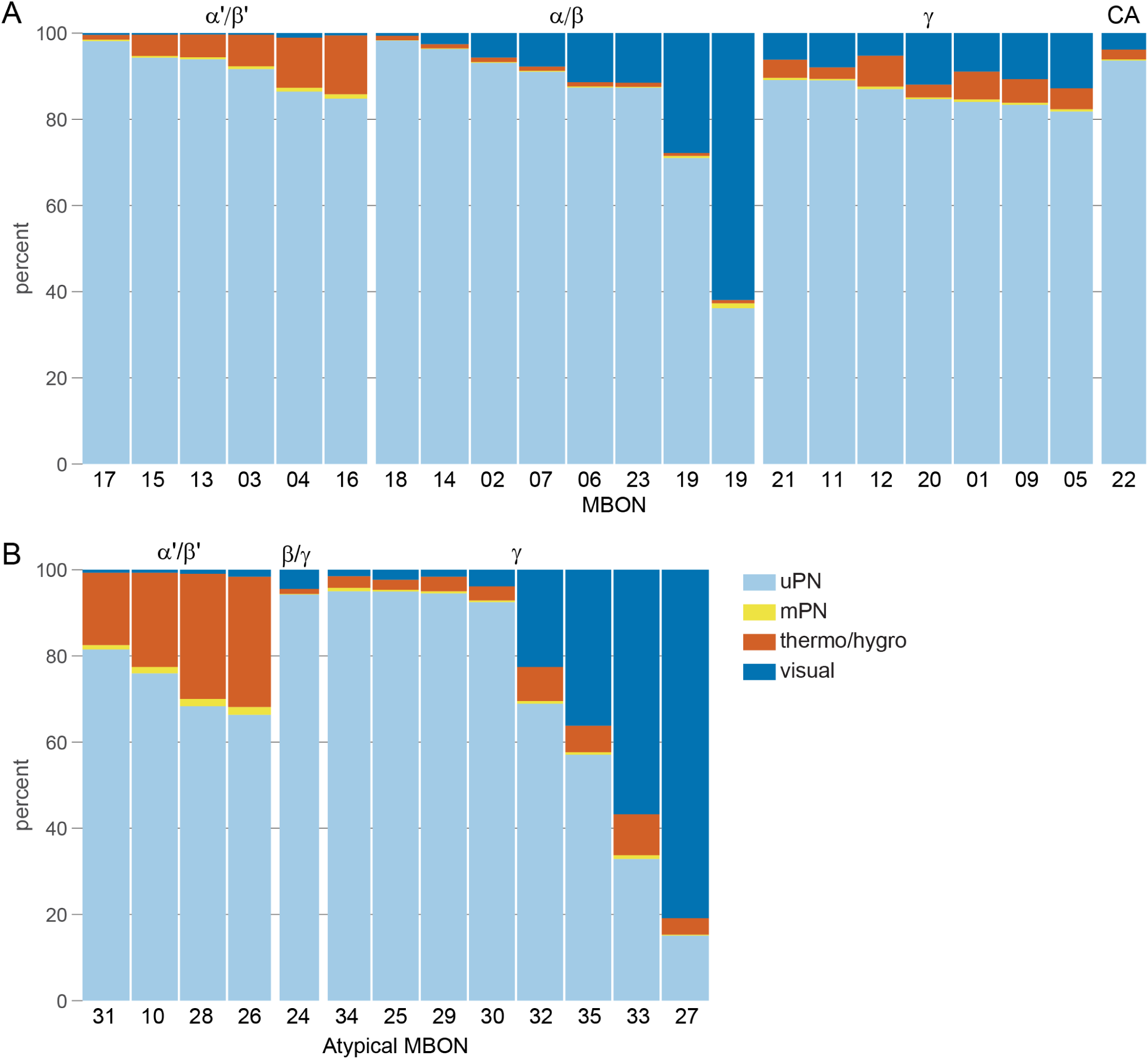
Structure in PN-KC-MBON connectivity. (A) Effective PN to MBON connectivity. MBONs within the α′/β′ lobe receive input primarily from uniglomerular olfactory PNs (uPNs), but they also show a gradation of input from thermo-hygrosensory PNs. MBONs from the α/β lobes show a similar, although stronger, variation in the amount of input they receive from visual PNs. MBONs from the γ lobe are more uniformly innervated. For cases where the MBON cell type contains more than one cell, results from the individual cells have been averaged in this figure, with the exception of the two MBON19 (α2p3p)s; both MBON19s are specialized for visual input, but to different extents. (B) Effective PN to atypical MBON connectivity. The gradation in non-olfactory input is stronger than for typical MBONs. In particular, the majority of the input to MBON33 (γ2γ3) and MBON27 (γ5d) is visual. In this figure and its supplements, the MBONs have been grouped by the MB lobe they primarily innervate and then ordered, within these groups, according to their selectivity. MBON24 (β2γ5) receives input from both the β and γ lobes. Percentages indicate the amount of input to each MBON from a particular PN type divided by the total sensory input to that MBON.

### Downstream targets of MBONs

The preceding analysis revealed segregated pathways through the MB that carry distinct sensory signals. We next investigated the extent to which these pathways remained segregated in the outputs of the MBONs by comparing the similarity of PN inputs between pairs of MBONs with the similarity of their output targets. For this purpose, we used cosine similarity, which measures the alignment between the inputs (or outputs) of the two neurons, without being affected by the total number of synapses they make. It also takes into account synapse counts, so that stronger connections influence the measure more than weaker connections, without using any arbitrary cutoff threshold. If two neurons have no shared partners, their cosine similarity will be 0, and if they target the exact same partners with the exact same relative strength, their cosine similarity will be 1.

The similarity of the PN input to pairs of MBONs reflects the selectivities seen in Figure 15 as well as revealing some more subtle structure (Figure 16, left panel). These similarities are, at least to some extent, preserved at the output level (Figure 16, right panel), particularly for the similarities among MBONs innervating the same lobes (Figure 16-figure supplement 1; results are quantified in Figure 16-figure supplement 2). This suggests that the parallel processing of different sensory modalities is, in some cases, preserved all the way from the PNs to the downstream targets of the MBONs. Beyond this organization, unbiased clustering of MBONs based solely on their output connectivity reveals additional groups of MBONs that have similar downstream targets but different effective PN input (Figure 16-figure supplements 3 and 4). For example, MBON06 (β1>α), and MBON18 (α2sc) innervate different dendritic compartments (Figure 16-figure supplement 4) but have a high cosine similarity score based on their output (Figure 16-figure supplement 3). Conversely, MBON20 (γ1γ2) and MBON25 (γ1γ2) have low cosine similarity based on their output although their dendrites innervate the same compartments.

**Figure 16.**
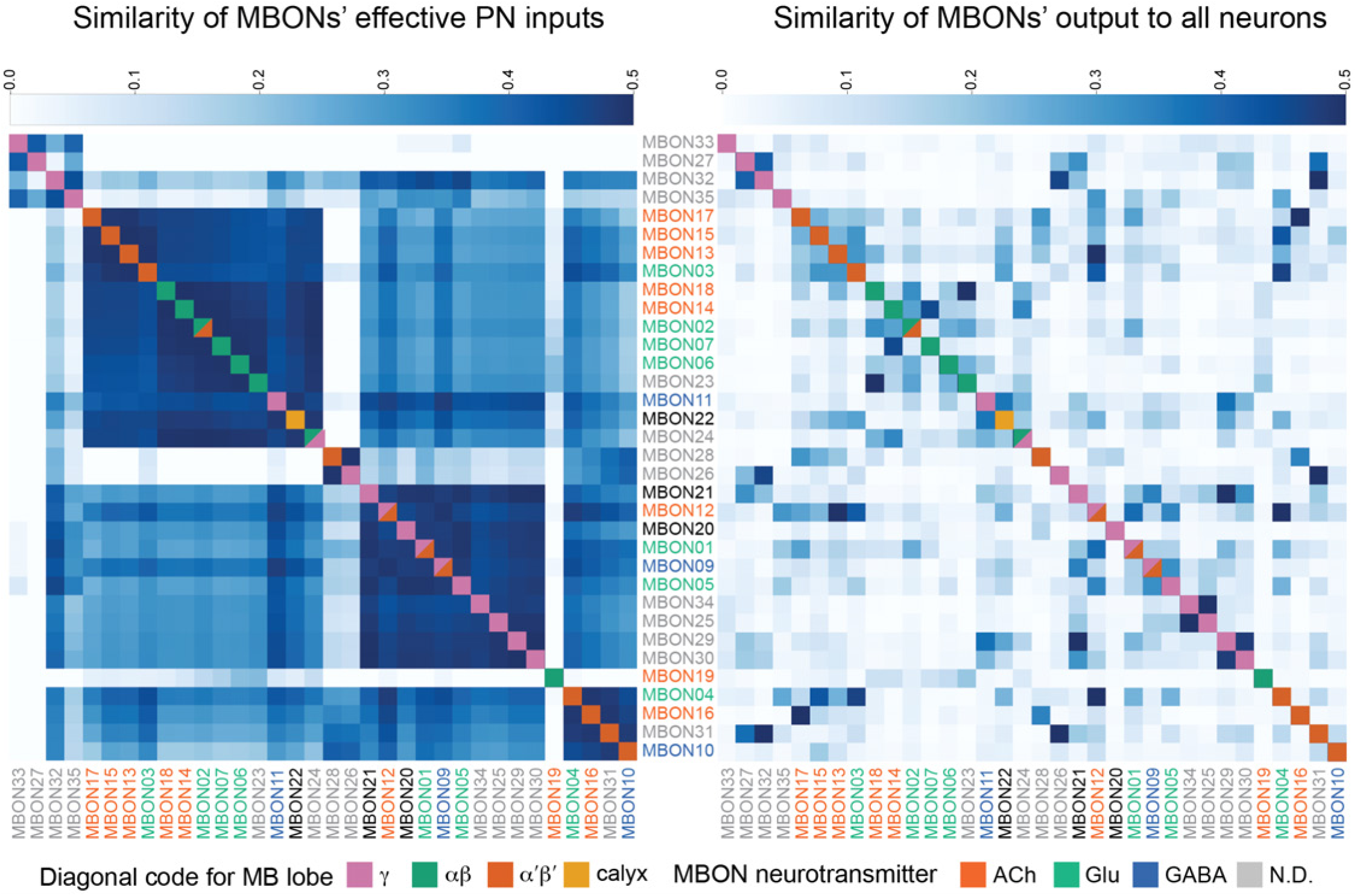
MBON input and output similarity structure. A comparison of MBON inputs and outputs through the lens of similarity structure. Left: cosine similarity of MBONs based on their effective PN inputs via KCs (computed as in Figure 15). Each cell of the heat map indicates the input similarity of the indicated pair of MBONs. A value of 0.5 indicates that the PN subpopulations conveying input to the two MBONs via KCs are half overlapping and half disjoint. Right: cosine similarity of MBONs based on their outputs to all neurons (unnamed neuronal fragments have been excluded). The resemblance between the left and right plots suggests that MBON outputs preserve some of the parallel structure in the PN-KC-MBON pathway. In each plot, the square in the diagonal is color-coded to indicate the MB lobe in which that MBON’s dendrites lie. The MBON names are color-coded to indicate their neurotransmitter, as indicated.

To look for other factors underlying the patterns of MBON output, we considered the sign of MBON output as implied by neurotransmitter identity. Acetylcholine is highly predictive of excitation, as is GABA for inhibition. The situation for glutamate is more ambiguous. There are established cases of glutamate having inhibitory (Liu and Wilson, 2013) and excitatory (Johansen et al., 1989) action. Most cells appear to express both inhibitory and excitatory receptors for glutamate; for example, all six MBON and all ten DAN cell types for which data exists express both inhibitory GluClalpha and excitatory NMDA type receptors (Aso et al., 2019). Activation of cholinergic and glutamatergic MBONs has been associated with opposing behaviors, avoidance and approach, respectively (Aso et al., 2014b; Owald et al., 2015; Perisse et al., 2016), but the extent to which these result from the action of these MBONs on shared targets has not been established.

The degree of overlap in outputs between pairs of MBONs organized by neurotransmitter type (Figure 17) reveals structure that can be summarized by grouping MBONs by their transmitters (Figure 17-figure supplement 1). Both cholinergic and glutamatergic MBONs show similarity in their outputs with other MBONs of the same transmitter type but, interestingly, there is a matching degree of similarity in the outputs across cholinergic and glutamatergic MBONs. This is not seen in the similarities between MBONs using either of these transmitters and GABAergic MBONs (Figure 17-figure supplement 1). Thus, assuming an inhibitory role for glutamate, the convergence of cholinergic and glutamatergic MBONs on a common downstream target is likely to produce a “push-pull” effect, in which competing excitatory and inhibitory influences on the common target drive opposite behaviors.

**Figure 17.**
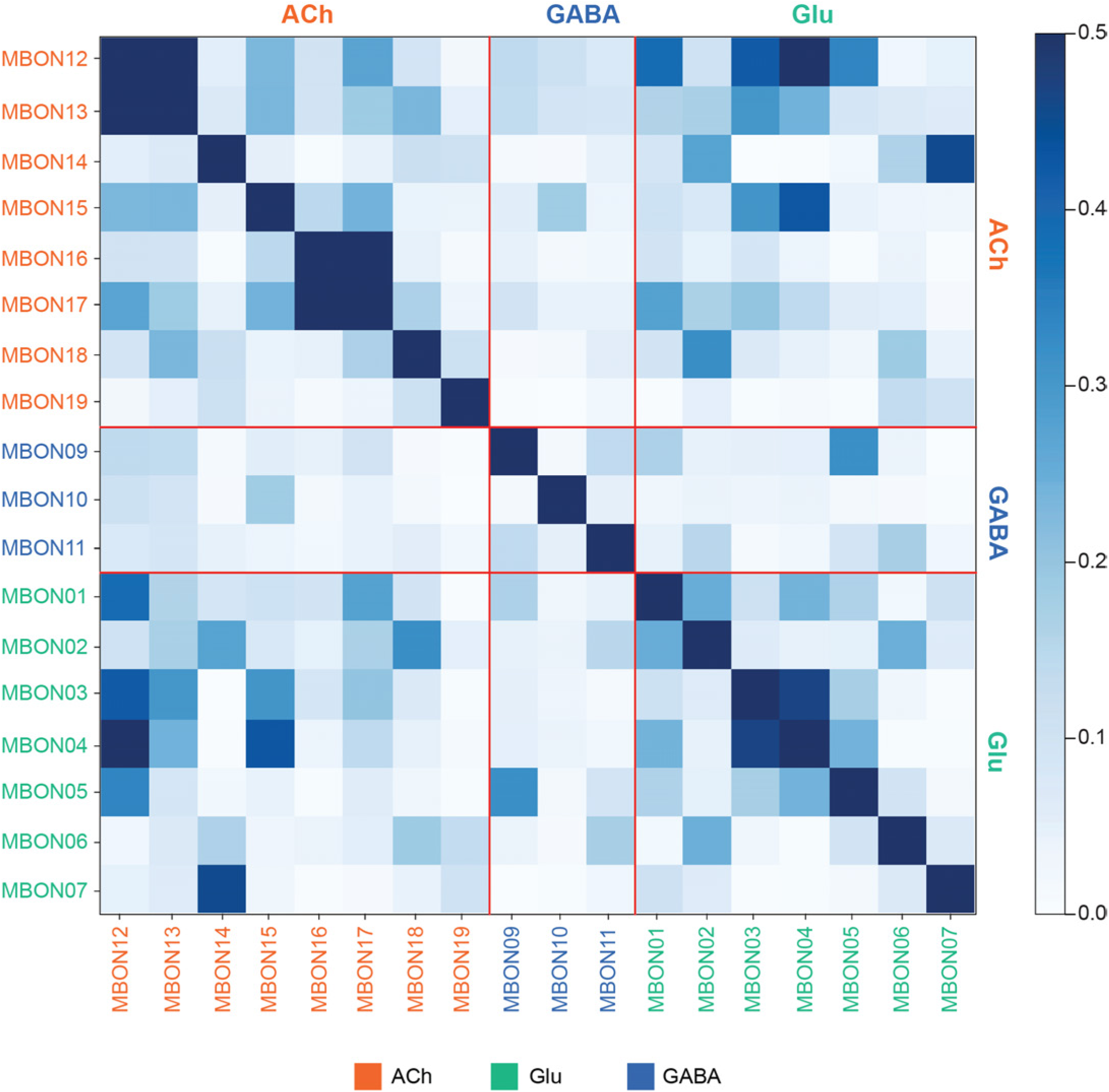
MBON output similarity by neurotransmitter. Cosine similarity of MBON outputs, for MBONs with known neurotransmitter, to all named neurons. MBONs have been grouped by neurotransmitter and then presented in numerical order. There exist many instances of convergent outputs between cholinergic and glutamatergic neurons, suggesting a widespread “push-pull” motif in MBON output convergence patterns. This phenomenon is emphasized in Figure 17-figure supplement 1 and notable examples are explored in further detail in Figure 20-figure supplement 1 and Figure 21-figure supplements 1 and 2. The MBON names are color-coded to indicate their neurotransmitter.

### The distribution of the MBON outputs differs between MBONs using different neurotransmitters

The analyses above reveal MBONs with similar downstream targets but do not identify those targets. We computed the propensity of MBONs, grouped by neurotransmitter type, to project to different brain regions (Figure 18); our results confirm and extend observations made by light microscopy (Aso et al., 2014a). All MBON types project strongly to the CRE, while cholinergic and glutamatergic MBONs (but not GABAergic MBONs) project strongly to the SMP, and to a lesser extent the SIP and SLP. This pattern suggests that the CRE, SMP, SIP, and SLP may be sites of “push-pull” interactions as described above. GABAergic MBONs are notable in that the vast majority of their projections are to the CRE, and several atypical MBONs are unique in their strong projections to the LAL. Overall, these results suggest strong biases in output connectivity patterns of MBON neurotransmitter types, viewed at the coarse level of brain area. Finer structure is also observable in the data. For instance, the downstream projection targets of MBONs exhibit spatial biases even within individual brain areas (Figure 18-figure supplement 1). Importantly, although MBONs of a given neurotransmitter type often exhibit related connectivity biases, they are certainly not homogeneous, consistent with unique behavioral roles for individual MBONs (Figure 18-figure supplement 2).

**Figure 18.**
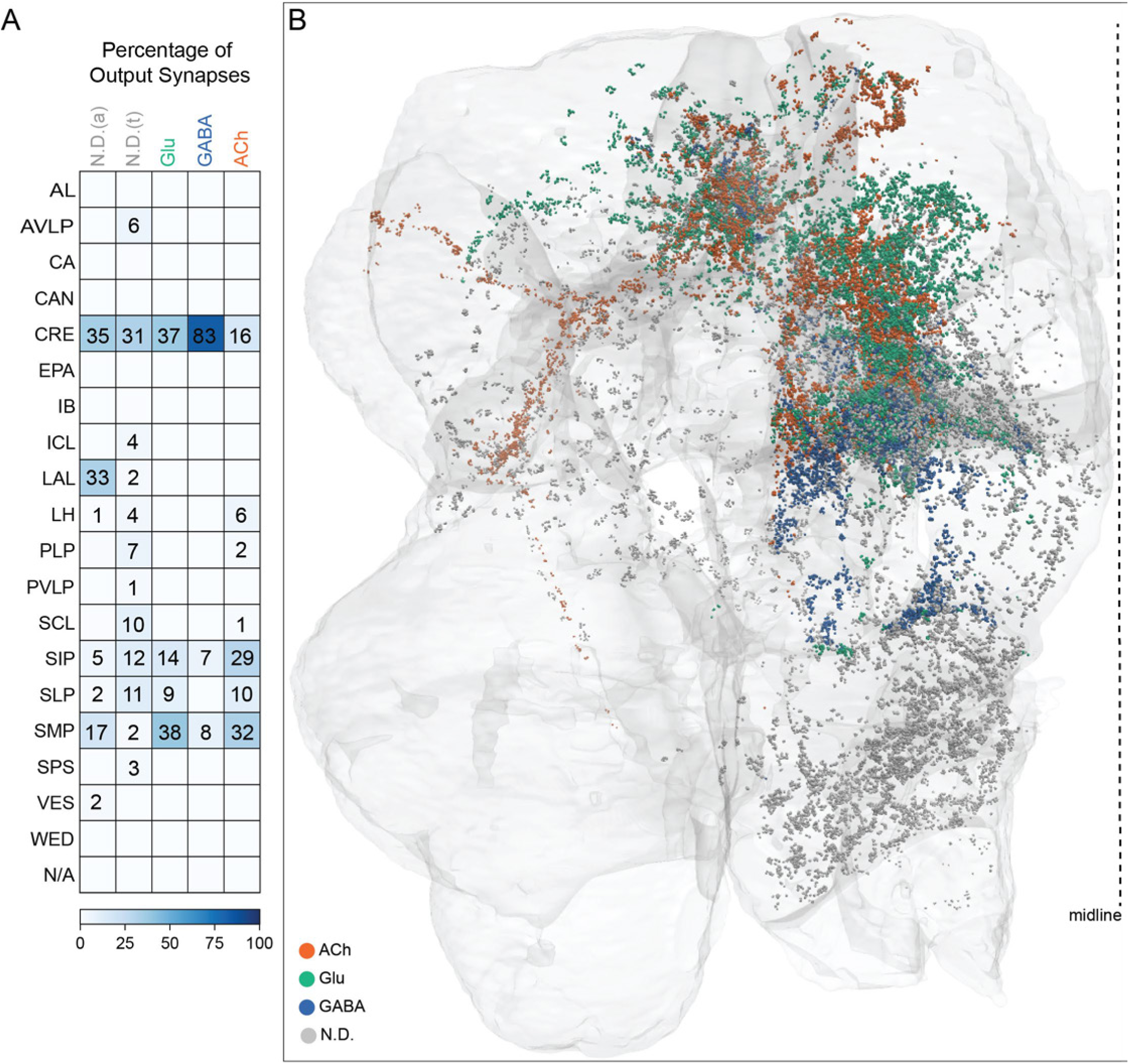
MBON output distribution by neurotransmitter. (A) Table indicating the percentage of output synapses of each MBON neurotransmitter type that reside in the given brain area; typical (t) and atypical (a) MBONs with undetermined transmitter were analyzed separately. (B) Visualization of the spatial position of MBON output synapses throughout the hemibrain volume, colored by their neurotransmitter type. Figure 18-figure supplement 1 shows the same data presented on separated brain areas.

### Neuronal pathways connecting the MB and the CX

The MB and central complex (CX), both highly structured centers, are known to carry out sophisticated computations. Communication from the MB to the CX is likely to be important for conveying information about learned associations of sensory cues and external rewards or punishment (Aso et al., 2014a, 2014b; Owald et al., 2015), novelty (Hattori et al., 2017) and sleep need (Sitaraman et al., 2015a; Dag et al. 2019), which, in turn, are expected to influence navigation, sleep and other activities governed by the CX. We describe here the complete network of strong (based on synapse number) neural pathways connecting the MB to the CX (to be discussed further in a companion manuscript on the connectome of the CX).

We found that direct communication from the MB to the CX is limited to two pathways (Figure 19). The most prevalent is MBONs connecting to the dendrites of fan-shaped body (FB) tangential neurons; 22 out of 34 MBON types make such direct connections and both typical and atypical MBONs participate. In addition, three atypical MBONs make connections in the LAL to the dendrites of three CX cell types that have axonal terminals in the nodulus (NO) (Figure 19-figure supplement 1A). The FB has been divided by anatomists into multiple layers, numbered ventrally to dorsally from one to nine (Figure 20; Video 26). Each tangential neuron’s arbors within the FB lie predominantly in a single layer and its dendrites project laterally in the CRE, SMP or SIP (morphologies are shown in Video 26). We found that MBONs preferentially target FB layers four (FBl4) and five (FBl5). Fifteen MBON types target FBl4 and account for 62% of all MBON to FB synapses; they innervate 58% of FB cell types found in this layer. Thirteen MBON types target FBl5 and account for 30% of all MBON to FB synapses; they innervate 43% of FB cell types found in this layer (Figure 20). Thirty-seven different FB cell types get direct synaptic input from MBONs, and some cell types get input from multiple MBON cell types; the fraction of each cell type’s input that comes from MBONs is shown in Figure 19. One cell type, CREFB4, has an unusually high level of MBON input, receiving nearly 20% of its synaptic input from a combination of MBON09 (γ3β′1) and MBON21 (γ4γ5) (Figure 19-figure supplement 1B and C). Among the MBONs that output onto FB neurons, MBON21, MBON33 (γ2γ3) and MBON34 (γ2) devote the highest fractions of their synaptic output connecting to FB neurons, at 16%, 9% and 8%, respectively.

**Figure 19.**
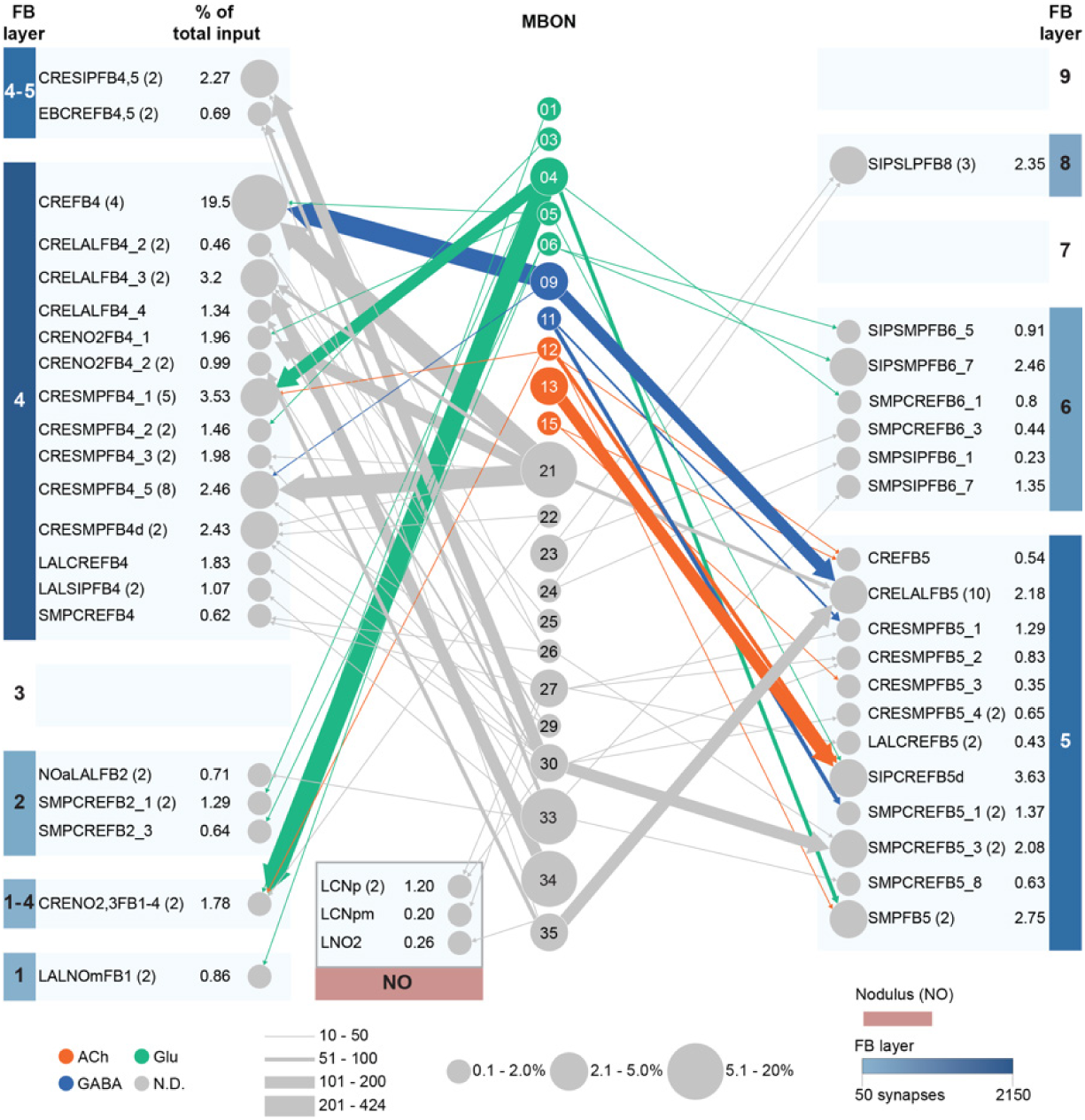
Direct connections from MBONs to the central complex (CX). In this diagram, the central column shows those MBONs that make direct connections of 10 or more synapses onto the dendrites of CX neurons. Such direct connections predominantly occur on fan-shaped body (FB) neurons. Three cell types of nodulus (NO) cell types are also direct targets (see Figure 22-figure supplement 1A); some FB cell types in layers 1, 2, 3 and 4 that are downstream of MBONs also have arbors in the NO. No direct targets were identified in other CX neuropils such as the ellipsoid body (EB) or protocerebral bridge (PB). The number in the circle indicates the MBON cell type; that is, 01 stands for MBON01. MBONs with numbers up to MBON23 are typical MBONs, while MBONs numbered between MBON24 and MBON35 are atypical. The color of the circle represents the MBON’s neurotransmitter while the size of the circle indicates what percent of that MBON’s total output (based on synapse number) goes directly to FB neurons. Numbers have been pooled for cases where there are multiple MBONs of the same cell type. The outer columns show FB cell types, arranged by the layer(s) in the FB where their output synapses are located. The name of the cell type and, in parentheses, the number of cells of that type are indicated for those with multiple cells. The percentage of that cell type’s input that comes from MBONs is given and is reflected by the size of the circle representing the cell type. All FB neuron types receive less than 5% of their total input from MBONs except for one layer 4 FB neuron type, CREFB4 (FB4R), which receives nearly 20% of its inputs from two MBONs, MBON09 (γ3β′1) and MBON21 (see Figure 22-figure supplement 1B and C). Direct MBON connections to the FB are concentrated in FB layers 4 and 5. Links to each of these FB and NO cell types in neuPrint, listed from FB layer 1 to FB layer 9 and then for the nodulus, are as follows: <u>LALNOmFB1 (FB1C)</u>, <u>CRENO2,3FB1-4 (FB1H), NOaLALFB2 (FB2A), SMPCREFB2_1 (FB2C), SMPCREFB2_2 (FB2L), CRESMPFB4_1 (FB4A), CRENO2FB4_1 (FB4C), CRESMPFB4_2 (FB4D), CRELALFB4_2 (FB4F_b), CRELALFB4_3 (FB4G), CRELALFB4_4 (FB4H), LALCREFB4 (FB4I), CRELALFB4_5 (FB4J), LALSIPFB4 (FB4L), CRENO2FB4_2 (FB4M), SMPCREFB4 (FB4N), CRESMPFB4d (FB4O), CRESMPFB4_4 (FB4P_a and FB4P_b), CREFB4 (FB4R), CRESIPFB4,5 (FB4X), EBCREFB4,5 (FB4Y), LALCREFB5 (FB5A), SIPCREFB5d (FB5AB), SMPCREFB5_1 (FB5C), CRESMPFB5_1 (FB5D), CRESMPFB5_2 (FB5E), CRESMPFB5_3 (FB5H), SMPCREFB5_3 (FB5I), SMPCREFB5 (FB5J), CRESMPFB5 (FB5K), CRESMPFB5_4 (FB5L), CRESMPFB5_5 (FB5M), CRELALFB5 (FB5V), SMPCREFB5_8 (FB5X), SMPSIPFB6_1 (FB6A), SMPCREFB6_1 (FB6P), SIPSMPFB6_5 (FB6Q), SMPSIPFB6_7 (FB6R), SIPSMPFB6_7 (FB6T), SMPCREFB6_3 (FB6V), SIPSLPFB8 (FB8F_a), LCNp, LCNpm, and LNO2.</u>

**Figure 20.**
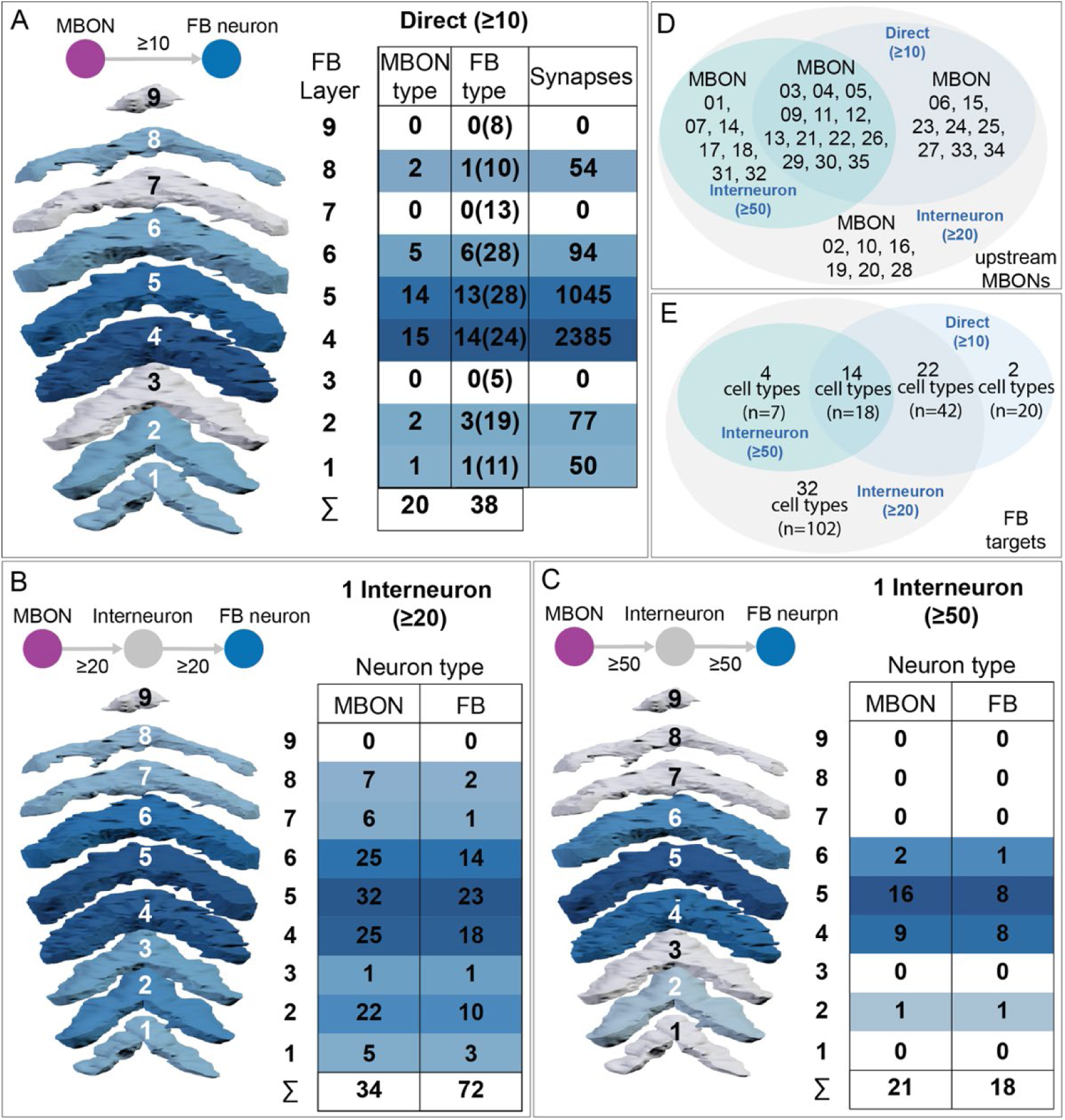
Summary of connections from MBONs to the FB. Heat maps are shown that compare the strength of connection between MBONs and FB neurons in each FB layer. (A) Direct connections. The number of MBON cell types within each layer that make synapses onto the dendrites of FB neurons, the number of FB neuron types (also within the indicated layers) they connect to, and the total number of MBON to FB neuron synapses are shown for each FB layer. The total number of cell types listed in neuPrint for each FB layer is shown in parentheses. The last row of the table shows the number of different MBON cell types that make direct connections to the FB; some MBONs connect to more than one layer. (B,C) Connections using a single interneuron. The number of MBON cell types connecting to each layer and the number of FB neuron types they connect to are shown. (B,C) Connections between the MBON and the interneuron as well as between the interneuron and the FB neuron are required to have 20 or more synapses (B) or 50 or more synapses (C). The last row of the table shows the number of different MBON cell types that make indirect connections to the FB at this threshold; some MBONs connect to more than one layer. (D) Venn diagram comparing MBON types that are connected to FB neurons directly (as in A); through 1 interneuron with threshold of 20 synapses (as in B); and through 1 interneuron with threshold of 50 synapses (as in C). (E) Venn diagram comparing FB neuron cell types that are targeted by MBONs directly (as in A), through one interneuron with threshold of 20 synapses (as in B), and through one interneuron with threshold of 50 synapses (as in C). The number of cell types and the total number of neurons (n) are indicated.

We also looked at MBON to CX connections that are mediated by a single interneuron. We separately determined strong connections, where we required at least 50 synapses from the MBON to the interneuron and then at least 50 synapses from the interneuron to the CX neuron, as well as weaker connections where these thresholds were set at 20 synapses. For these thresholds, we found that the only connected CX neurons were FB tangential neurons and two NO neurons (eight and two at thresholds of 20 and 50, respectively). We found that all 34 MBON types output to FB neurons when the threshold of 20 synapses at each connecting point was used (Figure 20B). Moreover, for this threshold all FB layers except layer 9 receive indirect input from MBONs. Increasing the threshold to 50 synapses results in a connectivity pattern that, at the level of which FB layers are connected, is very similar to that displayed by direct connections (compare Figure 20 panels A and C). The direct and indirect connections appear to target similar FB neurons. Over 90 percent of FB cell types that are directly postsynaptic to MBONs also get indirect input through a single interneuron; conversely, 76 percent of FB cell types connected by an interneuron, when using the 50-synapse threshold, also get direct connections. A single interneuron can get input from multiple MBONs and then itself make output onto multiple FB cell types. Figure 20-figure supplement 1 and Videos 27-29 show examples of this circuit motif.

Connections from the CX to the MB are much more limited. In particular, we identified no cases where CX neurons provide direct input to MB intrinsic cell types such as KCs, APL or DPM. Indirect connections are also rare. We did identify three cases where CX neurons were upstream of DANs. FB tangential neurons often have mixed arbors outside the FB and we found layer 6 neurons that are presynaptic to PAM09 (β1ped) and PAM10 (β1) and layer 7 neurons that are presynaptic to PPL105 (α′2α2), respectively (see Figure 34). In addition, one of the 11 PAM12 (γ3) neurons (862705904) is atypical in that it receives 12 percent of its input from the FB columnar neuron FB2-5RUB (Video 33). This same FB2-5RUB cell type makes strong inputs onto the atypical MBON30 (γ1γ2γ3); with 451 synapses, it is MBON30’s strongest input outside the MB lobes (see Figure 8-figure supplement 9, Video 15 and Figure 25-figure supplement 1).

### A network of convergence downstream of MBONs

We found that different MBONs frequently share downstream targets resulting in an extensive network of convergence. Such convergence would provide a mechanism to integrate the outputs from different MB compartments. At a threshold of 10 synapses, ∼1550 neurons are downstream targets of at least one MBON, with nearly 40 percent of these (∼ 600 neurons), getting input from at least two MBON types (Figure 21A). Downstream targets with multiple inputs are not unexpected given that the total of all MBON downstream targets is close to 10% of central brain neurons and individual MBONs tend to have many downstream partners (an average of 57 at the threshold we used). However, a null model in which MBON outputs are randomly sampled by central brain neurons, with weighted probabilities according to their total number of input synapses, yields ∼3200 downstream targets, only 14% of which receive input from at least two MBON types. This result indicates that MBON outputs are convergent beyond what would be expected from random connectivity alone. Both typical (Figure 21B) and atypical (Figure 21B′) MBONs participate in such convergent networks at roughly similar levels, after correcting for the smaller number of atypical MBONs, and individual cells often get input from both types of MBONs. As discussed below, MBONs are highly overrepresented among the downstream targets of other MBONs. Lateral horn (LH) neurons and lateral horn output neurons (TON) are also frequent targets of MBON convergence, being particularly overrepresented among those receiving input from seven or more MBONs. Figures 21-figure supplements 1 and 2 show examples of convergence neurons with seven and 11 upstream MBONs, respectively, and include MBONs with neurotransmitters of opposite sign, extreme examples of the “push-pull” arrangement introduced in Figure 17.

**Figure 21.**
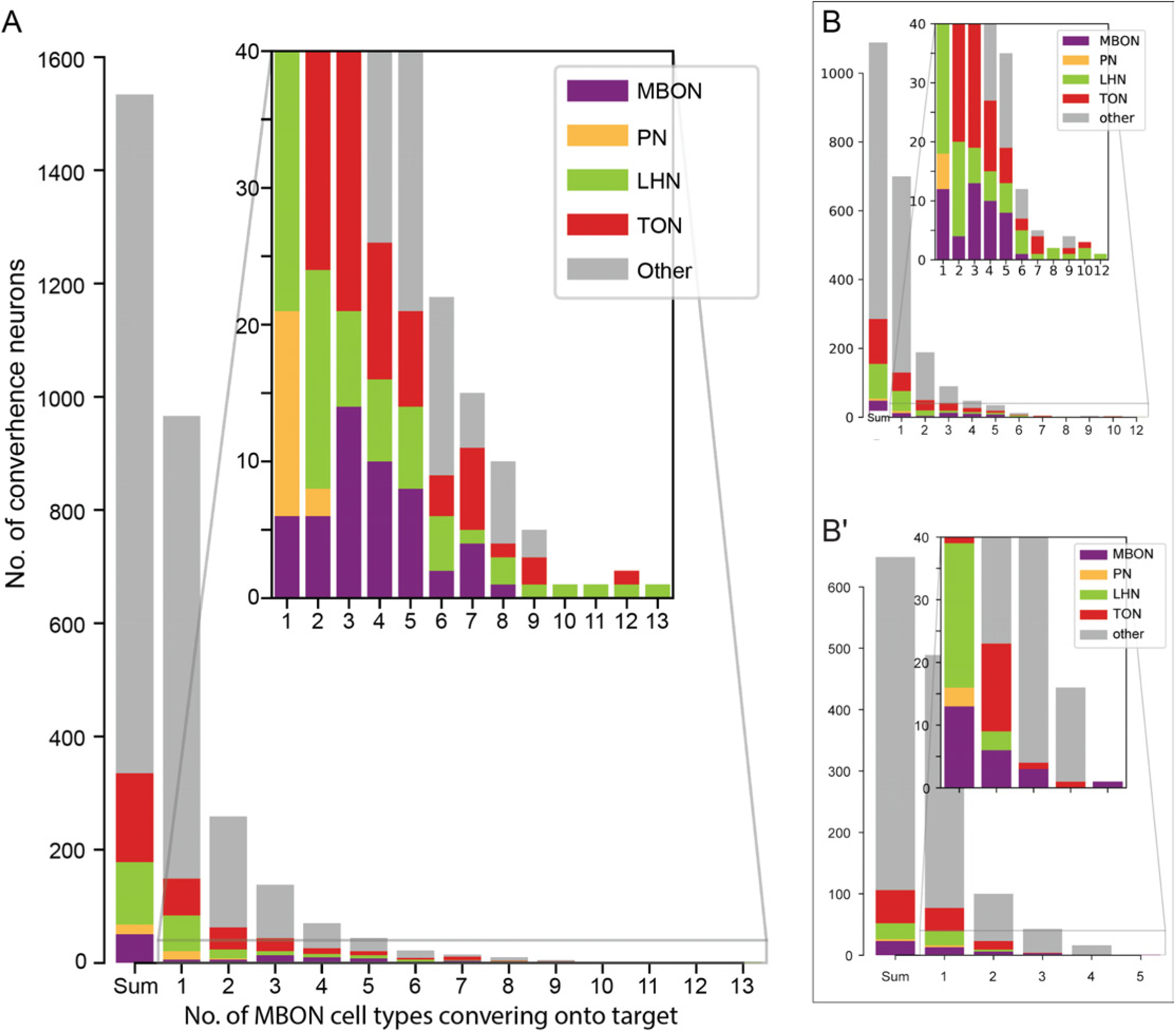
Downstream targets of MBONs often receive input from more than one MBON. (A) About 1550 neurons are downstream (including axo-axonal connections) of one or more of the 34 MBON cell types when a threshold of 10 synapses is used. Among them, about 600 are downstream of at least 2 MBON cell types and are therefore capable of integrating information from multiple MBONs. The inset shows an expanded view of the indicated portion of the plot. These neurons include a range of well-characterized neuron types, including MBONs, olfactory projection neurons (PNs), lateral horn neurons (LHNs), and third order olfactory neurons (TONs; downstream targets of PNs outside the LH) as well as less characterized neurons (other). Among the neurons that receive input from 10 or more MBON types, LHNs are heavily over-represented. (B, B′) The distribution shown in (A) has been divided into two separate histograms, one showing neurons downstream of multiple typical MBONs, in which atypical MBONs have not been counted (B), and one showing neurons downstream of multiple atypical MBONs, in which typical MBONs have not been counted (B′). The insets show expanded views of the indicated portions of each plot. Atypical MBONs show less convergence than the typical MBONs and do not preferentially converge onto LHNs.

### Three MBON types support a multilayer feedforward network in the MB lobes

MBON05 (γ4>γ1γ2), MBON06 (β1>α) and MBON11 (γ1pedc>α/β) have been previously shown, based on light microscopic data, to have a high fraction of their output directed to the dendrites of other MBONs within the MB lobes, providing pathways for a multi-layered feedforward MBON network (Aso et al 2014a). All three of these MBONs are putatively inhibitory based on their use of GABA or glutamate as neurotransmitters. We have now been able to map this network comprehensively and to look at the spatial distribution of the synapses from the three feedforward MBONs to their targets. It has been noted that in some cases, such synapses are clustered close to the root of the target MBON’s dendrites where they would be well positioned to cause a shunting effect (Perisse et al., 2016; Takemura et al., 2017; Felsenberg et al., 2018). We examined all feedforward connections between MBON05 (γ4>γ1γ2), MBON06 (β1>α) and MBON11 (γ1pedc>α/β) and their target MBONs and found that the same upstream neuron often has distinct synapse distributions on its different downstream MBONs (Figure 22), as previously observed for MBON11 (Perisse et al., 2016; Felsenberg et al., 2018). In several cases in the α lobe where such distributions had also been mapped by Takemura et al. (2017), the two EM datasets are consistent. To quantitate these distributions, we compared the distances from the root of the target MBON’s dendritic tree to the synapses onto it made by KCs and by feedforward MBONs. We found that, in most cases, the locations of synapses from feedforward neurons closely tracked those of KCs, showing no obvious spatial bias; an example is provided by the two feedforward MBONs that target MBON14 (α3): MBON06 (β1>α) and MBON11 (γ1pedc>α/β) (Figure 22B). An example of a biased spatial distribution is provided by MBON06, which feeds forward onto both MBON07 (α1) cells at locations that are shifted closer to the dendritic root (Figure 22D). Conversely, we found MBON11 (γ1pedc>α/β) synapses onto MBON03 (β′2mp) around the edge of its dendrite far away from its root (Figure 22E; consistent with Felsenberg et al., 2019), while MBON11’s synapses onto the two MBON07 cells are uniformly distributed (Figure 22D).

**Figure 22.**
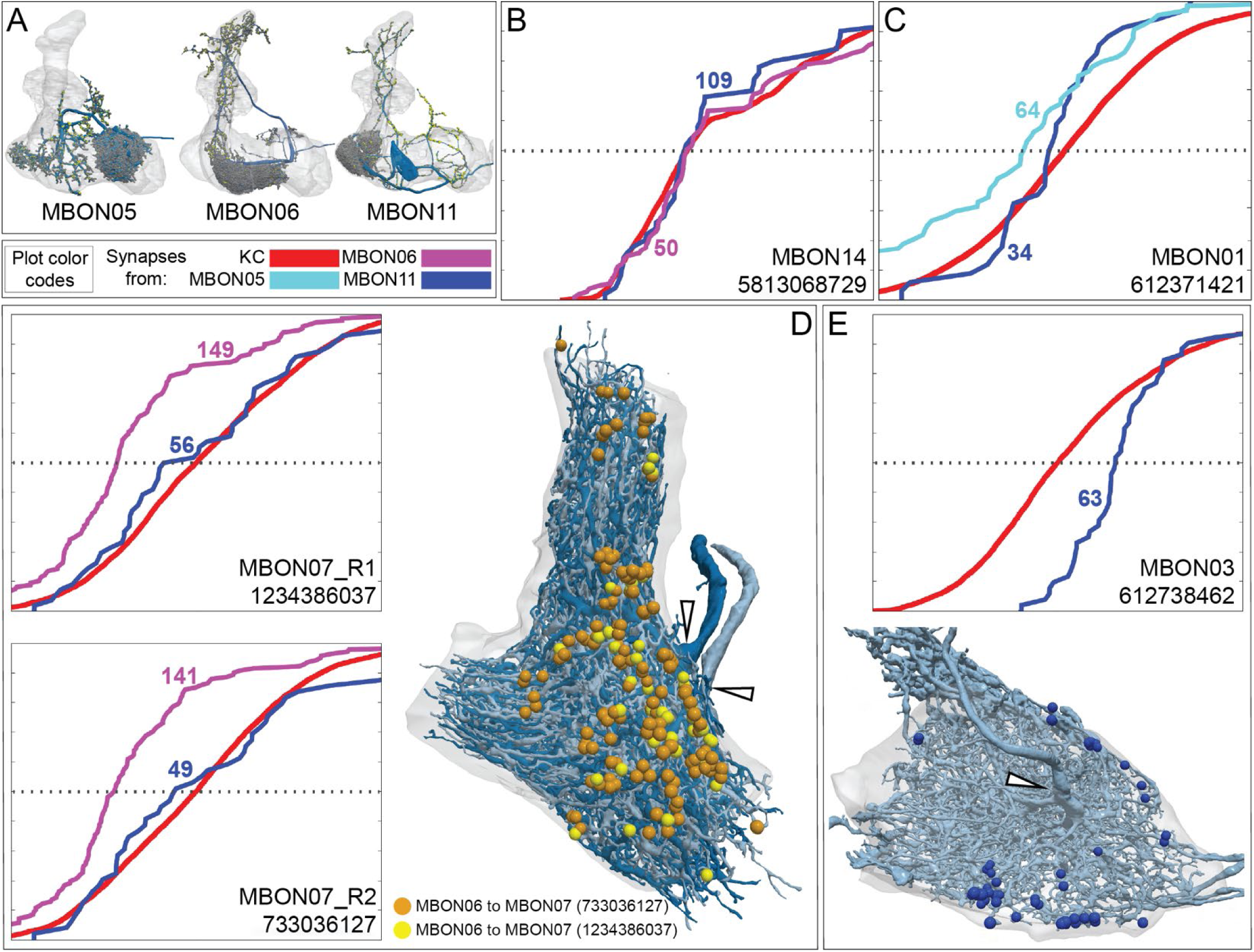
Spatial distribution of synapses from feedforward MBONs onto the dendrites of other MBONs. (A) The three MBON types that make feedforward connections to other compartments within the MB lobes are shown. Neurons are shown in blue with presynaptic and postsynaptic sites shown as yellow and grey dots, respectively. (B-E) The cumulative fraction of synaptic connections from a feedforward MBON to its target is plotted as a function of the normalized distance from the root of the target MBON’s dendritic tree, on a linear scale from 0 to 1. The name of the target neuron is given in the lower right corner of the panel. The horizontal dotted line indicates the distance from the root of the target MBON’s dendritic tree at which the half of the synapses from KCs to that MBON have occurred. The lines in these graphs are color-coded as indicated by key shown below panel A. The numbers indicate the total number of synapses between the neuron of the same color and the target MBON. (B) The synapses from MBON06 (β1>α) and MBON11 (γ1pedc>α/β) onto MBON14 in α3 have a similar spatial distribution as MBON14’s synapses from KCs. (C) MBON01’s dendrites in γ5 receives synaptic connections from MBON11 closer to MBON01’s dendritic root than KC inputs, a bias even more pronounced in the distribution of synapses MBON01 (γ5β’2a) receives from MBON05 (γ4>γ1γ2). (D) The two MBON07 cells in the α1 compartment each receive synaptic input from MBON06 (β1>α) at sites close to the root of their dendrites (hollow arrowheads); in contrast, MBON11’s synapses onto the MBON07s have a similar distribution as the input the MBON07s receive from KCs. (E) MBON03 (β′2mp) receives synaptic input from MBON11 shifted further from its dendritic root (hollow arrowhead) than the input it receives from KCs.

We also confirmed and extended the observation of Takemura et al. (2017) that MBON06 and MBON11 (γ1pedc>α/β) form axo-axonal connections in the α lobe in the same compartments where they make feedforward connections onto the dendrites of other MBONs (Figure 22-figure supplement 1); these axo-axonal connections add an additional layer of complexity to the MBON feedforward network.

### An extensive network of MBON to MBON connections outside the MB

MBONs make extensive contacts with one another outside the MB lobes, as summarized in diagrammatic form in Figure 23. Contacts between a typical and an atypical MBON or between two atypical MBONs can be axo-dendritic, whereas contacts between typical MBONs, with the exception of the three feedforward MBONs, are almost exclusively axo-axonal. Figure 23-figure supplement 1 shows all axo-axonal connections between typical MBON pairs that form at least 20 synapses. Such synapses are frequently observed between glutamatergic MBONs or from glutamatergic to cholinergic MBONs, as well as from GABAergic MBONs to glutamatergic MBONs, but not between cholinergic MBONs and GABAergic MBONs. We then examined where these synapses are located on the MBON’s axons. In several cases, axo-axonal synapses were highly localized and concentrated on a single axonal branch (Figure 23-figure supplement 2), suggesting that they might regulate synaptic transmission to only a subset of the postsynaptic MBON’s downstream targets. For example, all 110 synaptic connections from MBON09 (γ3β′1) to MBON01 (γ5β′2a) are confined to a small axonal branch of MBON01. We examined the morphology of these axo-axonal synapses and found them to be indistinguishable from axo-dendritic synapses made by the same MBON (Figure 23-figure supplement 3).

**Figure 23.**
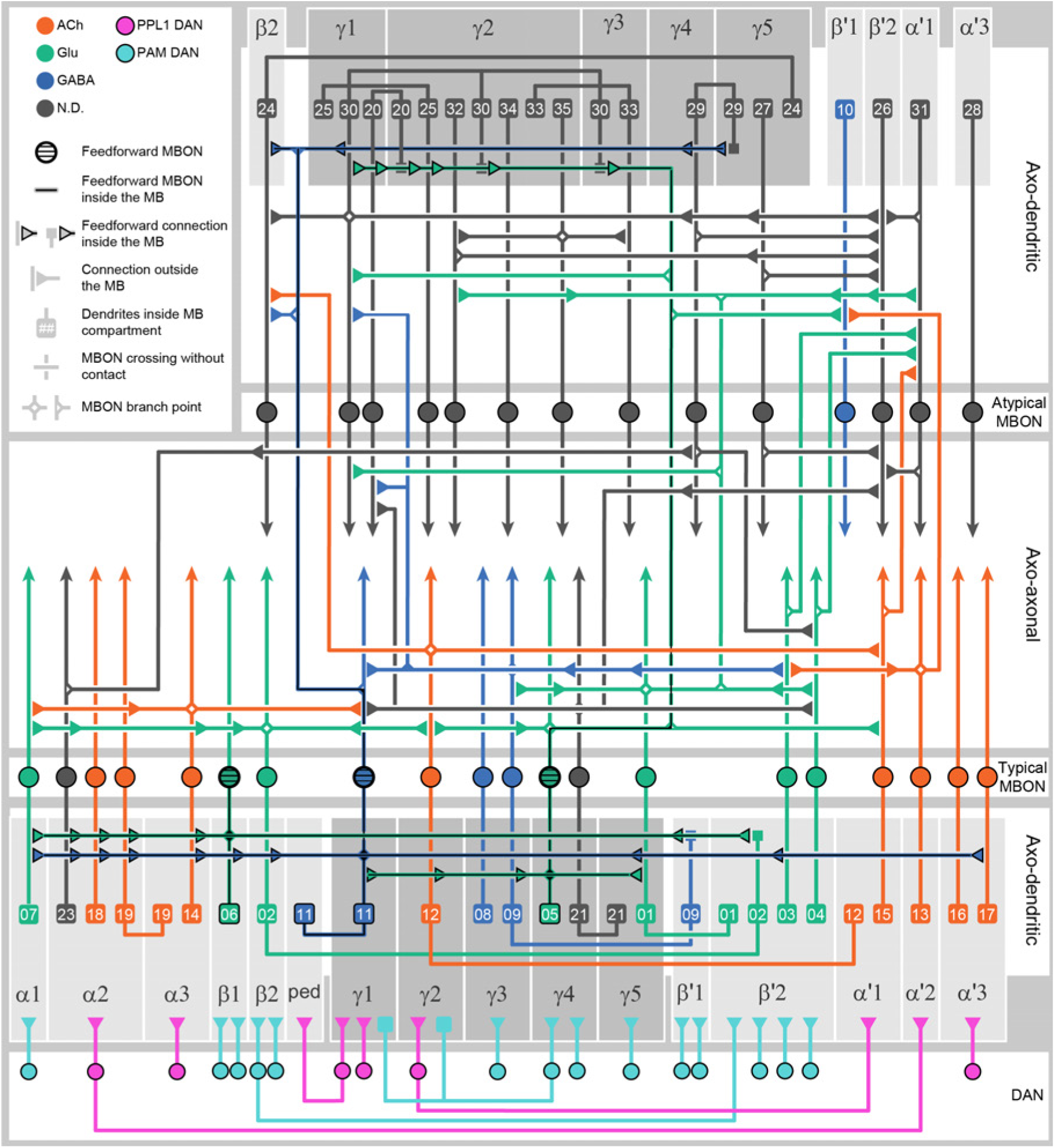
Diagram of the connections made between MBONs. A diagram showing MBON to MBON connections. At the bottom, grey boxes representing each of the MB compartments and the core of the distal pedunculus (pedc). DAN inputs, with PPL1 and PAM DANs color-coded, are shown below the compartments (see Figure 6-figure supplement 1 for the names of these cell types). MBON dendrites are represented as squares that contain the identification number of the MBON cell type, color-coded by neurotransmitter; see Figure 7 for more information about these MBONs. At the top, grey boxes representing the subset of MB compartments that house the dendrites of atypical MBONs are shown; the identification numbers of the MBON cell types are given in the squares. These MBONs are described in more detail in Figure 8. Outputs from MBONs are shown outside the grey boxes, with the exception of three MBONs—MBON05 (γ4>γ1γ2), MBON06 (β1>α) and MBON11 (γ1pedc>α/β)—indicated by the heavier outline and striped cell bodies, send axonal terminals (triangles) back into the MB lobes. These MBONs are described in more detail in Figure 22 and Videos 23, 24 and 25. Axo-dendritic and axo-axonal connections between MBONs are diagrammed separately, as indicated.

### Atypical MBONs form a multilayer feedforward network

Atypical MBONs receive significant input from both typical and atypical MBONs, forming a multilayer feedforward network that is diagrammed in Figure 24. Connections occur both inside and outside the MB lobes: panel A shows axo-dendritic connections outside the MB as well as both axo-dendritic and axo-axonal connections inside the MB, while panel B shows only axo-axonal connections outside the MB. This network contains three pairs of typical MBONs that form reciprocal connections: MBON06 (β1>α) with MBON11 (γ1pedc>α/β) through axo-axonal connections within the MB (Figure 24A), and MBON09 (γ3β′1) with both MBON03 (β′2mp) and MBON01 (γ5β′2a) through axo-axonal connections outside the MB (Figure 24B). Other than the loops formed by these MBONs, the network is exclusively feedforward. MBONs 1, 5, 6, 11 and 30 provide a significant fraction of the connections in this network. Although the neurotransmitter for MBON30 is currently unknown and the glutamatergic MBONs 1, 5 and 6 could have either an inhibitory or excitatory effect, it is possible that this network is dominated by inhibition. It is rare for atypical MBONs to make synapses onto typical MBONs and, when these occur, they are axo-axonal (Figure 24B). Only MBON29 and MBON30, non-LAL-innervating atypical MBONs with axons in the dorsal brain areas, make strong axo-axonal connections to typical MBONs, primarily to MBON04 (β′2mp_bilateral).

**Figure 24.**
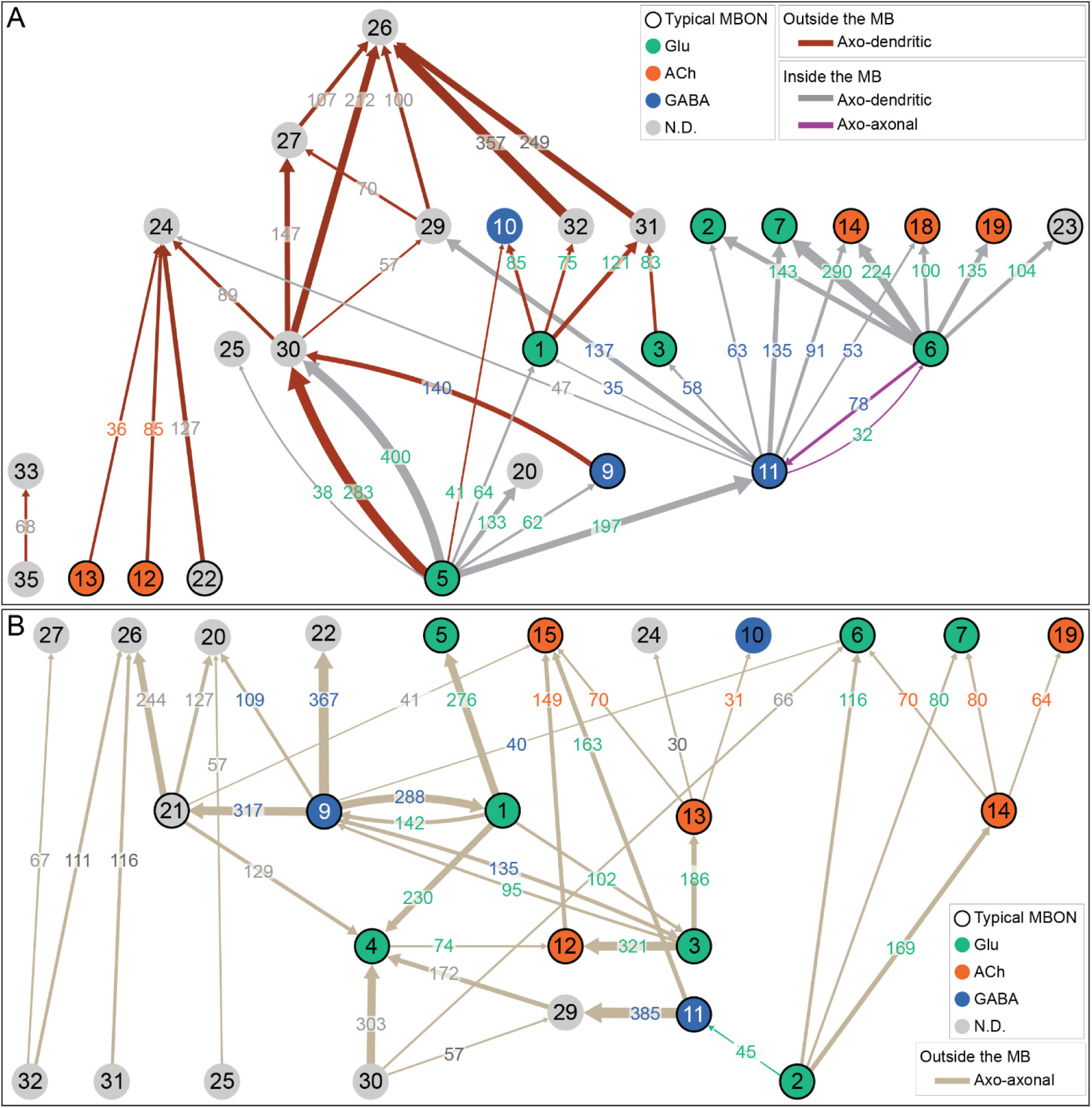
Connections between MBONs form a multi-layered feedforward network. Connections between MBONs, including both typical and atypical MBONs are shown, using a threshold of 30 synapses. The color of the arrow, as indicated in the color key, represents the nature and location of the synaptic connection. The number of synapses made in any given connection is indicated by the number in the arrow and the arrow thickness is proportional to that number. Numbers have been pooled for cases where there are two MBONs of the same cell type but only connections that occur in the right hemisphere are shown. The circles representing typical MBONs are colored according to their neurotransmitter and have a black border; atypical MBONs are shown without a border and in grey except for the atypical MBON10 that is GABAergic. (A) Axo-dendritic connections outside the MB lobes, axo-dendritic connections inside the MB lobes provided by feedforward MBONs (MBON05, MBON06 and MBON11), and the reciprocal axo-axonal connections between MBON6 and MBON11 within the MB lobes. (B) Axo-axonal synapses outside the MB lobes. Note that some MBONs, such as MBON26, form both axo-dendritic and axo-axonal connections outside the MB (see Figure 8-figure supplement 5E).

The network shown in Figure 24A is synaptically organized into 6 layers, with MBON26 (β′2d) forming the top or 6^th^ layer and MBON27 (γ5d), the 5^th^. Interestingly, of all the MBONs, these two are the most highly specialized for non-olfactory input; thermo-hygrosensory input in the case of MBON26 and visual for MBON27 (Figure 15; Figure 25-figure supplement 1). MBON26 and MBON27, as well as two of the neurons in layer 4, MBON31 (α′1) and MBON32 (γ2), send approximately 35, 60, 55 and 70 percent, respectively, of their synaptic output to the LAL, with MBON27 and MBON32 innervating the contralateral LAL. The numerous pathways by which information can reach these neurons, with modality-selective input that is subject to dopamine-modulated learning combined with both learned information carried by other MBONs and with non-MB inputs, reveals the potential for atypical MBONs to perform complex input integration. We address possible functional roles for such integration in the Discussion (Figure 38).

### Atypical MBONs provide a direct path to motor control

To drive changes in behaviour, such as approach or avoidance, the MB must ultimately communicate with motor neurons that lie in the ventral nerve cord (VNC). The brain is connected to the VNC by several hundred descending neurons (DNs) that pass through the neck to motor neuropils (Namiki et al., 2018). However, no direct MBON to DN connections have been reported, despite many DNs having dendrites in the same dorsal brain areas that serve as major MBON output sites (Figure 18; compare to Figure 4A of Namiki et al., 2018). Most DNs have their dendrites in more ventral brain regions; about a dozen lie in the LAL, a known CX (Hanesch et al., 1989; Heinze and Homberg, 2009; Lin et al., 2013; Namiki and Kanzaki, 2016; Wolff et al., 2015) and atypical MBON output site.

We report here the first identified direct neuronal paths from MBONs to DNs and thus to motor control. Four of the six LAL-innervating atypical MBONs are among the strongest inputs to the descending neuron (DN) DNa03, which in turn provides strong input to another DN, DNa02 (Figure 25). DNa02 also gets weaker, but direct, input from the ipsilateral MBON32 and from the MBON31s in both hemispheres. Asymmetrical activity in DNa02 has been strongly implicated in the steering of a fly’s walking direction (Rayshubskiy et al., 2020). The visual and thermo/hygrosensory sensory input these MBONs receive is predominantly ipsilateral, allowing them to convey directional sensory information that might then be used to drive approach or avoidance. The other two LAL-innervating MBONs, MBON33 and MBON35, also connect to neurons that we believe are DNs, but this needs to be confirmed by additional reconstruction.

**Figure 25.**
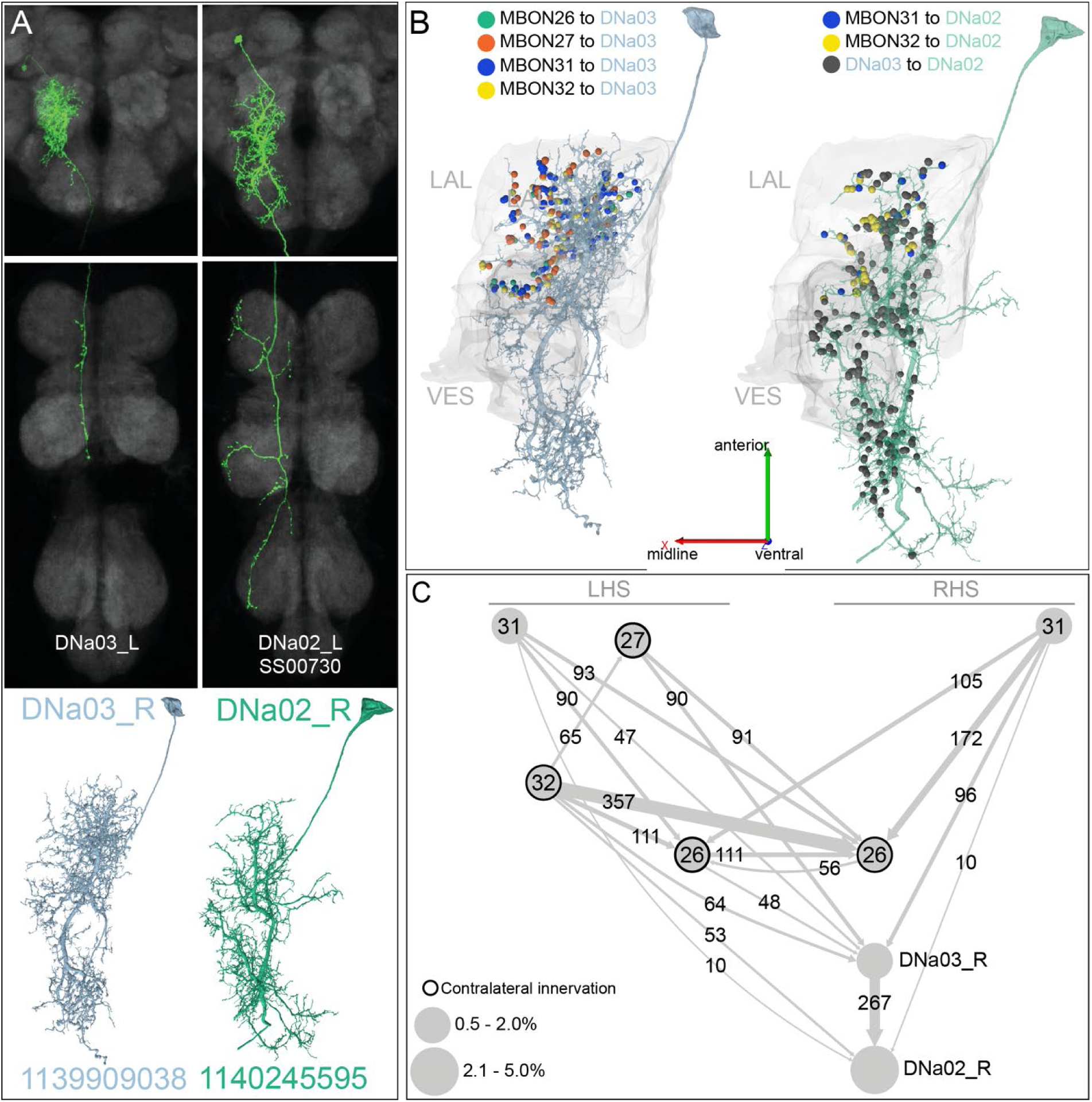
A descending pathway from atypical MBONs to the ventral nerve cord (VNC). Four LAL-innervating atypical MBONs form strong connections to descending neurons (DN). (A) A comparison between LM (Namiki et al., 2018) and EM morphologies of <u>DNa02</u> and <u>DNa03</u>. Note that the LM images show the cells in the left hemisphere while the EM images are from the right hemisphere and that the axons of the DNs extend outside the hemibrain volume. (B) Synaptic connections from MBON26, MBON27, MBON31 and MBON32 to DNa03 (left) and from MBON31, MBON32 and DNa03 to DNa02 (right) are shown, color coded. Note the connections from MBONs are restricted to the LAL while DNa03 makes synapses onto DNa02 throughout its arbor in the LAL and the VES. (C) Diagram of connectivity between MBONs, DNa03 and DNa02. Note that MBON26, MBON27 and MBON32 have contralateral axons (Figure 8-figure supplements 5, 6 and 11, Video 11, 12 and 17) and thus innervate the contralateral DNa02 and DNa03 while MBON31 has bilateral output (Figure 8-figure supplement 10, Video 16). Thus the connectivity diagram shows the inputs onto the DNa02 and DNa03 of the right hemisphere (RHS), while many of the neurons providing input have their cell bodies in the left hemisphere (LHS). Other inputs to DNa02 and DNa03 are summarized in Figure 25-figure supplement 1. DNa02 has been implicated in steering control (Rayshubskiy et al., 2020).

DNa03 appears to serve as a node for convergence of directional information from the optic lobes and the CX. DNa03 receives strong, direct visual input from multiple cells of a population of lobula VPNs (LT51 neurons; Figure 25-figure supplement 1 and Figure 25-figure supplement 2A and B). Unlike the VPN neurons that innervate the accessory calyces we described earlier, these LT neurons get input from layers of the lobula known to be involved in feature detection (see for example, Wu et al., 2016). Moreover, DNa03 gets strong input from a population of columnar FB neurons, PB1-4FB1,2,4,5LAL (PFL2; Figure 25-figure supplement 1 and Figure 25-figure supplement 2C and D) that are likely to convey orientation-sensitive information (discussed further in a companion manuscript in preparation on the CX connectome).

While six atypical MBONs innervate the LAL, the other eight have their arbors in more dorsal regions. We asked if any of these MBONs had identified DNs among their top 20 downstream targets. In this way, we were able to establish strong connections from MBON20 to two DNs, DNp42 and DNb05 (Figure 8-figure supplement 2). However, no identified DNs were found among the top downstream targets of any other atypical or typical MBON.

We do not mean to suggest that these are the only direct MBON to DN connections. At most half of the DNs expected from light level analyses (Namiki et al., 2018) have been identified in the hemibrain v1.1 dataset. We also identified weaker connections from atypical MBONs to other putative DNs as well as strong connections to LAL neurons that might turn out to be DNs upon more extensive analysis. However, it seems likely that the connections we describe here will remain among the strongest direct MBON to DN connections. Of course, all MBONs are likely to eventually drive DN activity through more indirect pathways, as it is through DNs that motor activity is largely controlled.

### Structure of DAN connectivity

DANs are divided into two major groups, PPL1 and PAM, that preferentially encode punishment and reward, respectively (reviewed in Modi et al., 2020). But there is growing evidence that DANs provide a wider range of information to the MB about novelty, locomotion, sleep state, reward or punishment omission, and safety (Aso and Rubin, 2016; Cohn et al., 2015; Dag et al., 2019; Felsenberg et al., 2018, 2017; Gerber et al., 2014; Handler et al., 2019; Hattori et al., 2017; Jacob and Waddell, 2020; Sitaraman et al., 2015b; Tanimoto et al., 2004). Understanding how DANs represent the external world and internal brain state requires a systematic investigation of their upstream inputs. Recent connectomic studies of DAN inputs in the larval brain (Eschbach et al., 2020a) and to a subset of DANs in the adult (Otto et al., 2020) revealed a surprising degree of heterogeneity. To understand what neuronal pathways contribute to the signals that DANs convey, as well as how these signals might produce learning-related changes in the strength of KC-to-MBON synapses, we characterized the inputs and outputs of DANs.

### Feedback from MBONs to DANs

We found extensive, direct synaptic connections between the axonal termini of MBONs and the dendrites of DANs (Figure 26). All six PPL1 DANs and 50 of the 150 PAM DANs receive direct MBON input; 20 of 22 typical MBON cell types, with representatives of all three neurotransmitter types, and seven of 14 atypical MBON cell types make direct connections to DANs. Connections were observed between MBONs and DANs that innervate the same compartment (Figure 26A), different compartments (Figure 26B) or both the same and different compartments (Figure 26C). Glutamatergic, and to a lesser extent, cholinergic, MBONs exhibit a strong bias toward providing feedback to DANs of the same compartment (self-feedback). A subset of PPL1 DANs was overrepresented among DANs receiving both within-compartment and cross-compartment connections (Figure 26C and D).

**Figure 26.**
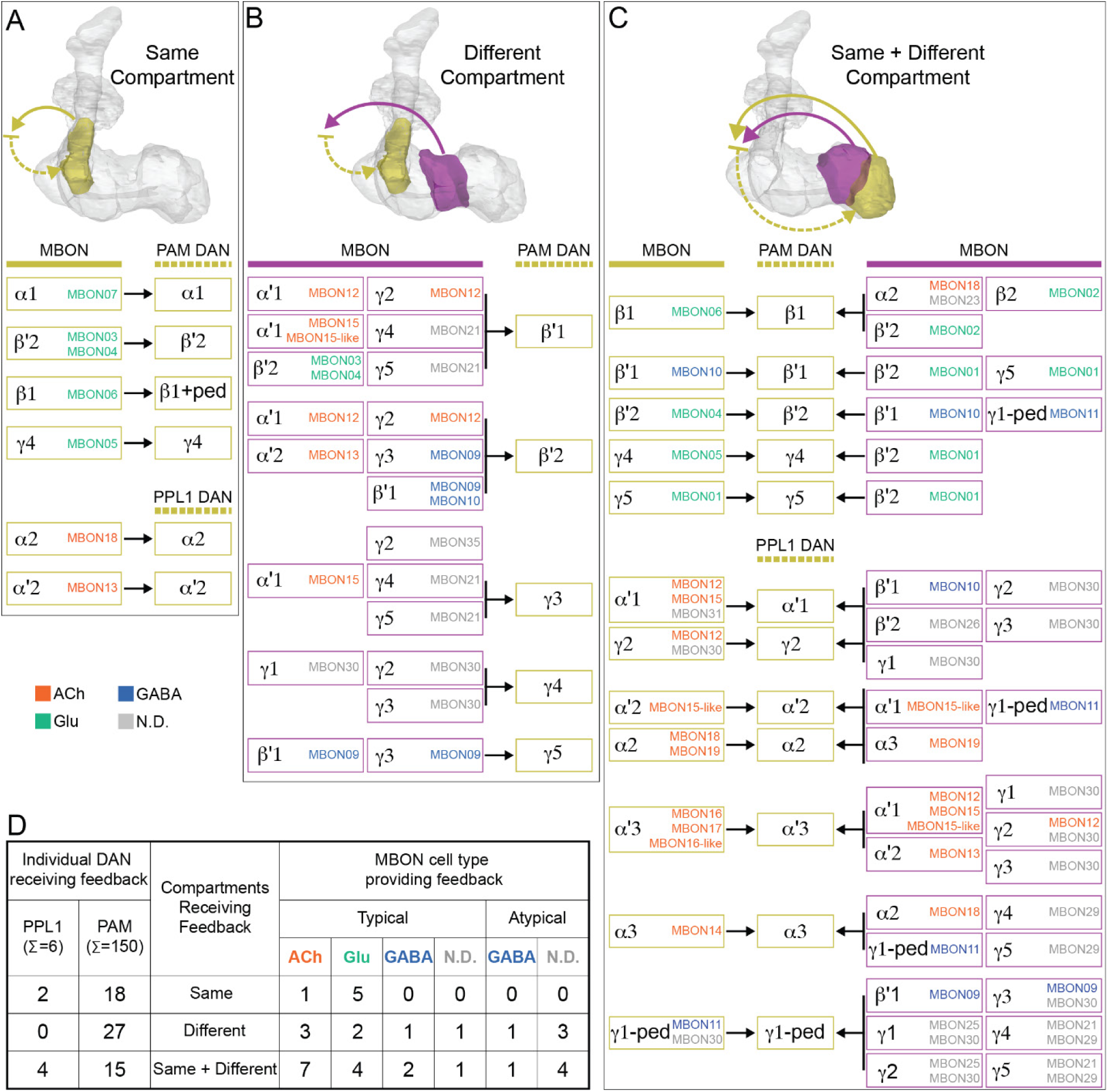
Feedback from MBONs to DANs. Input from MBONs onto DANs occurs in three distinct circuit motifs which are diagrammed. MBON connections to DANs from the same compartment are depicted as a yellow arrow, while MBON connections to DANs from a different compartment are depicted as a purple arrow. DANs downstream of these MBONs are depicted as dashed yellow arrows. (A) In the simplest case, the MBON and DAN form a reciprocal loop where the MBON innervates DAN(s) that send their axons back to the compartment where the MBON’s dendrites arborize. Below the diagram of this circuit motif, each of the instances we observed with this arrangement is presented; inputs onto PAM and PPL1 DANs are presented separately. The color of the MBON name indicates its neurotransmitter, when known. (B) In this case, MBON(s) synapse onto DAN(s) that send their axons to different MB compartment(s), providing a mechanism for cross-compartment communication. It is common for DANs to receive input from several MBONs. (C) This motif combines the previous two motifs, with DANs receiving input from both MBON(s) of same and different compartments. (D) The table summarizes DAN populations that participate in the three different feedback motifs. Note that all 6 PPL1 DANs receive MBON input, but only a subset of the 151 PAM DANs receives MBON input.

Prior studies have suggested a variety of roles for feedback from MBONs to the DANs that innervate the same compartment and modulate them. For example, the nutrient dependent consolidation of appetitive long-term memory requires MBON07 (α1) activation of PAM11 (α1) DANs (Ichinose et al., 2015) and inhibition from MBON 11 (γ1pedc>αβ) onto the PPL101-γ1pedc DANs (Pavlowsky et al., 2018) after training. The MBON 11 (γ1pedc>αβ) to PPL101-γ1pedc DAN connection has also been linked to the persistence of hunger-dependent food odor seeking (Sayin et al., 2019). In addition, feedback activation of PAM01 (γ5) by MBON01 (γ5β′2a) is required for the maintenance of short-term courtship memory (Zhao et al., 2018b) and for aversive memory extinction (Felsenberg et al., 2018; Otto et al., 2020).

In addition to direct connections, we also computed the effective connection strength from MBONs to DANs mediated by an interneuron (Figure 26-figure supplement 1) and the proportion of this feedback where an MBON feeds back onto a DAN that innervates the same compartment (Figure 26-figure supplement 2). Interestingly, the same-compartment bias of glutamatergic and, again to a lesser extent cholinergic, MBONs seen for direct connections is also present for these indirect connections. The presence of this motif in both direct and indirect MBON-DAN interactions is intriguing, and we consider its potential computational significance in the Discussion.

### Inputs to individual DANs

We characterized the inputs to the dendrites of DANs by computing cosine similarity matrices of these inputs for all pairs of the 157 DANs that innervate the MB (Figure 27). An enlarged view of PPL neurons is shown in Figure 27-figure supplement 1 and a simplified matrix, in which the data is collapsed by compartment is shown in Figure 27-figure supplement 2. Structure at the level of lobes, compartments, and even within compartments is evident. The heterogeneity of input to different DANs is also clearly visible. In some cases, DANs receive prominent input from particular brain areas (Figure 27-figure supplement 3, left panel). For instance, α′3 and α′2α2 DANs receive more than half their input from the SIP, γ3 and γ4<γ1γ2 DANs from the CRE, and γ5β′2a, γ5, γ1pedc, β′2a, β′2m, β′2p, and α3 DANs from the SMP. Tracing DAN inputs back to include those inputs mediated by an interneuron reveals even more complex and broadly distributed inputs (Figure 27-figure supplement 3, right panel), with some distinct differences from direct inputs such as vastly increased input from the CA. In general, these input correlations suggest a heterogeneity in DAN function that could in principle support more than a dozen different learning modalities. We identified 41 putative functional groups of DANs, and an analysis of the singular values of the DAN input connectivity matrix suggests at least 20 sub-compartmental zones within the MB lobes where distinct modulation by DANs might occur.

**Figure 27.**
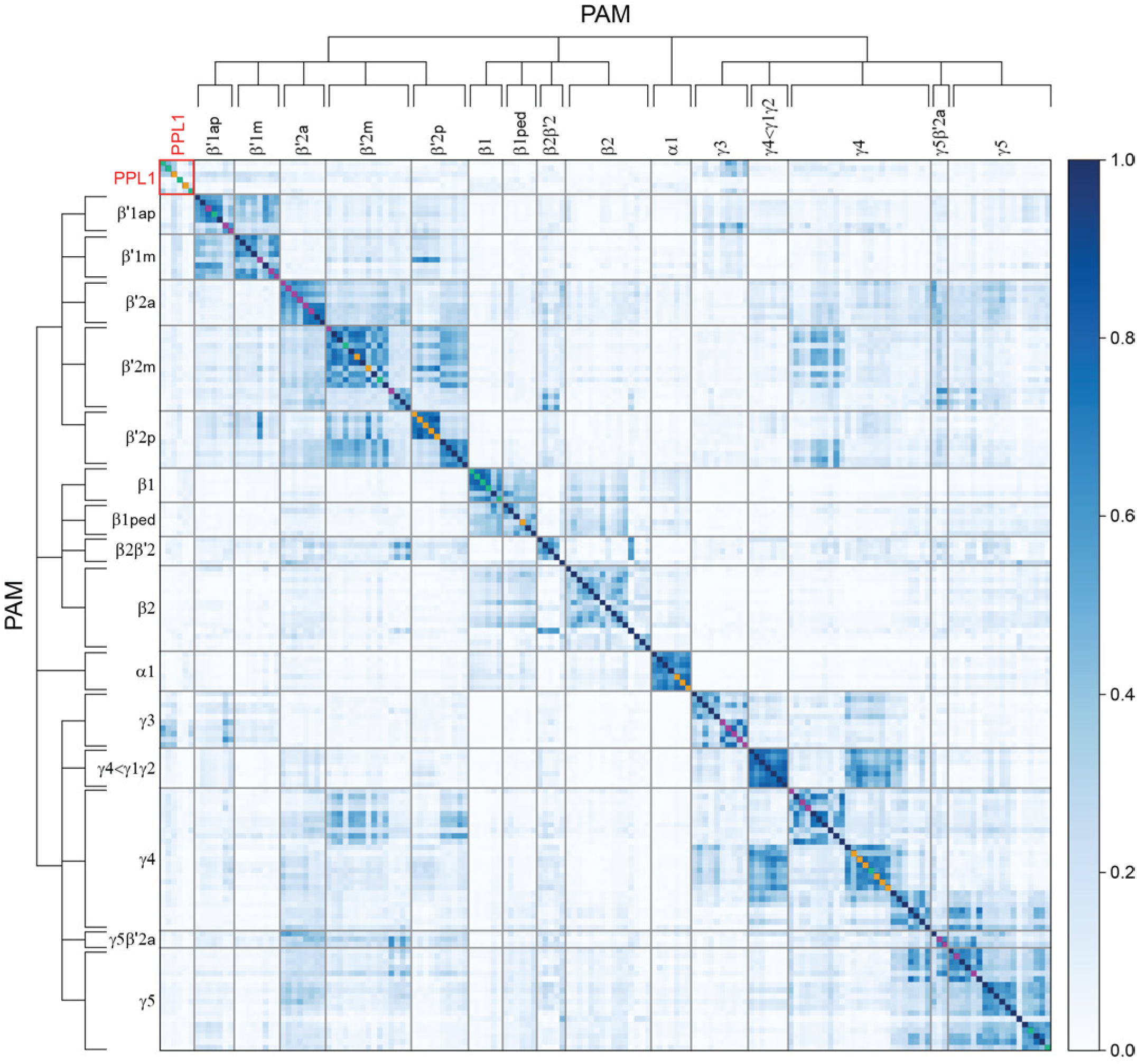
Similarity of input to individual DANs. Heatmap representing the similarity of inputs received by DANs. Each square represents the cosine similarity of inputs received by each PAM or PPL1 DAN. In order to focus on inputs received outside the lobes of the MB, inputs from KCs, other DANs, APL and DPM neurons have been excluded. DANs are grouped by type, and the ordering within each type is determined by spectral clustering. Colors on the diagonal indicate whether the given DAN receives feedback from MBONs in the same compartment (yellow), MBONs from different compartments (purple), both (green), or neither (dark blue). Figure 27-figure supplement 1 shows an expanded view of the PPL1 DAN portion of the heatmap. Figure 27-figure supplement 2 shows the average input similarity between DAN cell types computed after pooling the data for all cells of a given DAN cell type.

DAN inputs need to be organized so they can convey learning-related signals to the appropriate KC-MBON synapses within the MB. Such an organization should be reflected in the relationship between the structure of DAN inputs and MBON outputs. For each pair of compartments, we computed the similarity of the upstream inputs received by DANs innervating the two compartments and the corresponding similarity of the downstream targets of MBONs from those compartments (Figure 27-figure supplement 4). We identified a significant positive correlation between these two similarity measures, suggesting a form of “credit assignment” in which compartments whose MBONs control similar behaviors are innervated by DANs that receive similar reinforcement signals. This correlation suggests that the observed heterogeneity of DAN inputs is functionally relevant.

### PAM DANs cluster into morphological subtypes that are reflected in upstream connectivity

Grouping DANs into subtypes using hierarchical clustering based on either of two different criteria, morphology or upstream connectivity, gave strikingly similar results. Figure 28 shows such comparisons of morphology- and connectivity-based clusters for PAM01 (γ5), PAM12 (γ3) and PAM11 (α1) DANs. For α1, morphology and input clustering generate an identical dendrogram, whereas for γ5 and γ3 the major groupings are identical but some of the “within group” orders are shifted. Finding that morphology clustering reflects input clustering demonstrates that morphology is a good indicator of functionally-relevant DAN subtypes, as previously demonstrated for γ5 DANs (Otto et al., 2020).

**Figure 28.**
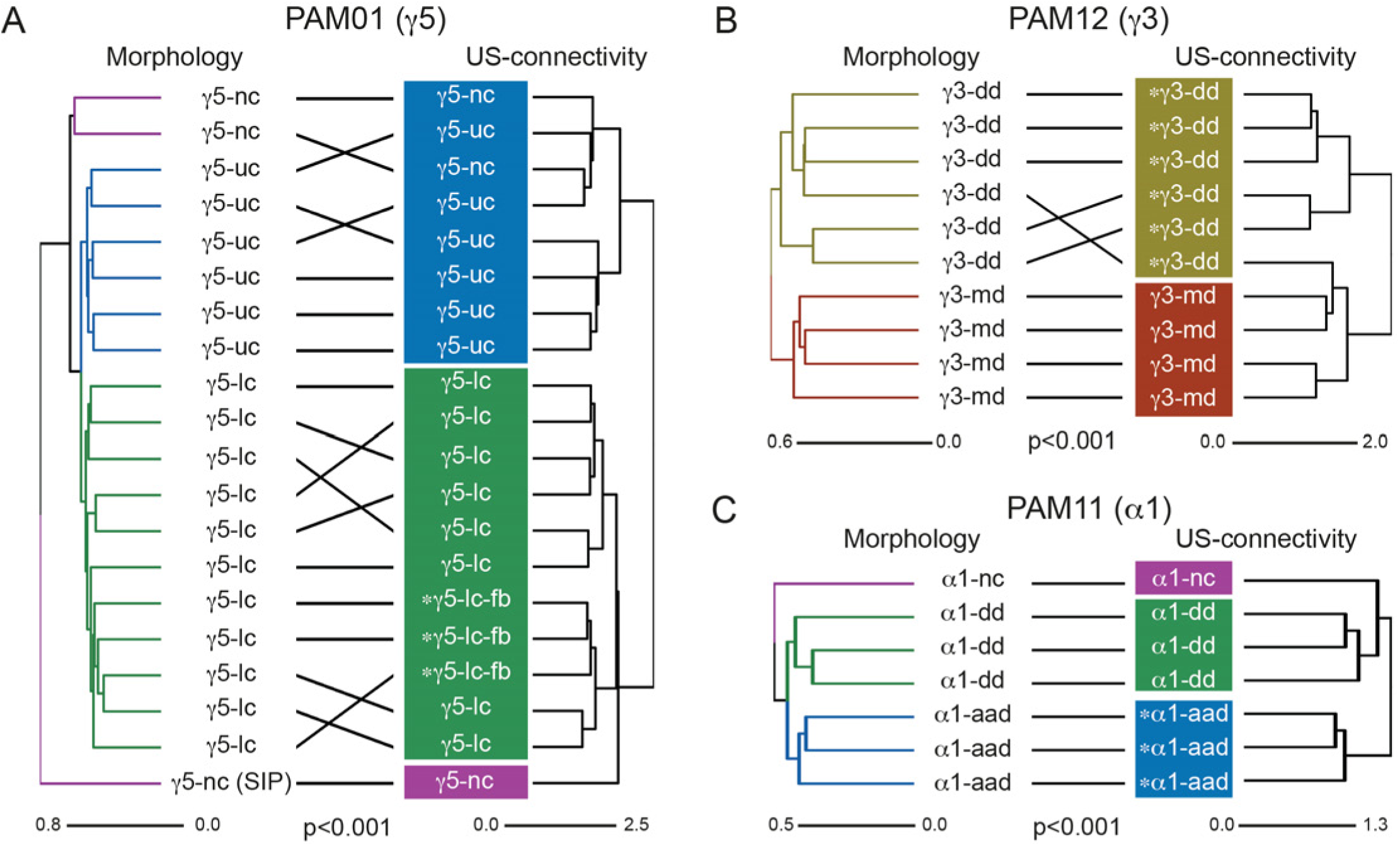
Morphologically defined DAN subtypes receive similar input Tanglegrams show that DAN subtyping based on hierarchical morphological clustering matches that generated by hierarchical clustering by input connectivity (compare this figure to DAN input similarity shown in Figure 27). We show data here for three compartments; we found that clustering by morphology or input connectivity gave similar results in all PAM DAN compartments (not shown). (A-C) Tanglegrams for right hemisphere PAM01 (γ5), PAM12 (γ3), and PAM11 (α1) DANs. The left hand dendrograms represent hierarchical clustering by morphology and colors mark neurons clustered together (morphologies shown in Figure 28-figure supplement 1 and Figure 30). Right hand dendrograms represent hierarchical clustering by dendritic input connectivity. Colored boxes mark DAN subtype clusters defined by input similarity (compare to Figure 27). Asterisks denote neurons that receive MBON feedback, as discussed in Figure 26 and 27. (A) PAM01 DANs cluster into subtypes. Two morphological types are categorized by their contralateral commissure tract: upper commissure (uc) and lower commissure (lc) (Figure 28-figure supplement 1 A, B). Three neurons within the lower commissure subtype receive MBON01 (γ5β’2a) feedback (asterisk) and have distinct connectivity and axon morphology features (Figure 28-figure supplement 1D). Two neurons within the first morphological subtype are non-canonical (nc) with significantly different dendritic or axonal branching patterns (Figure 28-figure supplement 1C). Subtypes are consistent with those described in Otto et al. 2020. Connectivity clustering produces three subtypes by grouping the two non-canonical neurons with the uc subtype. Connectivity and morphology clustering are not significantly independent (Mantel test: r=0.764, p< 10^-7^, n=10^7^). (B) PAM12 DANs cluster into two subtypes mostly defined by their dendritic field anatomy: dorsal dendrite (dd), and medial dendrite (md) (Figure 28-figure supplement 1 E-F). Clustering by input connectivity also produces two subtypes. Connectivity and morphology clustering are not statistically independent (Mantel test: r=0.838, p=2.76×10^-7^, n=3.6×10^6^). (C) PAM11 DANs cluster into three morphological subtypes. Two are distinguishable by their dendritic anatomy: dorsal dendrite (dd) and those with an additional anterior dendrite (aad). The third subtype is a single non-canonical α1 DAN, which has a restricted axonal field (Figure 28-figure supplement 1 G-I). Morphological clustering perfectly matches input clustering, and neither is statistically independent (Mantel test: r=0.779, p=3.97×10^-4^, n=5×10^4^).

Figure 28-figure supplement 1 illustrates the use of morphology to divide PAM01 (γ5), PAM12 (γ3) and PAM11 (α1) DANs into subtypes. The most distinguishing morphological features are the locations of dendritic and axonal fields, and the commissure in which a contralaterally projecting axon crosses the midline. Importantly, our clustering results—using cosine input similarity (Figures 27 and 29), input similarity and morphology (Figure 28)—all support the division of PAM cell types into subtypes in every compartment that is innervated by a population of DANs. As discussed in the next section, this sub-compartmental structure is likely to have important functional consequences.

**Figure 29.**
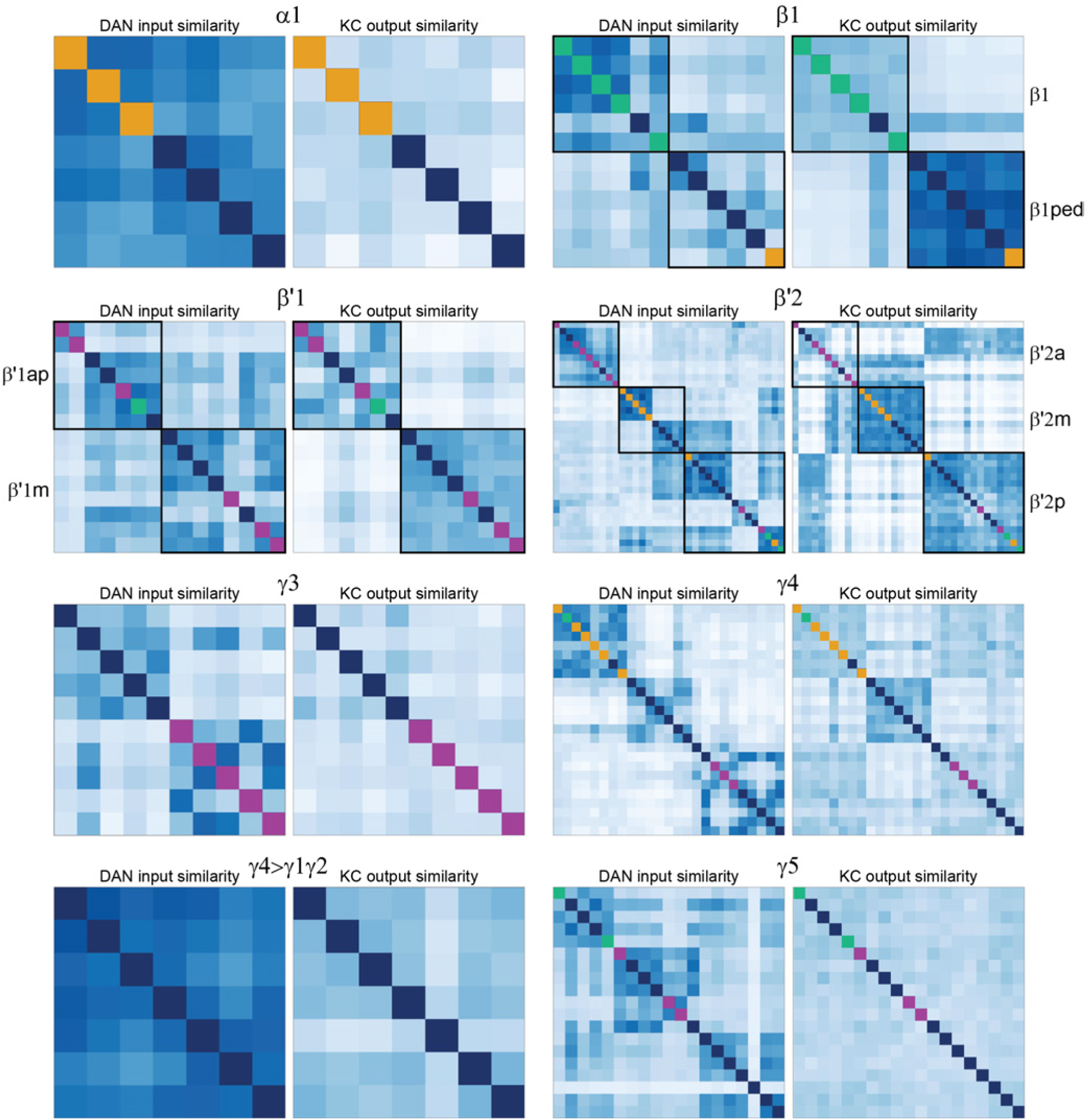
Relationship between DAN dendritic inputs and DAN axonal outputs to KCs. These plots explore whether DANs of the same cell type that have distinct inputs also connect to distinct populations of KCs within a compartment. Analysis of eight compartments is shown. Within each example, the left heatmap represents the similarity of input to DANs (identical to the corresponding block in Figure 27). The right heatmap represents the cosine similarity of the output synapses made by DANs onto KCs. The clustering observed in the right panels most likely results from individual PAM neurons only arborizing in a portion of their compartment (see neuronal morphologies Figure 28-figure supplement 1 and Figure 30). Such clustering may permit specific reinforcement signals to be conveyed to subsets of KCs within a compartment. See detailed discussion of DAN-KC connectivity in Figure 30 and Figure 30-figure supplements 1 and 2. The potential computational significance of this phenomenon is explored in Figure 38.

### PAM DAN subtypes selectively modulate subsets of KCs and specific MBONs within single compartments

We reasoned that DAN subtypes might specifically connect to particular types of KCs and MBONs within a single compartment. Previous work suggested that the major axonal arbors of γ5 DAN subtypes innervate distinct subregions within the compartment (Otto et al., 2020).

The dense anatomical reconstruction and connectivity of neurons in the hemibrain data set allowed us to examine whether such differences in morphology are reflected in connectivity and, if so, whether this is a general feature of all MB compartments that are innervated by multiple DANs. We first clustered DANs within a compartment based on the similarity of their dendritic inputs and then, without changing the order determined by that clustering, asked if we could observe structure in the pattern of their outputs onto KCs (Figure 29). In some compartments, for example β1, β′1 and γ4, such conserved structure was indeed observed indicating that DANs with similar inputs also output onto similar sets of KCs. In other compartments known to have clear subtypes based on their dendritic inputs and morphology, such as α1, γ3 and γ5, conserved structure was not identified by this analysis. Therefore, we investigated the DANs in these three compartments in more detail, as described below, and found clear examples of subtype specificity in their synapses onto KC cell types. We also found selectivity in DAN subtype synapses onto MBONs within compartments innervated by multiple MBON cell types, including the newly discovered atypical MBONs.

Each of the four defined PAM01 (γ5) DAN subtypes is differentially connected to distinct KC populations within the γ5 compartment. This is most evident for PAM01-nc and PAM01-fb DANs whose presynaptic arbors occupy distinct regions of the compartment (Figure 30A and B) where they contact KC populations representing different stimulus modalities (Figure 30D and E; Figure 30-figure supplements 1 and 2; Figure 15). For example, PAM01-nc DANs are the major input to the visual streams of γd and γs1 KCs (Figure 30D). In contrast, the PAM01-fb DANs synapse predominantly with the main olfactory KCγ population, and weakly with the γt and γs1 KCs (Figure 30E). This arrangement suggests that DAN modulation is specific to particular sensory modalities.

**Figure 30.**
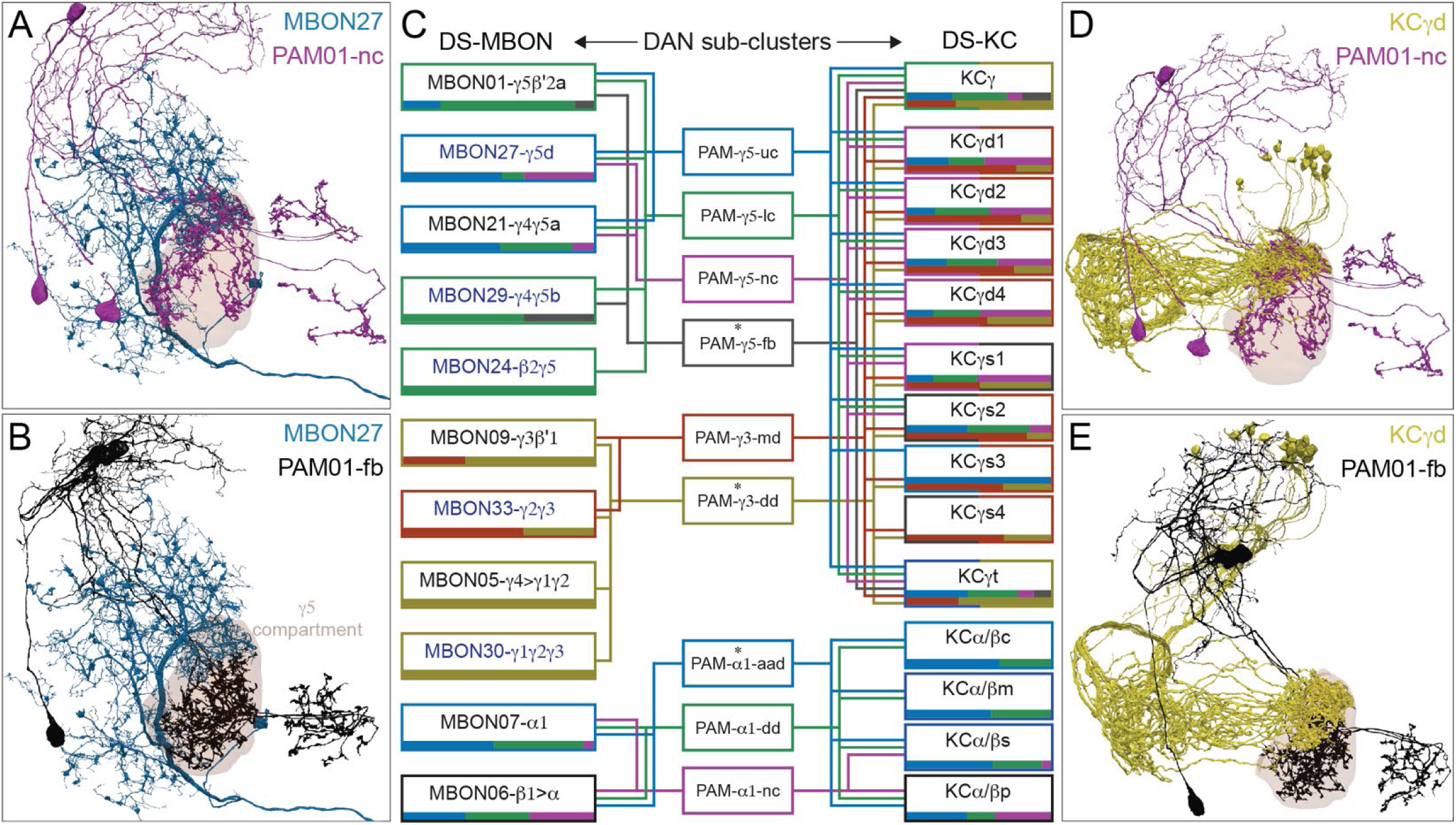
Different subtypes of the PAM01, PAM12, and PAM11 DANs synapse onto specific subpopulations of KCs and MBONs within their compartment. The axons from certain subtypes of DAN occupy different regions of, and together tile, a MB compartment. This permits for specific connectivity to KCs and MBONs within the compartment. A zonal subcompartment arrangement is a feature of many MB compartments and is illustrated here by the distinct morphologies of two subtypes of the γ5 compartment DANs (A, B, D, E). Implications for connectivity of this type of restricted DAN arborization are summarized for the γ5, γ3, and α1 compartments in panel C. Figure 30-figure supplements 2 and 3 show connectivity matrices for additional compartments in the right hemisphere. (A) The axons from PAM01-nc (magenta) overlap with dendrites of MBON27 (γ5d; dark blue). Note that the axons of the two PAM01-nc DANs (magenta) are confined to the dorsal and lateral region of the γ5 compartment (see also Figure 28-figure supplement 1C). (B) PAM01-fb (black) axons do not overlap with the dendrites of MBON27 (dark blue). (C) Schematic of the innervation of subtypes of γ5, γ3, and α1 DANs within the compartment showing their connection to MBONs (left) and KCs (right). Boxes for DAN subtypes and connectivity edges are colored according to subtype identity. Boxes around MBON types are colored by their strongest DAN subtype input. The boxes around KC names are colored to reflect their strongest DAN subtype input in each compartment (the two halves for KCγ types reflect their projections through the γ3 and γ5 compartments). Horizontal bars along the lower edge of the MBON and KC boxes are histograms, with bar colors corresponding to DAN subtypes, representing the proportion of each DAN subtype providing input to the respective MBON or KC subtype. For the KCγ types, the upper bar represents inputs in γ5 and the lower bar in γ3. Typical MBON names in black and atypical MBONs are in blue. Asterisks denote DAN subtypes receiving MBON feedback (compare with Figures 26, 27 and 28). Only cases where a DAN provides at least 0.1% of the MBON’s synaptic input or at least 1% of a KC type’s input from DANs are shown. KC cell types are biased in their representation of different sensory modalities (Figure 15-figure supplement 2). Note that the synapses onto the feedforward MBON06 (β1>α) are likely axo-axonal. (D) The axons from PAM01-nc (magenta) also overlap with the axonal processes of KCγd (yellow). (E) Axons from PAM01-fb (black) do not overlap with the axonal processes of KCγd (yellow).

DANs are known to make synapses onto the dendrites of MBONs (Takemura et al., 2017), but the physiological and behavioral roles of DAN to MBON connections are much less well understood than those of DAN to KC connections; in the one case where there is functional data, DAN activation induced a slow depolarization in the postsynaptic MBON (Takemura et al., 2017). We found that DANs also showed selectivity within individual compartments in their targeting of MBONs. For example, in γ5 the PAM01-fb DANs synapse onto MBON01 (γ5β’2a) and MBON29 (γ4γ5), but not MBON27 (γ5d), whose dendritic field preferentially occupies the dorsal region of the compartment. MBON27 nevertheless receives synapses from the other three classes of PAM01 DANs. Other MBONs in γ5 also receive preferential input from specific PAM01 DAN subtypes; for example, MBON24 (β2γ5) only receives synapses from the PAM01-lc DANs.

A similar but less complex subtype arrangement of DAN-KC and DAN-MBON connectivity is evident in the γ3 compartment, where there are two morphological DAN subtypes (Figure 28-figure supplement 1E and F; Figure 39). The PAM12-md DANs provide most of the DAN input to the γd KCs, whereas the γm and γt KCs are preferentially innervated by the PAM12-dd DANs (Figures 30E and 32). The three MBONs with processes in the γ3 compartment exhibit very different connectivity from γ3 DAN subtypes. PAM12-dd DANs provide all of the DAN input to MBON30 (γ1γ2γ3) and the majority of MBON09 (γ3β′1)’s input, whereas PAM12-md DANs provide the majority of MBON33 (γ2γ3)’s input.

**Figure 31.**
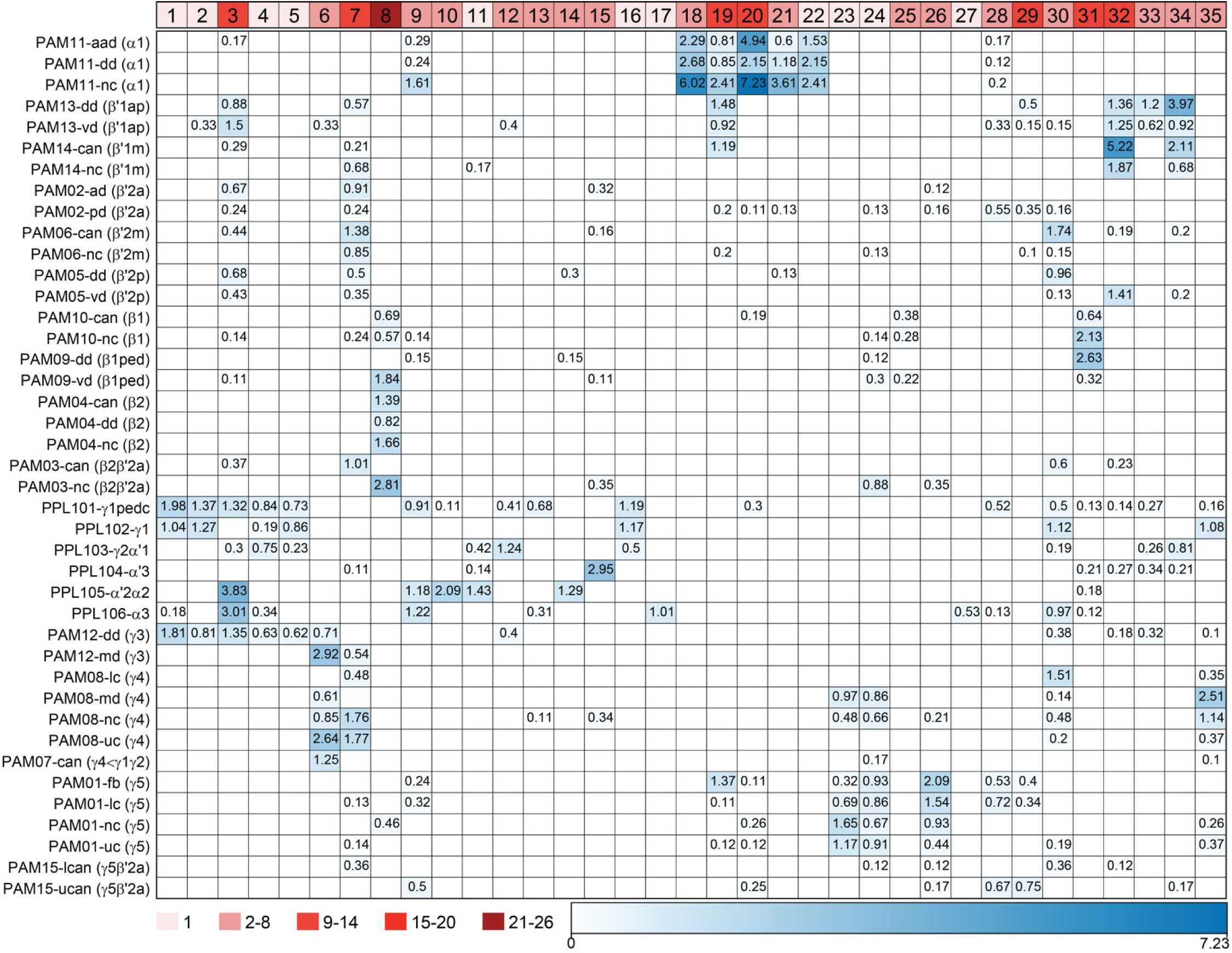
Shared input neurons reveal expected and unexpected across-compartment groupings of DAN subtypes. Matrix representing the connectivity to DAN subtypes provided by the top 50 strongest input neurons to each DAN subtype in the right hemisphere (n = 913 neurons, excluding MBONs and MB intrinsic neurons). These inputs are either unique individual neurons (clusters of one) or clusters of multiple neurons grouped according to their morphology. A selection of 35 input clusters are shown, numbered in boxes whose shading reflects the number of neurons that contribute to that cluster (all input clusters are listed in supplemental Table 2). Each cell represents the proportion of the total inputs to the dendrites of the DANs of the respective subtype (y-axis) provided by the input neuron cluster (x-axis). Selected input clusters and innervation patterns are further explored in Figures 32-37. Some DAN subtypes receive inputs from neurons projecting from heavily studied neuropils like the SEZ, the FB, and the LH; for example, clusters 10, 18, 19, 20, 21, 22, 23, 24, 25, 26, 27, 28, 31 (Figures 33-35). Many clusters selectively connect to groups of DAN subtypes of the PAM or PPL1 type specifically; for example, clusters 7, 8, 11, 12, 13, 14, 15, 29, 32 (Figure 36) and clusters 22, 23, 24, 26, 27 (Figure 35). Some input neuron clusters connect DAN subtypes with opposite/mixed valence; for example, clusters 3, 9, 30, 33, 35 (Figure 37) and cluster 28 (Figure 35; also see Otto et al. 2020). Input neuron clusters can also stand out by the DAN subtypes that they avoid; for example, clusters 3 and 30 (Figure 37). The matrix was thresholded to show inputs over 0.1% of total input to DAN dendrites and the threshold for strong connectivity was defined as 0.5%. The neuropil of the SEZ is not included in the hemibrain database. Therefore, more complete anatomy of DAN input neurons from this region were retrieved from FAFB (Zheng et al. 2018; see Figure 35).

**Figure 32.**
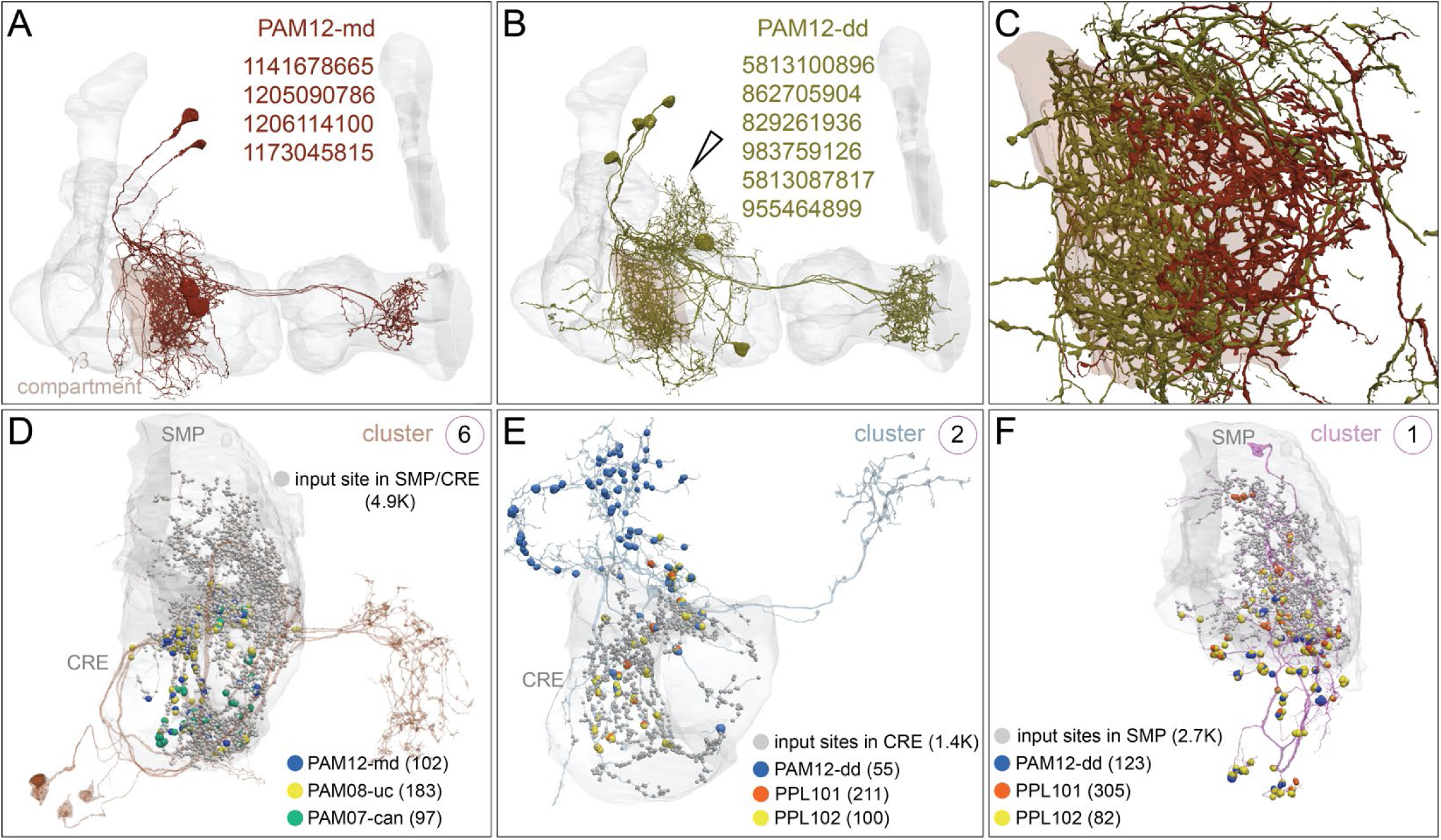
Morphologically distinct subtypes of PAM12-DANs share inputs with either PPL1 or PAM DANs. (A) The four PAM12-md DANs (maroon) innervate the γ3 compartment (brown) of the MB neuropil (grey). (B) The PAM12-dd subtype (green) has additional dorsal dendritic branches (open arrowhead) not found in PAM12-md. A seventh PAM12-dd that lacked its ipsilateral axon was excluded from the analysis. (C) The axonal fields of PAM12-md and PAM12-dd DAN subtypes tile the γ3 compartment (brown), suggesting they modulate different downstream neuronal connections (see Figure 30 for further analyses). (D-F) Strong inputs (see Figure 31) to PAM12-md also specifically synapse onto positive valence PAM DANs, whereas PAM12-dd shares inputs with negative valence PPL1 DANs; synapse numbers are given in parentheses. Circled number in the upper right of each panel refers to the input cluster number in Figure 31. (D) <u>Cluster 6</u> collectively provides the strongest dendritic input to PAM12-md DANs (blue dots) and is also the strongest input to two DAN subtypes innervating γ4, PAM08-uc DANs, (yellow dots) and canonical (can) PAM07-can (green dots). In addition, cluster 6 connects to other PAM08-DANs, but excludes the lower commissure (lc) subtype. (E) A single, morphologically distinct neuron (<u>cluster 2</u>) connects to the positive valence PAM12-dd (blue dots) and the negative valence PPL101 (γ1pedc) (orange dots) and PPL102 (γ1; yellow dots). (F) This single (<u>cluster 1</u>) neuron provides the strongest dendritic input to positive valence PAM12-dd (blue dots) and the negative valence PPL101 (γ1pedc; orange dots) and also synapses onto PPL102 (γ1; yellow dots) DANs. Connections contributing less than 1% of a neuron’s total dendritic input have been excluded.

The three α1 DAN subtypes (Figure 28-figure supplement 1 G-I) also exhibit preferential KC wiring (Figure 30). For example, PAM11-nc arbors only occupy a limited portion of the α1 compartment (Figure 28-figure supplement 1I) where they preferentially innervate the visual stream α/βp KCs, while avoiding α/βc and α/βm KCs. The α1 compartment houses the dendrites of only a single MBON type, whose two members each fill the compartment (Video 2) and therefore receive input from all three PAM11 subtypes.

DAN subtype-specific wiring is also evident in the α′/β′ lobes (Figure 30-figure supplement 2). As an example, consider input to the thermo/hygrosensory α′/β′ap1 KCs. In the β′2a compartment, only PAM02-pd (β′2a) provides input to these KCs. Similarly, in β′2p, PAM05 DANs make abundant synapses onto the α′/β′ap1 KCs, but do not contact the α′/β′m KCs. Previous work has implicated β′2 DANs in thirst-dependent naive and learned water-seeking (Lin et al., 2014b; Senapati et al., 2019) and the pattern of connectivity described above suggests that these roles are likely to be served by specific DAN subtypes modulating specific streams of KCs.

Our analyses suggest that selective DAN subtype innervation of modality-specific streams of KCs is a general feature of all MB compartments in the horizontal lobe (Figure 30-figure supplement 1). Furthermore, every MB compartment that is innervated by multiple DANs contains more than one DAN subtype and generally one DAN subtype in each compartment receives direct MBON input, whereas the other subtypes do not. With DAN subtypes synapsing onto selective KCs and MBONs, this allows different populations of synaptic weights to be independently adjusted within each compartment. This arrangement is likely to provide significantly more computational bandwidth to each compartment in the MB network. We further explore its function in the Discussion (Figure 38C-D).

### Shared input neurons suggest functional grouping of DAN subtypes across compartments

Clustering DANs based on input similarity revealed groups of DANs innervating different compartments that had a significant fraction of their upstream neurons in common (Figure 27). To explore the neuronal pathways that provide such shared input, we selected the 50 neurons that make the largest number of synapses onto each DAN subtype. We subjected this population of 913 neurons to hierarchical clustering based on morphology, which generated 235 input neuron clusters. We then analyzed how these input neuron clusters connect to the different DAN subtypes. Interestingly, several of the input clusters contained only a single neuron. Figure 31 shows 35 of these clusters, selected to illustrate the range of connectivity patterns; supplemental Table 2 shows data for all 235 clusters. In the following paragraphs we highlight some of our key findings.

Studies of aversive olfactory learning have established that key teaching signals are provided by the PPL101 (γ1pedc), PPL103 (γ2α′1) and PAM12 (γ3) DANs. Individually blocking output from these DANs impairs aversive memory formation and their forced activation can assign aversive valence to odors (Aso et al, 2010; Aso et al., 2012; Hige et al., 2015; Perisse et al; 2016; Yamagata et al., 2016; Jacob and Waddell, 2020). Our connectomics data shows that the PPL101 (γ1pedc), PPL103 (γ2α′1) and PAM12 (γ3) DANs share input, which supports the idea that they are driven in parallel in response to aversive/punishing cues. Several of the shared input pathways correspond to individual neurons that make synapses onto all of these aversively reinforcing DANs. These inputs provide the clearest evidence to date that this collection of aversively reinforcing DANs can be triggered together (assuming that at least some of the inputs are excitatory). Sometimes PPL101 (γ1pedc) input neurons also connect to other DANs, which again provides insight into the functional relevance of recruiting ensembles of DANs. For example, the individual PPL101(γ1pedc) input neurons in clusters 1 and 4 and the group in cluster 13 provide weaker input to the PPL106 (α3) DAN. When stimulated alone during odor exposure, the PPL106 (α3) DAN requires more trials to code aversive learning than PPL101 (γ1pedc), consistent with it being relatively inefficient (Aso and Rubin, 2016). However, multiple spaced aversive training trials strongly depress the conditioned odor-response of MBON14 (α3) (Jacob and Waddell, 2020). We also found that PPL101 (γ1pedc) frequently shares input with PPL102 (γ1). Although, these DANs have overlapping innervation in the γ1 compartment, to date most attention has been focused on the PPL101 (γ1pedc) DAN. It will therefore be important to determine the role of PPL102 (γ1).

Whereas each aversively reinforcing PPL1 DAN is a single neuron, there are 10-11 PAM12 DANs innervating γ3 (one PAM12 appears to be incomplete in this data set). An unexpected finding from this study is that the PAM12 (γ3) DAN population is comprised of at least two clear subtypes that are notable for the DANs with which they share input. Moreover, analysis of the inputs to the two PAM12 (γ3) DAN subtypes provides evidence that they convey opposite valence. Four single neuron clusters (clusters 1, 2, 4 & 5) preferentially connect to PAM12-dd (γ3) DANs and the negative valence PPL1 DANs, PPL101 (γ1pedc) and PPL102 (γ1), implying that PAM12-dd might also signal negative valence. PAM12-md (γ3) DANs have very different connectivity defined by strong input from the four neurons in cluster 6 (Figures 31 and 32D), which also provides input to 3 of the 4 subtypes of PAM08 (γ4) DANs and PAM07 (γ4<γ1γ2) DANs. These shared inputs suggest that PAM12-md signals positive valence. It is noteworthy that prior studies have implicated PAM12 DANs in both aversive and appetitive memory. At least some PAM12 DANs are shock responsive (Cohn et al., 2015; Jacob and Waddell, 2020) and can provide aversive reinforcement (Aso et al., 2012), but it has also been shown that their inhibition following sugar ingestion is required and even sufficient to assign positive value to odors during learning (Yamagata et al., 2016). Furthermore, a recent study of spaced aversive conditioning showed that the γ3 DANs are required for flies to gradually learn that the odor that is experienced without shock is safe (Jacob and Waddell, 2020). It will be important to clarify how the different γ3 DANs contribute to these learning processes.

Prior functional studies have suggested that different aversively reinforcing stimuli, such as electric shock, high and low heat, and bitter taste converge onto the PPL101 (γ1pedc) and PPL103 (γ2α′1) DANs (Das et al., 2014; Galili et al., 2014; Tomchik, 2013). One might therefore expect to find connectivity consistent with such convergence. Interestingly, rather than finding clear streams of input to these PPL1 DANs and the PAM12 (γ3) DANs, the identified individual input neurons emanate from discrete brain areas, consistent with these neurons conveying unique information. Only one neuron in cluster 28 projects from the SEZ and connects to PPL101 (γ1pedc) and PPL106 (α3). It may therefore relay the negative valence of bitter taste. Besides this SEZON we did not find other obvious inputs from primary sensory processing areas, or ascending pathways that might come from the ventral nerve cord and could for example relay shock from the legs. It therefore seems likely that most information conveyed to aversively reinforcing DANs has been pre-processed en route from the periphery.

Input connectivity also revealed insights into the overall organization of the PAM DANs, uncovering a remarkable heterogeneity and revealing a highly parallel architecture of rewarding reinforcement. Prior studies have established that PAM DANs are required for flies to learn using different types of reward, and that their artificial activation (or inactivation in the case of γ3 DANs) can assign positive valence to odors during learning (Liu et al., 2012; Burke et al., 2012; Perisse et al., 2013; Aso et al., 2014b; Huetteroth et al., 2015; Yamagata et al., 2015, 2016). In addition, a number of studies have implicated unique populations of PAM DANs in reinforcing different positive experiences, for example, the sweet taste and nutrient value of sugar (Huetteroth et al., 2015; Yamagata et al., 2015), water (Lin et al., 2014, Shyu et al., 2017), relative value (Perisse et al., 2013), the absence of expected shock (Felsenberg et al., 2018) and learned safety (Jacob and Waddell, 2020).

We found several input pathways that group the PAM DANs into both expected and unforeseen combinations. For example, the single neurons of clusters 23 (a SEZON) and 24 grouped the PAM08-md and PAM08-nc (γ4) DANs with the four subtypes of γ5 DANs. The cluster 24 neuron also provides input to a selection of β and β′ lobe innervating DANs. Cluster 7 neurons grouped a different collection of γ4 and γ5 with the PAM08-md (*γ*3) DANs and most PAM02 (β′2a), PAM05 (β′2p), PAM06 (β′2m) and PAM13 (β′1ap) and PAM14 (β′1m) DANs. These inputs are consistent with studies that have implicated the γ4, γ5 and β′2 compartments in reward learning, such as learning reinforced with water and the taste of sugar (Aso and Rubin, 2016; Burke et al., 2012; Huetteroth et al., 2015; Lin et al., 2014b; Liu et al., 2012; Shyu et al., 2017; Yamagata et al., 2015). In contrast, clusters 19 and 20 group unique subtypes of γ5 DANs with all three *α*1 DAN subtypes. Cluster 19 also provides strong input to both types of PAM13 DANs and the PAM14-can subtype. Prior work has shown that the γ5 and α1 DANs are required for the reinforcement of nutrient-dependent long-term sugar memory (Huetteroth et al., 2015; Yamagata et al., 2015; Ichinose et al., 2015). A similar complexity of combinatorial inputs is evident across most of the DAN subtypes in the PAM DAN population.

An unexpected finding was the strong and unique grouping of all subtypes of PAM04 (*β*2), two subtypes of PAM09 (β1) and the PAM10-vd (β1ped) subtype by the population of neurons in cluster 8. Two prior studies have shown that artificial activation of β1 and β′2 DANs together can form a long-lasting appetitive memory (Perisse et al., 2013; Huetteroth et al., 2015) and the cluster 8 input suggests that their activity is likely to be genuinely coordinated. The full importance of these DANs is not currently understood, but our analysis of MBON06 (β1>α) suggests that appetitive learning via these DANs modulates a key node of the MBON network (see Discussion).

The PAM11 (α1) DANs reinforce nutrient-dependent, long-term memory (Huetteroth et al., 2015; Ichinose et al., 2015; Yamagata et al., 2015). Their connectivity differs from other DAN cell types in that they receive input from a few clusters of input neurons that are very selective for PAM11 DANs but do not strongly discriminate between the three α1 DAN subtypes (Figures 31 and 33). For example, cluster 22 is a single neuron that provides strong, highly selective input to all three PAM11 subtypes and appears to convey input from the SEZ (Figure 33B). Clusters 19 and 20 convey information from the LH (Figure 33 E and F). When α1 DAN input neurons do show connections to other DANs, this input is much weaker and largely confined to positive valence DANs (Figure 31).

**Figure 33.**
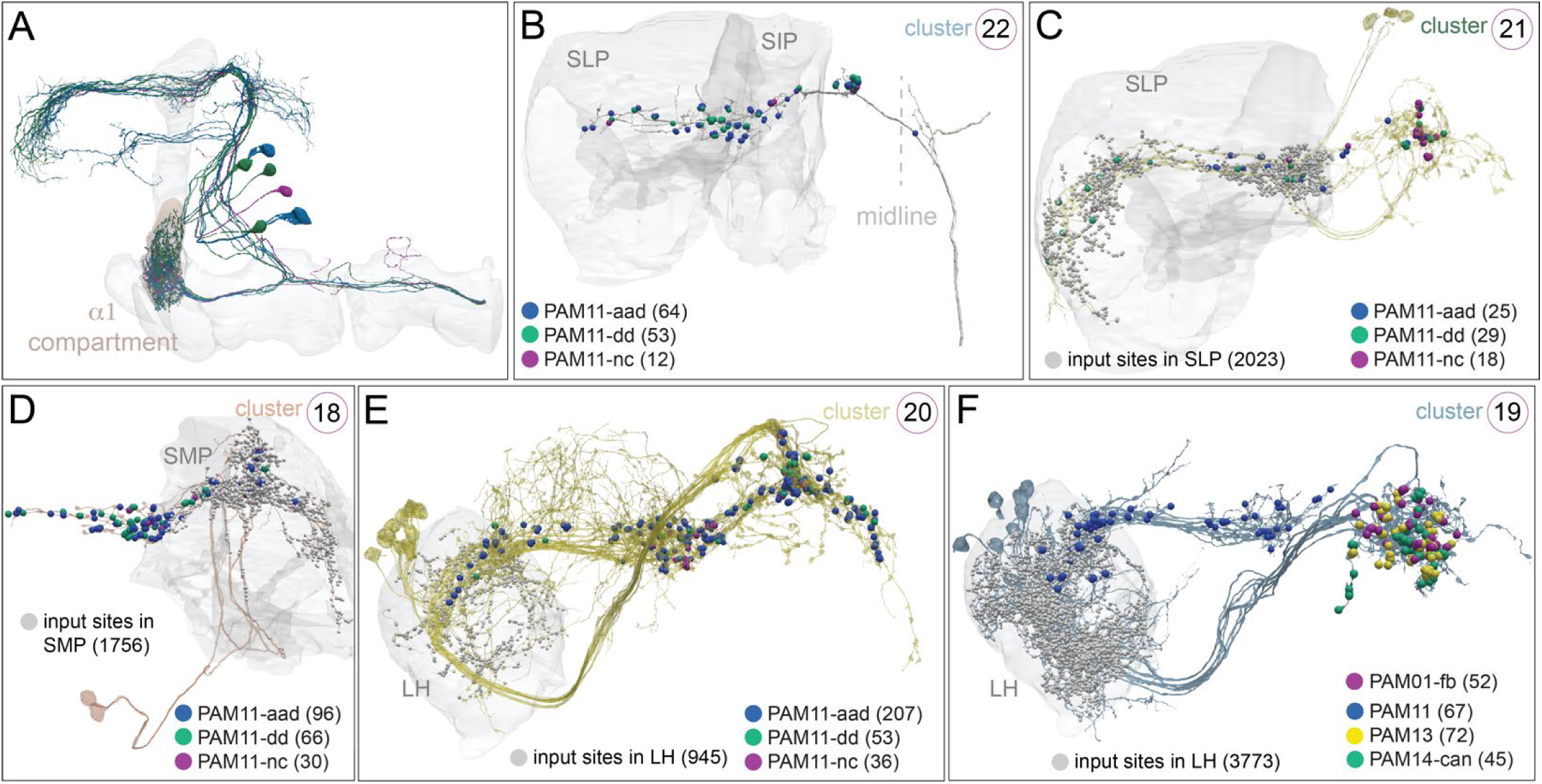
Examples of local neurons, SEZONs, and LHONs that target PAM11 DANs. (A) All PAM11 DAN subtypes are shown, color-coded as in Figure 28 and 28-figure supplement 1; the MB lobes are shown in grey and the α1 compartment is shaded brown. (B-F) Examples of neurons from clusters in Figure 30 that provide strong input to PAM11 DANs or to PAM11 and other positive valence PAM DANs are shown; cluster identity is indicated by the circled number in the upper right of each panel. Neuropils where DAN input clusters receive their inputs are shown in grey (except for SEZONs whose dendrites are not in the hemibrain volume) and position of output synapses to PAM11 subtypes are shown color-coded; synapse numbers are given in parentheses. Figure 33-figure supplement 1 shows more information about the neurons constituting these clusters. (B) The single neuron in <u>cluster 22</u>, a SEZON, makes synapses to all three subtypes of PAM11 DANs; PAM11 DANs are the only DAN targets of cluster 22. The same applies to its contralateral partner (see Figure 35-figure supplement 1A). (C) The local interneurons that make up <u>cluster 21</u> receive inputs in the SLP and connect to all three PAM11 subtypes, but not to other DANs. (D) The local interneurons that make up <u>cluster 18</u> (299626157, 361700223) receive inputs in the SMP and connect to PAM11 DANs, but not to other DANs. (E) The LHON <u>cluster 20</u> connects to all three PAM11 subtypes and weakly to PAM01 and other positive valence PAM DANs (compare to Frechter et al. 2018, Dolan et al. 2019, and Otto et al. 2020). (F) The LHONs <u>cluster 19</u> connect to all three PAM11 subtypes and also to PAM13 (β′1ap), PAM14 (β′1m) and PAM01-fb (γ5) DANs. The connections to the PAM11 and the other PAM DANs are located on different primary branches of the cluster 19 neurons. Connections contributing less than 0.5% of a neuron’s total dendritic input have been excluded.

We uncovered several additional features of the organization of the DAN system. Some input clusters synapse with multiple cell types, but only with specific DAN subtypes within them (Figure 36). There are many PAM specific clusters, such as clusters 7, 29, 32 (Figure 36-figure supplement 1) as well as PPL1 specific clusters, such as clusters 13, 14 and 15 (Figure 36-figure supplement 2). We also identified several clusters of neurons that innervate subsets of both PAM and PPL1 DANs (clusters 3, 9, 30, 33, 35; Figure 36 and Figure 36-figure supplement 1). Some clusters, such as 3, 7 and 30, have such dense innervation that their most notable feature might be the DANs that they do not innervate.

**Figure 34.**
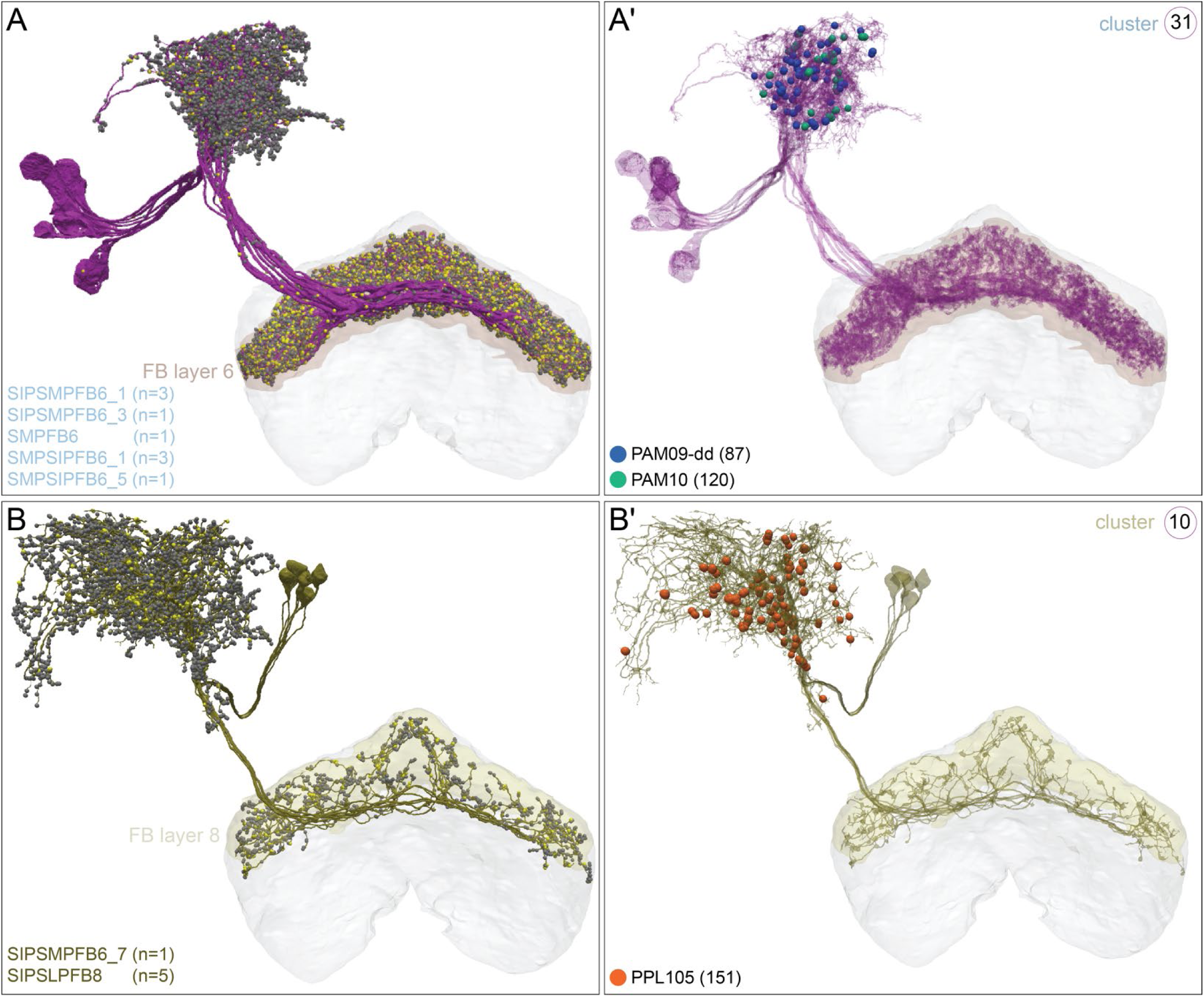
Distinct FB neurons provide input to β lobe PAM or PPL105 DANs. Two morphological clusters (31 and 10, Figure 31) of neurons from FB are among the strongest DAN inputs. These FB neurons have arbors of mixed polarity both in the SMP and in the FB, as can be seen in panels A and B where their presynaptic and postsynaptic sites are shown in yellow and grey, respectively. (A and A′) <u>Cluster 31</u> neurons are shown in magenta and consists of nine FB layer 6 neurons of five cell types. These neurons synapse onto the dendrites of three potentially positive valence PAM DAN subtypes: PAM09-dd (β1pedc; blue dots), and two β1 DAN subtypes, PAM10-can and PAM10-nc(dd) (combined in figure; green dots); synapse numbers are given in parentheses. Remarkably, seven of the nine neurons of cluster 31 are connected, via an interneuron, to MBON19 (α2p3p; see Figure 20-figure supplement 1C and Video 29). (B and B′) <u>Cluster 10</u> neurons are shown in yellow and consist of five layer 8 (cell type: SIPSLPFB8) and one layer 6 FB neurons and make synapses in the SMP onto PPL105 (α′2α2; red dots; synapse number is given in parentheses); the negative valence PPL105 is the only DAN they innervate. Connections contributing less than 0.5% of a neuron’s total dendritic input have been excluded.

**Figure 35.**
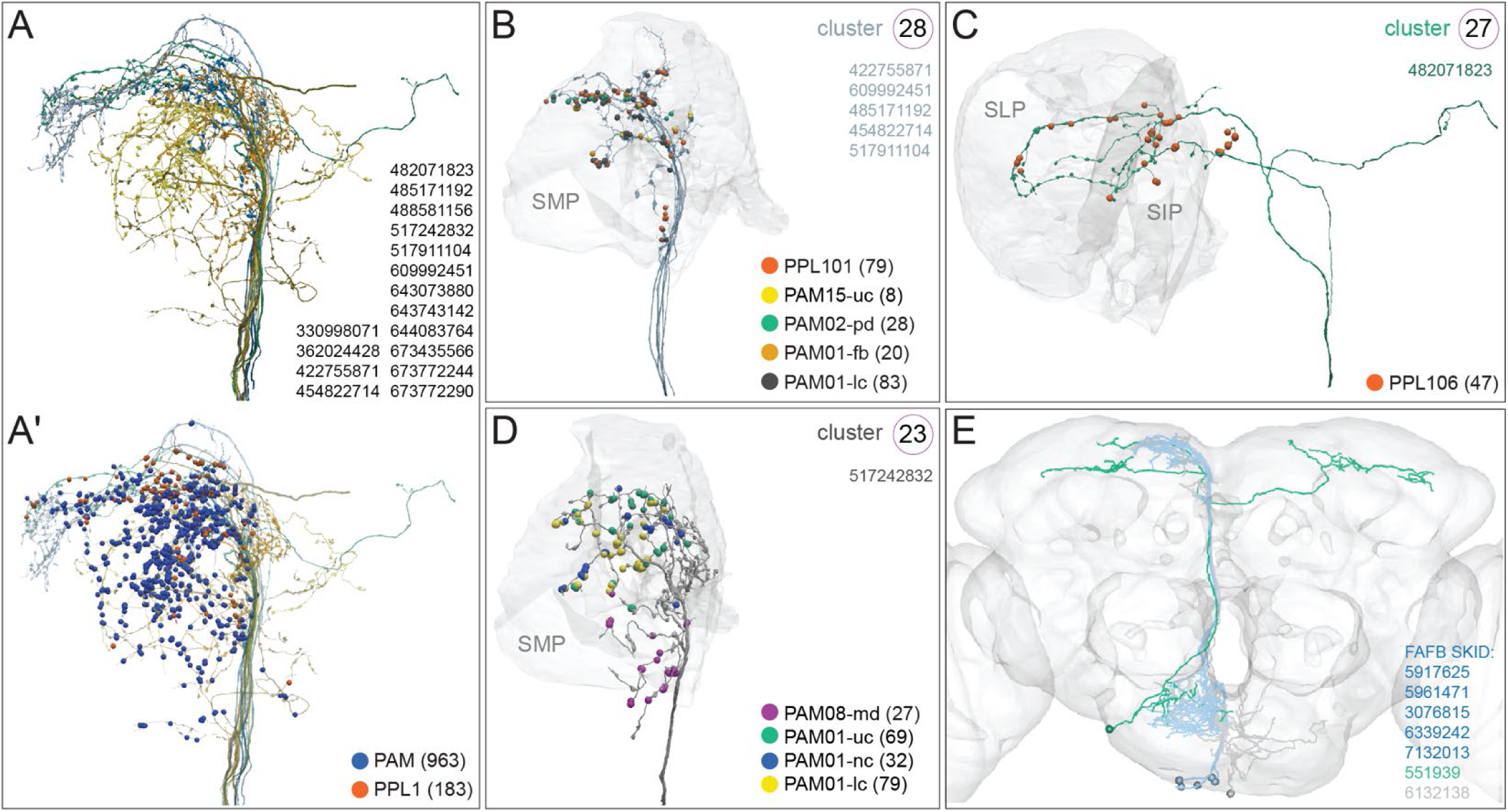
Output neurons from different areas of the SEZ tile the SMP where they synapse onto select DAN subtypes. (A) Axons from multiple SEZON cell types in clusters 22, 23, 24, 26, 27, and 28 (see Figure 31) that provide strong inputs to DAN subtypes are shown together. (A′) The same SEZON axons shown with synapses connecting to PAM DANs (blue, from 5 clusters) and to PPL1 DANs (red, from 2 clusters); synapse numbers are given in parentheses. (B) The five neurons of SEZON <u>cluster 28</u> (light blue; these neurons correspond to the SEZON01 cluster 5.4.1 of Otto et al., 2020) branch in the SMP and synapse onto the PPL101 (red), PAM01-lc (black), PAM01-fb (orange), PAM02-pd (green), and PAM15-uc (yellow) subtypes; synapse numbers are given in parentheses. (C) The single neuron in SEZON <u>cluster 27</u> (green) branches in the SMP and SIP and synapses only onto PPL106; synapse number is given in parentheses. (D) The single neuron in SEZON <u>cluster 23</u> (grey) branches in the SMP and connects to positive valence DANs: PAM01-lc (yellow), PAM01-uc (green), PAM01-nc (blue), and PAM08-md (magenta); synapse numbers are given in parentheses. (E) The axons of SEZON neurons can be matched to those in the FAFB EM volume (Zheng et al., 2018) to permit the retrieval of their dendritic field, which resides in tissue that is missing from the hemibrain dataset; skeleton ID numbers from FAFB are shown. FAFB matches to hemibrain SEZONs are shown and colored to match their hemibrain counterparts (colors correspond to (B-D)). Their dendritic fields differ, suggesting these SEZON clusters are likely to convey distinct information. Connections contributing less than 0.5% of a neuron’s total dendritic input have been excluded.

**Figure 36.**
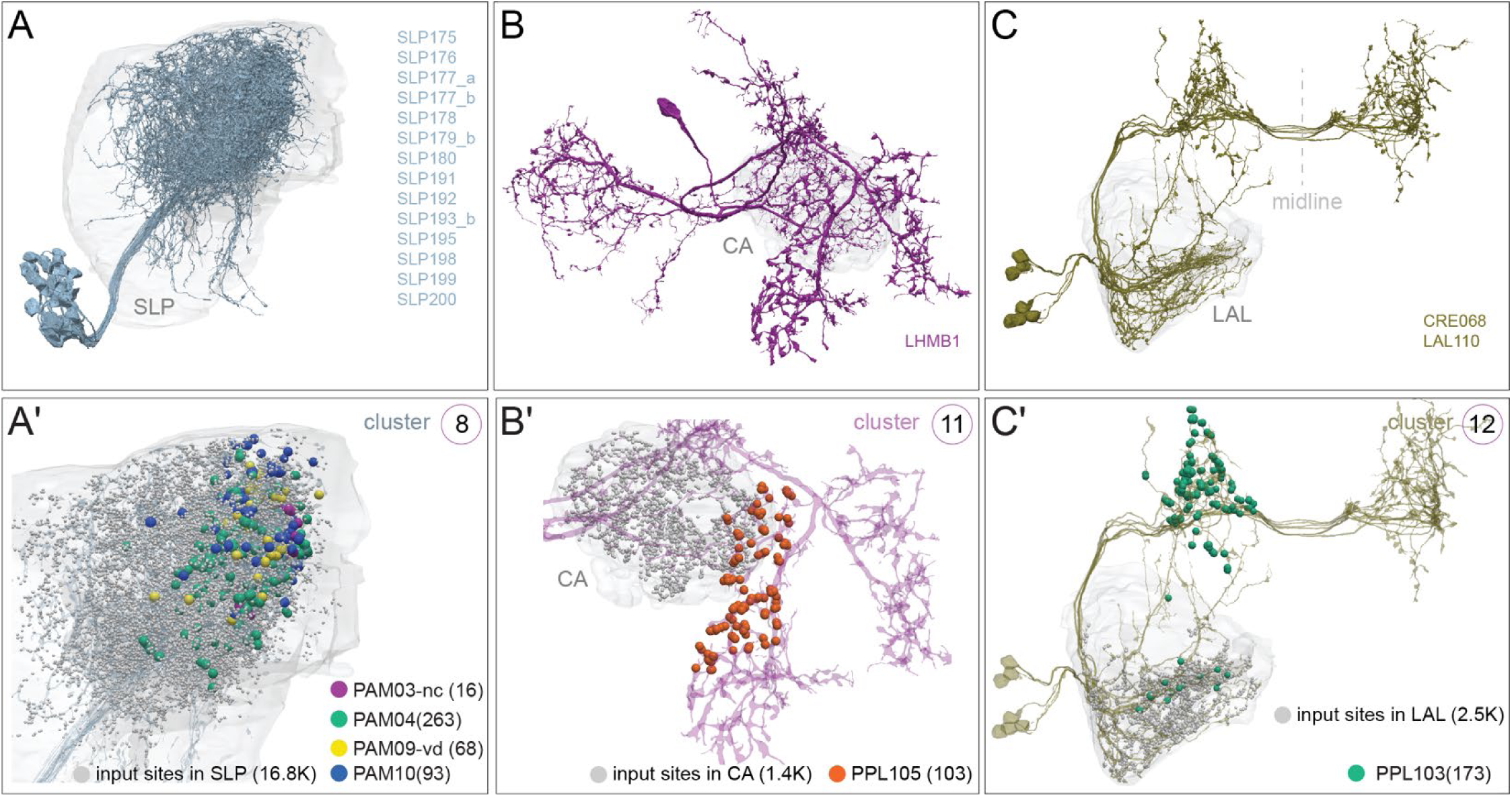
A number of input neurons specifically target DAN groups of the PAM or PPL1 clusters. Neurons providing strong input to PAM or PPL1 DANs are shown. Neuropil areas of inputs to these neurons are shown in grey in (A-C) and synapses to DANs are shown color-coded in (A′-C′); synapse numbers are given in parentheses. The threshold for connectivity is 0.5% of the DANs total inputs (Figure 31). (A, A′) <u>Cluster 8</u> is comprised of 26 neurons of 14 similar cell types which provide input to subtypes of β-lobe PAM DANs in the SLP: PAM10 (β1; blue); PAM09-vd (β1pedc; yellow); all PAM04 subtypes (β2; green); and the PAM03-nc (β2β′2a; magenta) subtype. (B, B′) The single neuron in <u>cluster 11 receives inputs in the CA and connects to PPL105 (α′2α2; red). (C, C′) The seven neurons in cluster 12 (LAL110, CRE068) receive inputs in the (LAL) and provide the strongest input to PPL103 (γ2α′1; green). Connections contributing less than 0.5% of a neuron’s total dendritic input have been excluded.</u>

Two discrete classes of dorsal fan-shaped body (FB) neurons make up clusters 10 and 31, which provide selective input to DANs of opposite valence (Figure 31 and 34). The arbors of the FB neurons where they synapse to these DANs are of mixed polarity and could simply serve as a local relay for other inputs. We also note that (Hu et al., 2018) reported that more ventral layers of the FB respond to electric shock and their artificial activation could substitute for negative valence reinforcement, but it remains unclear what role, if any, more dorsal FB layers might play.

Neurons from the SEZ provide a major source of DAN inputs. While studies have shown that pleasant and unpleasant tastes can reinforce learning by engaging different DAN subsets (Das et al., 2014; Huetteroth et al., 2015; Kirkhart and Scott, 2015; Masek et al., 2015), the relevant input pathways are only recently emerging from connectomic studies. A comparison of inputs to PPL101 (γ1pedc), PAM01 (γ5) and PAM05/6 (β′2) DANs identified a set of SEZONs. The hemibrain dataset provides access to the projections of all SEZONs (although characterizing their dendritic arbors required identifying the corresponding neurons in the FAFB volume) and confirms that they are a major source of DAN input (clusters 22-28). Some single neuron SEZON clusters tend to connect to either exclusively positive valence (clusters 22, 23) or negative valence (cluster 27) DANs, consistent with them relaying the valence of different tastants to the MB. Some clusters contain multiple SEZONs (clusters 25, 26 and 28); clusters 25 and 26 innervate only positive valence DANs, while SEZONs in cluster 28 innervate either PPL101 or DANs of both positive and negative valence.

As with the PPL1 DANs, most neurons providing strong input to PAM DANs do not project from primary sensory areas in the brain. PAM DANs are therefore also likely to receive highly processed value-based information. However, as discussed above for PPL101 and previously noted for the PAM01 (γ5) and PAM02 (β′2a) DANs (Otto et al., 2020), we found that several PAM DANs receive significant input from SEZONs. Since blocking some of these neurons has been shown to impair sugar and/or bitter taste learning (Otto et al., 2020), SEZON input suggests that taste may provide fairly direct valence-related signals to the DANs. Various subtypes of PAM01, PAM02, PAM03, PAM08, PAM09, PAM10 and PAM15 DANs all receive SEZON input, some of which is highly selective. The single SEZON in cluster 22 exclusively synapses onto the three subtypes of PAM11 (α1) DANs, whereas another (cluster 27) synapses selectively with the PPL106 (α3) DAN. We speculate that the prevalence of SEZON innervation to the DANs indicates the ecological relevance of evaluating taste for the fly, and that these very selective pathways to PAM11 (α1) and PPL106 (α3) might represent stimuli that are of unique importance. PAM11 DANs have been previously implicated in the nutrient-dependent reinforcement of long-term memories. In addition to the recently reported involvement of SEZONs in sugar learning (Otto et al., 2020), an earlier study demonstrated the importance of a physical and functional connection between octopaminergic neurons and the PAM01 (γ5) and PAM02 (β′2a) DANs (Burke et al., 2012). Also within the top 50 inputs, we identified synapses from the OA-VPM3 neurons onto PAM01 (γ5), PAM01 (γ4) and to a lesser extent PAM02 (β′2a) DANs. It will be important to understand the relationship between the OA and SEZON inputs to the rewarding DANs.

Our work provides a comprehensive description of the substructure of DANs, uncovering a complexity that goes far beyond the previously defined 21 cell types (Aso et al., 2014a). Understanding the functional significance of this complexity will require more complete knowledge of the information about the outside world and internal brain state that is conveyed by the hundreds of neurons that provide input to DANs. Our analyses of the DAN connectivity, together with what we learned about the segregation of sensory modalities in different classes of KCs, the selective connectivity of MBONs to KCs, and the discovery of a new class of MBONs, opens a window onto a much broader and richer landscape of MB circuitry underlying what a fly might be able to learn, remember and use to guide its behavior.

## Discussion

The hemibrain dataset we have analyzed contains the most comprehensive survey of the cell types and connectivity of MB neurons available to date. This connectome allowed us to probe in fine detail the circuitry underlying canonically proposed functions of the MB, including the representation of olfactory information by KCs, computation of valence by MBONs, and reinforcement of associations by DANs. We found patterns in the input to DANs, MBON-to-DAN and MBON-to-MBON connectivity that suggest how associative learning in the MB can affect both the acquisition of new information through learning and the expression of previously learned responses. The connectome also reveals circuitry that supports non-canonical MB functions, including selective structure in non-olfactory pathways, a network of atypical MBONs, extensive heterogeneity in DAN inputs, and connections to central brain areas involved in navigation and movement.

### Advantages and limitations of fly connectomics

Our analysis of the hemibrain connectome relied heavily on an extensive catalog of previously identified and genetically isolated cell types and on decades of study illuminating the link between mushroom body physiology and fly behavior. It is worth emphasizing the interdependency of anatomy, physiology and behavior as we enter the post-connectomic era in fly research. Some of the neurons we have described that appear similarly connected may turn out to have diverse functions due to different physiology (Groschner et al., 2018) and, conversely, neurons that are morphologically distinct may turn out to have similar functions. In addition, a set of inputs with high synapse counts might appear, at the connectome level, to represent a major pathway for activating a particular neuron, but this will not be true if these inputs rarely fire at the same time. Likewise, a set of highly correlated inputs can be effective even if their individual synapse counts are modest. Finally, the connectome does not reveal gap junction connections or identify more distant non-synaptic modulation. These caveats should be kept in mind when interpreting connectome data (Bargmann and Marder 2013).

### Comparison with the larval MB

The cell type constituents and circuit motifs of the MB in the adult fly have many similarities with its precursor at the larval stage of development (Aso et al., 2014a, Eichler et al., 2017, Takemura et al., 2017, Eschbach et al., 2019, 2020a,b). Both the larval and adult MBs support associative learning and, in both, PNs from the antennal lobe that convey olfactory information provide the majority of the sensory input, complemented by thermal, gustatory and visual sensory information that is segregated into distinct KC populations. However, the multi-layered organization of non-olfactory inputs in the main and accessory calyces (including integration of diverse input sources by LVINs) suggests that the KC representation in the adult is more highly enriched and specialized for non-olfactory sensory features. It is worth noting that the earliest born types of each of the three main adult KC classes (KCγs, KCγd, KCγt, KCα′β′ap1, KCαβp) appear to be specialized for non-olfactory sensory cues, and in most cases their dendrites lie in the accessory calyces.

In the first instar larval MB, the only larval stage for which a connectome is available (Eichler et al., 2017), there are roughly 70 mature KCs, which is increased nearly 30-fold in the adult. This enrichment likely increases odor discrimination and olfactory memory capacity. The larval MB has only eight compartments in its horizontal and vertical lobes. Although the increase in the adult to 15 compartments is only about a factor of two, the extent of their DAN modulation is greatly expanded. The larva has only seven confirmed DANs and five additional cells of unknown neurotransmitter thought to provide modulatory input, a factor of 15-fold less than the adult. Whereas each larval KC innervates all 8 compartments, individual adult KCs innervate only five out of 15 compartments. Therefore the DANs in the larva are capable of modulating all KCs, whereas in the adult, DANs modulate specific subset of KCs. Subcompartmental targeting of DAN modulation in the adult increases this difference even more.

MBONs feedback onto DANs and converge onto common downstream targets in both the larva (Eschbach et al. 2020a,b) and adult, implying some shared computational strategies. However, the greatly increased DAN complexity in the adult fly and the presence of subcompartmental organization of DAN axon targets not present in the larva suggest a substantial increase in the specificity of the learning signals involved in memory formation and raises the possibility of modality-specific learning signals to complement the multimodal KC representation.

### Sensory input to KCs

Previous theoretical work has emphasized the advantages of mixing sensory input to the KCs so that they provide a high-dimensional representation from which MBONs, guided by DAN modulation, can derive associations between sensory input and stimulus valence (Litwin-Kumar et al., 2017). The connectome data modifies this viewpoint in two ways. First, although olfactory input to the KCs is highly mixed, various structural features reduce the dimensionality of the KC odor representation. A recent analysis of EM data from an adult *Drosophila* MB identified groups of PNs thought to represent food odors that are preferentially sampled by certain KCs (Zheng et al., 2020). Consistent with this observation, our analysis revealed subtype-specific biases in PN sampling by KCs, including an overrepresentation of specific glomeruli by α/β and α′/β′ KCs. These biases, as well as other structural features, appear to arise from the stereotyped arrangement of PN axons within the calyx and their local sampling by KCs. This may reflect a developmental strategy by which the KC representation is organized to preferentially represent particular PN combinations. Further analyses of KC connectomes across hemispheres and animals, as well as experimental studies, will help evaluate the impact of this structure.

The hemibrain connectome also revealed a second, more dramatic, structural feature of sensory input to the MB: non-olfactory input streams corresponding to visual and thermo-hygro sensation are strongly segregated. This organization may reflect the nature of stimulus-valence associations experienced by flies. When valence depends on a specific combination of stimulus attributes, such as binding to particular sets of olfactory receptors, MBONs need to be able to sample those combinations to successfully identify the stimulus. However, if different sensory modalities can be used separately to identify valence, mixing may not be necessary. For example, either the visual appearance of an object or its odor may individually be sufficient to identify it as a food source (for a discussion of multi-sensory learning in flies, see Guo and Guo, 2005). We asked whether there is an advantage in having separate modality-specific sensory pathways, as seen in the connectome data, when valences can be decomposed in this way.

To address this question we considered a model in which KC input is divided into two groups, visual and olfactory (Figure 38A). In one version of the model (“shared KC population”), all KCs receive sparse random input from both types of PNs, corresponding to a high degree of mixing across modalities. In the other (“separate KC populations”), half of the KCs receive sparse random input exclusively from visual PNs and half only from olfactory PNs, corresponding to no cross-modal mixing. We define two tasks that differ in the way valences are defined. In the first (factorizable) version of the task, each olfactory stimulus and each visual stimulus is assigned a positive or neutral subvalence, and the net valence of the combined stimulus is positive if either of these components is positive (olfactory OR visual). In the second (unfactorizable) version, a valence is randomly assigned independently to each olfactory+visual stimulus pair. We evaluated the ability of a model MBON, acting as a linear readout, to determine valence from these two different KC configurations. Separate modality-specific KC populations are indeed beneficial when the valence can be identified from either one modality or the other (the ’factorizable’ case in Figure 38B). Dedicating different KC subtypes to distinct sensory modalities allows the predictive value of each modality to be learned separately. This result suggests that the divisions into KCs specialized for visual, thermo/hygro and olfactory signals may reflect a property of natural valences across these modalities.

**Figure 37.**
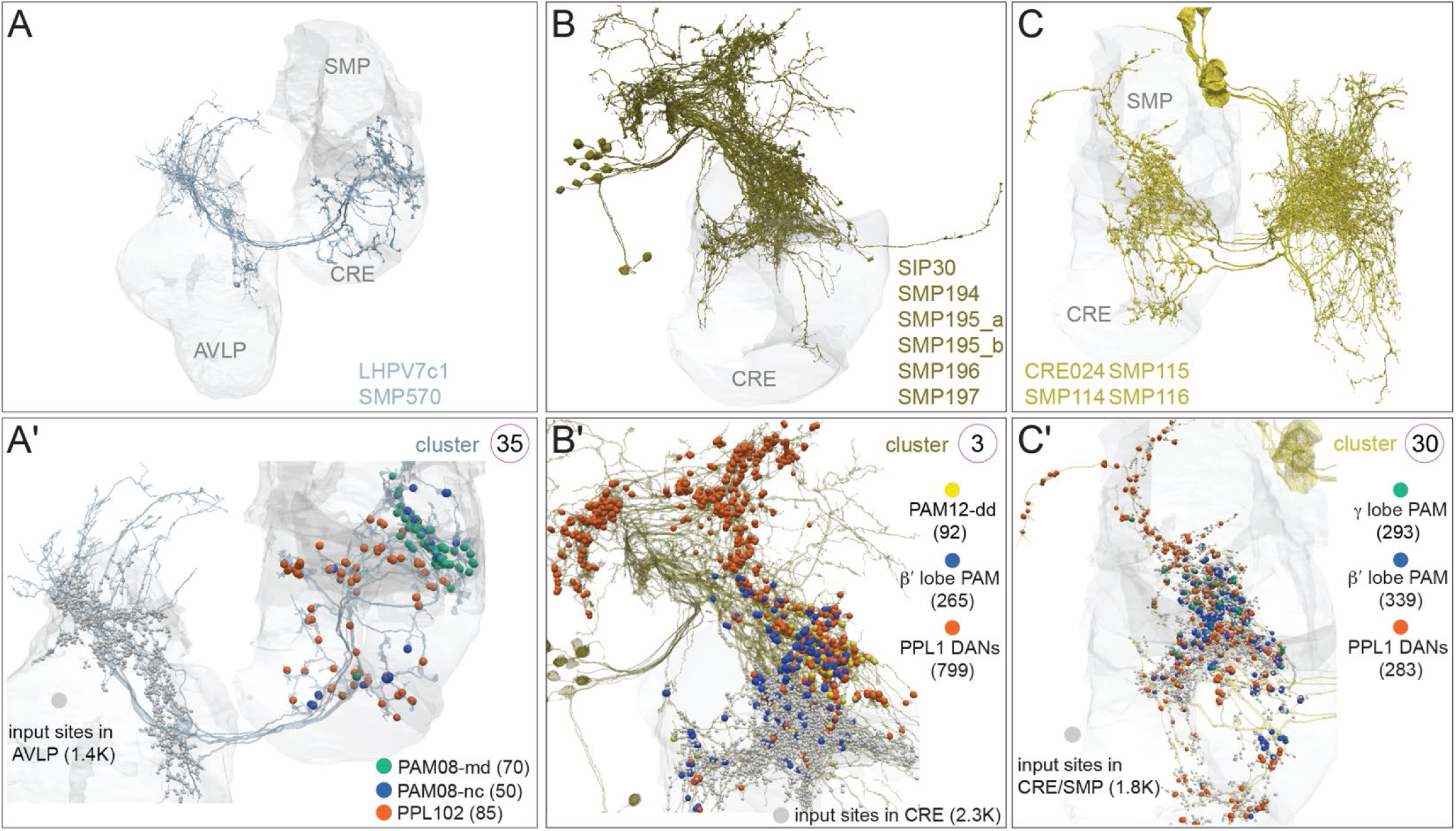
Neurons that provide strong input to both PAM and PPL1 DANs. Neuropil areas indicated in grey and synapses to DANs are color coded; synapse numbers are given in parentheses. (A, A′) The four neurons of <u>cluster 35</u> receive inputs in the AVLP and provide strong input to the PAM08-md (γ4; green) and PAM08-nc (blue synapses) subtypes as well as the PPL102 (γ1) DAN (red synapses). (B, B′) The 13 neurons of <u>cluster 3</u> receive some inputs in the CRE; they constitute the strongest input to PPL105 (α′2α2) and PPL106 (α3) and also connect to PPL101 (γ1pedc), PPL102 (γ1), and PPL103 (γ2α′1) (all PPL1 DAN synapses combined, red), all of the β′ lobe PAM DAN subtypes (blue synapses) and the PAM12-dd (γ3) subtype (yellow synapses). (C, C′) <u>Cluster 30</u> has four neurons that receive inputs in the SMP and CRE and are upstream of negative valence PPL1 neurons (PPL101, PPL102, PPL105, and PPL106; red synapses) and PAM DANs innervating the β’ (blue synapses) and γ (green synapses) lobe (PAM08, PAM01, PAM03, PAM05, PAM06). While both cluster 3 and 30 connect to both PPL1 and PAM DANs, cluster 30 connects to γ lobe PAM DANs, in contrast to cluster 3 favoring β′ lobe PAM DANs.

**Figure 38.**
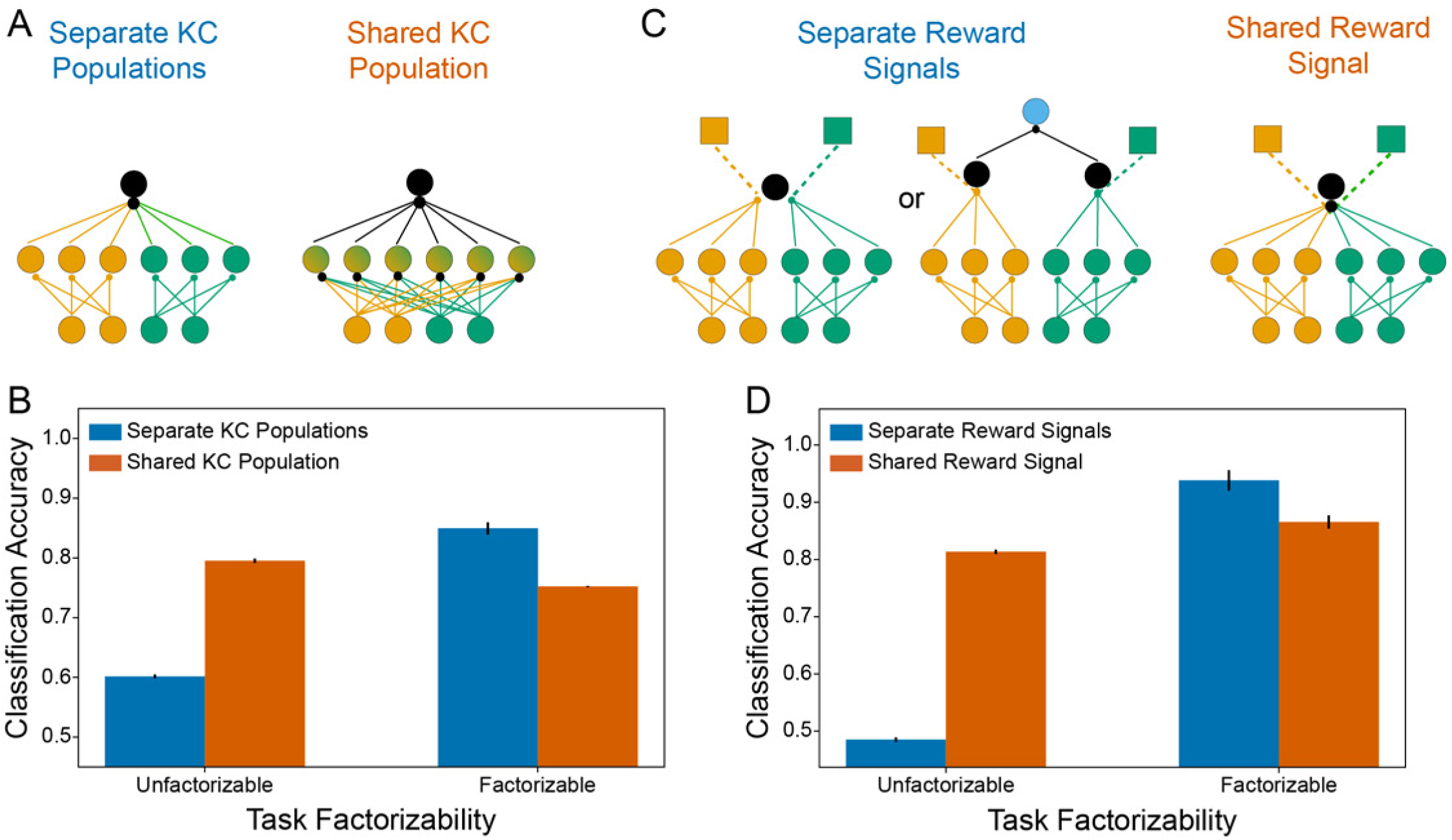
Effects of segregated or mixed-modality signals. (A) Schematic of two possible architectures for an MB-like circuit, in which KCs are either specialized for one sensory modality (left) or receive mixed input (right). Bottom layer circles represent PN inputs, middle layer circles represent KCs, and black output circles represent MBONs. Green and yellow shades indicate different sensory modalities. (B) Task performance in a stimulus discrimination model for the architectures shown in A. Stimuli consist of random binary patterns in two sensory modalities. In the “factorizable” task, the stimulus from each modality has an assigned valence (positive or negative) and the overall valence is positive only if both constituent valences are positive. In the “unfactorizable” task, each stimulus is randomly assigned its own unique valence, irrespective of its constituent modalities. A linear readout, modeling an MBON, is fit to discriminate the valences of the stimuli based on the activations of the KC population. The linear fit procedure corresponds to the experienced-based plasticity of KC-MBON synapses. Task performance is defined as the binary classification accuracy—that is, the frequency with which the thresholded response of the MBON corresponds correctly to the valence of the input—of this MBON readout. 25 stimuli are presented in the unfactorizable task and 200 in the factorizable task, to make overall task difficulty comparable. Error bars represent standard error over 20 simulations. (C) Schematic of possible architectures for DAN-dependent plasticity in an MB-like circuit. In the leftmost model, DANs (colored squares) modulate KC-MBON synapses for KCs of a particular sensory modality. The second-to-left model depicts two MBONs, each integrating input from KCs of a particular sensory modality, and each with its own corresponding DAN, that additively converge onto a common output (blue). These two models are functionally equivalent, and hence are grouped together under “separate reward signals.” In the rightmost model, there is a single MBON, and DANs modulate all the KC-MBON synapses (right). (D) Task performance in a stimulus discrimination model (same as B) for the architectures shown in C. For each task the optimal KC structure from B is used (“separate” for the factorizable task, “shared” for the unfactorizable task).

### Sub-compartmental modulation by DANs

On the basis of light-level studies, DAN modulation of KC-to-MBON synapses has been considered to operate at the resolution of MB compartments. However, taken with other recent studies (Lee et al., 2020; Otto et al., 2020), the morphology and connectivity data indicate that functionally distinct PAM-DAN subtypes operate within a MB compartment. DAN subtypes receive different inputs and likely modulate different KC-MBON synapses within a compartment.

A prior analysis showed that PAM01-fb DANs were required to reinforce the absence of expected shock during aversive memory extinction, whereas a different set of γ5 DANs were needed for learning with sugar reinforcement (Otto et al., 2020). It is conceivable that another γ5 DAN subtype is required for male flies to learn courtship rejection (Keleman et al., 2012). The connectome data revealed DAN subtypes in every compartment that is innervated by PAM DANs.

The γ3 compartment provides an interesting example of subcompartmental targeting of modulation by DANs and of KC input onto MBONs (Figure 39). There are two subtypes of PAM DANs and three types of MBONs in the γ3 compartment. MBON09 and MBON30 primarily receive olfactory information from γm KCs whereas MBON33 primarily receives visual information from γd KCs. PAM12-dd and PAM12-md DANs appear to modulate KC inputs to MBON09/MBON30 or MBON33, respectively. Although existing driver lines do not separate the dd and md subtypes, PAM-γ3 DAN activity is supressed by sugar (Yamagata et al., 2016) and activated by electric shock (Jacob and Waddell, 2020). We found that PAM12-dd DANs share input with PPL1 DANs conveying punishment signals, while PAM12-md DANs are co-wired with PAM08 (γ4) DANs conveying reward signals. Thus, synaptic transmission from two sets of modality-specific KCs to different MBONs can be independently modulated by DANs signaling different valences, all within a single compartment.

**Figure 39.**
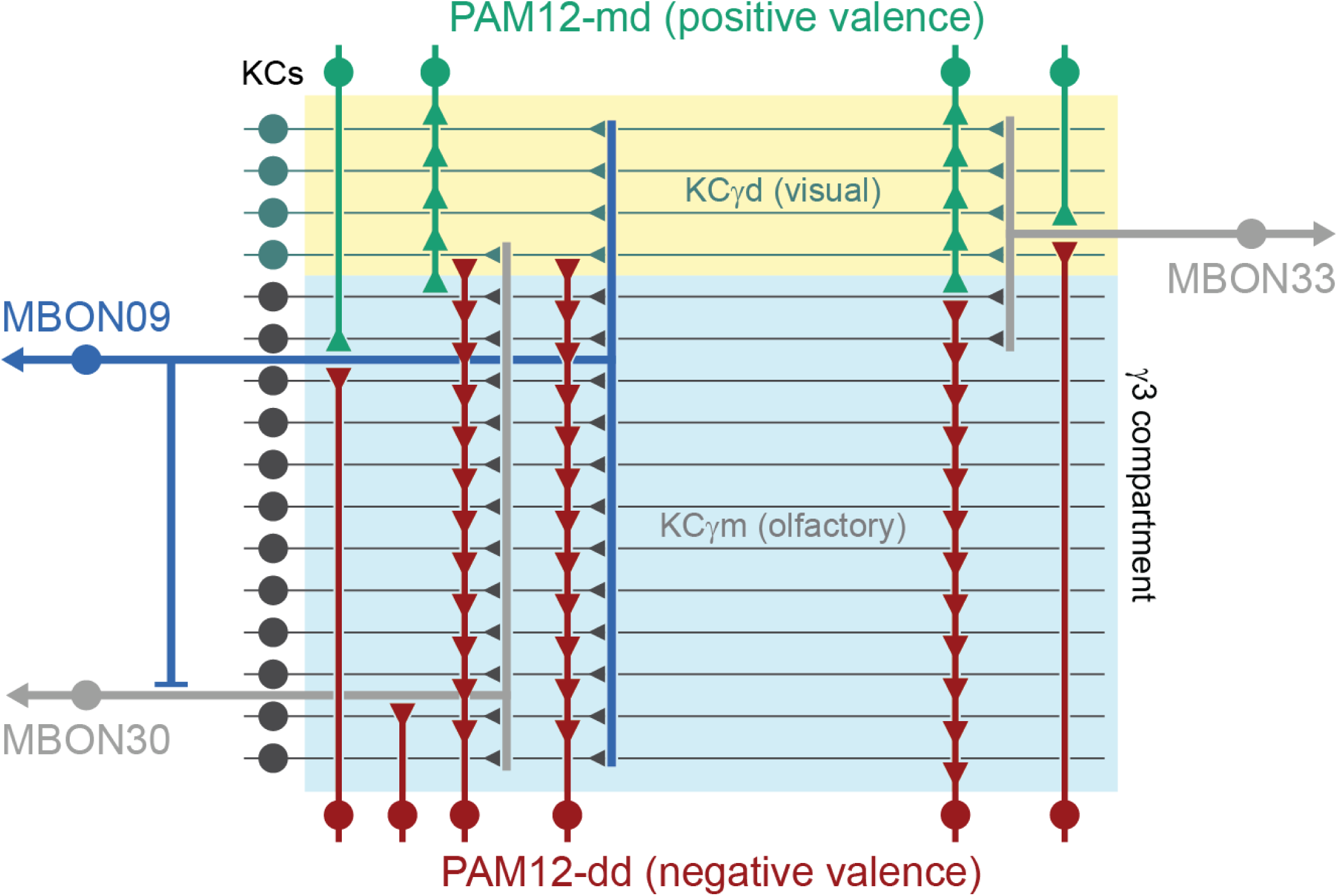
Schematic of subcompartment DAN and MBON innervation within γ3. The two PAM12 (γ3) DAN subtypes likely represent opposite valence based on their shared input with either aversively reinforcing PPL1-DANs or appetitively reinforcing PAM08 (γ4) DANs. Whereas PAM12-md DANs selectively innervate the visual γd KCs, the PAM12-dd DANs innervate both γd and γm (olfactory) KCs. The dendrite of atypical MBON30 is restricted to the olfactory γm (olfactory) KCs and its DAN input is exclusively from the PAM12-dd subtype. In contrast, the dendrite of atypical MBON33 preferentially collects γd KC input and receives input from both PAM12-md and PAM12-dd DANs. The GABAergic MBON09 (γ3β′1) pools γm and γd KC input, receives DAN input from PAM12-md and selectively synapses onto MBON30.

To explore the implications of DAN modulation that is specific to sensory modality, we extended the model presented above (Figure 38A) by including two DANs, one conveying the visual component of valence and the other the olfactory component (Figure 38C). KCs were divided into visual and olfactory modalities, and we considered two configurations for DAN modulation, one, “shared reward signal”, that is compartment-wide and non-specific, and the other, “separate reward signals”, in which each DAN only induces plasticity onto KC synapses matching its own modality. This latter case models a set of DANs that affect synapses from visual KCs onto the MBON and another set that affect olfactory synapses (alternatively, it could model two MBONs in different compartments that are modulated independently and converge onto a common target). We find that, when KCs are divided into separate populations and separate modalities can be used to identify valence, it is beneficial to modulate the pathways individually (Figure 38D).

### Functional implications of DAN input heterogeneity and MBON feedback to DANs

Our analysis revealed that DANs receive very heterogeneous inputs but, nonetheless, some DANs both within and across compartments often share common input. This combination of heterogeneity and commonality provides many ways of functionally combining different DAN subtypes. For example, we expect that it allows DANs to encode many different combinations of stimuli, actions and events in a state-dependent manner and to transmit this information to specific loci within the MB network.

In addition to heterogeneous inputs from a variety of brain regions, the DAN network receives a complex arrangement of within and across compartment monosynaptic input from a variety of MBONs, using both excitatory and inhibitory neurotransmitters. We found that nearly all MB compartments contain at least one direct within-compartment MBON-DAN feedback connection. MBON feedback onto DANs allows previously learned associations that modify MBON activity to affect future learning. MBONs that feedback onto the same DANs that modulate them could, if the result of learning is reduced DAN activity, prevent excess plasticity for an already learned association. In cases where MBON activity excites the DAN, the self-feedback motif could assure that learning does not stop until the MBON has been completely silenced. More generally, MBON inputs to DANs imply that dopaminergic signals themselves reflect learned knowledge and the actions it generates. This could, in turn, allow MBON modulation of DAN activity to support a number of learning paradigms beyond pure classical conditioning, including extinction, second-order conditioning, operant conditioning and reinforcement learning.

Flies can perform second-order conditioning, in which a stimulus that comes to be associated with reinforcement may itself act as a pseudo-reinforcement when associated with other stimuli (Tabone and de Belle, 2011). This computational motif of learning the value of sensory states and using the inferred value as a surrogate reinforcement to guide behavioral learning is the core principle behind a class of machine learning techniques known as actor-critic algorithms. These algorithms consist of two modules: the “actor,” whose job is to map sensory inputs to behavioral outputs and the “critic,” whose job is to map sensory inputs to their inferred values and provide these values as a learning signal for the actor. In the mammalian basal ganglia, the dorsal and ventral striatum, the latter of which strongly influences the activity of dopamine neurons in areas including VTA, have been proposed to represent actor and critic modules, respectively. In the mushroom body, this perspective suggests a possible additional ‘critic’ function for some MBONs beyond their known ’actor’ role in directly driving behaviors. Consistent with this view, activation of individual MBONs can excite DANs in other compartments (Cohn et al., 2015; Felsenberg et al., 2017). We found strong direct monosynaptic connections between some MBONs and DANs in other compartments. Functional studies will be needed to determine which MBONs, if any, participate in an actor-critic arrangement, and which circuit mechanisms—for example, release from inhibition, or reduction of excitation—are at work.

Another potential role of cross-compartment MBON-DAN feedback is to gate the learning of certain associations so that the learning is contingent on other associations having already been formed. Such a mechanism could support forms of memory consolidation in which long-term memories are only stored after repeated exposure to a stimulus and an associated reward or punishment. Prior studies have linked plasticity in the γ1 and γ2 compartments to short-term aversive memory (Aso et al., 2012; Hige et al., 2015; Perisse et al., 2016; Sejourne et al., 2011) and plasticity in the α2 and α3 compartments to long-term aversive memory (Aso and Rubin, 2016; Awata et al., 2019; Jacob and Waddell, 2020; Pai et al., 2013; Sejourne et al., 2011). The cross-compartmental MBON to DAN connections we observed suggest an underlying cicuit mechanism for this “transfer” of short to long term memory. Aversion drives PPL101 (γ1pedc) and depresses the conditioned odor-drive to the GABAergic MBON11 (γ1pedc>α/β). MBON11 is strongly connected with PPL1-γ1pedc, but only much more weakly connected with PPL105 and PPL106. Depression of MBON11 will therefore release the PPL105 and PPL106 DANs from MBON11-mediated inhibition, increasing their activity in response to the conditioned odor and making them more responsive during subsequent trials. The net result is that short-term aversive learning by MBON11 (γ1pedc>α/β) promotes long-term learning in MBON18 (α2sc) and MBON14 (α3) by releasing the inhibition on the dopamine neurons that innervate the α2 and α3 compartments (Sejourne et al., 2011; Jacob and Waddell, 2020). Indeed pairing inactivation of MBON11 with odor presentation can form an aversive memory that requires output from PPL1-DANs (Ueoka et al., 2017) and optogenetic stimulation of MBON11 during later trials of odor-shock conditioning impairs long-term memory formation (Awata et al., 2019).

Cross-compartment MBON-DAN feedback may also enable context-dependent valence associations, such as the temporary association of positive valence with neutral stimuli when a fly is repeatedly exposed to aversive conditions. Multiple, spaced aversive conditioning trials were recently shown to form, in addition to an aversive memory for the shock-paired odor, a slowly emerging attraction, a ‘safety’ memory, for a second odor that was presented over the same training period without shock (Jacob and Waddell, 2020). MBON09 (γ3β′1) appears to play a critical role in the formation of this safety memory. The synapses from KCs conveying the shock-paired odor are depressed in that the portion of MBON09’s dendrite that arborizes in γ3, while the synapses from KCs conveying the safety odor are depressed in its β′1 arbor. These combined modulations should gradually release downstream neurons from MBON09 feedforward inhibition, consistent with the proposed mechanism for PAM13/14 (β′1) and PAM05/06 (β′2m and β′2p) DANs becoming more responsive to the safe odor (Jacob and Waddell, 2020). The connectome data revealed that MBON09 is directly connected to the PAM13/14 (β′1ap) and PAM05 (β′2p) DANs, consistent with release from inhibition underlying the delayed encoding of safety.

DANs also make direct connections with MBONs in each MB compartment, and optogenetic activation of PAM11 DANs can directly excite MBON07 with slow dynamics (Takemura et al., 2017). The excitation of the MBON by the DAN during memory retreival can temporally suppress expression of the memory, since the memory is conveyed to the rest of the brain by the lowered activity of the downstream MBON. But the memory itself, which is stored by depressed KC to MBON synapses, is not extinguished (Takemura et al., 2017; Schleyer et al., 2020). This observation raises the possibility that MBON-DAN feedback may play a role in regulating memory expression (Krashes et al., 2009; Senapati et al., 2019).

### MBON-MBON interactions

Feedforward MBON-MBON connections were postulated, based on behavioral and light microscopic anatomical observations, to propagate local plasticity between compartments (Aso 2014a, b). The GABAergic MBON11 is poised to play a key role in such propagation as it is connected with 17 other MBONs (at a threshold of 10 synapses). Thus local depression of KC-MBON11 synapses by the shock responsive PPL1-γ1pedc DAN would be expected to result in disinhibition of MBONs in other compartments. Indeed enhanced CS+ responses of MBON01 and MBON03 after odor-shock conditioning have been ascribed to release from MBON11 inhibition (Owald et al., 2015; Perisse et al., 2016; Felsenberg et al., 2018). The consequence of releasing other strongly connected MBONs (MBON07, MBON14 and MBON29) from MBON11 inhibition awaits future study. As discussed above, MBON11 is also connected to DANs innervating its cognate compartment and to DANs innervating other compartments. MBON11 may therefore coordinate MBON network activity via both direct and indirect mechanisms. We also found analogous feedforward inhibitory connections from MBON09 (γ3β′1) to MBON01 and MBON03. Aversive learning will therefore reduce both MBON11- and MBON09-mediated inhibition of these MBONs, which would then further skew the MBON network towards directing avoidance of the previously punished odor (Figure 40).

**Figure 40.**
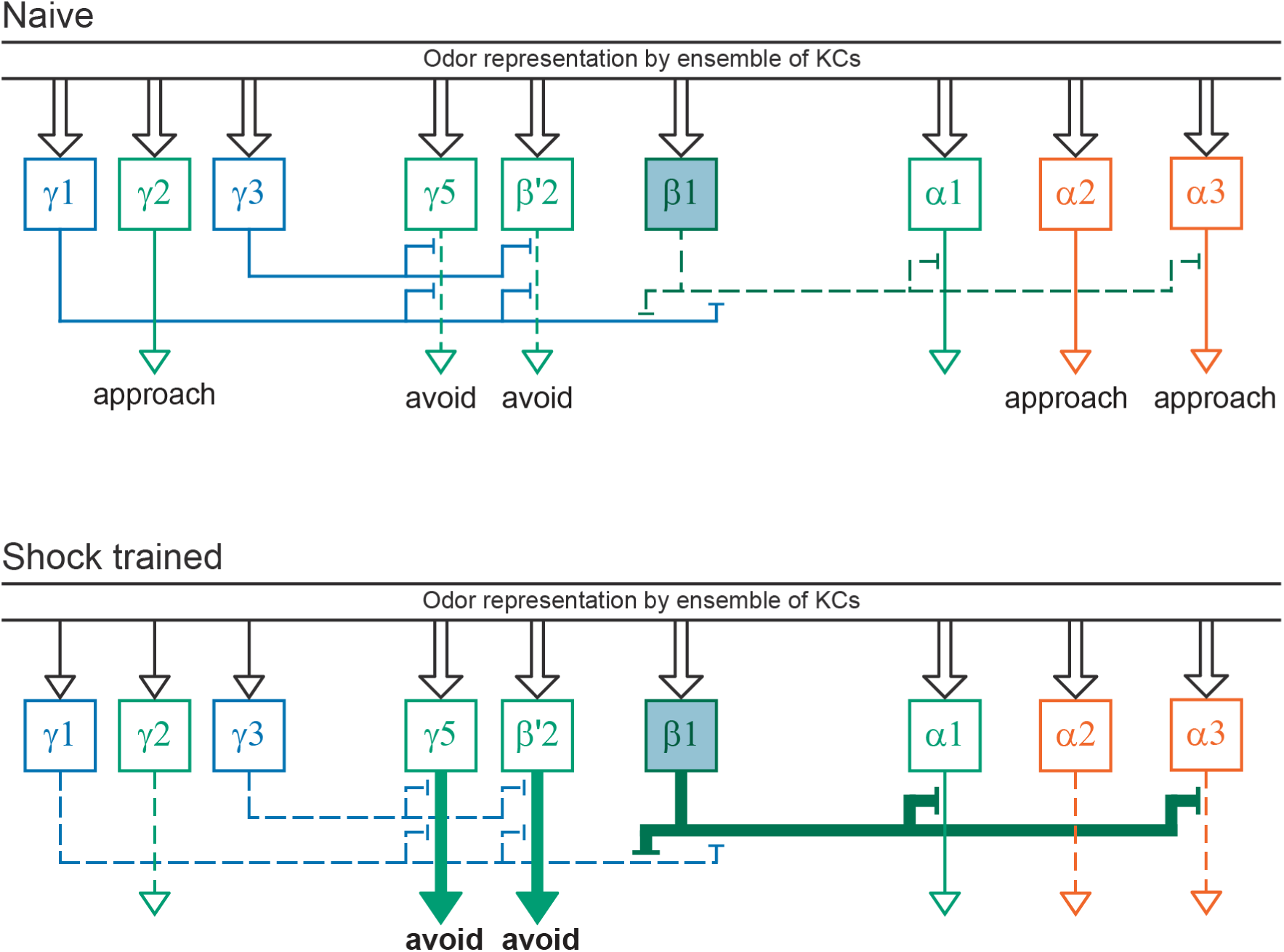
MBON network representation of aversive memory. Shock responsive DANs innervating γ1, γ2 and γ3 depress odor-specific KC synapses onto the respective MBONs. Depression of MBON11 (γ1pedc>α/β) and MBON09 (γ3β′1) releases their feedforward inhibition onto the avoidance-directing γ5 and γ′2 MBONs. In addition, MBON11 release increases the inhibitory effect of the MBON06 (β1>α) onto the approach-directing α1 and α3 compartments. Repetitive aversive training trials, and a release from inhibition also allows the DANs in α2 and α3 to depress odor-specific KC synapses onto the respective approach directing MBONs. These changes together leave the network in a configuration to direct strong odor-specific avoidance behavior.

Disinhibition likely also plays an important role in appetitive memory (Figure 41). The connectivity of MBON06 (β1>α) revealed here indicates that local plasticity in the β1 compartment can propagate to other MBONs. Similar to MBON11, MBON06 is directly connected with nine other MBONs with a threshold of 10 synapses. MBON06 gradually increases its response to repeated odor exposure (Hattori et al., 2017) and odor-evoked responses of β1 DANs vary with metabolic state (Siju et al. 2020). More compellingly, artificially triggering PAM09 (β1) DANs can assign appetitive valence to odors (Perisse et al., 2013; Huetteroth et al., 2015; Aso and Rubin, 2016). However, the role of MBON06 in appetitive memory and how the PAM09 (β1) DANs modulate KC synapses to MBON06 have not been further investigated. The glutamatergic MBON06 (β1>α) makes a large number of reciprocal axoaxonic connections with the GABAergic MBON11 (Takemura et al., 2017), whose activation favors approach (Aso et al., 2014b; Perisse et al., 2016). This reciprocal network motif and the positive sign of behavior resulting from β1 DAN-driven memory suggests that MBON06 released glutamate is likely to be inhibitory to MBON11 and its other downstream targets. MBON06 also makes twice as many connections onto the glutamatergic MBON07 and the cholinergic MBON14 as does MBON11. MBON14 (α3) and MBON07 (α1) have established roles in appetitive memory (Placais et al., 2013; Huetteroth et a., 2015; Yamagata et al., 2015; Ichinose et al., 2015; Widmer et al., 2018). Assuming that PAM09 (β1) DANs encode appetitive memory by depressing synapses from odor-specific KCs onto MBON06 (β1>α), MBON06 suppression will release the feedforward inhibition of MBON06 onto MBON07 (α1), freeing it to participate in driving the PAM11-aad (α1) DANs (Ichinose et al., 2015). Release of MBON06 inhibition should also simultaneously potentiate the responses of MBON14 (α3) to the conditioned odor (Placais et al., 2013). Lastly, releasing the strong inhibition from MBON06 frees MBON11 to provide weaker inhibition that would further favor odor-driven approach (Perisse et al., 2016; Sayin et al., 2019).

**Figure 41.**
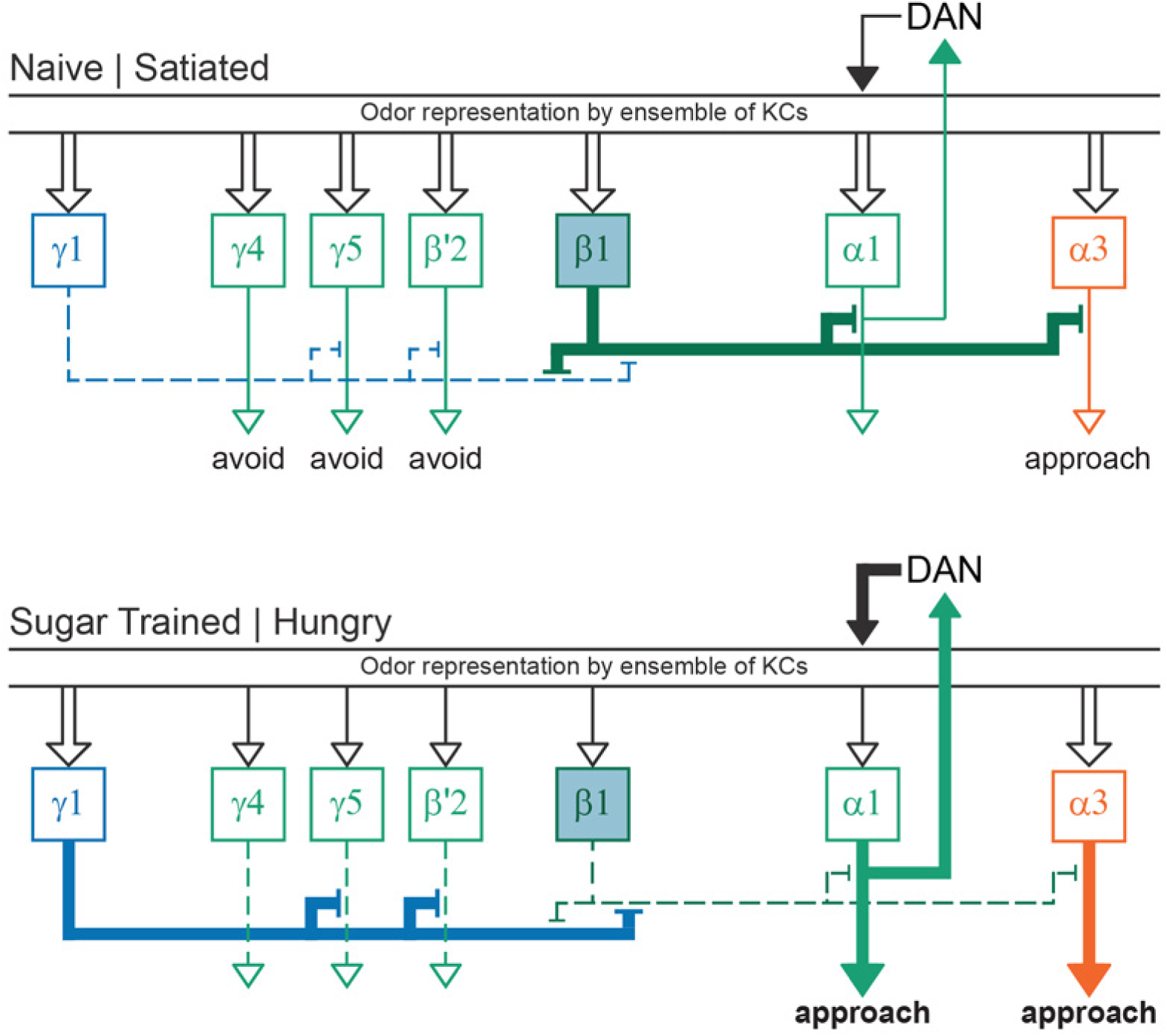
MBON network representation of appetitive memory. Sugar-responsive DANs innervating γ4, γ5, β′2 and β1 depress odor-specific KC synapses onto the respective MBONs. Depression of the odor-specific response of MBON06 (β1>α) reduces odor-specific feedforward inhibition onto the MBON07 (α1) and MBON14 (α3). This frees α3 to direct odor-driven approach behavior and increases activity in a recurrent α1 MBON07-PAM11aad DAN loop to consolidate long-term appetitive memory. MBON11 (γ1pedc>α/β) is sensitive to satiety state and is more responsive when the fly is hungry. Hunger therefore further promotes appetitive memory formation and expression by increasing inhibition onto MBON01 (γ5β′2a) and β′2 MBONs and by further reducing the feedforward inhibitory effects of MBON06 (β1>α) MBON within α1 and α3. These changes together leave the network in a hunger state-dependent configuration that directs strong odor-specific approach behavior.

Aversive learning will also alter the function of the MBON06:MBON11 network motif (Figure 40). Aversive reinforcement through the PPL101 (γ1pedc) DAN depresses KC-MBON11 connections. This depression releases MBON06 from MBON11-mediated suppression and allows MBON06 to then suppress output through MBON07 and MBON11, further favoring odor-driven avoidance.

### The influence of internal state in the network

Several studies have described the influence of internal states such as hunger and thirst on the function and physiology of the MB. In essence, states appear to modulate the DAN-MBON network so that the fly preferentially engages in the pursuit of its greatest need (Senapati et al., 2019). Since our current knowledge suggests these deprivation states employ volume release of modulatory peptides or monoamines to control specific DANs and their downstream MBONs (Krashes et al., 2009; Lin et al., 2014b; Lewis et al., 2015; Tsao et al., 2018; Sayin et al., 2019; Siju et al., 2020), the connectome we have analyzed does not provide a complete description of this circuit. Nevertheless, direct connectivity does provide some interesting new insight concerning hunger (Figure 41) and thirst-dependent control.

The MBON06 (β1>α):MBON11 (γ1pedc>α/β) cross-inhibitory network motif is likely to be relevant for the dependence of learning and memory expression on hunger state (Figure 41). PPL101 (γ1pedc) DANs and MBON11 (γ1pedc>α/β) are sensitive to nutrient/satiety status, with MBON11 being more responsive in hungry flies (Krashes et al., 2009; Placais and Preat, 2013; Perisse et al., 2016; Pavlowsky et al., 2018). Therefore, the hungry state favors the activity of the MBONs that are normally repressed by MBON06.

Thirst-dependent seeking of water vapor requires the activity of DANs innervating the β′2 compartment (Lin et al., 2014b). Our current work shows that these DANs are likely to directly modulate thermo/hygrosensory KCs. In addition, a recent study showed that thirst-dependent expression of water memory required peptidergic suppression of the activity of both the PPL103-γ2α′1 and PAM02 (β′2a) DANs (Senapati et al., 2019). Interestingly, blocking the PAM02 (β′2a) DANs released memory expression in water-sated flies whereas blocking the PPL103-γ2α′1 DANs had no effect. However, if the PPL103-γ2α′1 DANs were blocked together with PAM02 (β′2a) they further facilitated water memory expression. The connectome suggests circuit mechanisms that could reconcile these observations: MBON12 (γ2α′1) provides strong cholinergic input to the PAM02 (β′2a) DANs, suggesting that PPL103-γ2α′1 DANs might facilitate water memory expression by suppressing MBON12’s excitatory input onto PAM02 (β′2a) DANs.

### A possible role for MB output in the control of movement

A role for the MB in guiding locomotion and navigation in ants and other insects has been proposed (Ardin et al., 2016; Collett and Collett, 2018; Kamhi et al., 2020; Kim et al., 2019; Mizunami et al., 1998; Le Möel F, Wystrach A. 2020; Paulk and Gronenberg, 2008; Sun et al., 2020). The strong and direct connections we observed from the majority of MBONs to the CX, the fly’s navigation and locomotion control center, provide one circuit path for the MB to exert influence on motor behaviors. Discovering how this input is utilized by the CX will require additional experimental work.

Optogenetic activation of *Drosophila* MBONs can promote attraction or avoidance by influencing turning at the border between regions with and without stimulating light (Aso et al 2014b). The effect of MBON activation is additive: coactivation of positive-valence MBONs produced stronger attraction, whereas coactivation of positive and negative-valence MBONs cancelled each other out. Because the fly needs to balance the outputs of different compartments, we expect that those downstream neurons that integrate inputs from multiple MBONs will have a privileged role in motor control.

The activity of some DANs has been shown to correlate with motor activity (Berry et al., 2015; Cohn et al, 2015), and the optogenetic activation of PAM-β′2 or PPL1-α3 DANs can attract flies, indicating that DAN activity can itself in some circumstances drive motor behavior. The circuit mechanisms generating the correlation between DAN activity and motor behavior remain to be discovered. Downstream targets of MBONs provide extensive input to DANs, and we found that neurons downstream of multiple MBONs are twice as likely as other MBON targets to provide such direct input to DANs.

We also discovered a direct pathway mediated by atypical MBONs that connects to the descending neurons that control turning, an observation that provides additional support for the importance of the MB in the control of movement. The atypical MBONs that connect directly to the descending steering system, MBON26, MBON27, MBON31 and MBON32, appear to be among the most integrative neurons in the MB system in the sense that they combine direct KC input from the MB compartments with both input from many other MBONs and non-MB input. At the level of the descending neurons, the highly processed signals from these MBONs are combined with inputs from many other sources, including the central complex, to affect a decision to turn. This high degree of integration presumably reflects the complexity and importance of this decision, with many factors involved that might act individually or in combination.

Visual input to the MB is over-represented in the output to the descending neurons, predominantly through MBON27. Short- and long-term learning based on features in a visual scene has been reported to involve the CX (Liu et al., 2006; Neuser et al., 2008). Plasticity in the CX enables visual feature input from the sky and surrounding scenery to be mapped flexibly onto the fly’s internal compass (Fisher et al., 2019; Kim et al., 2019). The visual input conveyed to the MB and, presumably, the learning at the synapses between visual KCs and MBON27, may be of lower resolution, encoding broader features such as color and contrast (Guo, 2005; Tang, 2001; Zhang et al., 2007). An early study demonstrated that the MB is dispensable for flying *Drosophila* to learn shapes but that it is required for them to generalize their learning if the visual context changes between training and testing (Liu et al., 1999). Memory of visual features and the ability to generalize context could allow visual landmarks to help guide navigation either through the CX or by directly influencing descending neurons. The thermo/hygrosensory features conveyed by MBON26 could play a similar role, as could the large amount of odor-related information present in this MBON output pathway.

## Concluding remarks

The MB has an evolutionarily-conserved circuit architecture and uses evolutionarily-conserved molecular mechanisms of synaptic plasticity. The dense connectome analysed in this report has uncovered many unanticipated circuit motifs and suggested potential circuit mechanisms that now need to be explored experimentally. In *Drosophila,* we currently have access to many of the required tools such as cell-type-specific driver lines, genetically encoded sensors and microscopy methods to observe whole brain neuronal activity and fine ultrastructure. These features make the fly an excellent system in which to study many general issues in neuroscience, including: the functional diversity of dopaminergic neurons that carry distributed reinforcement signals, the interactions between parallel memory systems, and memory-guided action selection, as well as the mechanisms underlying cell-type-specific plasticity rules, memory consolidation, and the influence of internal state. We expect studies of the MB to provide insight into general principles used for cognitive functions across the animal kingdom.

As mentioned in the introduction, the MB shares many features with the vertebrate cerebellum, and our results should be informative for studies of the cerebellum proper as well as other cerebellum-like structures such as the dorsal cochlear nucleus and the electrosensory lobe of electric fish. A distinctive feature of these systems, and of the MB, is that learning is driven by a particular mechanism; for example DAN modulation in the MB or complex spiking driven by climbing fiber input in the cerebellum. Studies of learning in cortical circuits have traditionally focused on Hebbian forms of learning driven by the ongoing input and output activity of a neuron. However, recent results from both hippocampal (Bittner et al., 2017) and cortical (Gambino et al., 2014; Lacefield et al., 2019; Larkum, 2013) circuits have stressed the importance of plasticity that is driven by dendritic plateau potentials or bursts that resemble the distinct learning events seen in cerebellar and MB circuits. Thus, the form of plasticity seen in the MB and its control by output and modulatory circuits may inform studies of learning in the cerebral cortex as well.

## Materials and Methods

### Connectivity and Morphological Data

The analyses reported in this paper were all based on the hemibrain:v1.1 dataset (Scheffer et al., 2020). Analyses were done by querying this dataset using the neuPrint user interface or, for more complex queries, by directly querying the neuPrint backend Neo4J graph database (Clements et al., 2020). In some analyses and diagrams, a threshold was applied so as to only consider connections representing more than a certain number or percentage of synapses. Any such thresholds are stated in the figure legends; otherwise, all connections were included in the analysis. Each individual neuron has a unique numerical identifier (generally 8 to 10 digits) that refers to that neuron as seen in this dataset. Neurons were also grouped into putative cell types and given names as described in Scheffer et al. (2020) and we use those v1.1 names here. Even though most of the cell types in the MB were already known, we still found a few new cell types, which we named using established naming schemes. We further refined morphological groupings with relevant information on connectivity (see below).

Each neuron’s unique identifier is permanent, while we expect cell type classifications and neuron names to be continuously updated in response to new biological information. For that reason, we present the unique identifiers for all the neurons we discuss, either by stating them in the main text, figures or figure legends, or by providing links in the figure legends to the appropriate data in neuPrint. The version of the dataset that we used for the analyses reported here will be archived under the name “hemibrain:v1.1” and will remain available in neuPrint, even after additional data releases are made. As described in Scheffer et al., (2020), in addition to the “named neurons” (whose status is indicated as “Traced” in neuPrint) there are also small fragments that, while assigned a unique identifier, are not connected to any named neuron. These small fragments were excluded from our analysis.

### Morphological Clustering

Our overall method of cell type classification is described in Scheffer et al., (2020) and was applied to generate Figures 3 and 6.

Kenyon Cell Morphological Clustering. Analysis of Kenyon cell morphologies (Figures 4 and 5) was carried out in R using the <u>natverse</u> toolkit (Bates et al., 2020a). Briefly, skeletons were downloaded from neuprint.janelia.org using the <u>neuprint_read_neurons</u> function from the <u>neuprintr</u> package, which healed any gaps in the skeleton and rerooted on the soma. When required, neurons were simplified to one major branch using <u>nat::simplify_neuron</u>. Prior to NBLAST, neurons were scaled to units of microns, resampled to 1 µm step size and converted to <u>dotprops</u> format with k=5 neighbors. In order to reduce the weight given to the many fine branches present in EM reconstructions, <u>nat::prune_twigs</u> was applied with twig_length=2. Standard NBLAST clustering was then carried out using the <u>nat.nblast</u> package as previously described (Costa et al., 2016). For KC cell typing, we carried out a stepwise manual NBLAST clustering followed by manual review, which resulted in 17 reassignments. These manual reassignments were almost exclusively cases in which a KC more closely resembled one KCα/β subtype in the vertical lobe and either the subsequent or previously generated subtype in the medial lobe, possibly because it was born during the developmental transition from generation of KCα/βs to KCα/βm or from KCα/βm to KCα/βc.

Morphological Clustering of DANs. DAN cell types reported in the hemibrain:v1.1 were further divided by morphological clustering into subtypes using NBLAST and manual inspection as described in Otto et al., (2020). To generate the results presented in Figures 28, 30 and 31, the same steps as described above, but no simplification was carried out, and nat::prune_twigs was applied with twig_length=5 - a combination that was manually reviewed to preserve subtle differences in dendrites and axons. For Figures 28 and Figure 28-figure supplement 1, hierarchical clustering was performed on DANs of each type/compartment using a wrapper function for base R clustering functions with nat.nblast (nat.blast::nhclust), taking Euclidean distance matrices of similarity scores, with average linkage clustering criterion. The number of clusters and the content was manually reviewed.

Clustering by morphology of the top dendritic inputs to each DAN subtype (Figures 31-37) was performed as follows: The 50 input neurons with the most input synapses to each DAN subtype was performed using hierarchical clustering with average linkage criterion on all-by-all NBLAST similarity scores, which were obtained as described above. In addition, a multi-step approach was needed because of the morphological diversity of the upstream population (see Otto et al., 2020). Only designated right hemisphere dendritic input neurons were considered; MB intrinsic neurons (MBONs, KCs, DPM, and APL) and unnamed neuron fragments were excluded. Before clustering, neurons were simplified by nat:: prune_twigs with twig length = 5. Input neurons were initially split into 25 coarse clusters, largely representative of neuropil of origin. These primary clusters were then individually sub-clustered to yield 235 clusters using an iterative manual review process, taking into account within cluster differences in connectivity. Thirty-five representative clusters were selected and used for the analyses shown in Figures 31 - 37.

### Connectivity analysis

Connectivity-based clustering: Neurons with practically indistinguishable shapes but with different connectivity patterns can often be split into connectivity subtypes within a morphology type. To generate the cell type assignments reported in Scheffer et al., (2020) we made extensive use of a tool for cell type clustering based on neuron connectivity, called CBLAST. In an iterative process, using neuron morphology as a template, neurons were regrouped after more careful examination of neuron projection patterns and their connections. This is especially useful in the case of a dataset like ours, in which noise and missing data make it difficult to rely solely on connectivity to find a good partitioning automatically. CBLAST clusters neurons together using a similarity feature score defined by how the neuron distributes inputs and outputs to different neuron types. In some cases, this readily exposes incompleteness (for example, due to the finite size of the volume) in some neurons. Based on these interactions, we made decisions and refined the clusters manually, iterating until further changes are not observed. CBLAST usually generates clusters that are consistent with the morphological groupings of the neurons, with CBLAST often suggesting new sub-groupings as intended.

In a number of instances we obtained high-level groupings of neurons based on their input or output connectivity and without regard to morphology: Figure 10-figure supplement 3; Figure 11-figure supplements 2, 4 and 5; Figure 13-figure supplements 1 and 2; Figure 16; Figure 16-figure supplements 1 and 3; Figure 17; Figure 17-figure supplements 1; Figure 27; Figure 27-figure supplements 1 and 2. This was achieved by performing spectral clustering (Shi and Malik, 2000) on the input (or output) connectivity of a set of neurons using cosine similarity as the clustering metric. Spectral clustering is particularly appropriate in cases where there is no clear hierarchical structure to the data but there are clearly defined groupings. Spectral clustering requires specifying the number of clusters in advance -- in some cases, we specified this value manually, and in others, we automatically determined it by choosing the value that yielded the optimal silhouette score, a measure of clustering quality. All other parameters of the implementation were taken from the default values used by the SpectralClustering method in Python’s scikit-learn package.

For the hierarchical clustering by input connectivity of PAM DANs shown in Figure 28, KC-DAN connectivity was excluded to prevent bias. Before clustering the number of synapses between input neurons and each DAN was normalized by the remaining input to that DAN (minus KC input). Hierarchical clustering with the r-base function hclust was performed for DANs of each type/compartment using the Manhattan distance between upstream connectivity profiles of DANs with Ward’s clustering criterion.

For determination of the DAN to KC and DAN to MBON connectivity shown in Figure 30 and Figure 30-figure supplements 2 and 3, synapse counts between subtypes of DANs and subclasses of KCs were normalized by the total synapse count between DANs and that KC type, and then thresholded at 0.5%. Synapse counts between DAN subtypes of the right hemisphere and MBONs with dendrites in the right hemisphere were normalized by the total number of synapses connections made to that MBON type, and then thresholded at 0.1%.

For producing the connectivity data shown in Figures 31-37, connectivity information was retrieved from neuPrint with the neuprintr function neuprint_connection_table (natverse) for each morphological cluster of upstream neurons to each DAN subtype innervating the right MB. Only synapses in the right hemisphere were used due to incomplete connectivity in the left hemisphere and to prevent bias between PPL and PAM DANs. Connectivity to DAN subtypes was thresholded as indicated in the figure legends.

Comparison of clustering based on morphology and connectivity (Figure 28): Tanglegrams were generated to facilitate visual comparison of dendrograms of morphology- and connectivity-based clustering using the tanglegram function of the dendextend package (see Otto et al., 2020). Dendrogram layouts were determined to minimize edge crossing using dendextend::untangle with <u>method=“step2side</u>” (Galili, 2015). The Mantel test (Legendre and Legendre 2012) implemented in vegan::mantel (https://github.com/vegandevs/vegan) was used to evaluate the similarity of morphology- and connectivity-based clustering. Pearson’s correlation between the distance matrices of these two observed datasets was calculated, then one of the matrices was shuffled all possible ways or at least 10^7^ times and each event tested for correlation with the observed data. The number of events where the correlation was higher than between the two original datasets was divided by the amount of comparisons to create a p-value. When p-values were lower than the significance level, the null model of independence between the two feature spaces was rejected.

### Calculation of multi-step effective connectivity

In several analyses, we computed the “effective” connectivity through multi-synaptic pathways between a set of source and target neurons: Figure 10-figure supplement 2; Figure 11-figure supplement 1; Figure 26-figure supplement 1 and 2; Figure 27-figure supplement 3. Although our procedure generalizes to pathways of any length, we only performed it for two-step (or “one-hop”) pathways. To do so, we determined the set of interneurons either postsynaptic to the source population or presynaptic to the target population. Starting with the matrices of source-interneuron connectivity and interneuron-target connectivity, we normalized each so that the sum of inputs to each postsynaptic cell summed to 1. Then we multiplied the two matrices to yield an estimate of effective source-target connectivity. This procedure reflects the assumption that an output synapse from an interneuron conveys information about its inputs to varying degrees, which are proportional to the number of input synapses coming from each input.

### Data presentation

The 3D renderings of neurons presented in the Figures were generated using the visualization tools of NeuTu (Zhao et al., 2018a); grey-scale images of EM data were taken from NeuTu. Annotations were added using Adobe Illustrator. Color depth MIP masks of MCFO or FAFB skeletons were generated using the ColorMIP_Mask_Search plugin for Fiji (https://github.com/JaneliaSciComp/ColorMIP_Mask_Search) (Otsuna et al., 2018). Cytoscape (cytoscape.org) was used to produce the node layout of connectivity diagrams of connections between neurons, which were then edited in Adobe Illustrator. Videos were produced using Blender (blender.org) and a set of custom Python scripts written by Philip Hubbard (Hubbard, 2020). Narration was recorded using Camtasia (techsmith.com) and text and narration were added to videos using Adobe Premiere Pro.

## Acknowledgements

This work was supported by the Howard Hughes Medical Institute, by the MRC (MC-U105188491 to GSXEJ), by the Wellcome Trust (203261/Z/16/Z to GSXEJ, GMR and SW), by the Simons Foundation (Simons Collaboration on the Global Brain to JL, YA, LFA, AL-K and GMR), by the NSF (NeuroNex Award to LFA, AL-K and JL) and by a Department of Energy Computational Science Graduate Fellowship (DE-SC0020347) to JL. We thank Tanya Wolff for her help in the analysis of MBON to CX connections. We thank Tanya Wolff and Vivek Jayaramn for detailed comments on the manuscript. Kazunori Shinomiya for help generating Figure 2-figure supplement 1. Philip Hubbard for providing software and advice on video production and for generating segments of videos 21, 22 and 26. Emily Joyce for narrating videos. Konrad Heinz and Joseph Hsu provided proofreading assistance and the The Janelia FlyLight project team provided the images used in LM-EM comparisons (Figure 10-figure supplement 1).

---

**Figure 2-figure supplement 1.**
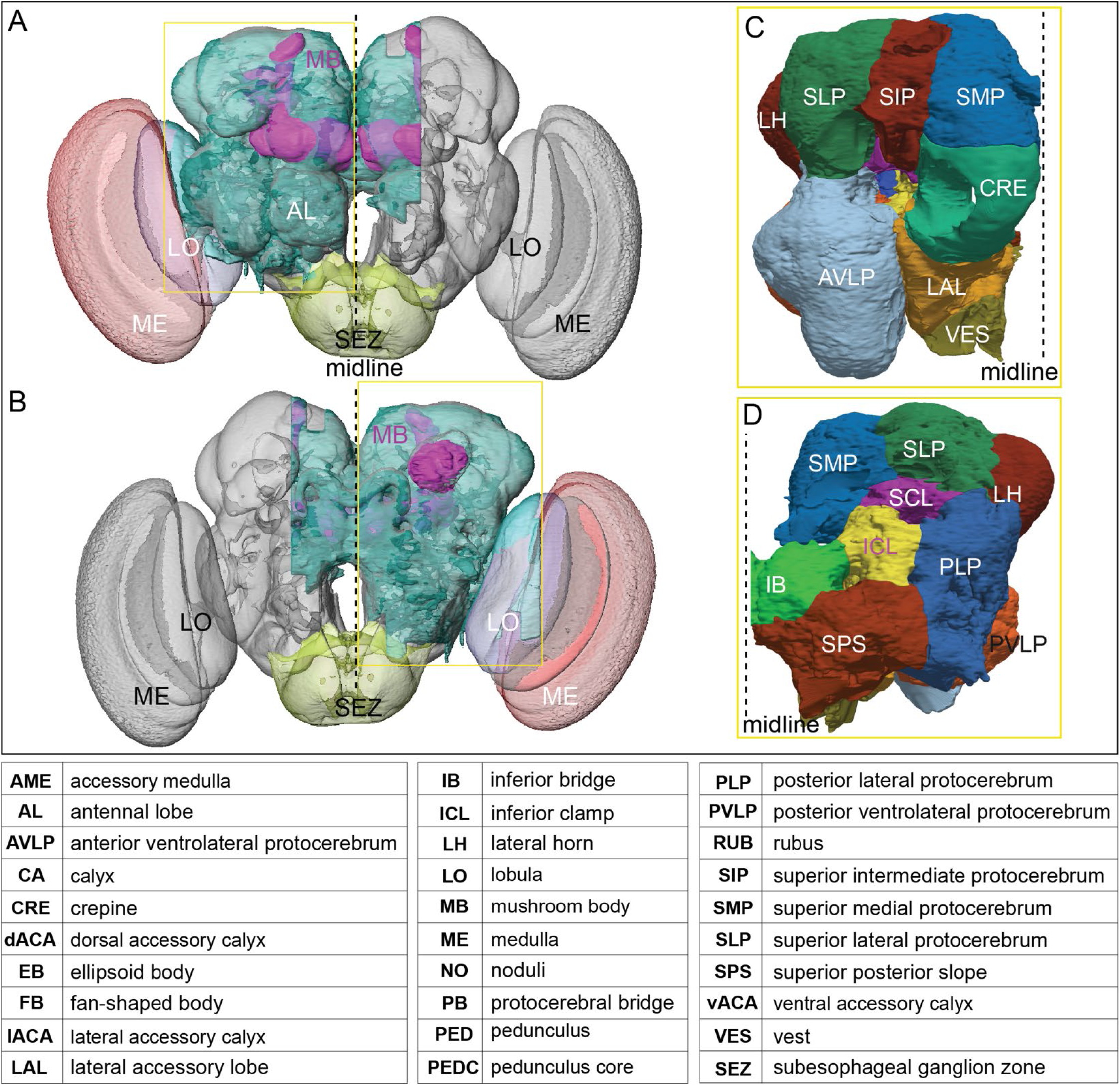
The extent of the hemibrain volume and key to brain area nomenclature. (A) The portion of the central brain (light blue) that was imaged and reconstructed to generate the hemibrain volume (Scheffer et al., 2020) is superimposed on a frontal view of a grayscale representation of the entire *Drosophila* brain (JRC 2018 unisex template; Bogovic et al., 2019). The mushroom body (MB) is shown in purple. The midline is indicated by the dotted black line. The brain areas LO, ME and SEZ, which largely lie outside the hemibrain, are labeled. (B) The same structures viewed from the posterior side of the brain. (C,D) Labeled brain regions in the area indicated by the yellow boxes in panels A and B, respectively. The table shows the abbreviations and full names for brain regions discussed in this paper. See Scheffer et al., (2020) for details.

**Figure 3-figure supplement 1.**
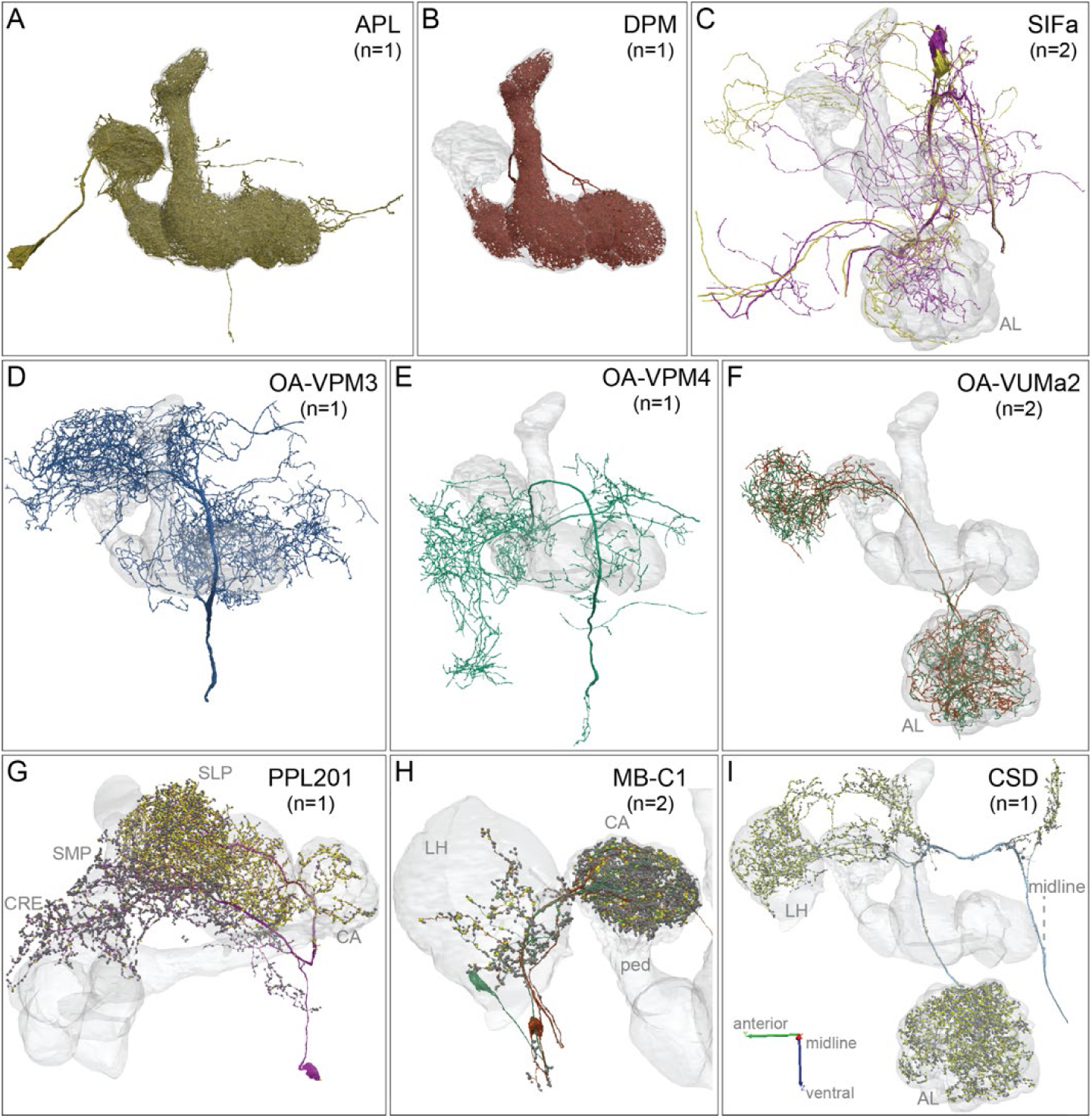
APL, DPM, SIFamide, OA neurons and other modulatory neurons. Neuronal morphologies are shown. (A) The anterior posterior lateral neuron, <u>APL</u>, innervates the entire MB (Video 4). APL is GABAergic and provides negative feedback important for sparse coding of odor identities (Lin et al., 2014a; Liu and Davis, 2009; Papadopoulou et al., 2011; Tanaka et al., 2008). (B) The dorsal paired medial neuron, <u>DPM</u>, innervates the lobes in their entirety and the distal pedunculus (Video 4). The DPM neuron has been proposed to use the neuropeptide amnesiac (Waddell et al., 2000), serotonin (Lee et al., 2011) and GABA (Haynes et al., 2015) as neurotransmitters and has been reported to be gap-junctionally coupled to APL (Wu et al., 2011). DPM has an important role in memory consolidation (Haynes et al., 2015; Keene et al., 2006; Pitman et al., 2011; Yu et al., 2005). The connectivity between DPM and APL and other neurons within the α lobe was described by Takemura et al. (2017), and our observations for the full MB are consistent with that description. (C) Two neurons that express the neuropeptide SIFamide, <u>SIFa</u>, are shown here and in Video 5 (Park et al., 2014; Verleyen et al., 2004). (D-F) Three types of neurons that express the neurotransmitter octopamine (OA) are shown here and in Video 5 (Busch et al 2009; Aso et al. 2014a): (D) <u>OA-VPM3</u>, (E) <u>OA-VPM4</u> and (F) <u>OA-VUMa2</u>. (G) <u>PPL201</u> is a PPL2ab cluster dopaminergic neuron with axonal terminals in the CA, lateral horn (LH) and SLP; it has also been referred to as PPL2a (Mao and Davis, 2009; Tanaka et al., 2008; Zheng et al., 2018). In the CA, 16% of KCα/β and 25% of KCα*′*/β*′* and KCγm receives synapses from PPL201; however, KCs devoted to visual information are only very sparsely innervated (5% of KCα/βp and 3% of KCγd). The dendrites of this neuron are in the SMP and CRE. (H) Two GABAergic <u>MB-C1</u> neurons (shown in different colors) innervate the LH and CA (Mao and Davis, 2009; Tanaka et al., 2008; Zheng et al., 2018). (I) The serotonergic <u>CSD</u> neuron innervates the AL, LH and CA (Dacks et al., 2009). In panels G-I, yellow dots indicate presynaptic sites (sites where these neurons make synapses onto other neurons) and dark grey dots indicate postsynaptic sites (sites where these neurons receive input from other neurons).

**Figure 4-figure supplement 1.**
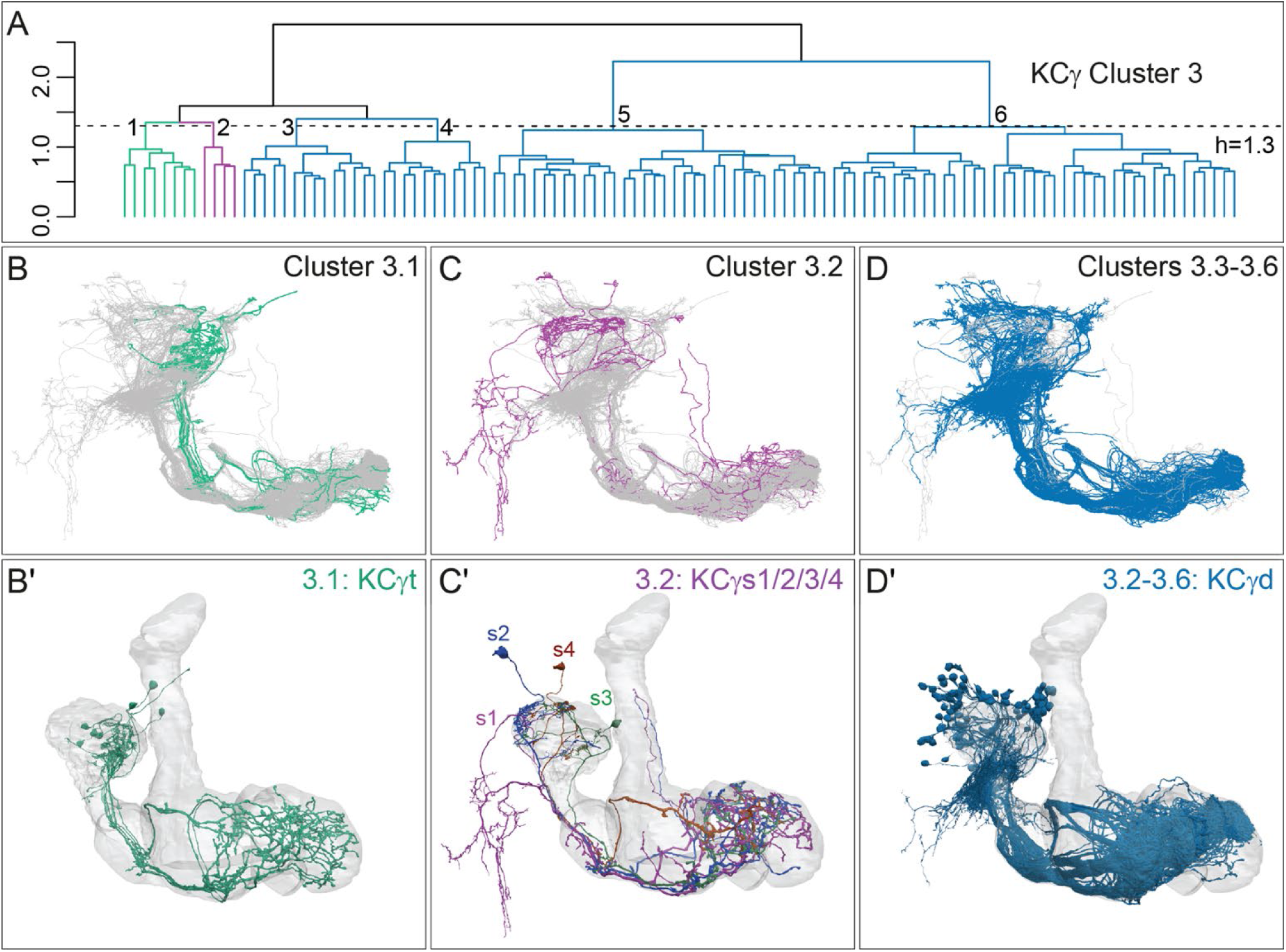
Successive rounds of whole neuron morphological hierarchical clustering reveal novel KCγ subtypes. (A) Morphological hierarchical clustering based on NBLAST scores for all-by-all comparison of KCγ cluster 3 (from Figure 4B) cut at height 1.3 (dashed line); 6 sub-clusters are produced corresponding to 3 neuronal subtypes. (B) KCγt skeletons (cluster 3.1) shown in green, with remaining KCγ found in cluster 3 shown in grey. (B*′*) Space-filling morphologies of the same KCγt with the MB shown in grey. The eight KCγt neurons innervate the anterior of the CA, where most thermo/hygrosensory PNs project. (C) KCγs skeletons in cluster 3.2 shown in magenta, with remaining KCγ found in cluster 3 shown in grey. (C*′*) Space-filling morphologies of the four unique neurons of subtype KCγs, KCγs1/2/3/4, shown in magenta, dark blue, green, rust, respectively with the MB shown in grey. Each KCγs likely derives from a different neuroblast. KCγs1 and KCγs2 both have unusually complex axons that wrap around the surface of the gamma lobe and feature a single dorsal branch; KCγs1 innervates all three accessory calyces and also exhibits a striking dendritic elaboration that extends into the PLP (Figure 10), while KCγs2 has a large dendritic claw in the lateral accessory calyx (lACA) and smaller dendritic branches in the ventroanterior CA (Marin et al., 2020; Figure 12-figure supplement 2). KCγs3 and KCγs4 have axon morphologies more similar to those of canonical KCγ neurons, but KCγs3 receives input in the dorsal accessory calyx (dACA) and ventral accessory calyx (vACA), and KCγs4 receives input in the dACA and anterior CA. (D) KCγd dendrites innervate the vACA and their axons contribute to the dorsal layer of the γ lobe. The 99 KCγd found in sub-clusters 3.3, 3.4, 3.5 and 3.6 are shown in blue; the other γ KCs found in cluster 3 are shown in grey. (D*′*) Space-filling morphologies of the same γd KCs shown in blue, with the MB shown in grey.

**Figure 4-figure supplement 2.**
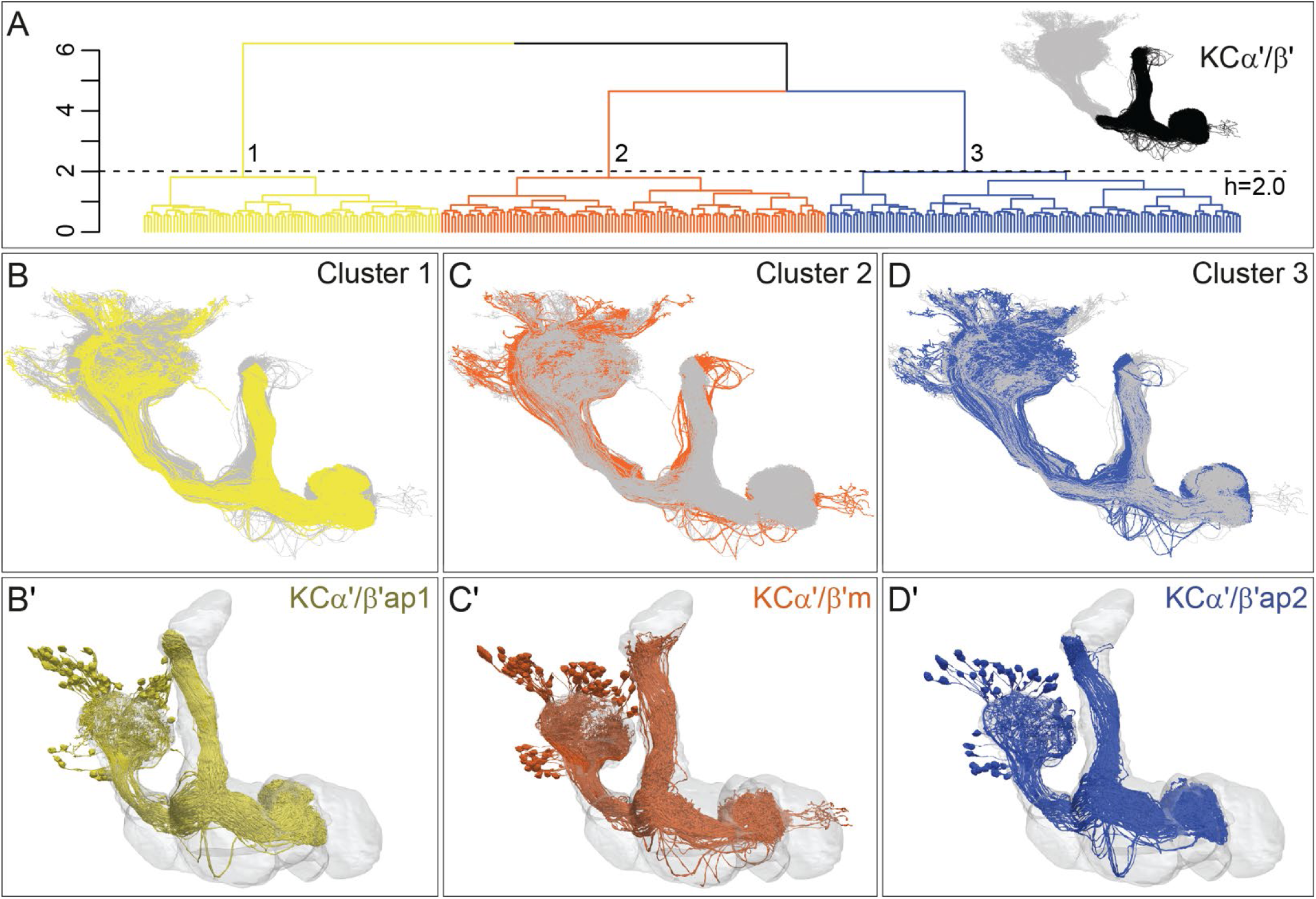
Three distinct morphological subtypes of KCα*′*/β*′*. (A) Morphological hierarchical clustering based on NBLAST scores for all-by-all comparison of α*′*/β*′* KCs (simplified and pruned to include axon lobes only; black area in inset) is shown. When cut at height 2.0 (dashed line), α*′*/β*′* KCs are split into three subtypes. (B) KCα*′*/β*′*ap1 skeletons from cluster 1 are shown in yellow with the remaining α*′*/β*′* KCs shown in grey. (B*′*) Space-filling morphologies of the same KCα*′*/β*′*ap1 are shown in yellow, with the MB shown in grey. (C) KCα*′*/β*′*m skeletons from cluster 2 are shown in rust with the remaining α*′*/β*′* KCs shown in grey. (C*′*) Space-filling morphologies of the same KCα*′*/β*′*m shown in rust, with the MB shown in grey. (D) KCα*′*/β*′*ap2 skeletons from cluster 3 are shown in dark blue with the remaining α*′*/β*′* KCs shown in grey. (D*′*) Space-filling morphologies of the same KCα*′*/β*′*ap2 are shown in dark blue, with the MB shown in grey.

**Figure 4-figure supplement 3.**
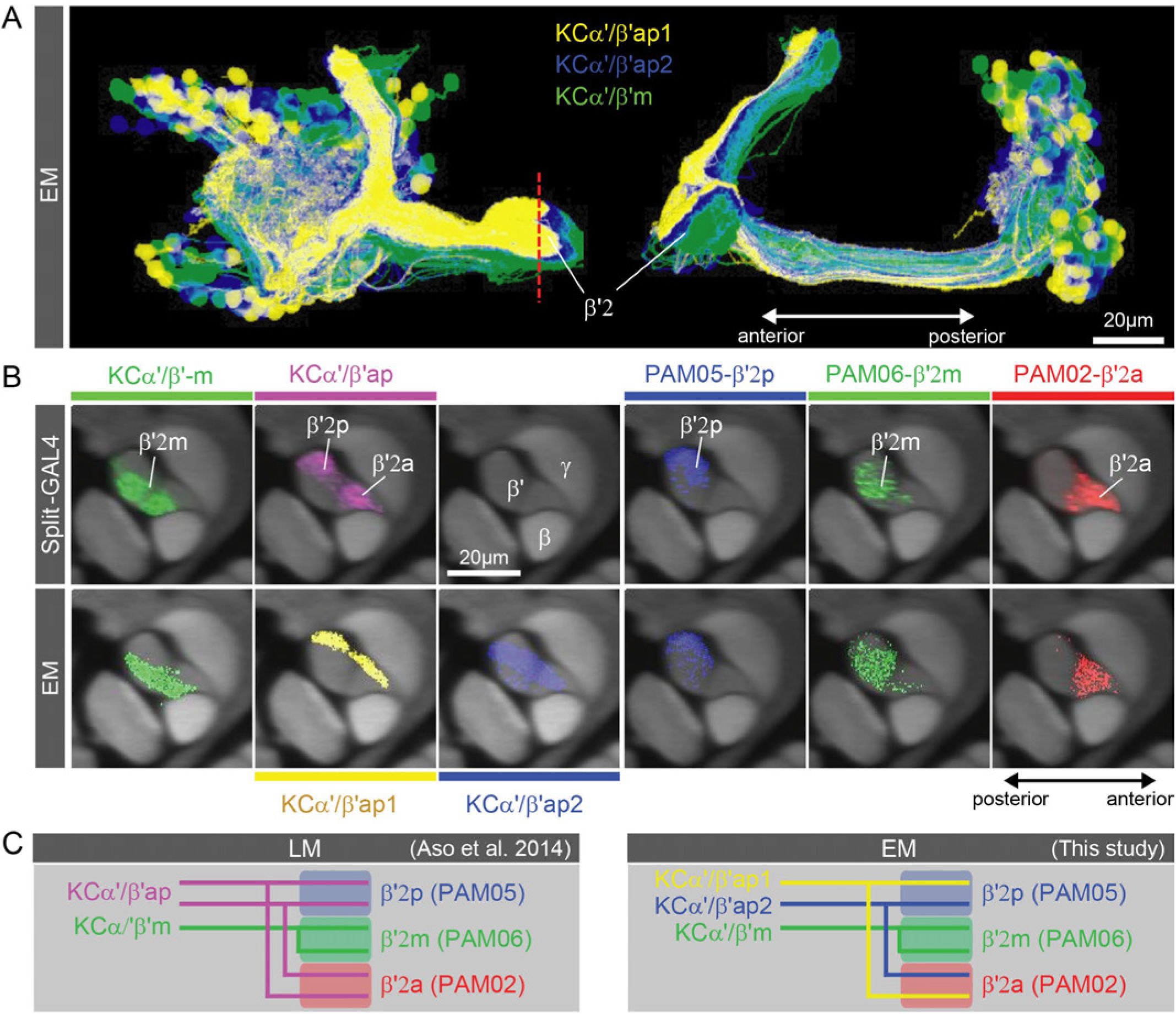
Further explanation of KC subtype nomenclature. (A) Three subtypes of α*′/*β*′* KCs in the hemibrain EM data set are shown in the coordinates of the JF2018 standard brain in frontal (left) and sagittal view (right). The red dashed line indicates the position of the cross section of the MB β*′* lobe shown in (B). (B) Innervation patterns of KCs and PAM DANs in the β*′*2 compartment. The reference neuropil stain from the JF2018 standard brain is shown in all panels in grey. Top: Two classes of KCs and three classes of PAM cluster DANs were defined by split-GAL4 drivers (Aso et al., 2014a) and light microscopic images of those split-GAL4 lines are shown: KCα*′*β*′*m, MB418B; KCα*′*β*′*ap, MB463B; reference neuropil stain; PAM05-β*′*2p, MB056B; PAM06-β*′*2m, MB032B; PAM02-β*′*2a, MB109B. Bottom: The positions of the indicated KC and PAM cell types are based on the hemibrain EM dataset, after transfer into the JF2018 standard brain coordinate system and color-coded. (C) Comparison of LM and EM data sets. Each α*′*β*′*ap bifurcates and innervates both β*′*2a and β*′*2p, coincident with the PAM02 and PAM05 innervation patterns, respectively. Morphological clustering revealed three classes of KCα*′*β*′*, subdividing α*′*β*′*ap into two classes: α*′*β*′*ap1 and α*′*β*′*ap2. KC α*′*β*′*ap1, KCα*′*β*′*ap2 and KCα*′*β*′*m likely correspond to KC α*′*β*′*a, KC α*′*β*′*m and KCα*′*β*′*p, respectively, in Tanaka et al. (2008). However, we followed the nomenclature in Aso et al. (2014a) to minimize confusion with established names based on split-GAL4 driver lines.

**Figure 4-figure supplement 4.**
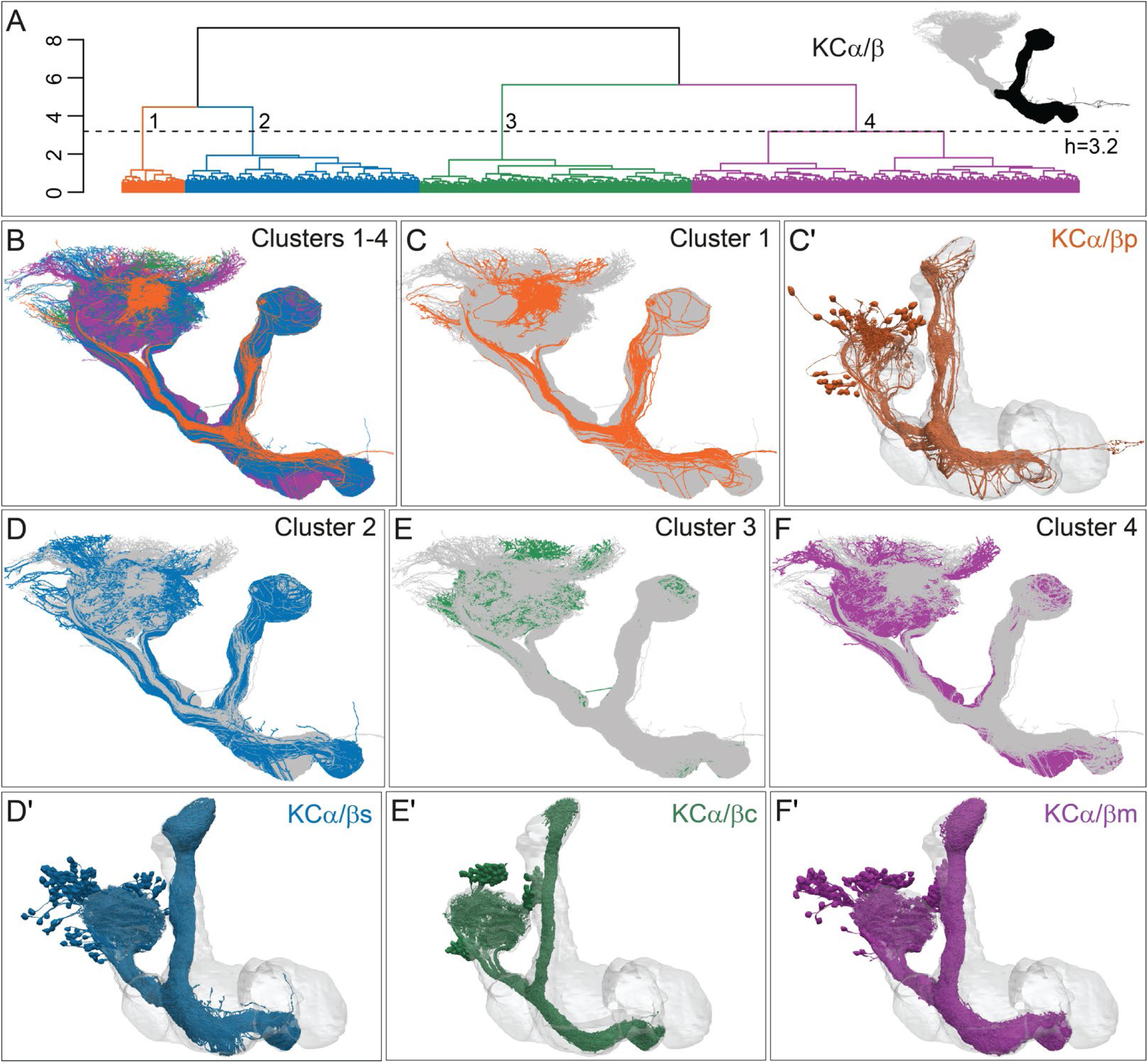
Four distinct morphological subtypes of KCα/β. (A) Morphological hierarchical clustering based on NBLAST scores for all-by-all comparison of KCα/β carried out on that portion of their axons found in the lobes (black area in inset) is shown. When cut at height 3.2 (dashed line), α/β KCs are split into four subtypes. (B) KCα/β skeletons shown color-coded to match the clustering in (A). (C) KCα/βp, cluster 1, shown in rust, with the remaining α/β KCs shown in grey. (C’) Space-filling morphologies of the same KCα/βp shown in rust, with the MB shown in grey. (D) KCα/βs skeletons, cluster 2, shown in blue, with the remaining α/β KCs shown in grey. (D*′*) Space-filling morphologies of the same KCα/βs shown in blue, with the MB shown in grey. (E) KCα/βc skeletons, cluster 3, shown in green, with the remaining α/β KCs shown n grey. (E*′*) Space-filling morphologies of the same KCα/βc shown in green, with the MB shown in grey. (F) KCα/βm skeletons, cluster 4, shown in magenta, with the remaining α/β KCs shown in grey. (F*′*) Space-filling morphologies of the same KCα/βm shown in magenta, with the MB shown in grey.

**Figure 5-figure supplement 1.**
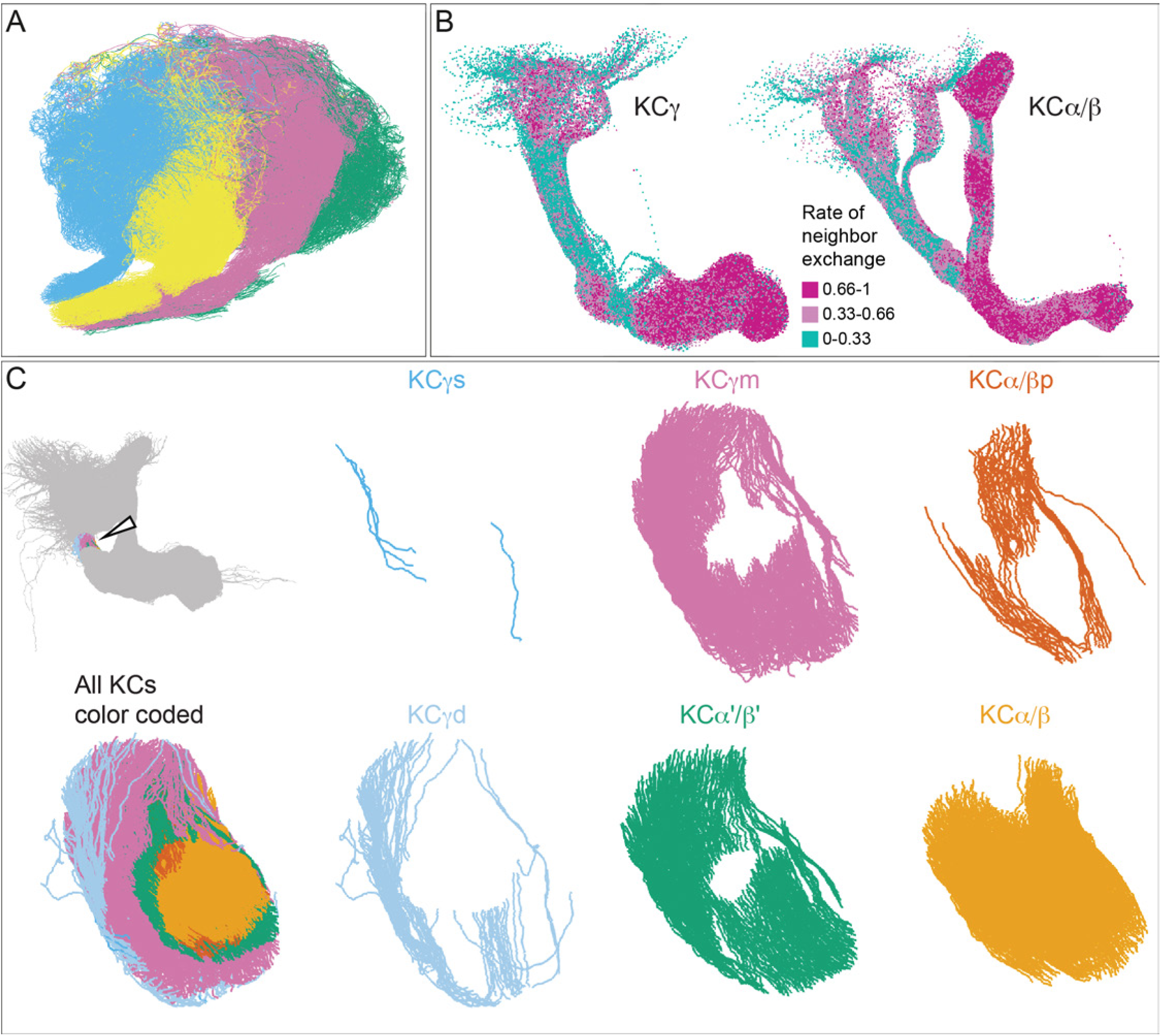
Additional organizational features of KC projections. (A) Posterior view of the CA showing the four clonal units that make up α*′*/β*′* KCs (Figure 5B) as revealed by NBLAST clustering (Figure 5A). (B) Visualization of the rate of change in KC neighbors quantified as the fraction of 10 nearest neighbors that change compared with a position 5μm closer to the soma. Large values imply a rapid change in the neighbors of individual KC fibers, which are observed all the way along γm KCs in the γ lobe; much lower rates of rearrangement are seen in the pedunculus and in segments of the α/β KCs in the vertical lobe. (C) Cross-sections through the mushroom body pedunculus, at the point indicated by the hollow arrowhead, showing the organization of KC types by birth order, with the first-born types (KCγs and KCγd) generally more peripheral and the last-born type (KC α/β) more central.

**Figure 5-figure supplement 2.**
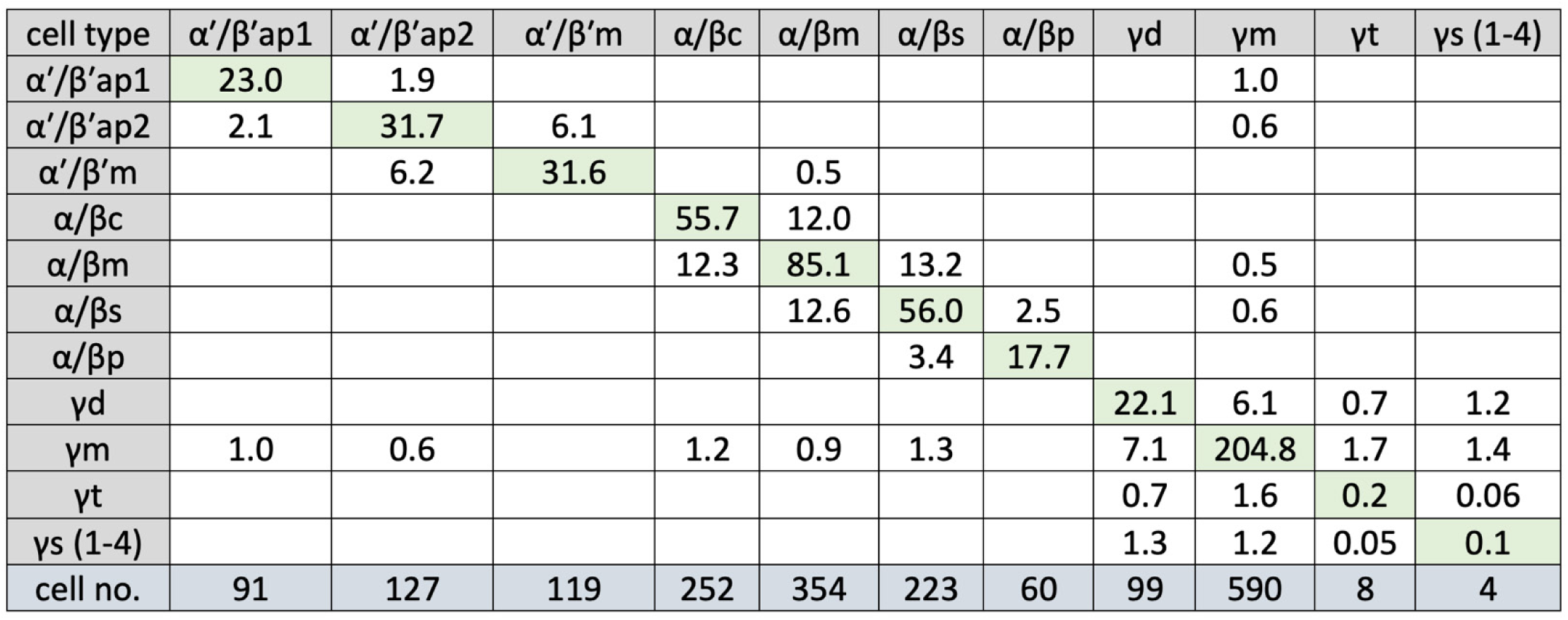
KC-to-KC synapses. The total number of synapses/1000 made between the indicated KC types in the MB (including the lobes, pedunculus and calyces) are shown. Only connections totalling >500 synapses are shown, except for those involving γt and γs where connections greater than 50 synapses were included because of the small numbers of cells of these types. The four γs cell types were pooled. The y-axis shows postsynaptic cell types and the x-axis presynaptic cell types. The number of KCs of each type is shown.

**Figure 6-figure supplement 1.**
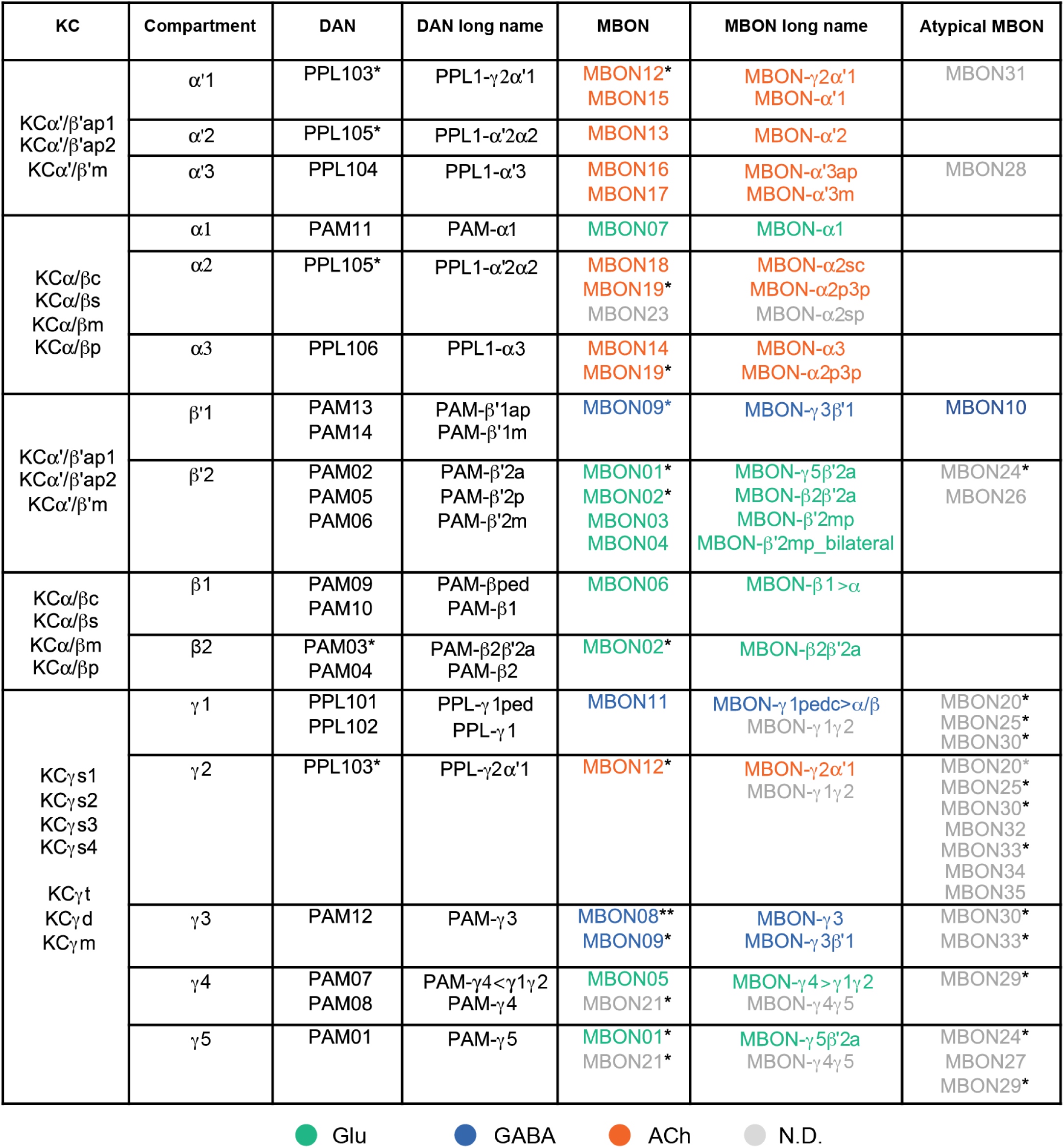
Table of cell types found in each MB compartment. This table shows which KCs, DANs and MBONs are found in each of the 15 compartments of the MB lobes. Both the short names used throughout this paper and the longer names used in Aso et al. (2014a) are shown. MBONs or DANs that innervate more than one compartment are marked with an asterisk. MBON names are color-coded based on neurotransmitter as indicated.

**Figure 8-figure supplement 1.**
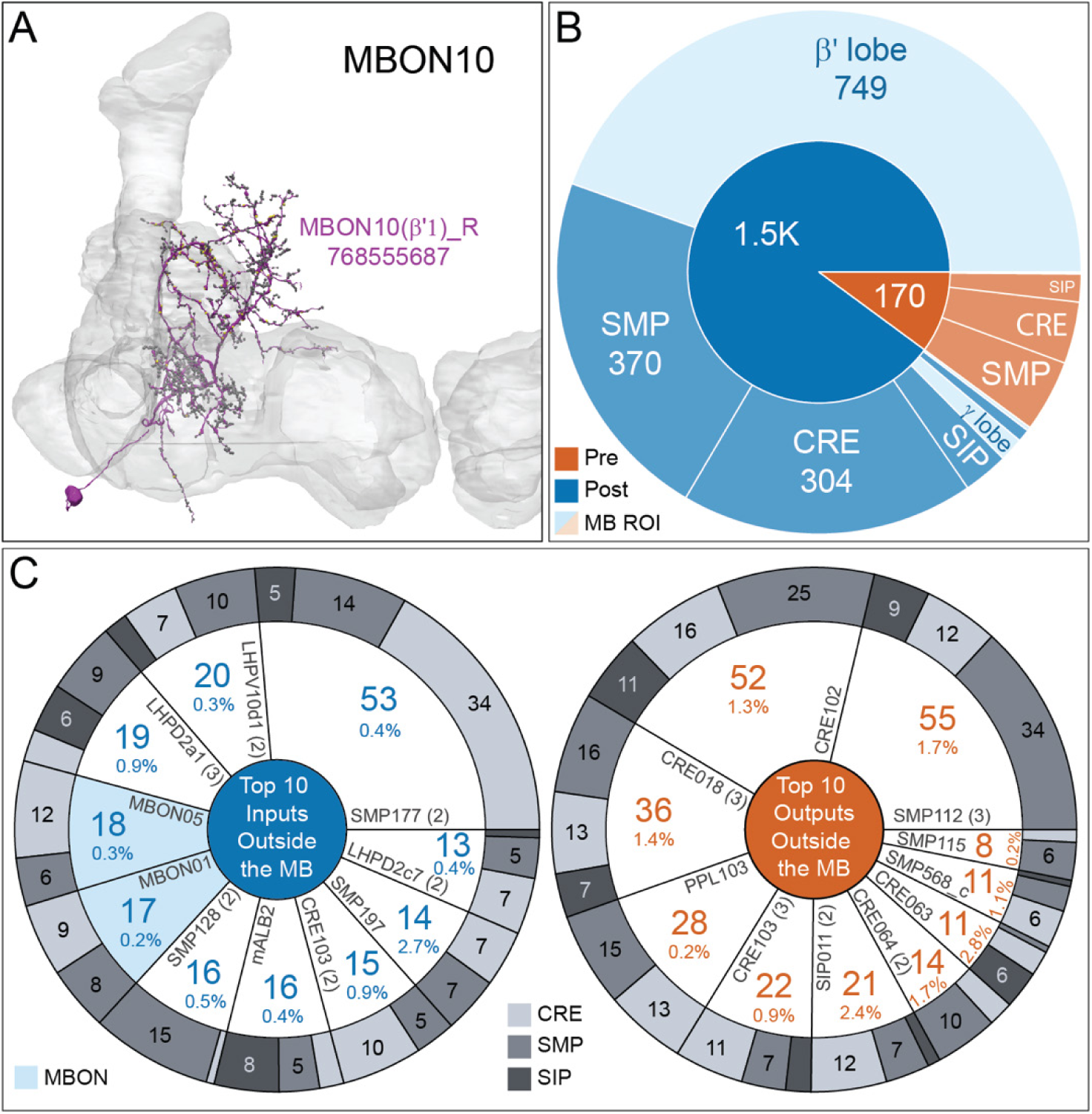
Atypical MBON10. (A) Atypical MBON10 is the only atypical MBON cell type with more than one cell per brain hemisphere. One of the three MBON10s is shown here; all three cells are shown in Video 7. Presynaptic sites are shown in yellow and postsynaptic sites are shown in grey. (B) A pie chart showing the distribution of MBON10’s presynaptic sites (indicated in orange) and postsynaptic sites (in blue). The inner circle shows the total number of MBON10’s pre- and postsynaptic sites; the numbers represent the MBON10 shown in A. The outer circle of the pie chart shows the brain regions in which these sites occur; regions of the MB are shown in lighter colors. (C) MBON10’s top ten inputs by neuron type outside the MB lobes (left) are shown; links to these cell types in neuPrint are: <u>SMP177</u>, <u>LHPV10d1</u>, <u>LHPD2a1</u>, <u>MBON05</u>, <u>MBON01</u>, <u>mALB2</u>, <u>SMP128</u>, <u>CRE103</u>, <u>SMP197</u>, and <u>LHPD2c7</u>. The numbers in each slice represent synapse numbers, while the percentages indicate the percentage of that neuron’s synaptic output that goes to MBON10; the shading of the outer ring of the pie chart indicates where these synapses are located. Note that two typical MBONs synapse onto MBON10, MBON01 (γ5β’2a) and MBON05 (γ4>γ1γ2). MBON10’s top ten outputs by neuron type outside the MB lobes (right) are shown; links to these cell types in neuPrint are: <u>SMP112</u>, <u>CRE102</u>, <u>CRE018</u>, <u>PPL103</u>, <u>CRE103</u>, <u>SIP011</u>, <u>CRE064</u>, <u>SMP568_c</u>, <u>CRE063</u>, and <u>SMP115</u>. The numbers in each slice represent synapse numbers, while the percentages indicate the fraction of that neurons input that is provided by MBON10; the shading of the outer ring of the pie chart indicates where these synapses are located.

**Figure 8-figure supplement 2.**
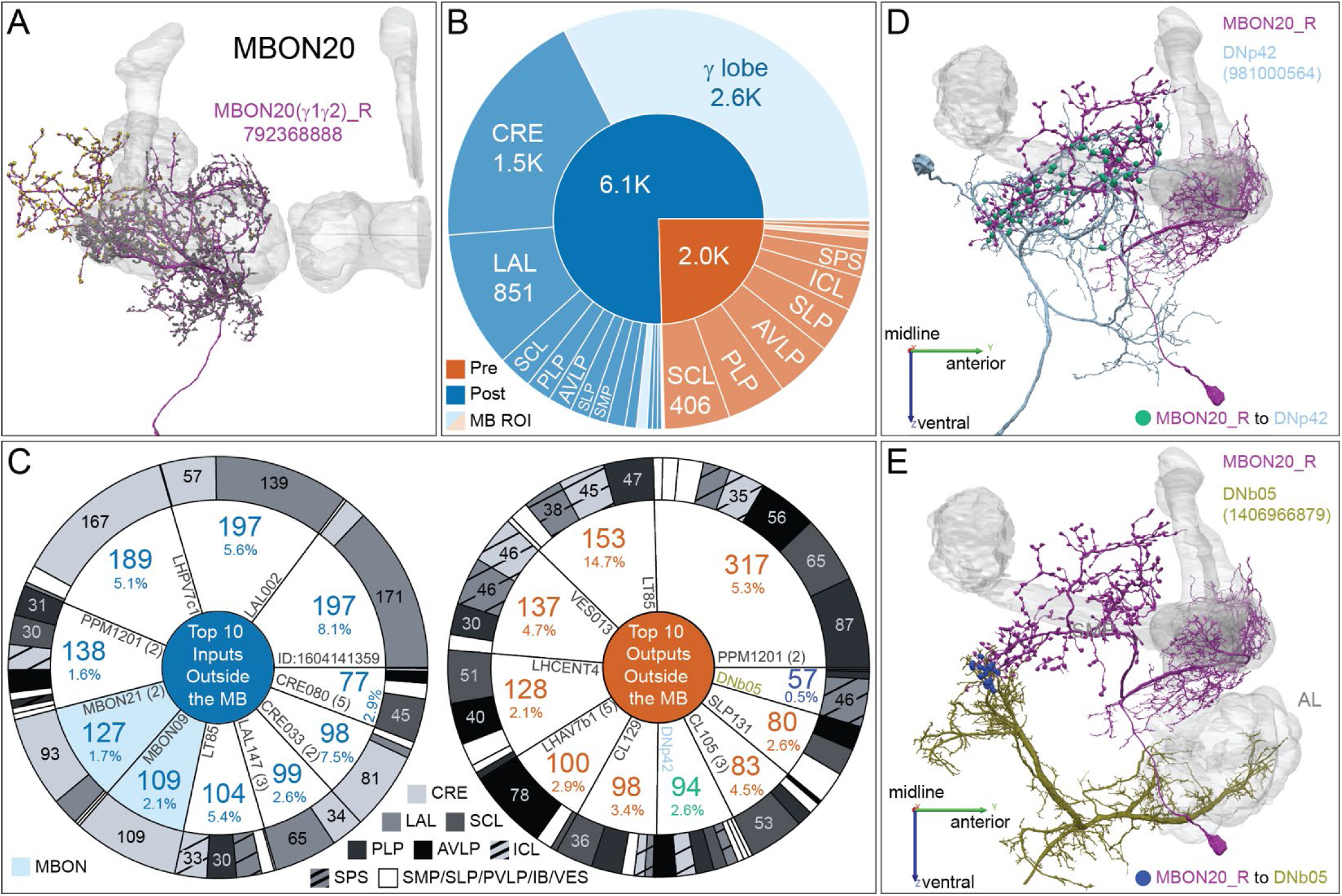
MBON20. (A) Atypical MBON20 (γ1γ2) is shown here and in Video 8. Presynaptic sites are shown in yellow and postsynaptic sites are shown in grey. MBON20 receives 45 percent of its input from sites within MB lobes and 85 percent of those input are in the γ1γ2 compartments. (B) A pie chart showing the distribution of MBON20’s presynaptic sites (indicated in orange) and postsynaptic sites (in blue). The inner circle shows the total number of MBON20’s pre- and postsynaptic sites. The outer circle of the pie chart shows the brain regions in which these sites occur; regions of the MB are shown in lighter colors. (C) MBON20’s top ten inputs by neuron type outside the MBON lobes are shown; links to these cell types in neuPrint are: <u>1604141359</u>, <u>LAL002</u>, <u>LHPV7c1</u>, <u>PPM1201</u>, MBON21, MBON09, <u>LT85</u>, <u>LAL147</u>, <u>CRE033</u>, and <u>CRE080</u>. The numbers in each slice represent synapse numbers, while the percentages indicate the percentage of that neuron’s synaptic output that goes to MBON20; the shading of the outer ring of the pie chart indicates where these synapses are located. MBON20’s top ten outputs by neuron type outside the MB lobes (right) are shown; links to these cell types in neuPrint are: <u>PPM1201</u>, <u>LT85</u>, <u>VES013</u>, <u>LHCENT4</u>, <u>LHAV7b1</u>, <u>CL129</u>, <u>DNp42</u>, <u>CL105</u>, <u>SLP131</u>, and <u>DNb05</u>. The numbers in each slice represent synapse numbers, while the percentages indicate the fraction of that neuron’s input that is provided by MBON20; the shading of the outer ring of the pie chart indicates where these synapses are located. Synapse numbers in highlighted text indicate neurons further discussed in panel D and E and are color-coded to match the neurons in these panels. Note MBON20 makes strong reciprocal connections with LT85, which is a LO tangential neuron conveying visual information, and PPM1201. (D-E) Although not all DNs have been identified, there are at least two individual DNs that are strongly connected to MBON20. (D) DNp42, connects to MBON20 with 94 synapses in dorsal brain regions, including SCL and PLP. (E) DNb05, connects to MBON20 with 57 synapses in SPS. Note DNb05 also receives input from PNs in the AL.

**Figure 8-figure supplement 3.**
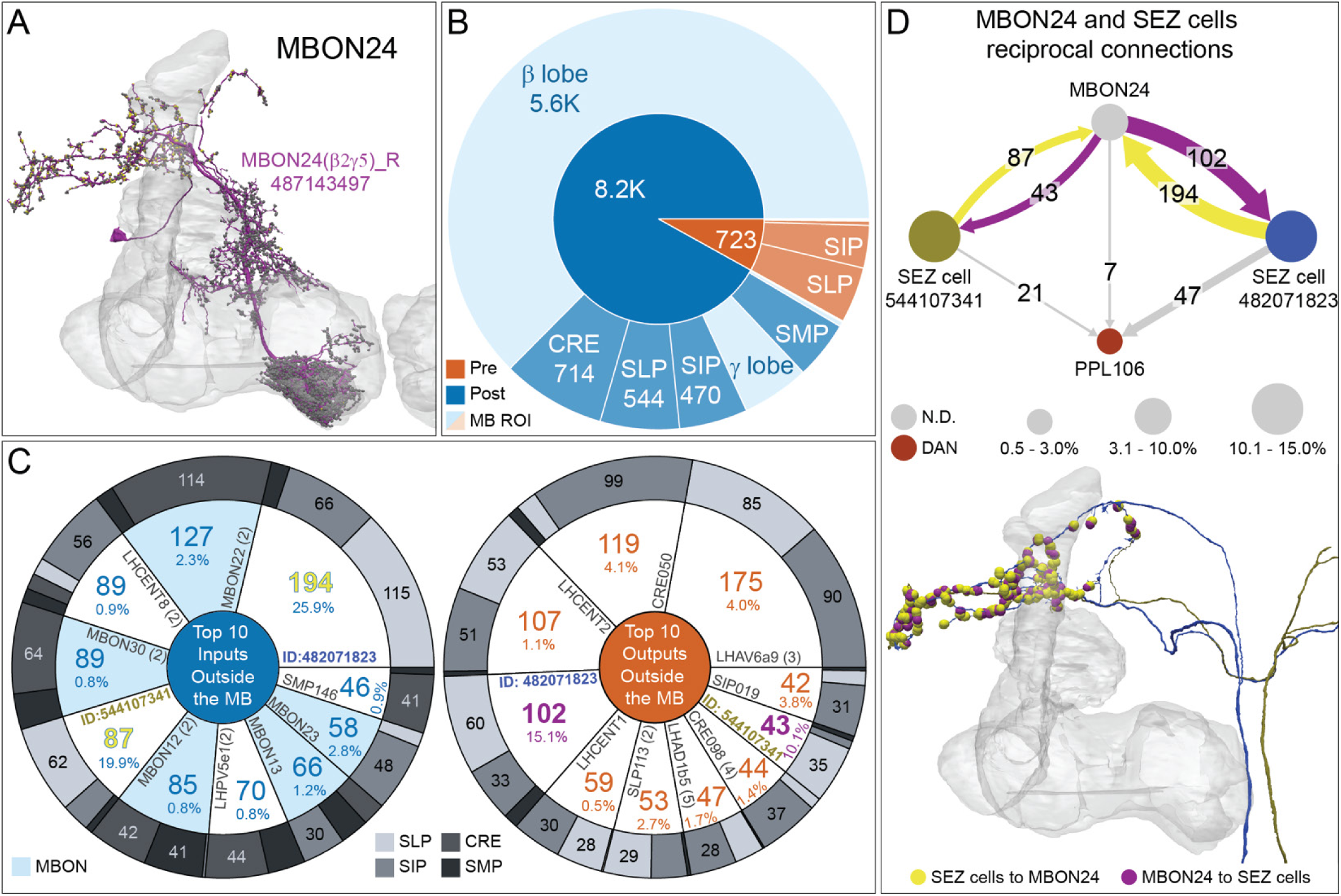
MBON24. (A) Atypical MBON24 (β2γ5) is shown here and in Video 9. Presynaptic sites are shown in yellow and postsynaptic sites are shown in grey. MBON24 receives 75 percent of its inputs from sites within the MB lobes and 90 percent of these inputs are in the β2 compartment. (B) A pie chart showing the distribution of MBON24’s presynaptic sites (indicated in orange) and postsynaptic sites (in blue). The inner circle shows the total number of MBON24’s pre- and postsynaptic sites. The outer circle of the pie chart shows the brain regions in which these sites occur; regions of the MB are shown in lighter colors. (C) MBON24’s top ten inputs by neuron type outside the MB lobes (left) are shown; links to these cell types in neuPrint are: <u>482071823</u>, <u>MBON22</u>, <u>LHCENT8</u>, <u>MBON30</u>, <u>544107341</u>, <u>MBON12</u>, <u>LHPV5e1</u>, <u>MBON13</u>, <u>MBON23</u>, and <u>SMP146</u>. The numbers in each slice represent synapse numbers, while the percentages indicate the percentage of that neuron’s synaptic output that goes to MBON24; the shading of the outer ring of the pie chart indicates where these synapses are located. Note that four of MBON24’s top input neurons are typical MBONs (indicated in light blue). MBON24’s top ten outputs by neuron type outside the MB lobes (right) are shown; links to these cell types in neuPrint are: <u>LHAV6a9</u>, <u>CRE050</u>, <u>LHCENT2</u>, <u>482071823</u>, <u>LHCENT1</u>, <u>SLP113</u>, <u>LHAD1b5</u>, <u>CRE098</u>, <u>544107341</u>, and <u>SIP019</u>. The numbers in each slice represent synapse numbers, while the percentages indicate the fraction of that neurons input that is provided by MBON24; the shading of the outer ring of the pie chart indicates where these synapses are located. Synapse numbers in highlighted text indicate neurons further discussed in panel D and are color-coded to match the diagrams in panel D. (D) Two of the top inputs to MBON24, and MBON24’s reciprocal connections back to them, are shown. These cells send processes outside of the hemibrain volume, presumably to the subesophageal zone (SEZ), as has been confirmed for the more strongly connected of the two (482071823) by reconstructing its counterpart in the FAFB volume (Figure 35 C and E). The same two neurons also provide strong input to a PPL106, contributing more than 0.5% of its total input, and are part of cluster 27 of DAN inputs shown in Figure 31. These SEZ cells also receive strong input from MBON14 (387 synapses).

**Figure 8-figure supplement 4.**
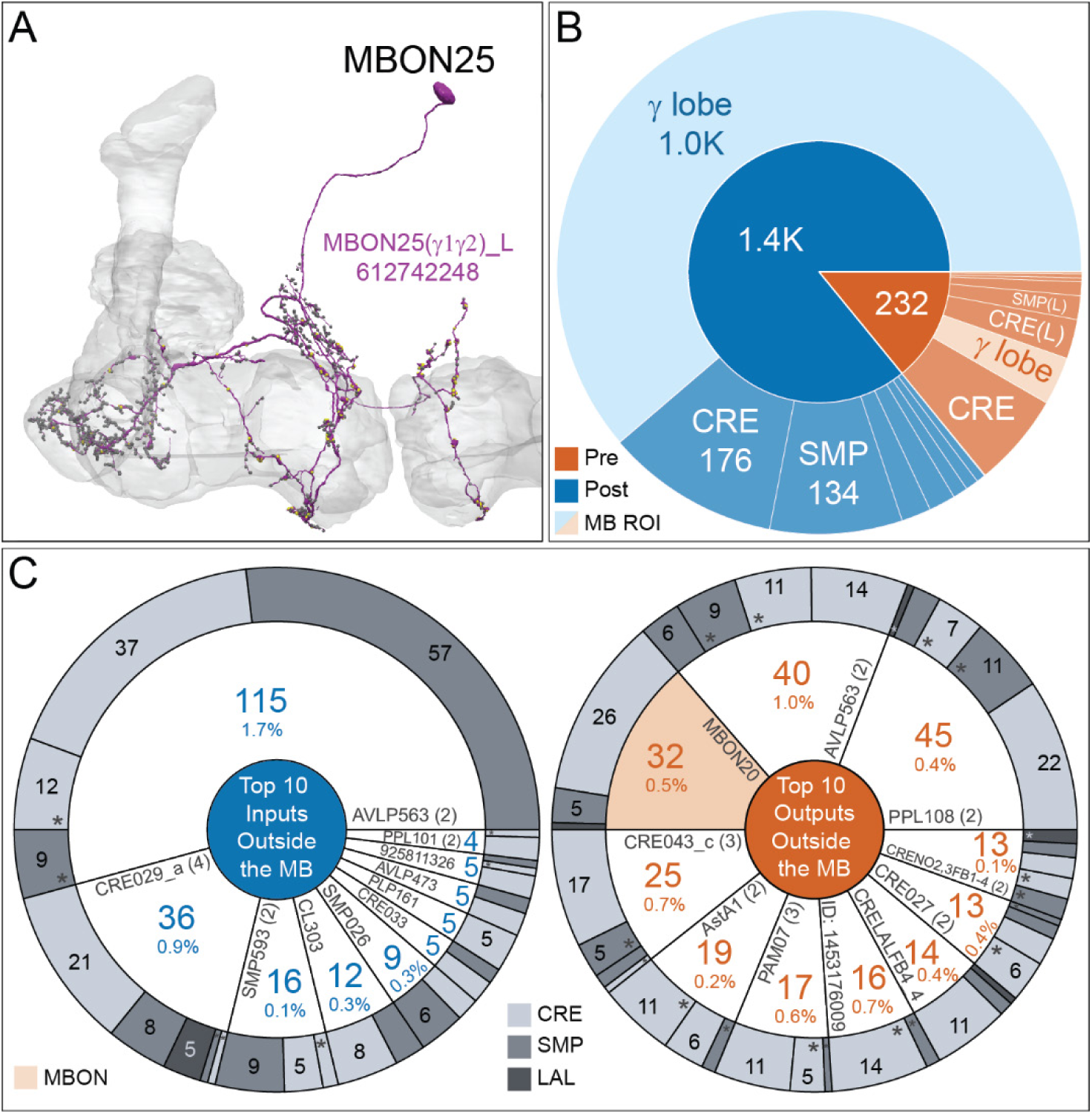
Atypical MBON25. (A) Atypical MBON25 (γ1γ2) is shown here and in Video 10. Presynaptic sites are shown in yellow and postsynaptic sites are shown in grey. MBON25 receives input in the γ1 and γ2 compartments. (B) A pie chart showing the distribution of MBON25’s presynaptic sites (indicated in orange) and postsynaptic sites (in blue). The inner circle shows the total number of MBON25’s pre- and postsynaptic sites. The outer circle of the pie chart shows the brain regions in which these sites occur; regions of the MB are shown in lighter colors. The CRE is MBON25’s major output area and MBON25 extends bilaterally to the left hemisphere CRE. (C) MBON25’s top ten inputs by neuron type outside the MB lobes (left) are shown; links to these cell types in neuPrint are: <u>AVLP563</u>, <u>CRE029_a</u>, <u>SMP593</u>, <u>CL303</u>, <u>SMP026</u>, <u>AVLP473</u>, <u>CRE033</u>, <u>PLP161</u>, <u>925811326</u>, <u>PPL102</u>, and <u>PPL101</u>. The numbers in each slice represent synapse numbers, while the percentages indicate the percentage of that neuron’s synaptic output that goes to MBON25; the shading of the outer ring of the pie chart indicates where these synapses are located. MBON25’s top ten outputs by neuron type outside the MB lobes (right) are shown; links to these cell types in neuPrint are: <u>PPL108</u>, <u>AVLP563</u>, <u>MBON20</u>, <u>CRE043_c</u>, <u>AstA1</u>, <u>PAM07</u>, <u>1453176009</u>, <u>CRELALFB4_4 (FB4H), CRE027, and CRENO2,3FB1-4 (FB1H). The numbers in each slice represent synapse numbers, while the percentages indicate the fraction of that neuron’s input that is provided by MBON25; the shading of the outer ring of the pie chart indicates where these synapses are located (asterisks indicate that these synapses are located in the left hemisphere). MBON25 makes axo-axonal connections outside the MB onto MBON20 (light orange shading); MBON20 and MBON25 both innervate the γ1 and γ2 compartments.</u>

**Figure 8-figure supplement 5.**
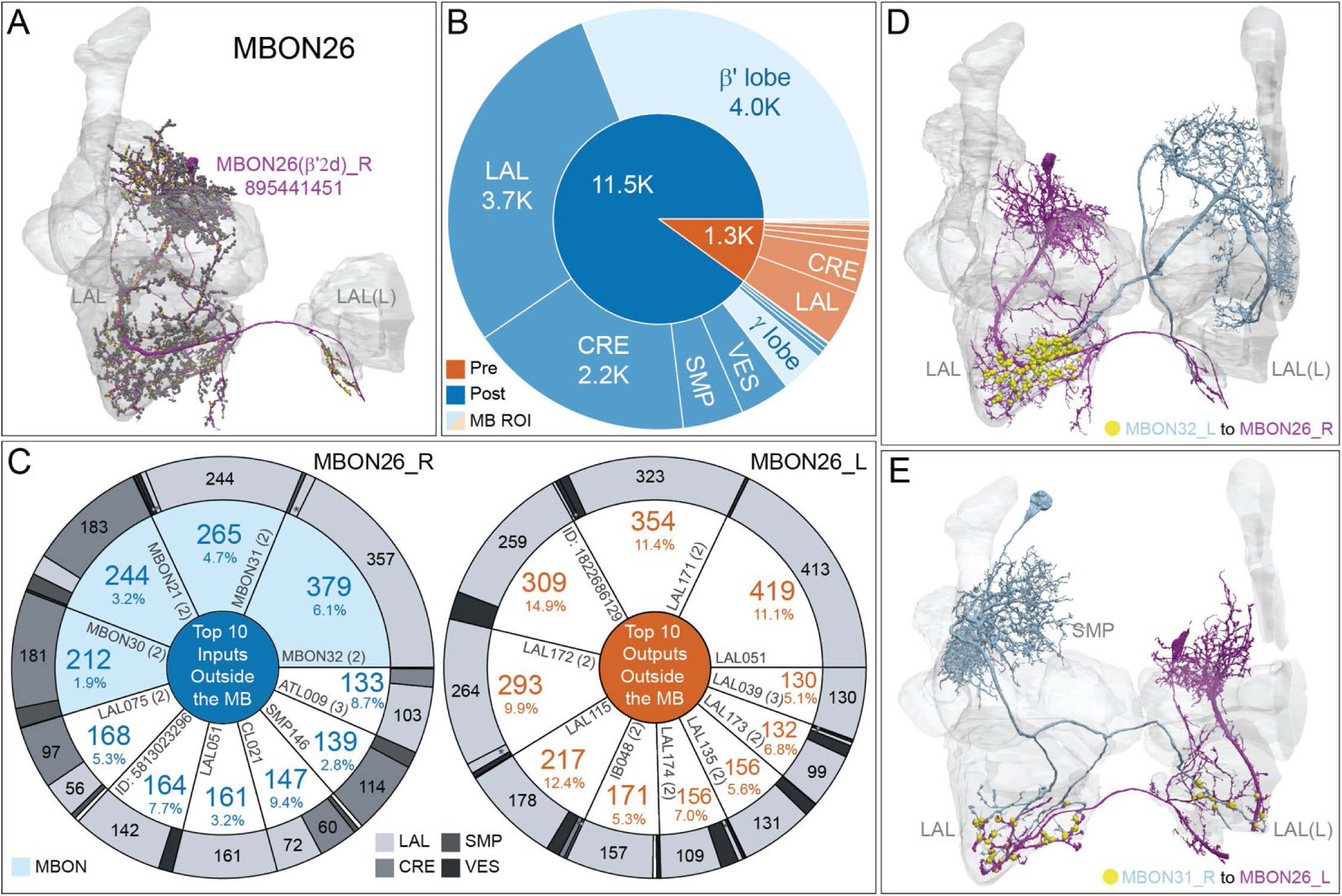
Atypical MBON26. (A) Atypical MBON26 (β*′*2d) is shown here and in Video 11. Presynaptic sites are shown in yellow and postsynaptic sites are shown in grey. MBON26 receives input from the β*′*2 compartment. (B) A pie chart showing the distribution of MBON26’s presynaptic sites (indicated in orange) and postsynaptic sites (in blue). The inner circle shows the total number of MBON26’s pre- and postsynaptic sites. The outer circle of the pie chart shows the brain regions in which these sites occur; regions of the MB are shown in lighter colors. Note MBON26_R’s output in LAL(L) is incomplete due to the left LAL(L) being only partially contained within the hemibrain volume (see panel D); we therefore display the output of MBON26_L in LAL(R) in the output pie chart to indicate what we believe to be MBON26’s full output in the LAL. (C) MBON26’s top ten inputs by neuron type outside the MB lobes (left) are shown; links to these cell types in neuPrint are: <u>MBON32</u>, <u>MBON31</u>, <u>MBON21</u>, <u>MBON30</u>, <u>LAL075</u>, <u>5813023296</u>, <u>LAL051</u>, <u>CL021</u>, <u>SMP146</u>, and <u>ATL009</u>. The numbers in each slice represent synapse numbers, while the percentages indicate the percentage of that neuron’s synaptic output that goes to MBON26; the shading of the outer ring of the pie chart indicates where these synapses are located. MBON26’s top ten outputs by neuron type outside the MB lobes (right) are shown; links to these cell types in neuPrint are: <u>LAL051</u>, <u>LAL171</u>, <u>1822686129</u>, <u>LAL172</u>, <u>LAL115</u>, <u>IB048</u>, <u>LAL174</u>, <u>LAL135</u>, <u>LAL173</u>, and <u>LAL039</u>. The numbers in each slice represent synapse numbers, while the percentages indicate the fraction of that neuron’s input that is provided by MBON26; the shading of the outer ring of the pie chart indicates where these synapses are located (asterisk indicates that these synapses are located in the left hemisphere). Note that MBON26’s strongest inputs come from MBON21 (γ4γ5) and three atypical MBONs, MBON30 (γ1γ2γ3), MBON31 (α*′*1) and MBON32 (γ2), (indicated in light blue). MBON26 is at the top layer of an MBON feedforward hierarchy network and also gets strong input from MBON27 (γ5d) and MBON29 (γ4γ5) (Figure 24; Figure 24-figure supplement 1). (D,E) The connections from MBON31 and MBON32 to MBON 26 occur almost exclusively in the LAL.

**Figure 8-figure supplement 6.**
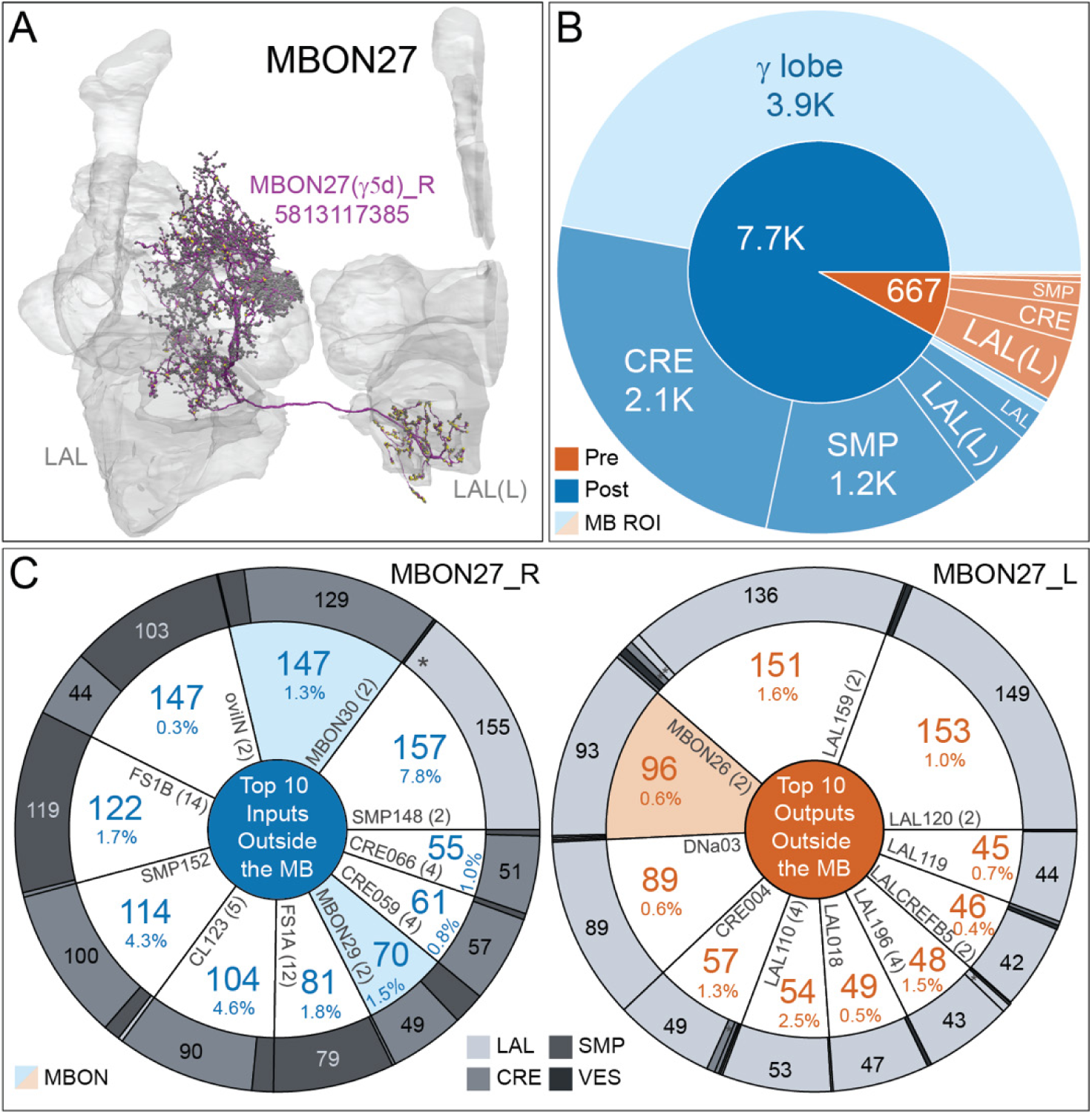
Atypical MBON27. (A) Atypical MBON27 (γ5d) is shown here and in Video 12. Presynaptic sites are shown in yellow and postsynaptic sites are shown in grey. MBON27 receives about half its input inside the MB, from the γ5 compartment, and outputs to both the ipsi- and contralateral LALs. (B) A pie chart showing the distribution of MBON27’s presynaptic sites (indicated in orange) and postsynaptic sites (in blue). The inner circle shows the total number of MBON27’s pre- and postsynaptic sites. The outer circle of the pie chart shows the brain regions in which these sites occur; regions of the MB are shown in lighter colors. The LAL is MBON27’s major output area. (C) MBON27’s top ten inputs by neuron type outside the MB lobes (left) are shown; links to these cell types in neuPrint are: <u>SMP148</u>, <u>MBON30</u>, <u>oviIN</u>, <u>FS1B</u>, <u>SMP152</u>, <u>CL123</u>, <u>FS1A</u>, <u>MBON29</u>, <u>CRE059</u>, and <u>CRE066</u>. The numbers in each slice represent synapse numbers, while the percentages indicate the percentage of that neuron’s synaptic output that goes to MBON27; the shading of the outer ring of the pie chart indicates where these synapses are located. MBON27’s top ten outputs by neuron type outside the MB lobes (right) are shown; links to these cell types in neuPrint are: <u>LAL120</u>, <u>LAL159</u>, <u>MBON26</u>, <u>DNa03</u>, <u>CRE004</u>, <u>LAL110</u>, <u>LAL018</u>, <u>LAL196</u>, <u>LALCREFB5 (FB5A), and LAL119. The numbers in each slice represent synapse numbers, while the percentages indicate the fraction of that neurons input that is provided by MBON27; the shading of the outer ring of the pie chart indicates where these synapses are located (asterisk indicates that these synapses are located in the left hemisphere). Note that MBON27 receives input from two atypical MBONs, MBON29 (γ4γ5) and MBON30 (γ1γ2γ3), and outputs onto another atypical MBON, MBON26 (β*′*2d; indicated by the light blue and light orange shading in input and output pie charts respectively).</u>

**Figure 8-figure supplement 7.**
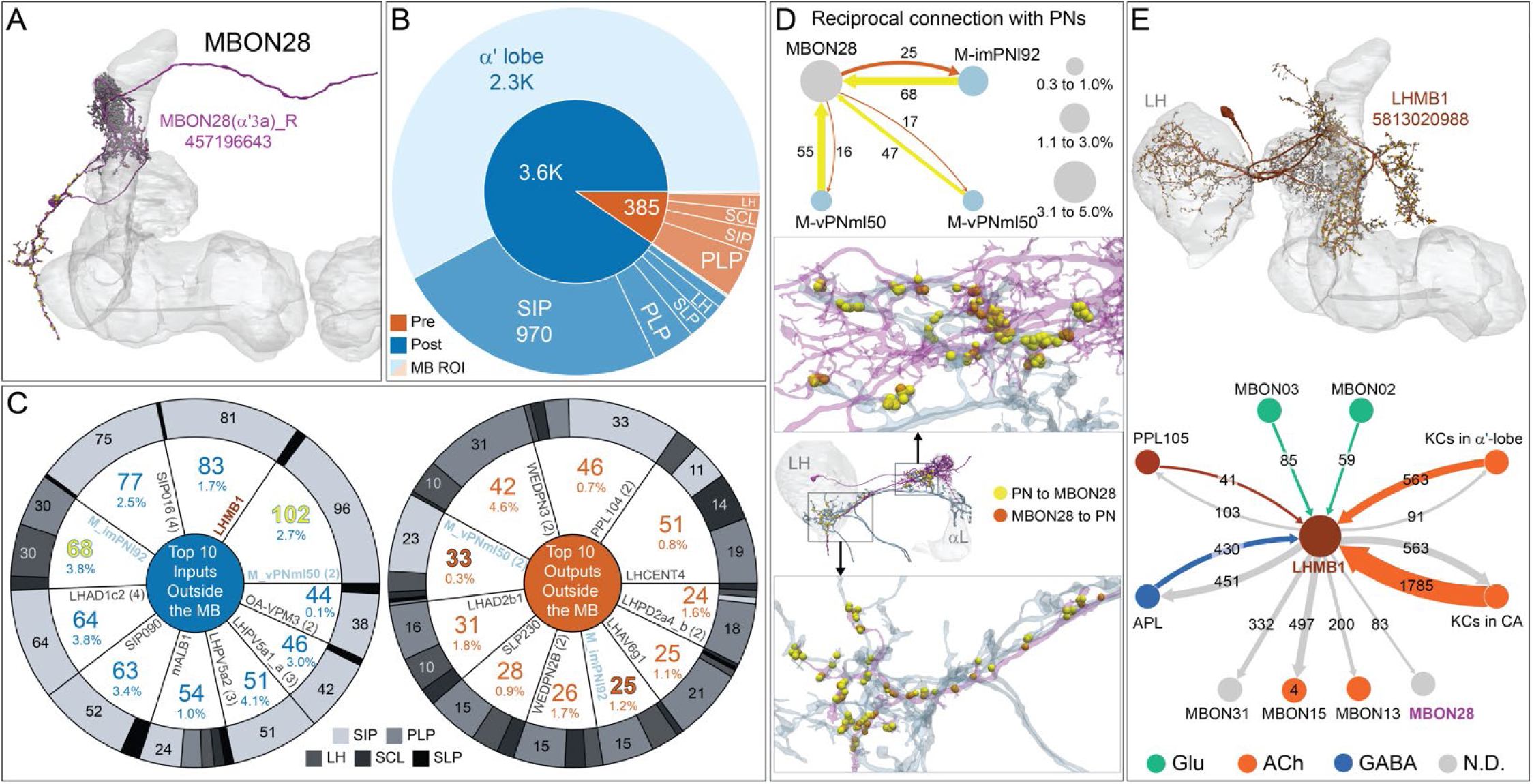
Atypical MBON28. (A) Atypical MBON28 (α′3) is shown here and in Video 13. Presynaptic sites are shown in yellow and postsynaptic sites are shown in grey. MBON 28 receives about 60 percent of its input from the α′3 compartment within the MB lobes; within the α′3 compartment its arbors have a similar morphology to those of MBON16, but MBON28’s dendrites also extend outside the MB lobe. (B) A pie chart showing the distribution of MBON28’s presynaptic sites (indicated in orange) and postsynaptic sites (in blue). The inner circle shows the total number of MBON28’s pre- and postsynaptic sites. The outer circle of the pie chart shows the brain regions in which these sites occur; regions of the MB are shown in lighter colors. The PLP is MBON28’s major output area but it also sends output to the LH. (C) MBON28’s top ten inputs by neuron type outside the MB lobes (left) are shown; links to these cell types in neuPrint are: <u>M_vPNml50</u>, <u>LHMB1</u>, <u>SIP016</u>, <u>M_imPNl92</u>, <u>LHAD1c2</u>, <u>SIP090</u>, <u>mALB1</u>, <u>LHPV5a2</u>, <u>LHPV5a1_a</u>, and <u>OA-VPM3</u>. The numbers in each slice represent synapse numbers, while the percentages indicate the percentage of that neuron’s synaptic output that goes to MBON28; the shading of the outer ring of the pie chart indicates where these synapses are located. MBON28’s top ten outputs by neuron type outside the MB lobes (right) are shown; links to these cell types in neuPrint are: <u>LHCENT4</u>, <u>PPL104</u>, <u>WEDPN3</u>, <u>M_vPNml50</u>, <u>LHAD2b1</u>, <u>SLP230</u>, <u>WEDPN2B</u>, <u>M_imPNl92</u>, <u>LHAV6g1</u>, and <u>LHPD2a4_b</u>. The numbers in each slice represent synapse numbers, while the percentages indicate the fraction of that neurons input that is provided by MBON28; the shading of the outer ring of the pie chart indicates where these synapses are located. Synapse numbers in highlighted text indicate neurons further discussed in panel D and are color-coded to match the diagrams in panel D. (D) MBON28 makes strong reciprocal connections with three inhibitory multi-glomerular PNs. The size of the circle representing the PN indicates the percentage of that PN’s output that goes to MBON28. The size of the MBON28 circle represents the percentage of its inputs outside the MB provided by the three PNs. This is the only MBON that receives PN input among its top ten inputs. These reciprocal connections are distributed in two major areas: one near the MB, the other near or inside the LH. (E) Left: MBON28’s second strongest input neuron, LHMB1, is shown here and in Video 13; LHMB1 innervates both the LH, the CA and the α lobe. Right: Connectivity diagram showing LHMB1’s main outputs and inputs. LHMB1 makes synapses onto MBONs on their dendrites inside the lobes as well as outside the MB through axo-axonal connections, and receives input from MBON02 (β2β′2a) and MBON03 (β′2mp), and reciprocally connects to APL and KCs in both the α lobe and CA.

**Figure 8-figure supplement 8.**
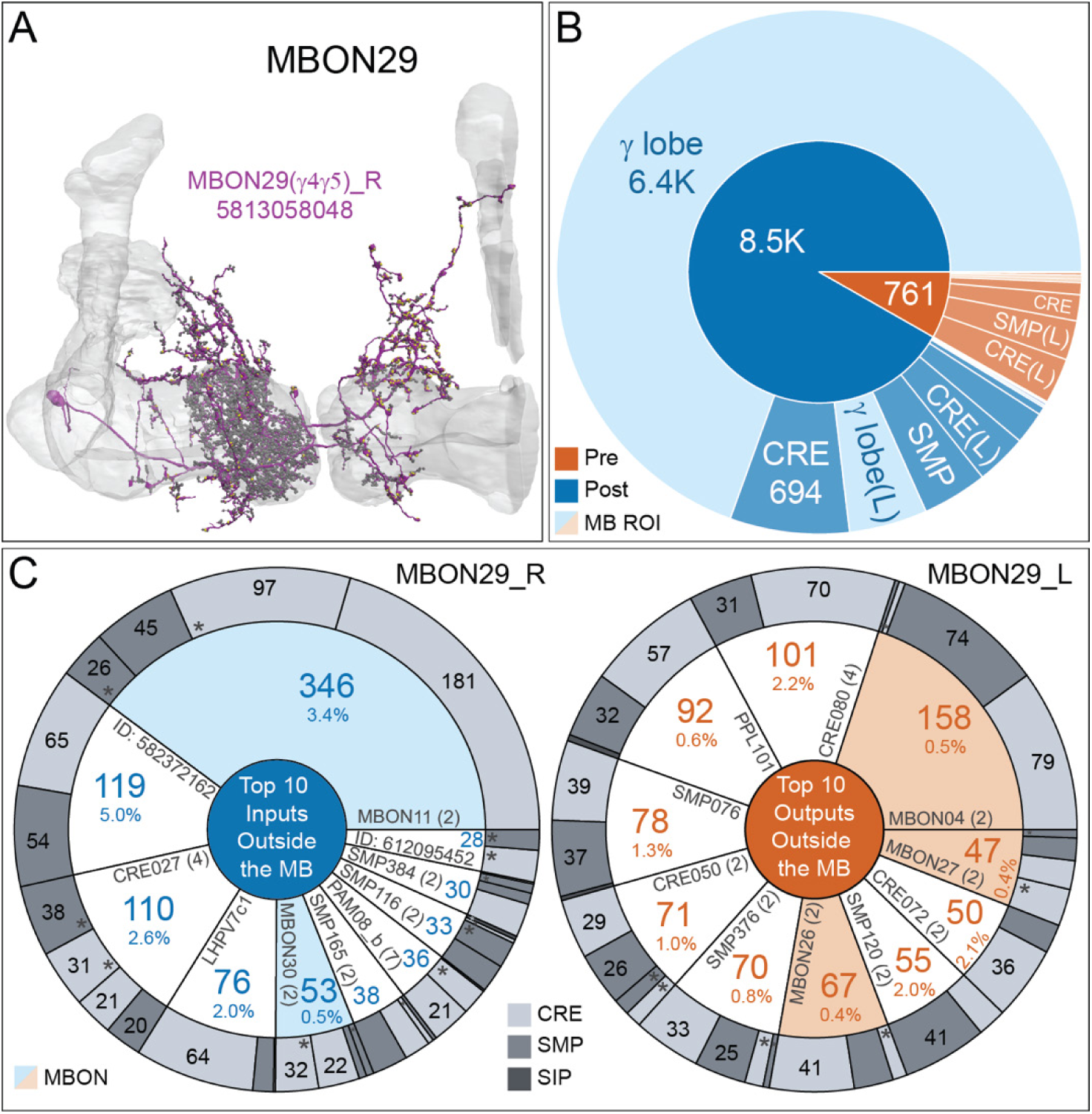
Atypical MBON29. (A) Atypical MBON29 (γ4γ5) is shown here and in Video 14. Presynaptic sites are shown in yellow and postsynaptic sites are shown in grey. MBON29 receives about 80% of its input from the γ4 and γ5 compartments within the MB lobes. (B) A pie chart showing the distribution of MBON29’s presynaptic sites (indicated in orange) and postsynaptic sites (in blue). The inner circle shows the total number of MBON29’s pre- and postsynaptic sites. The outer circle of the pie chart shows the brain regions in which these sites occur; regions of the MB are shown in lighter colors. (C) MBON29’s top ten inputs by neuron type outside the MB lobes (left) are shown; links to these cell types in neuPrint are: <u>MBON11</u>, <u>582372162</u>, <u>CRE027</u>, <u>LHPV7c1</u>, <u>MBON30</u>, <u>SMP165</u>, <u>PAM08_b</u>, <u>SMP116</u>, <u>SMP384</u>, and <u>612095452</u>. The numbers in each slice represent synapse numbers, while the percentages indicate the percentage of that neuron’s synaptic output that goes to MBON29; the shading of the outer ring of the pie chart indicates where these synapses are located. MBON29’s top ten outputs by neuron type outside the MB lobes (right) are shown; links to these cell types in neuPrint are: <u>MBON04</u>, <u>CRE080</u>, <u>PPL101</u>, <u>SMP076</u>, <u>SMP376</u>, <u>CRE050</u>, <u>MBON26</u>, <u>SMP120</u>, <u>CRE072</u>, and <u>MBON27</u>. (Note that to ensure completeness of the contralateral arbor, MBON29_L was used.) The numbers in each slice represent synapse numbers, while the percentages indicate the fraction of that neuron’s input that is provided by MBON29; the shading of the outer ring of the pie chart indicates where these synapses are located (asterisks indicate that these synapses are located in the left hemisphere). Note that MBON29 receives very strong input from MBON11 (γ1pedc>α/β)_right and _left, which together contribute 9% of MBON29’s total input outside MB and also receives input from MBON30 (γ1γ2γ3; indicated by the light blue shading in the input pie chart). MBON29 makes axo-axonal synapses onto MBON04 (β′2mp_bilateral) as well as onto the atypical MBONs, MBON26 (β′2d) and MBON27 (γ5d; indicated by the light orange shading in the output pie chart).

**Figure 8-figure supplement 9.**
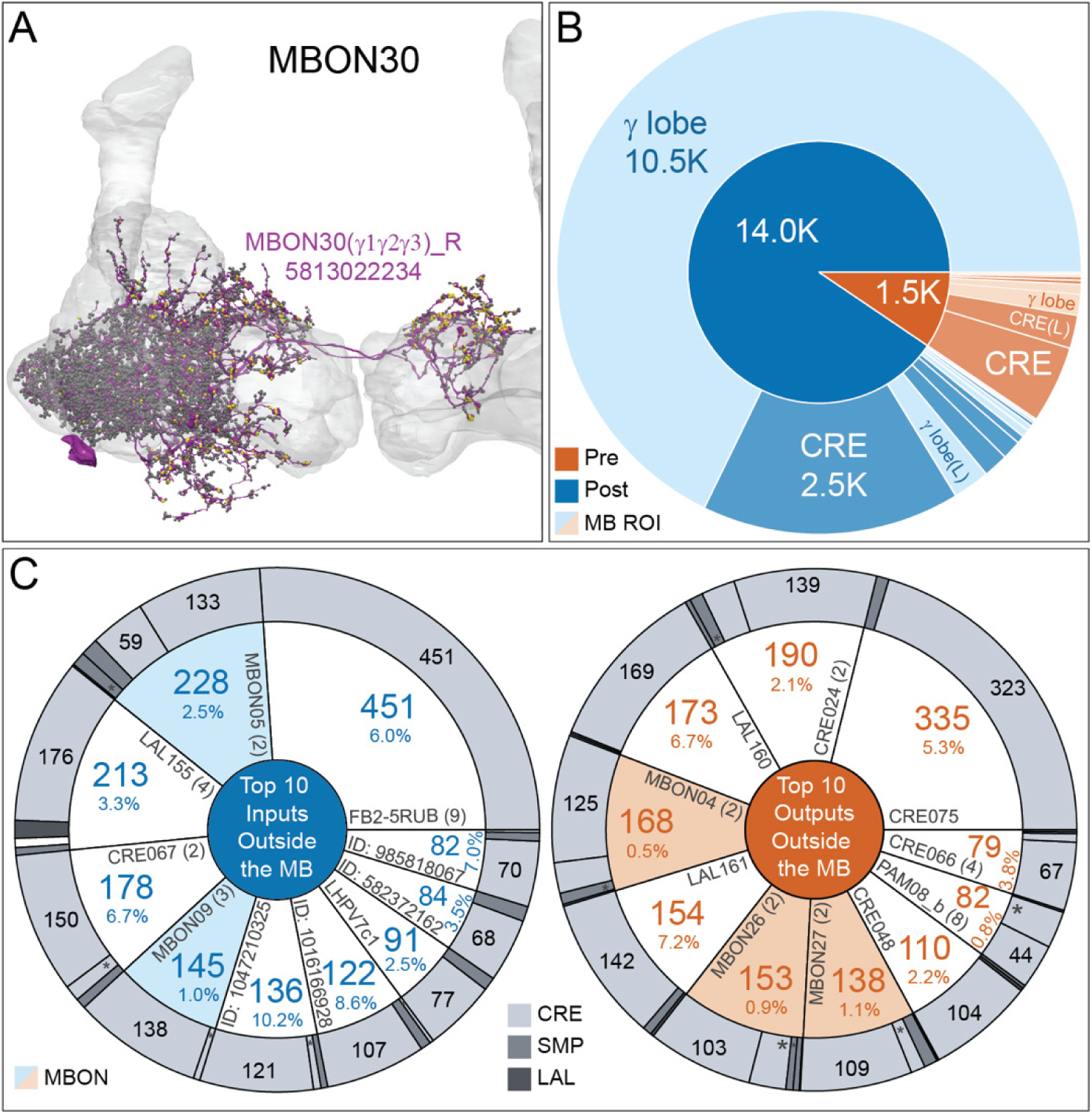
Atypical MBON30. (A) Atypical MBON30 (γ1γ2γ3) is shown here and in Video 15. Presynaptic sites are shown in yellow and postsynaptic sites are shown in grey. MBON30 receives about 80% of its input within the MB lobes, primarily from the γ1, γ2, and γ3 compartments. (B) A pie chart showing the distribution of MBON30’s presynaptic sites (indicated in orange) and postsynaptic sites (in blue). The inner circle shows the total number of MBON30’s pre- and postsynaptic sites. The outer circle of the pie chart shows the brain regions in which these sites occur; regions of the MB are shown in lighter colors. The CRE is MBON30’s major output area. (C) MBON30’s top ten inputs by neuron type outside the MB lobes (left) are shown; links to these cell types in neuPrint are: <u>FR1</u>, <u>MBON05</u>, <u>LAL155</u>, <u>CRE067</u>, <u>MBON09</u>, <u>1047210325</u>, <u>1016166928</u>, <u>LHPV7c1</u>, <u>582372162</u>, and <u>985818067</u>. The numbers in each slice represent synapse numbers, while the percentages indicate the percentage of that neuron’s synaptic output that goes to MBON30; the shading of the outer ring of the pie chart indicates where these synapses are located. MBON30 gets strong input from nine fan-shaped body cells of cell type <u>FB2-5RUB</u> (FR1; collectively they make 451 synapses onto MBON30’s arbors in a small brain area called the rubus (see Video 15). MBON30’s top ten outputs by neuron type outside the MB lobes (right) are shown; links to these cell types in neuPrint are: <u>CRE075</u>, <u>CRE024</u>, <u>LAL160</u>, <u>MBON04</u>, <u>LAL161</u>, <u>MBON26</u>, <u>MBON27</u>, <u>CRE048</u>, <u>PAM08_b</u>, and <u>CRE066</u>. The numbers in each slice represent synapse numbers, while the percentages indicate the fraction of that neuron’s input that is provided by MBON30; the shading of the outer ring of the pie chart indicates where these synapses are located (asterisks indicate that these synapses are located in the left hemisphere). Note that MBON30 receives input from MBON05 (γ4>γ1γ2) and MBON09 (γ3β′1) and makes synapses onto the atypical MBONs MBON27 (γ5d) and MBON28 (α′3) as well as to typical MBON04 (β′2mp_bilateral; indicated by the light blue and light orange shading in input and output pie charts, respectively).

**Figure 8-figure supplement 10.**
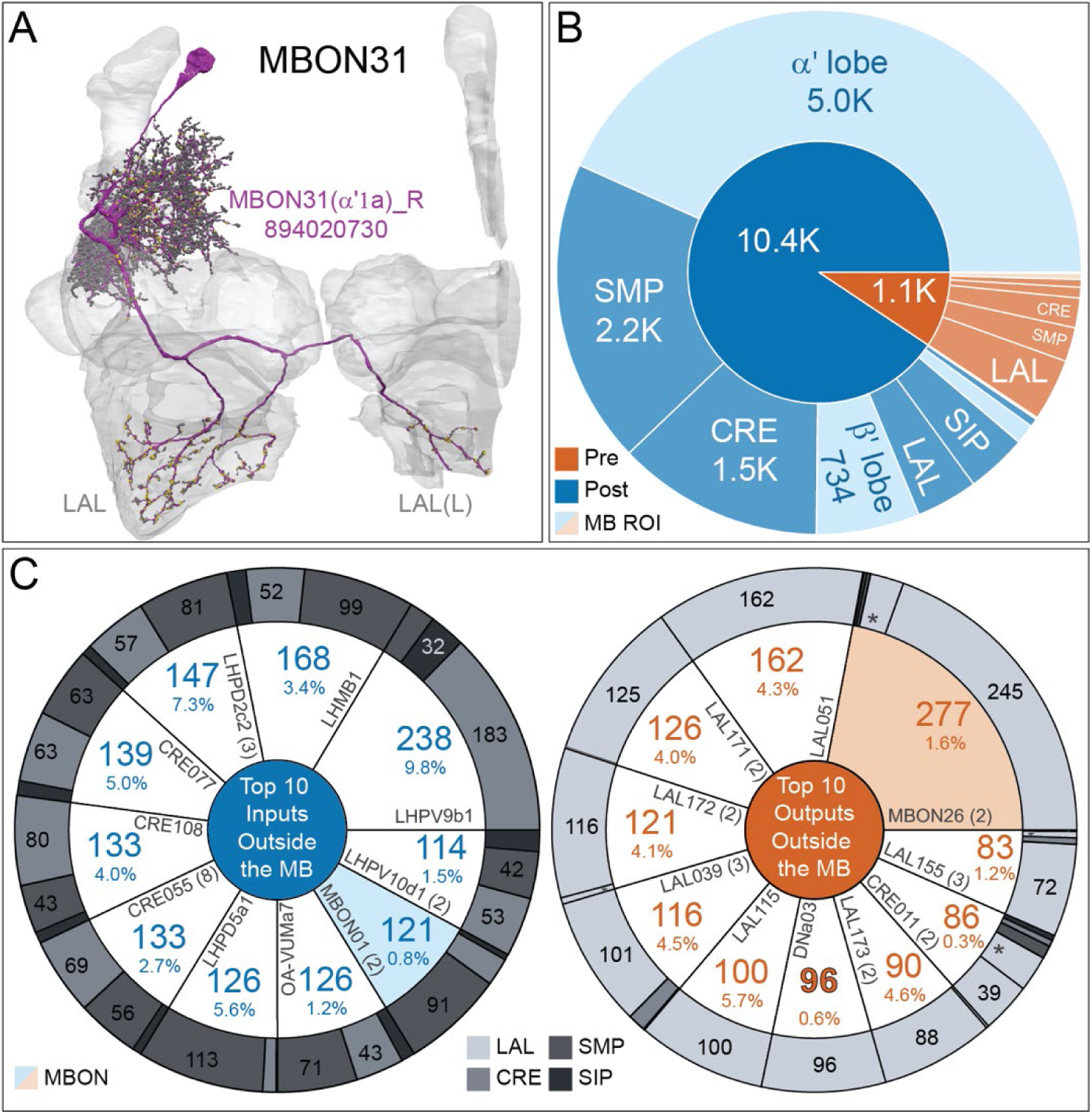
MBON31. (A) Atypical MBON31 (α′1) is shown here and in Video 16. Presynaptic sites are shown in yellow and postsynaptic sites are shown in grey. MBON31 gets about half of its inputs from the α′1 compartment inside the MB and sends outputs to LAL. (B) A pie chart showing the distribution of MBON31’s presynaptic sites (indicated in orange) and postsynaptic sites (in blue). The inner circle shows the total number of MBON31’s pre- and postsynaptic sites. The outer circle of the pie chart shows the brain regions in which these sites occur; regions of the MB are shown in lighter colors. (C) MBON31’s top ten inputs by neuron type outside the MB lobes (left) are shown; links to these cell types in neuPrint are: <u>LHPV9b1</u>, <u>LHMB1</u>, <u>LHPD2c2</u>, <u>CRE077</u>, <u>CRE108</u>, <u>CRE055</u>, <u>LHPD5a1</u>, <u>OA-VUMa7</u>, <u>MBON01</u>, and <u>LHPV10d1</u>. The numbers in each slice represent synapse numbers, while the percentages indicate the percentage of that neuron’s synaptic output that goes to MBON31; the shading of the outer ring of the pie chart indicates where these synapses are located. MBON31’s top ten outputs by neuron type outside the MB lobes (right) are shown; links to these cell types in neuPrint are: <u>MBON26</u>, <u>LAL051</u>, <u>LAL171</u>, <u>LAL172</u>, <u>LAL039</u>, <u>LAL115</u>, <u>DNa03</u>, <u>LAL173</u>, <u>CRE011</u>, and <u>LAL155</u>. The numbers in each slice represent synapse numbers, while the percentages indicate the fraction of that neuron’s input that is provided by MBON31; the shading of the outer ring of the pie chart indicates where these synapses are located (asterisk indicates that these synapses are located in the left hemisphere). MBON31 top 10 inputs include five cell types that convey information from the LH and MBON01 (γ5β’2a); its strongest downstream target is the atypical MBON, MBON26 (β′2d).

**Figure 8-figure supplement 11.**
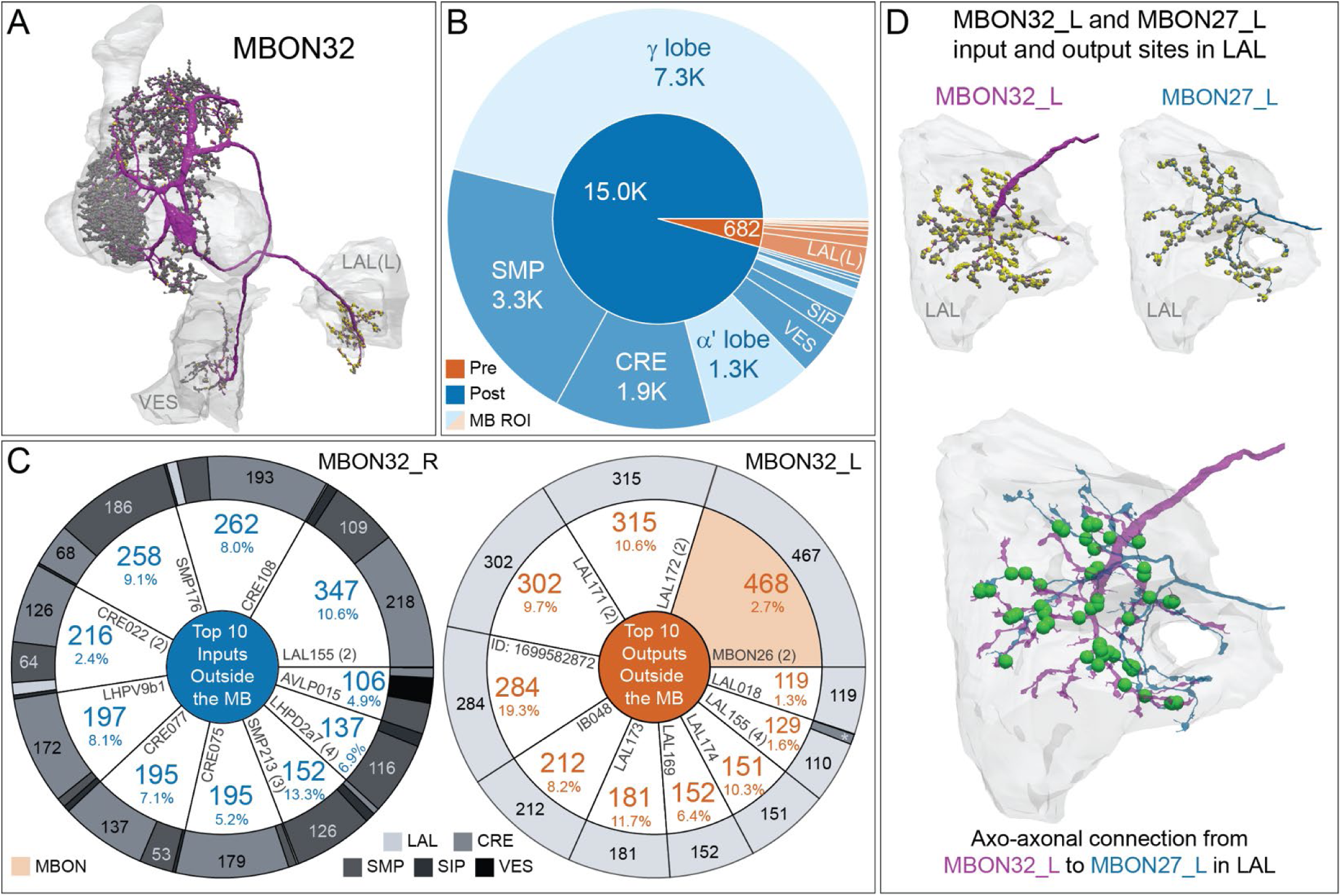
Atypical MBON32. (A) Atypical MBON32 (γ2) is shown here and in Video 17. Presynaptic sites are shown in yellow and postsynaptic sites are shown in grey. MBON32 gets about 50 percent of its synaptic input inside the MB lobes, primarily from the γ2 compartment, and sends about half its output to the contralateral LAL. (B) A pie chart showing the distribution of MBON32’s presynaptic sites (indicated in orange) and postsynaptic sites (in blue). The inner circle shows the total number of MBON32’s pre- and postsynaptic sites. The outer circle of the pie chart shows the brain regions in which these sites occur; regions of the MB are shown in lighter colors. (C) MBON32’s top ten inputs by neuron type outside the MB lobes (left) are shown; links to these cell types in neuPrint are: <u>LAL155</u>, <u>CRE108</u>, <u>SMP176</u>, <u>CRE022</u>, <u>LHPV9b1</u>, <u>CRE077</u>, <u>CRE075</u>, <u>SMP213</u>, <u>LHPD2a7,</u> and <u>AVLP015</u>. The numbers in each slice represent synapse numbers, while the percentages indicate the percentage of that neuron’s synaptic output that goes to MBON32; the shading of the outer ring of the pie chart indicates where these synapses are located. MBON32’s axon targets the contralateral LAL and the left brain hemisphere is incomplete in the hemibrain volume; therefore, we used MBON32_L’s arbors in the right hemisphere to ascertain MBON32’s downstream targets. The top ten outputs by neuron type outside the MB lobes (right) are shown; links to these cell types in neuPrint are: <u>MBON26</u>, <u>LAL172</u>, <u>LAL171</u>, <u>1699582872</u>, <u>IB048</u>, <u>LAL173</u>, <u>LAL169</u>, <u>LAL174</u>, <u>LAL155</u>, and <u>LAL018</u>. The numbers in each slice represent synapse numbers, while the percentages indicate the fraction of that neuron’s input that is provided by MBON32; the shading of the outer ring of the pie chart indicates where these synapses are located (asterisks indicate that these synapses are located in the left hemisphere). (D) We use the MBON32_L to illustrate MBON32’s downstream targets in the right hemisphere LAL. MBON27 (γ5d)’s axon also targets contralateral LAL (see Figure 8-figure supplement 6); the arbors of MBON32 and MBON27 in the LAL are shown separately in the upper portion of panel (D) with their pre- (yellow) and postsynaptic (grey) sites indicated. MBON32_L makes axo-axonal synapses onto MBON27_L, indicated by the green dots (lower portion of panel (D)).

**Figure 8-figure supplement 12.**
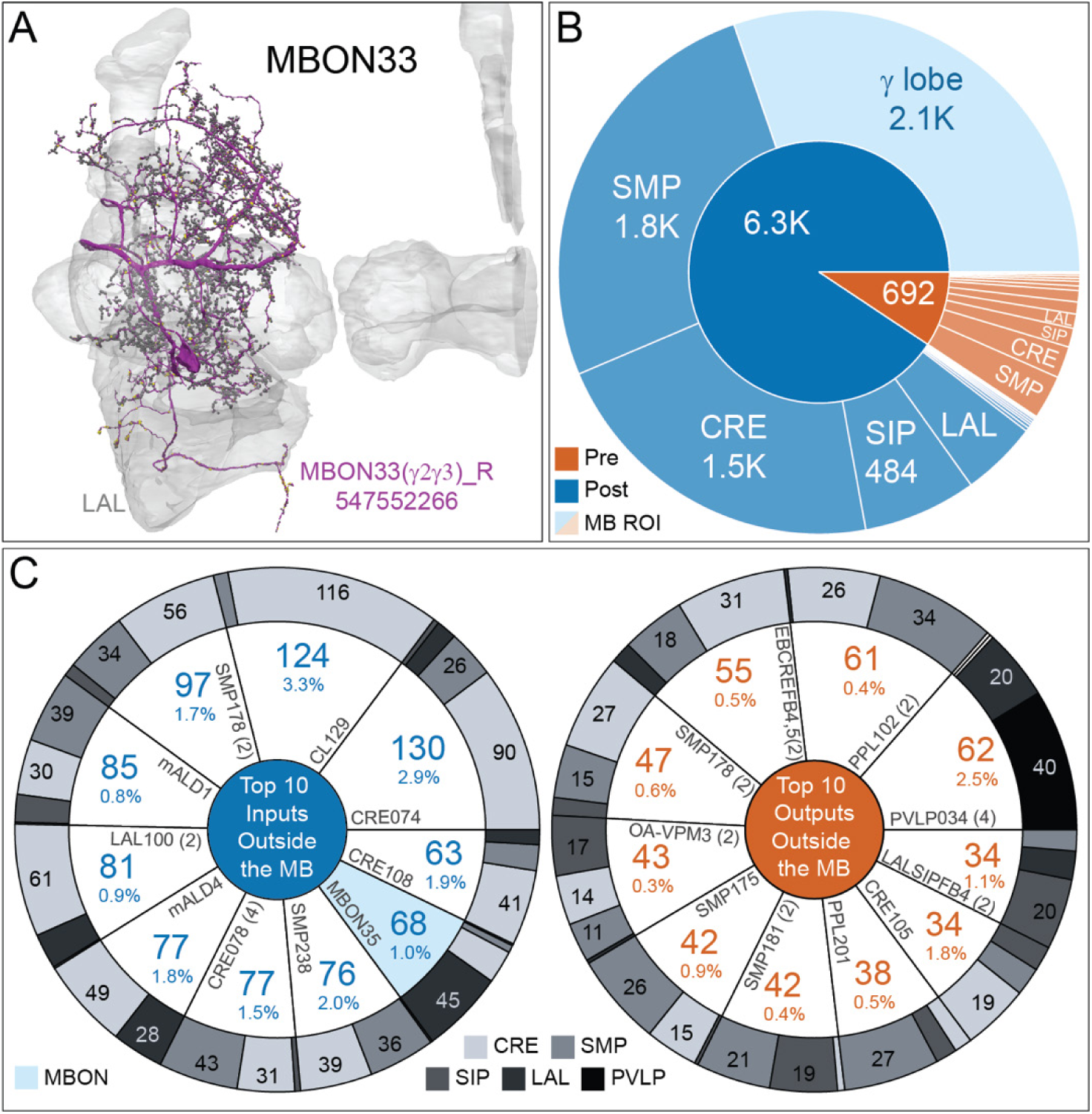
Atypical MBON33. (A) Atypical MBON33 (γ2γ3) is shown here and in Video 18. Presynaptic sites are shown in yellow and postsynaptic sites are shown in grey. MBON33 gets about one-third of its input synapses within the γ lobe of the MB, mostly in the γ2 and γ3 compartments, and sends about 9% of its output to the LAL. (B) A pie chart showing the distribution of MBON33’s presynaptic sites (indicated in orange) and postsynaptic sites (in blue). The inner circle shows the total number of MBON33’s pre- and postsynaptic sites. The outer circle of the pie chart shows the brain regions in which these sites occur; regions of the MB are shown in lighter colors. (C) MBON33’s top ten inputs by neuron type outside the MB lobes (left) are shown; links to these cell types in neuPrint are: <u>CRE074</u>, <u>CL129</u>, <u>SMP178</u>, <u>mALD1</u>, <u>LAL100</u>, <u>mALD4</u>, <u>CRE078</u>, <u>SMP238</u>, <u>MBON35</u>, and <u>CRE108</u>. The numbers in each slice represent synapse numbers, while the percentages indicate the percentage of that neuron’s synaptic output that goes to MBON33; the shading of the outer ring of the pie chart indicates where these synapses are located. MBON33’s top ten outputs by neuron type outside the MB lobes (right) are shown; links to these cell types in neuPrint are: <u>PVLP034</u>, <u>PPL102</u>, <u>EBCREFB4,5 (FB4Y)</u>, <u>SMP178</u>, <u>OA-VPM3</u>, <u>SMP175</u>, <u>SMP181</u>, <u>PPL201</u>, <u>CRE105</u>, and <u>LALSIPFB4 (FB4L)</u>. The numbers in each slice represent synapse numbers, while the percentages indicate the fraction of that neuron’s input that is provided by MBON33; the shading of the outer ring of the pie chart indicates where these synapses are located.

**Figure 8-figure supplement 13.**
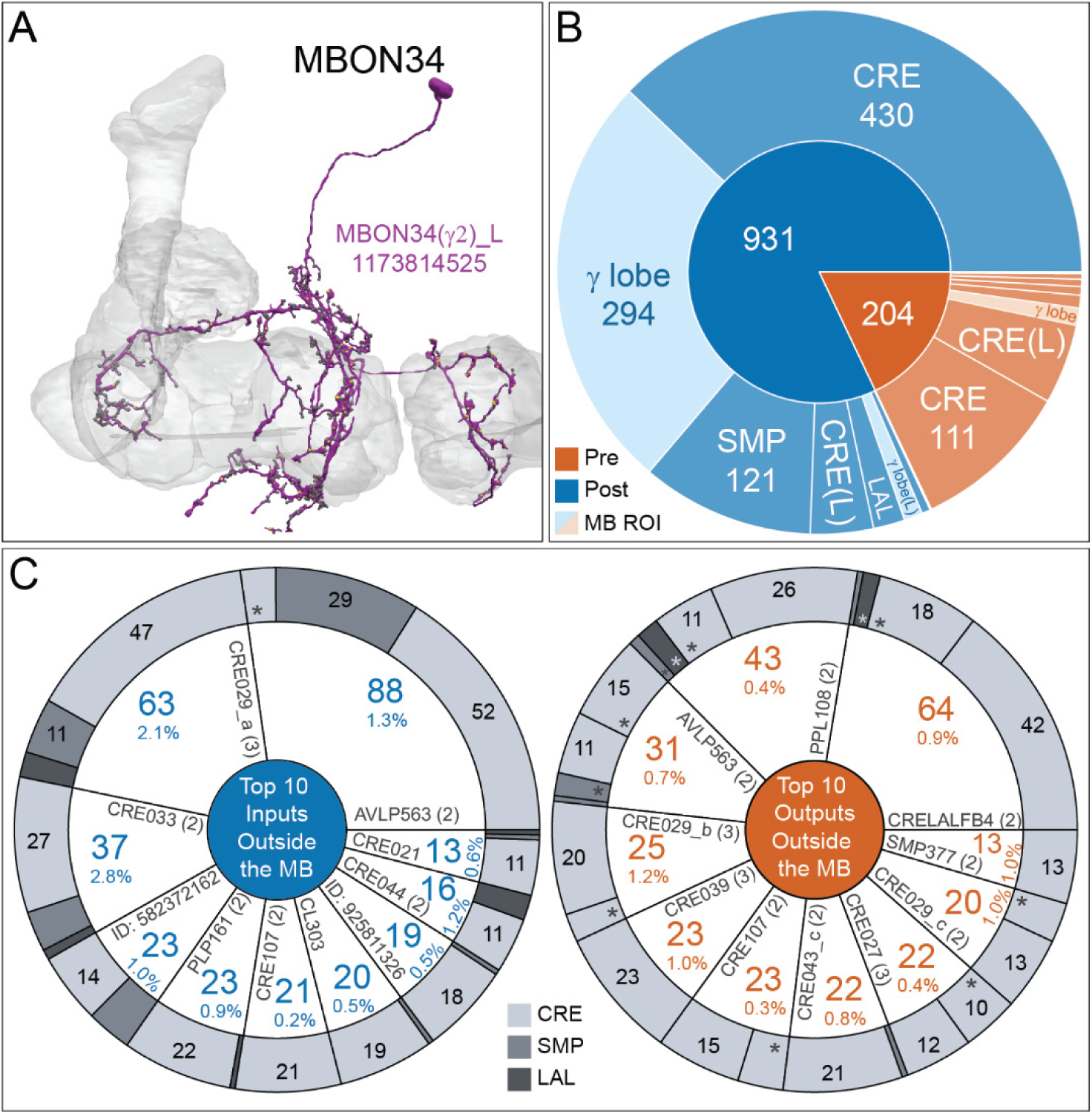
MBON34. (A) Atypical MBON34 (γ2) is shown here and in Video 19. Presynaptic sites are shown in yellow and postsynaptic sites are shown in grey. MBON34 has a small arbor and gets about one-third of its synapses from the γ2 compartment inside the MB lobes. (B) A pie chart showing the distribution of MBON34’s presynaptic sites (indicated in orange) and postsynaptic sites (in blue). The inner circle shows the total number of MBON34’s pre- and postsynaptic sites. The outer circle of the pie chart shows the brain regions in which these sites occur; regions of the MB are shown in lighter colors. (C) MBON34’s top ten inputs by neuron type outside the MB lobes (left) are shown; links to these cell types in neuPrint are: <u>AVLP563</u>, <u>CRE029_a</u>, <u>CRE033</u>, <u>582372162</u>, <u>PLP161</u>, <u>CRE107</u>, <u>CL303</u>, <u>925811326</u>, <u>CRE044</u>, and <u>CRE021</u>. The numbers in each slice represent synapse numbers, while the percentages indicate the percentage of that neuron’s synaptic output that goes to MBON34; the shading of the outer ring of the pie chart indicates where these synapses are located. MBON34’s top ten by neuron type outputs outside the MB lobes (right) are shown; links to these cell types in neuPrint are: <u>CRELALFB4_4</u> (FB4H), <u>PPL108</u>, <u>AVLP563</u>, <u>CRE029_b</u>, <u>CRE039</u>, <u>CRE107</u>, <u>CRE043_c</u>, <u>CRE027</u>, <u>CRE029_c</u>, and <u>SMP377</u>. The numbers in each slice represent synapse numbers, while the percentages indicate the fraction of that neuron’s input that is provided by MBON34; the shading of the outer ring of the pie chart indicates where these synapses are located (asterisks indicate that these synapses are located in the left hemisphere).

**Figure 8-figure supplement 14.**
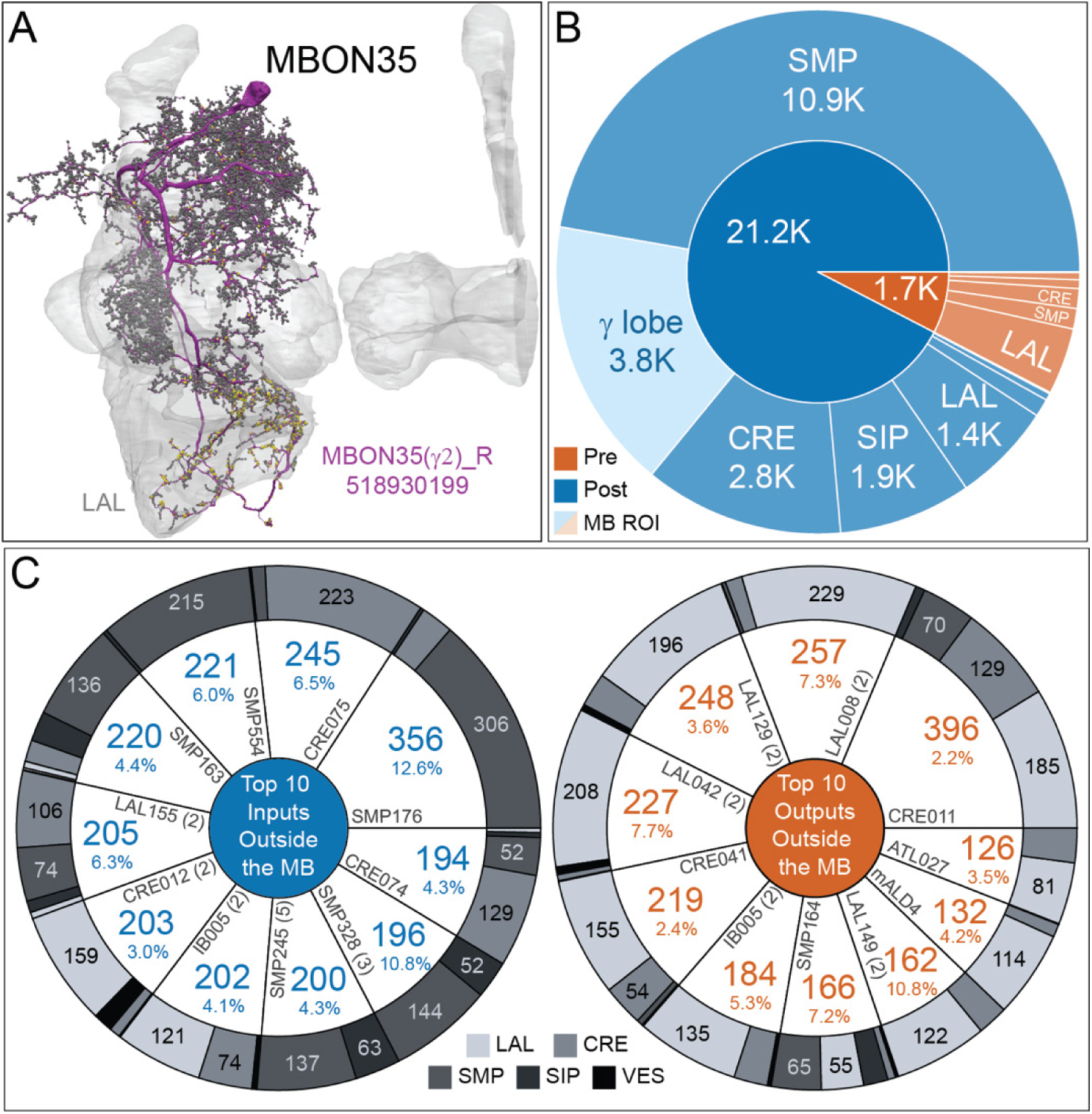
Atypical MBON35. (A) Atypical MBON35 (γ2) is shown here and in Video 20. Presynaptic sites are shown in yellow and postsynaptic sites are shown in grey. MBON35 gets about 20% of its input from the γ2 compartment inside the MB lobes and sends about half its output to the LAL. (B) A pie chart showing the distribution of MBON35’s presynaptic sites (indicated in orange) and postsynaptic sites (in blue). The inner circle shows the total number of MBON35’s pre- and postsynaptic sites. The outer circle of the pie chart shows the brain regions in which these sites occur; regions of the MB are shown in lighter colors. (C) MBON35’s top ten inputs by neuron type outside the MB lobes (left) are shown; links to these cell types in neuPrint are: <u>SMP176</u>, <u>CRE075</u>, <u>SMP554</u>, <u>SMP163</u>, <u>LAL155</u>, <u>CRE012</u>, <u>IB005</u>, <u>SMP245</u>, <u>SMP328</u>, and <u>CRE074</u>. The numbers in each slice represent synapse numbers, while the percentages indicate the percentage of that neuron’s synaptic output that goes to MBON35; the shading of the outer ring of the pie chart indicates where these synapses are located. MBON35’s top ten outputs by neuron type outside the MB lobes (right) are shown; links to these cell types in neuPrint are: <u>CRE011</u>, <u>LAL008</u>, <u>LAL129</u>, <u>LAL042</u>, <u>CRE041</u>, <u>IB005</u>, <u>SMP164</u>, <u>LAL149</u>, <u>mALD4</u>, and <u>ATL027</u>. The numbers in each slice represent synapse numbers, while the percentages indicate the fraction of that neuron’s input that is provided by MBON35; the shading of the outer ring of the pie chart indicates where these synapses are located.

**Figure 8-figure supplement 15.**
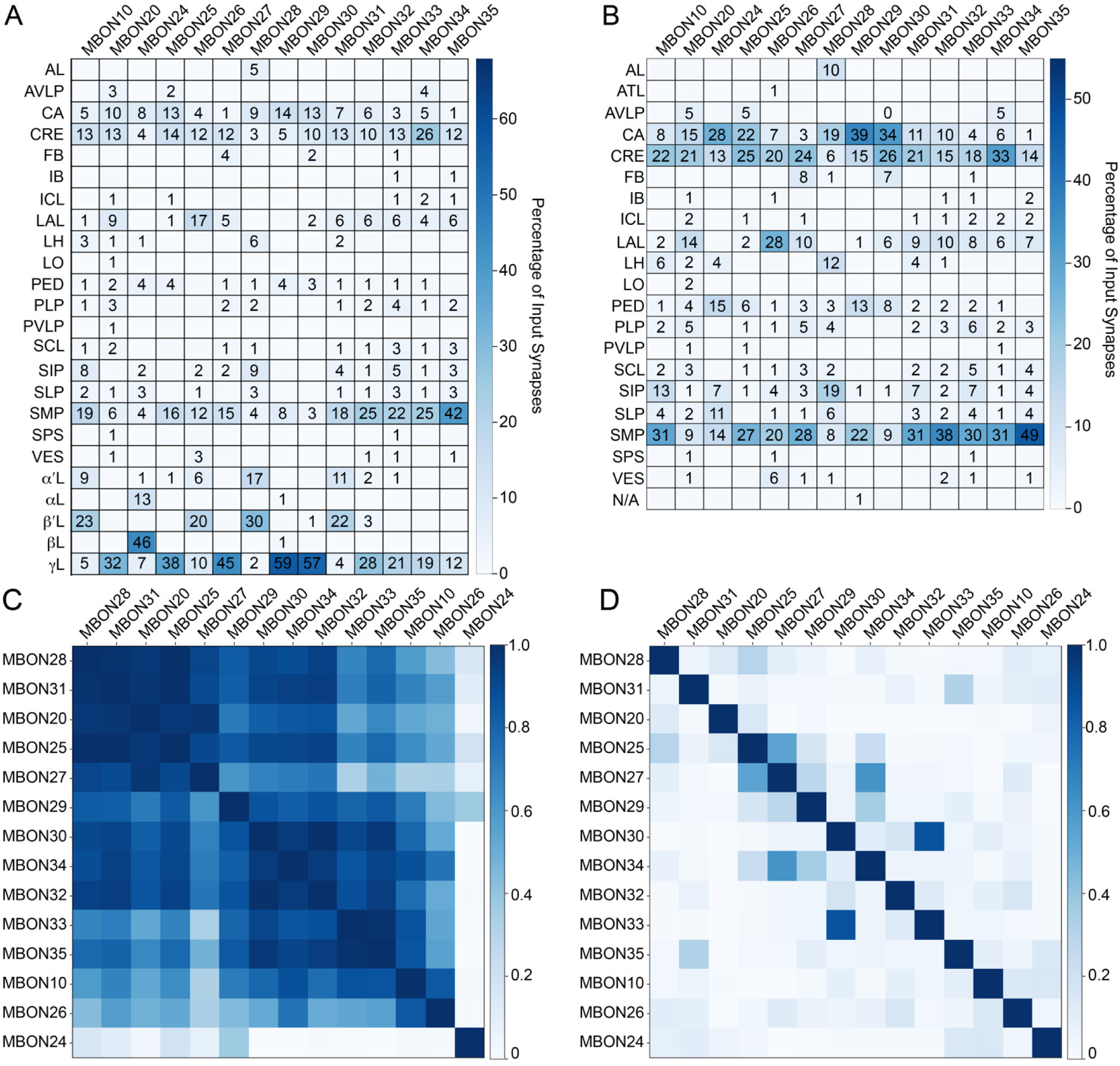
Atypical MBON input distribution by brain region and similarity of inputs to different MBONs. (A-B) Distribution of brain regions providing input to atypical MBON types. The value in each box indicates the effective input to the indicated MBON from the indicated brain region; that is, based on where the neurons making synapses onto the MBONs get their own inputs. Effective input is a measure that takes into account both the strength of the connection of each of the neurons that provides input to the MBON and the connection strength of other neurons to each of those input neurons in each brain region. Effective input is computed by matrix-multiplying the inputs to the MBONs and the inputs to those MBON-presynaptic neurons (normalizing both matrices so that inputs to all neurons sum to 1). Blank boxes indicate values of less than 1%. Input received inside MB lobes is included in A and excluded in B. (C) Cosine similarity between MBON types based on their input at the level of brain regions, with the MB excluded. (D) Cosine similarity between each MBON based on their inputs, with the MB excluded and the computation of the degree of input overlap performed at the level of individual presynaptic neurons. The ordering of MBONs in C and D was determined using spectral clustering.

**Figure 9-figure supplement 1.**
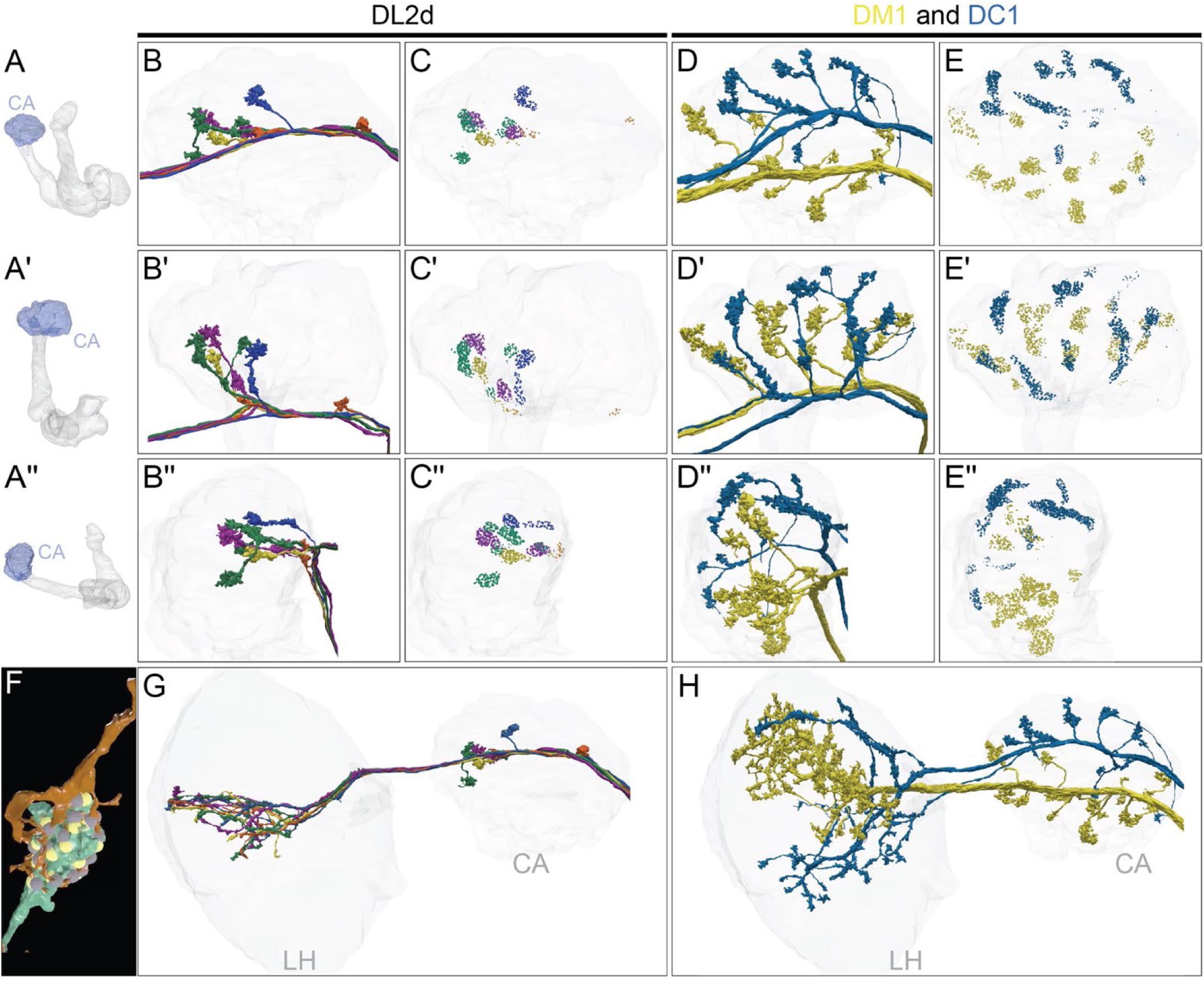
Distribution of the termini of olfactory PNs in the CA. (A-A′′) MB images showing the orientation of the CA (blue) in panels B-E′′: (A) frontal view. (A′) top view. (A′′) side view. (B-B′′) Four individual DL2d uniglomerular PNs (uPNs), each shown in a different color, innervate the CA, shown in faint grey. The axons of individual PNs split into two to three branches, each terminating in a large bouton that forms part of a highly stereotyped “claw-like” synapse with the KCs (F; Video 21). DL2d PNs terminate primarily in the anterior-lateral CA. (C-C′′) Synaptic connections, color-coded to correspond to the PNs in B-B′′, from DL2d PNs onto KCs. (D-D′′) DM1 and DC1 are shown; these uniglomerular PNs are unusual in that there is only a single cell of each type per brain hemisphere. Both DM1 and DC1 have widely distributed boutons, which occupy non-overlapping areas of the CA (Jeanne et al., 2018); DM1’s boutons are more dorsal and DC1’s more ventral. (E-E′′) Synaptic connections from the DM1 and DC1 PNs onto KCs, color-coded by PN type. (F) An example PN to KC synapse with a claw-like structure. <u>DA1 lPN</u> (1734350908; green) connects to <u>KCgm</u> (600356751; Video 21; orange); note how the KC dendrite wraps around the PN bouton. Presynaptic sites of the PN and postsynaptic sites of the KC are shown in yellow and grey respectively. (G-H) Distribution of the axons of DL2d, DM1 and DC1 in the LH and CA. (G) The four DL2d PNs innervate the middle of the LH. Note their axons terminate in the same area of the LH, but lack the large terminal boutons seen in the CA. (H) DM1 and DC1 axons remain segregated in the LH.

**Figure 9-figure supplement 2.**
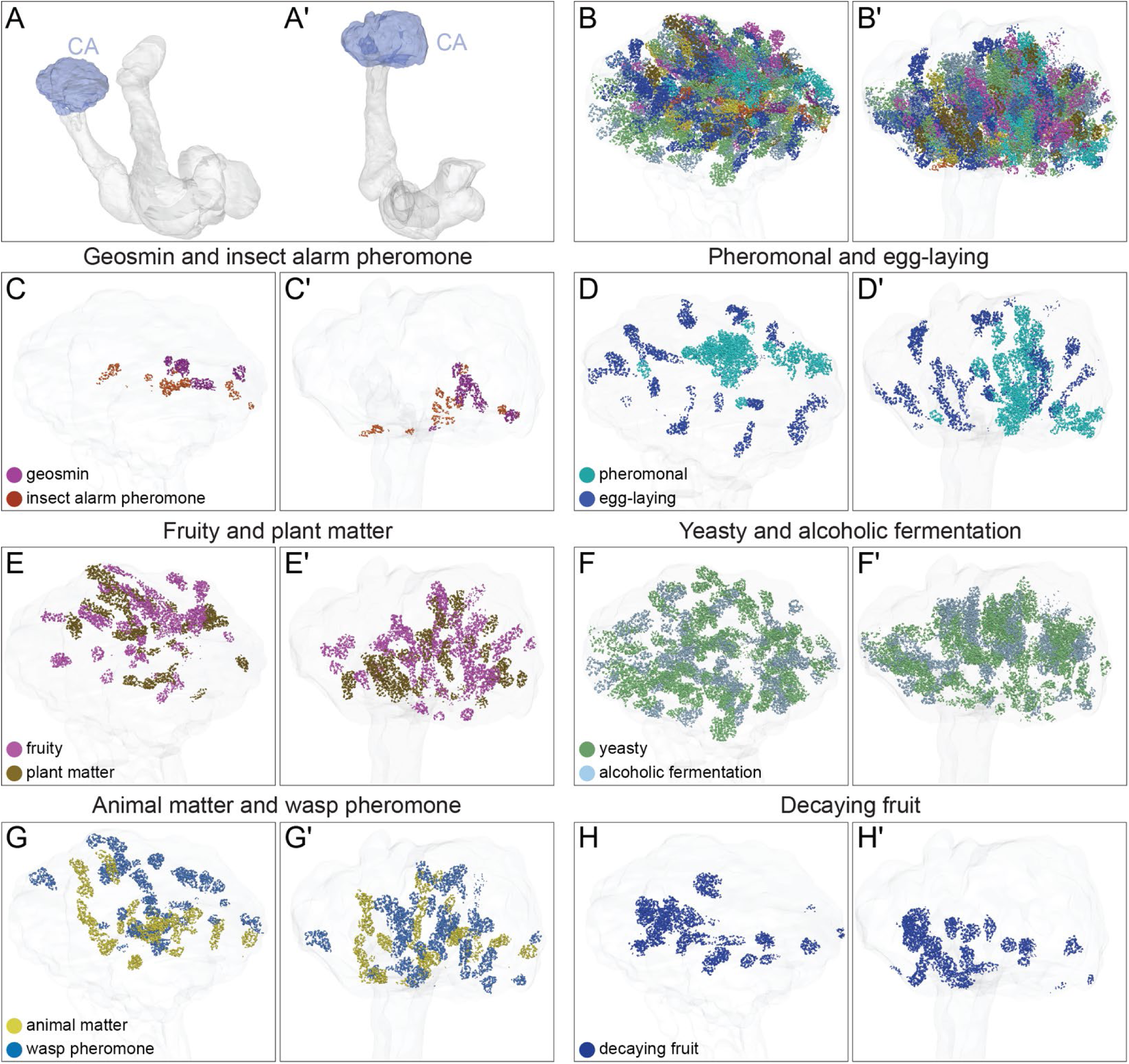
Spatial arrangement in the CA of synaptic input from different PN groups. The synaptic terminals of PNs in the CA are shown. PNs that convey distinct types of olfactory information are shown separately as indicated. (A, A′) MB images indicating the orientation of the figures shown in remaining panels: (A) frontal view; (A′) top view. The CA is shaded blue. (B, B′). Synapses KCs receive from all the olfactory PN groups that are shown individually in panel C-H′. (C, C′) PNs that convey the presence of geosmin and insect alarm pheromone, both highly aversive odors, are clustered. (D, D′) PNs that convey pheromones and odors involved in egg-laying site selection occupy the center and periphery of the CA, respectively. (E, E′) PNs that convey fruity and plant matter information are distributed in the dorsal part of the CA. (F, F′) PNs that convey yeasty and alcoholic fermentation odors are distributed widely across the CA. (G, G′) PNs that convey wasp pheromone and animal matter information. (H, H′) PNs that convey decaying fruit information.

**Figure 9-figure supplement 3.**
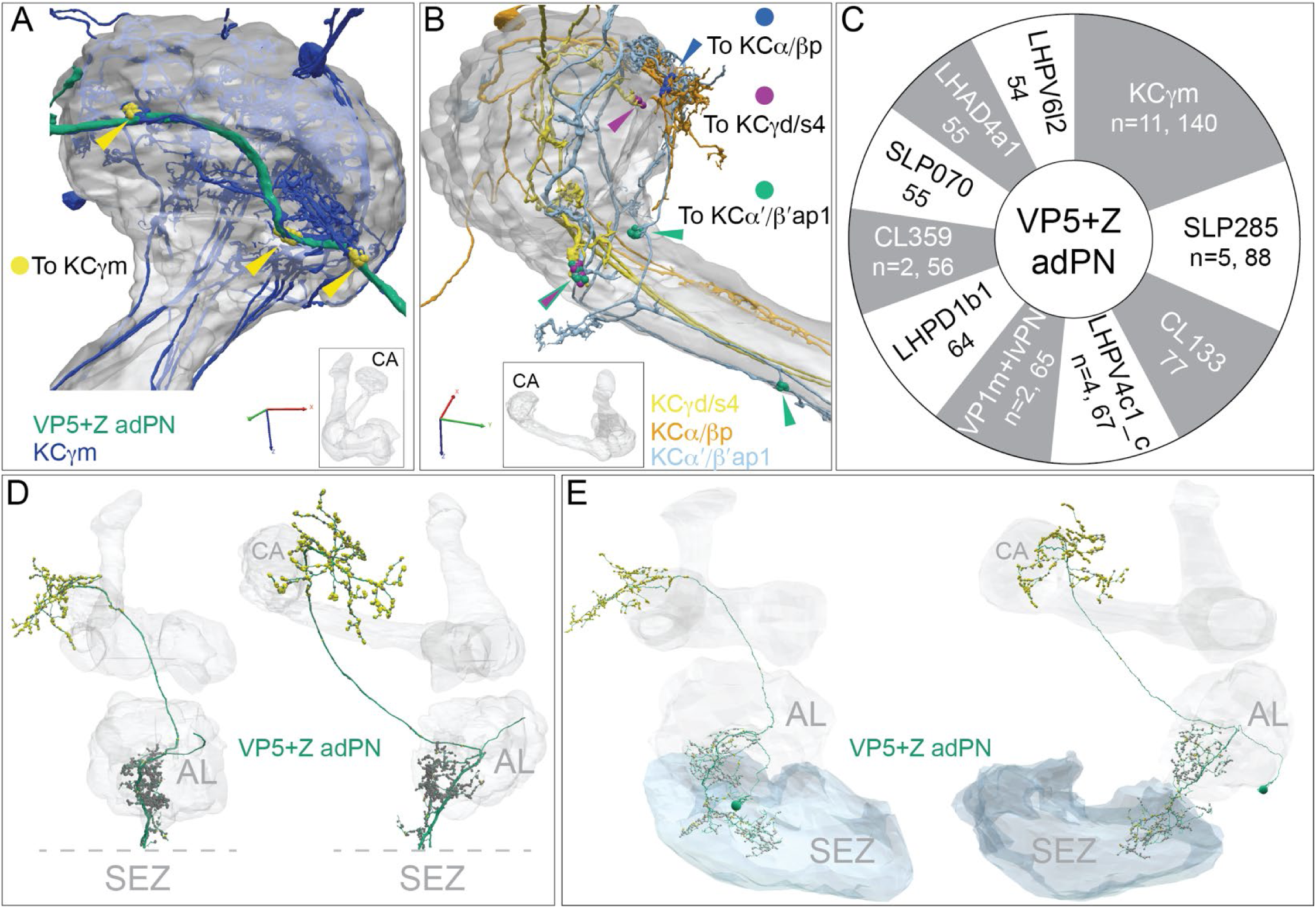
Gustatory input to a subset of KCs. The PN, <u>VP5+Z adPN</u>, has been reported to receive sensory input with mixed modalities, with hygrosensory input from the VP5 glomerulus in the AL and gustatory input from the SEZ (Marin et al., 2020). We found that it connects to 16 KCs that extend a portion of their dendritic arbors outside the main CA. (A) <u>Eleven gm</u> KCs connect at three locations to the axon of VP5+Z adPN, just anterior to the main CA (indicated by yellow arrowheads), forming a total of 140 synapses. The same 11 γm KCs also have dendritic claws inside the CA (their dendrites are dark blue where they lie outside the CA and faint blue where they lie within the CA) where they receive input from many <u>olfactory PN</u> types; their synapses from VP5+Z adPN do not have a claw-like structure. VP5+Z adPN is the top PN input to these KCs. (B) Connections from VP5+Z adPN to other KC types are shown: <u>three KCgd/s4</u> (purple, n=30); <u>two KCa/bp</u> (green, n=13) and <u>two KCa’b’ap1</u> (blue, n=12). (C) Top 10 downstream targets of VP5+Z adPN by neuron type. Note that the 11 γm KCs are its top target. (D) VP5+Z adPN in the hemibrain (left: front view, right: side view) with presynaptic sites shown in yellow and postsynaptic sites in grey. Note the SEZ is not in the hemibrain volume and its putative position is indicated below the dashed lines, which mark the ventral extent of the hemibrain volume. (E) VP5+Z adPN traced in FAFB, with its complete dendrite in the SEZ where it receives strong input from gustatory receptor neurons (S. Engert and K. Scott, personal communication). The same two views as in panel D are shown; presynaptic (yellow) and postsynaptic (grey) sites are indicated.

**Figure 10-figure supplement 1.**
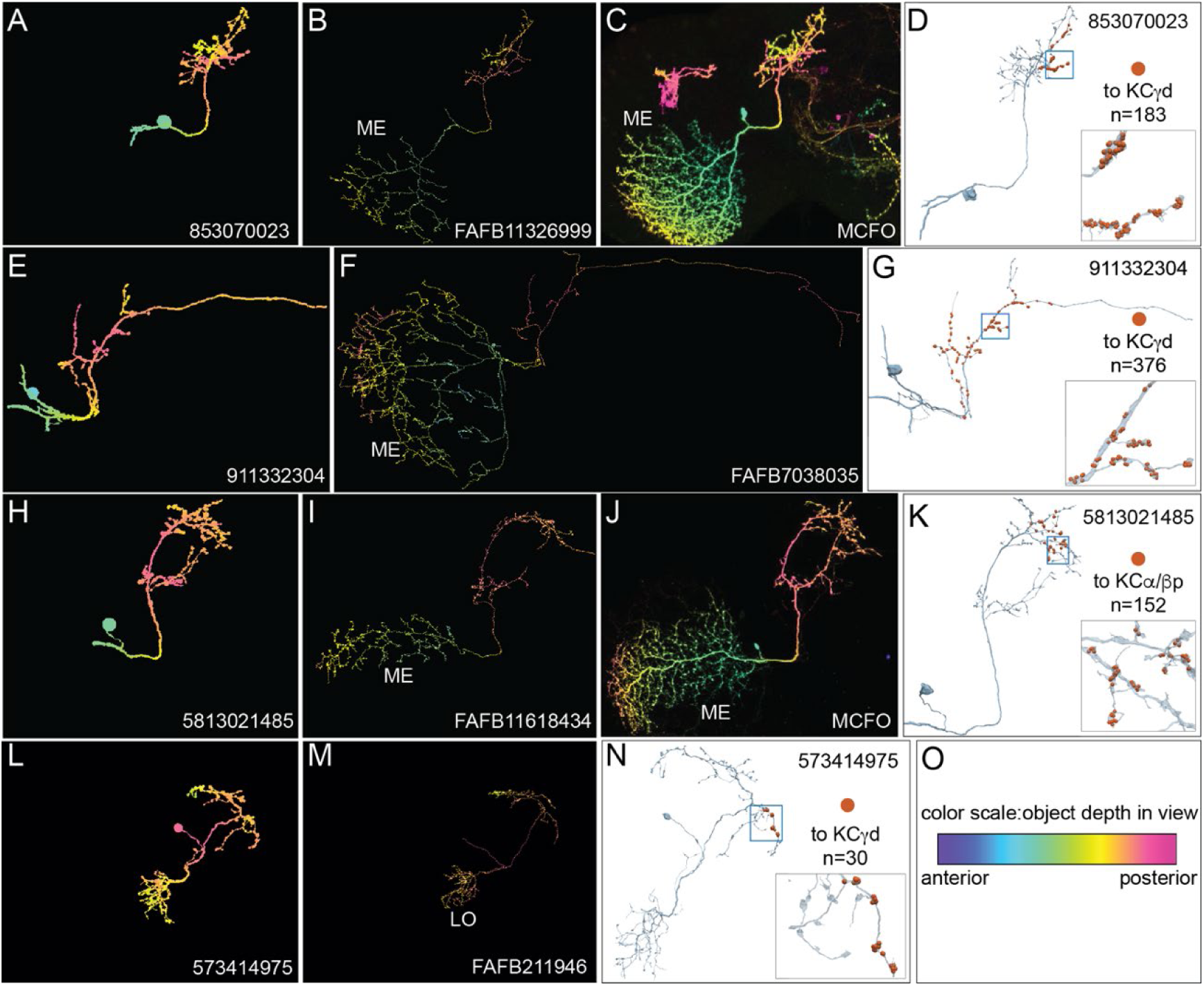
Identification of VPNs. The arbors of the ME VPNs generally extend into portions of the optic lobes that are not contained in the hemibrain volume. In order to determine the locations in the optic lobes where their dendrites are located, we relied as much as possible on matching the portion of their neuronal arbors contained in the hemibrain with morphologies of complete neurons from two other datasets: neurons that we traced in FAFB (Zheng et al., 2018) or light microscopic images of single neurons derived using Multi-Color Flp-Out (Nern et al., 2015) of GAL4 lines. In other cases, we based our classification on the similarity of the morphology of the neuron to other VPNs and the position where its arbor left the hemibrain volume. This figure shows three examples of matching ME VPNs to their putative FAFB and MCFO counterparts (A-K) and one example showing an LO VPN that is totally contained within the hemibrain volume (L-N). The images shown in panels A-C, E,F, H-J and L,M are maximum intensity projections in which the color of the neuronal arbors reflects their anterior-posterior depth in the brain (see scale in panel O; Bailey and Clark, 1998; Ropinski et al., 2006; Otsuna et al., 2018). (A-C) An example of a ME VPN innervating the vACA from the hemibrain (A) and its likely equivalents in FAFB (B) and MCFO light data (C). (D) The reconstructed neuron from the hemibrain showing the positions on its axon of the 183 synapses that it makes onto a total of 20 γd KCs indicated by the orange dots; inset, enlarged view of the boxed area. (E-F) A second example of a vACA- innervating ME VPN from the hemibrain dataset (E) and its likely match in FAFB (F). (G) The reconstructed neuron from the hemibrain with the positions on its axon of the 376 synapses that it makes onto a total of 27 γd KCs and the single γs KC indicated by the orange dots; inset, enlarged view of the boxed area. (H-J) An example of a dACA-innervating ME VPN (H) and its likely matches in FAFB (I) and MCFO (J). (K) The reconstructed neuron from the hemibrain showing the positions on its axon of the 152 synapses it makes onto a total of 41 α/βp KCs; inset, enlarged view of the boxed area. (L-N) A vACA- innervating Lo VPN (L) and its likely matches in FAFB (M). Note how the entire arbor appears to be present in the hemibrain. (N) The reconstructed neuron from the hemibrain showing the positions on its axon of the 30 synapses that it makes onto a total of 2 γd KCs; inset, enlarged view of the boxed area. (O) Color-depth scale.

**Figure 10-figure supplement 2.**
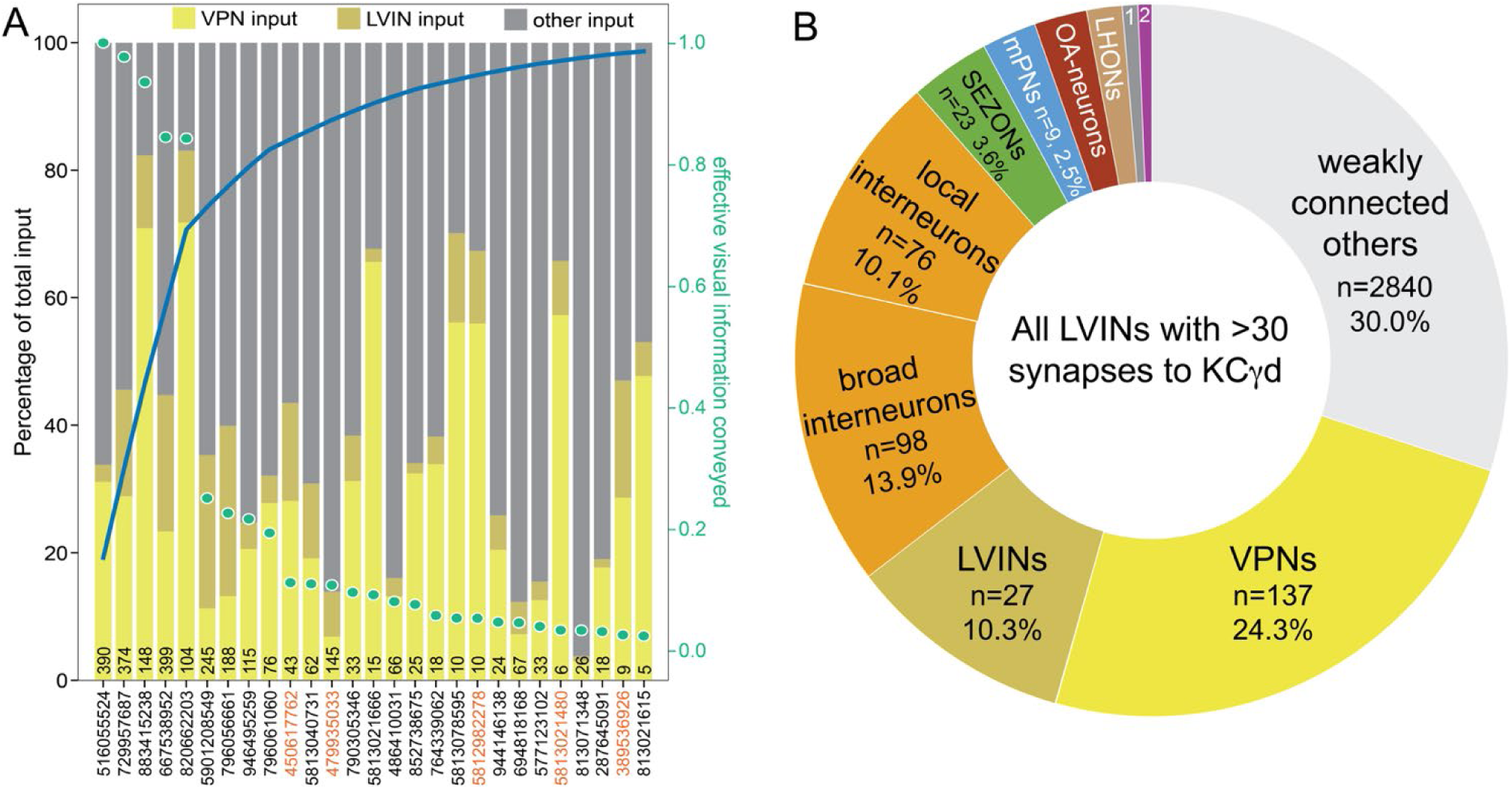
LVINs ranked by amount of visual information conveyed. (A) Plot of interneurons carrying visual information to the γd and γs1 KCs ranked by the strength of their effective contribution of visual information. For a given interneuron, this quantity is computed by multiplying the number of synapses that interneuron makes onto γd and γs1 KCs by the fraction of its input synapses that come from visual projection neurons. The green dots show this effective visual information quantity. Values are normalized to the most strongly connected LVIN, which was assigned a value of 1.0 (green scale on the right side of the plot). The blue line shows the cumulative amount of effective visual information conveyed by LVINs; over 80% of the input is delivered by the top ten LVINs. Links to these 10 LVINs in neuPrint are as follows: <u>CL063</u>, <u>PLP095</u>, <u>PLP145</u>, <u>PLP120</u>, <u>CL258</u>, <u>CL200</u>, and <u>AVLP043</u>. The color-coded bars indicate the percentage of that neuron’s input that comes from VPNs, other LVINs, and other neurons. The black number at the base of each bar shows the number of synapses made by that LVIN to KCs in the vACA. The LVINs whose ID numbers are shown in orange also provide input to the α/βp KCs in the dACA. Note that many of the lower ranked LVINs that receive high levels of VPN input do not make strong connections to KCs and are therefore presumably primarily performing some other role in integrating visual information. (B) Effective input delivered by the 15 LVINs that are the most strongly connected to the 99 γd and one γs1 KCs that are found in the vACA; each of these LVINs makes a total of 30 or more synapses to vACA KCs. Effective input is calculated by multiplying the indicated neuronal population’s synaptic input to each of the 15 LVINs times the fraction of the 99 γd and one γs1 KCs’ total input coming from that LVIN, and then summing across all 15 LVINs. Effective input is expressed as the percentage of the total input from these 15 LVINs to the 99 γd and one γs1 KCs that originated with the indicated neuronal population. To calculate the values presented, we considered the 408 neurons that collectively account for 70% of the total input synapses to these 15 LVINs and placed them into the categories shown in the pie chart; we did not attempt to classify the large number of diverse neurons (weakly connected others) that make up the other 30% of the input. VPNs, at 24.3%, provide the strongest category of effective input to vACA KCs mediated by LVINs. This visual input is in addition to direct input that VPNs make onto the dendrites of the vACA; taking the direct and indirect pathways together, more than half the total input, and ∼70% of the sensory input to the vACA is visual. We classified interneurons that did not receive visual input as local if their arbors were confined to the brain area around the vACA, as we observed for the LVINs, or broad, if they arborized in several brain areas. Although there is no direct input to γd or γs1 KCs from the SEZ, a group of 13 putative <u>SEZONs</u> contributes 2.6% of the LVINs effective input. Other inputs include <u>OA neurons</u> (n=6, 2.4%), <u>LHONs</u> (n=22, 1.6%), <u>5-HT neurons</u> (n=6, 0.7%; numbered sector 1), and APL and DPM (0.6% combined; numbered sector 2).

**Figure 10-figure supplement 3.**
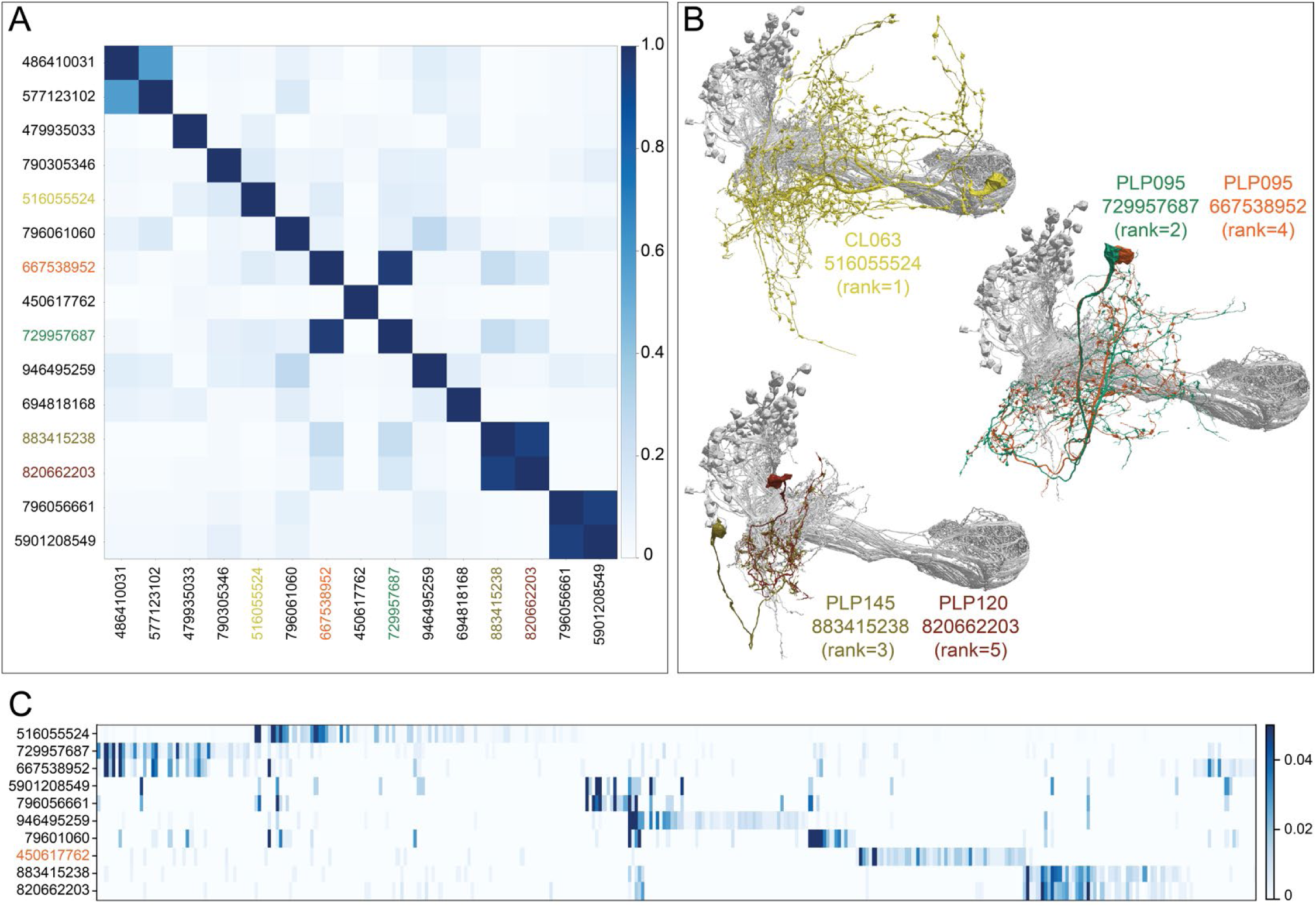
Further description of LVINs upstream of vACA KCs. (A) A plot showing the similarity of the 15 LVINs most strongly connected to the vACA KCs, based on the cosine similarity of their inputs. The neuron IDs are color coded to match the morphologies shown in (B). (B) The morphologies of the top five LVINs are shown; KCs are in grey. The orientation of the images is the same as in the inset shown in Figure 10B. (C) A plot showing the VPN inputs for each of the top ten LVINs, using the ranking of effective visual input from Figure 10-figure supplement 1A. Each vertical blue bar represents a different VPN and the intensity of the color reflects the fraction of the corresponding LVIN’s total VPN input that is contributed by that VPN. The order of the VPNs and LVINs is determined using spectral clustering.

**Figure 10-figure supplement 4.**
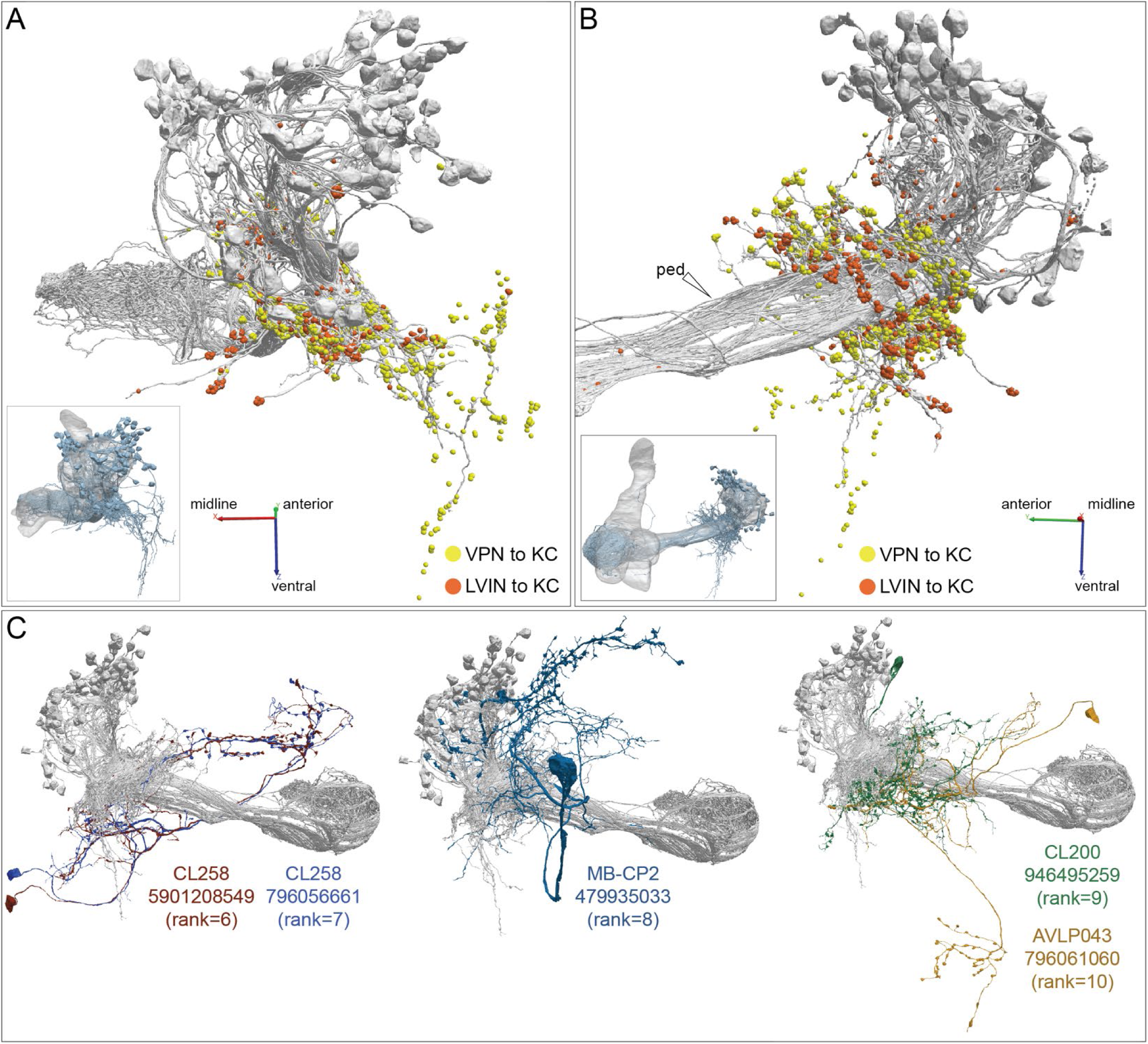
Additional views of the morphologies of VPN and LVIN inputs to the vACA. (A-B) Additional views of the distribution of VPN and LVIN inputs onto γd and γs1 KCs (grey); synapses are color-coded as indicated. The insets show the orientation of the view shown. Note the spatial segregation of these two types of connections. VPNs onto KCyd dendrites in an area ventral to the CA, previously recognized as the vACA (Butcher et al., 2012), as well as in a diffuse ring surrounding the base of the CA; synapses from LVINs are restricted to the ring. An additional view is shown in Figure 10B. (C) The morphologies of the LVINs ranked sixth to tenth in conveying the highest visual input to the vACA are shown; KCs are in grey. The orientation of the images is the same as the inset shown in Figure 10B.

**Figure 10-figure supplement 5.**
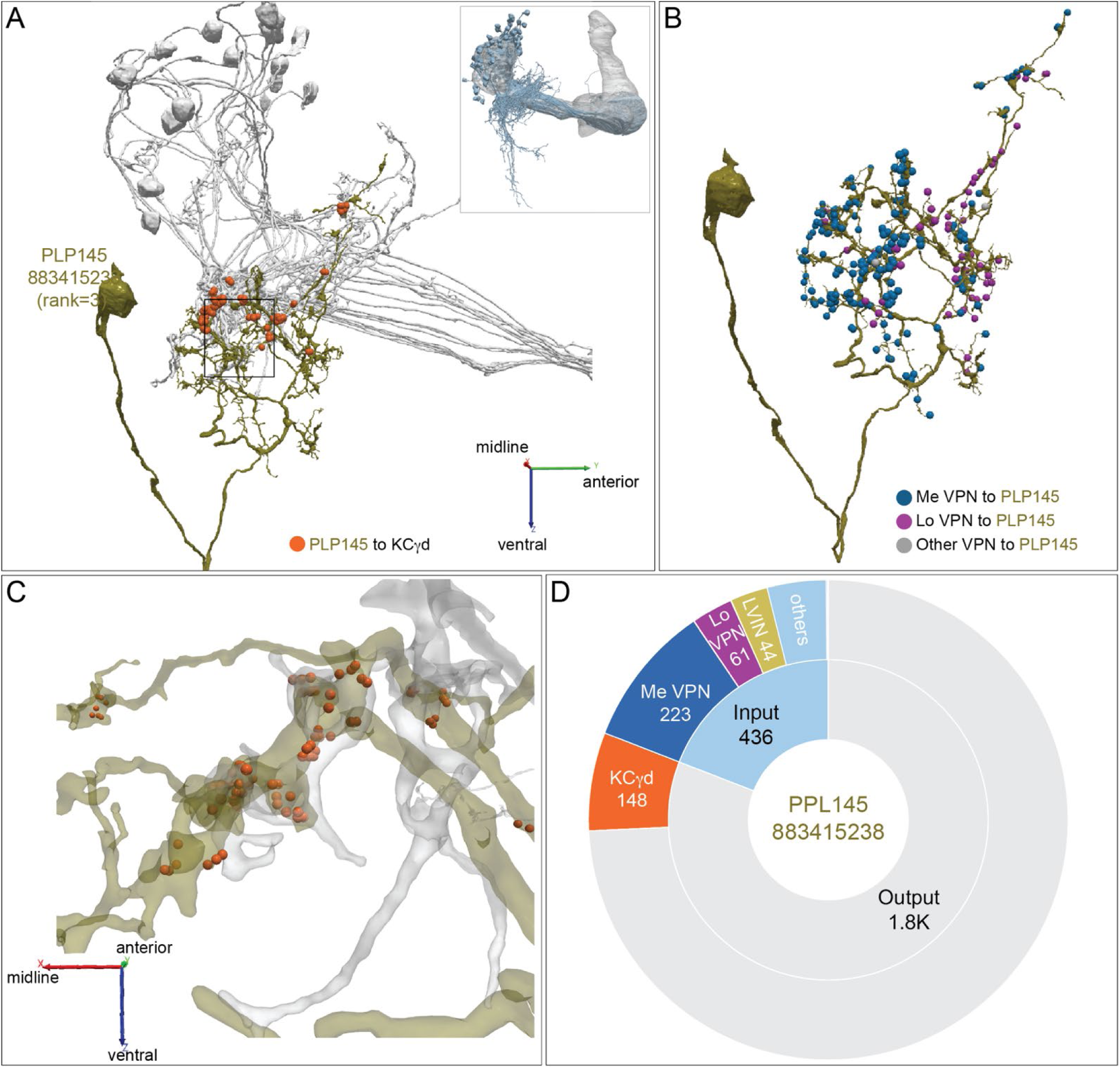
Distribution of VPN inputs onto an LVIN. (A) The LVIN <u>PLP145</u> (883415238; green) that conveys the third highest amount of visual input to the vACA is shown along with the positions where it makes synaptic output (orange dots) onto a subset of γd KCs (grey). The inset shows the orientation of the MB in this image. (B) PLP145 receives synaptic input from three classes of VPNs: ME, LO, other, shown color-coded. Other VPNs include cells that could not be placed into the ME or LO groups based on their arbors in the hemibrain volume or that appear to have dendrites in both ME and LO. (C) Enlarged and rotated view of the boxed area in (A). Note that synaptic connections from PLP145 to γd KCs lack the claw-like structure typical of olfactory PN to KC connections in the CA. (D) Pie chart showing the portion of the total input to PLP145 (light blue) that comes from direct ME (dark blue) and LO VPNs (purple), other LVINs (yellow) and other neurons (blue); the portion of PLP145’s total output (grey) that goes to KCγd (orange) is also shown.

**Figure 11-figure supplement 1.**
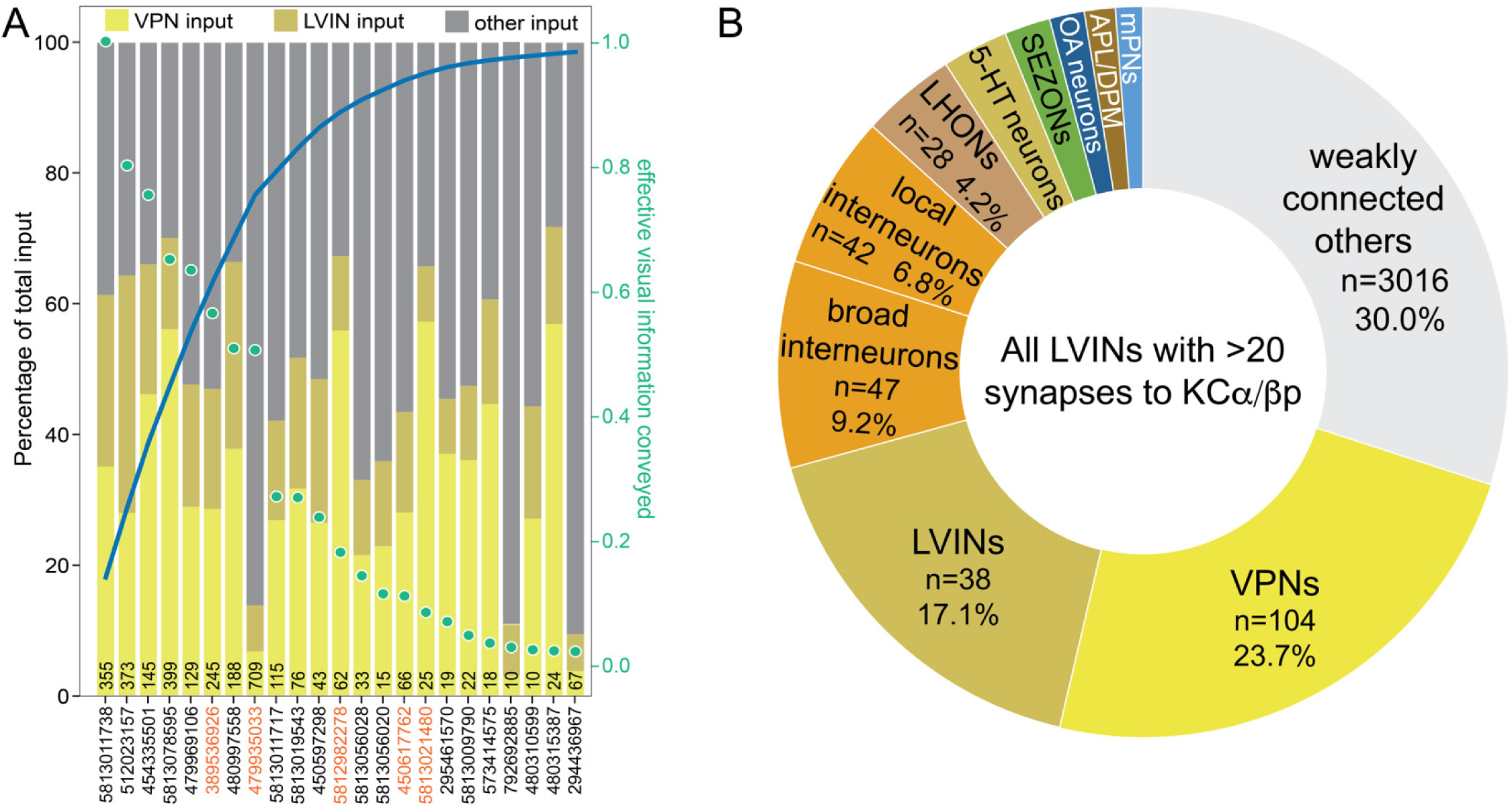
VPN and LVIN inputs to the dACA. (A) Plot of interneurons carrying visual information to the α/βp KCs ranked by the strength of their effective contribution of visual information. For a given interneuron, this quantity is computed by multiplying the number of synapses that interneuron makes onto α/βp KCs by the fraction of its input synapses that come from visual projection neurons. The green dots show this effective visual information quantity. Values are normalized to the most strongly connected LVIN, which was assigned a value of 1.0 (green scale on the right side of the plot). The blue line shows the cumulative amount of effective visual information conveyed by LVINs; over 80% of the input is delivered by the top ten LVINs. The color-coded bars indicate the percentage of that neuron’s input that comes from VPNs, LVINs, and other neurons. The black number at the base of each bar shows the number of synapses made by that LVIN to KCs in the dACA. The LVINs whose ID numbers are shown in orange also provide input to the γd KCs in the vACA. Links to these 10 LVINs in neuPrint are as follows: <u>SLP371</u>, <u>SLP360</u>, <u>CL357</u>, <u>SLP362</u>, and <u>MB-CP (LHPV3c1)</u>. Note that many of lower ranked LVINs that receive high levels of VPN input do not make strong connections to KCs and are therefore presumably performing some other role in integrating visual information. (B) Multiple sensory pathways contribute to the LVINs that connect to α/βp KCs. Effective input delivered by local neurons that each of the 19 LVINs that make at least a total of 20 synapses to α/βp KCs are shown, all of which are LVINs except one OA-neuron (<u>329566174</u>) which makes weak connections to most KCα/βp (50 of 60 total α/βp KCs). We did not find other interneurons strongly connected to the vACA KCs that were not LVINs; that is, interneurons that did not also receive VPN input. Effective input is calculated by multiplying the indicated neuronal population’s synaptic input to each of the 19 LVINs times the fraction of the α/βp KCs’ total input coming from that LVIN, and then summing across all 19 LVINs. Effective input is expressed as the percentage of the total input from these 19 LVINs to α/βp KCs that originated with the indicated neuronal population. The number of individual cells in each population is also indicated. Seventy percent of the most strongly connected upstream neurons (total of 350) to the 19 LVINs have been classified. The strongest inputs are VPNs and LVINs contributing combined 40.8% effective input. Local (those confined to the neuropil that is adjacent to dACA) and broad (those that expand through multiple neuropils) interneurons contribute a total of 16.0% of the effective input. Other prominent inputs are <u>LHONs</u>, <u>5-HT neurons</u> (n=2, 2.9%), <u>SEZONs</u> (n=7, 2.0%), <u>OA-neurons</u> (n=3, 1.5%), APL and DMP (1.4%) and <u>mPNs</u> (n=5, 1.2%).

**Figure 11-figure supplement 2.**
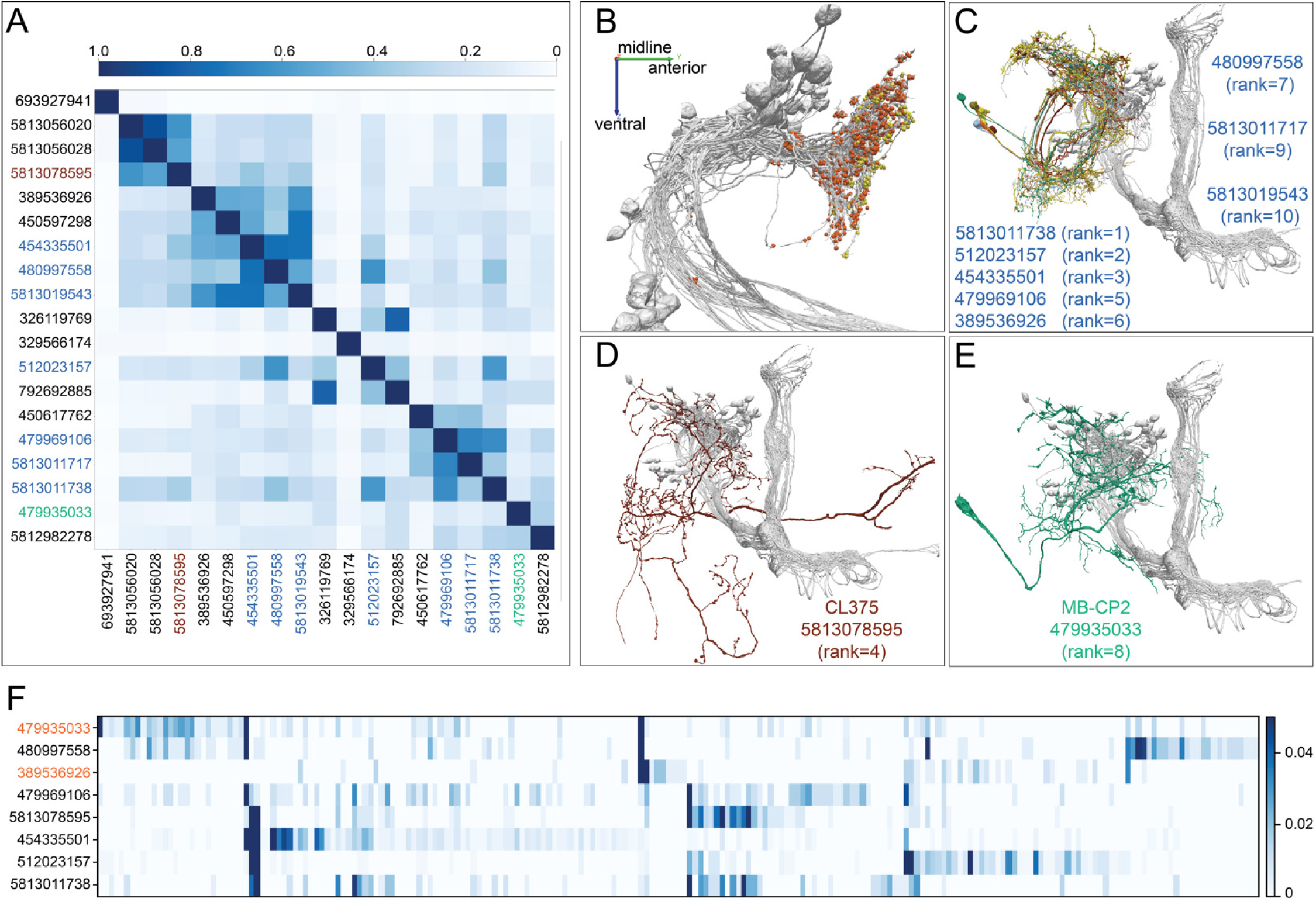
LVINs that conveys visual input onto α/βp KCs. (A) A plot showing the cosine similarity of inputs to each of the 19 LVINs that provide more than 20 synapses to α/βp KCs described in Figure 11-figure supplement 1B. (B) An additional view of the distribution of VPN and LVIN inputs onto α/βp KCs. Note that LVIN input locations tend to be more posterior than those of VPNs. (C) Of the top ten LVINs based on visual input delivered to α/βp KCs, eight are neurons with similar morphologies that fall into three cell types, SLP360, SLP363 and SLP371, shown here. (D) The LVIN conveying the fourth highest visual input to the vACA. (E) MB-CP2, the LVIN conveying the eighth highest visual input to the vACA; MB-CP2 (LHPV3c1) also provides 1% of input to KCs in the CA (see Figure 9). (F) A plot showing the similarity of VPN inputs for each of the top ten LVINs (using the ranking from panel A). Each vertical blue bar represents a different VPN and the intensity of the color reflects the percentage of the corresponding LVIN’s total VPN input that is contributed by that VPN. The order of the VPNs and LVINs was determined using spectral clustering.

**Figure 11-figure supplement 3.**
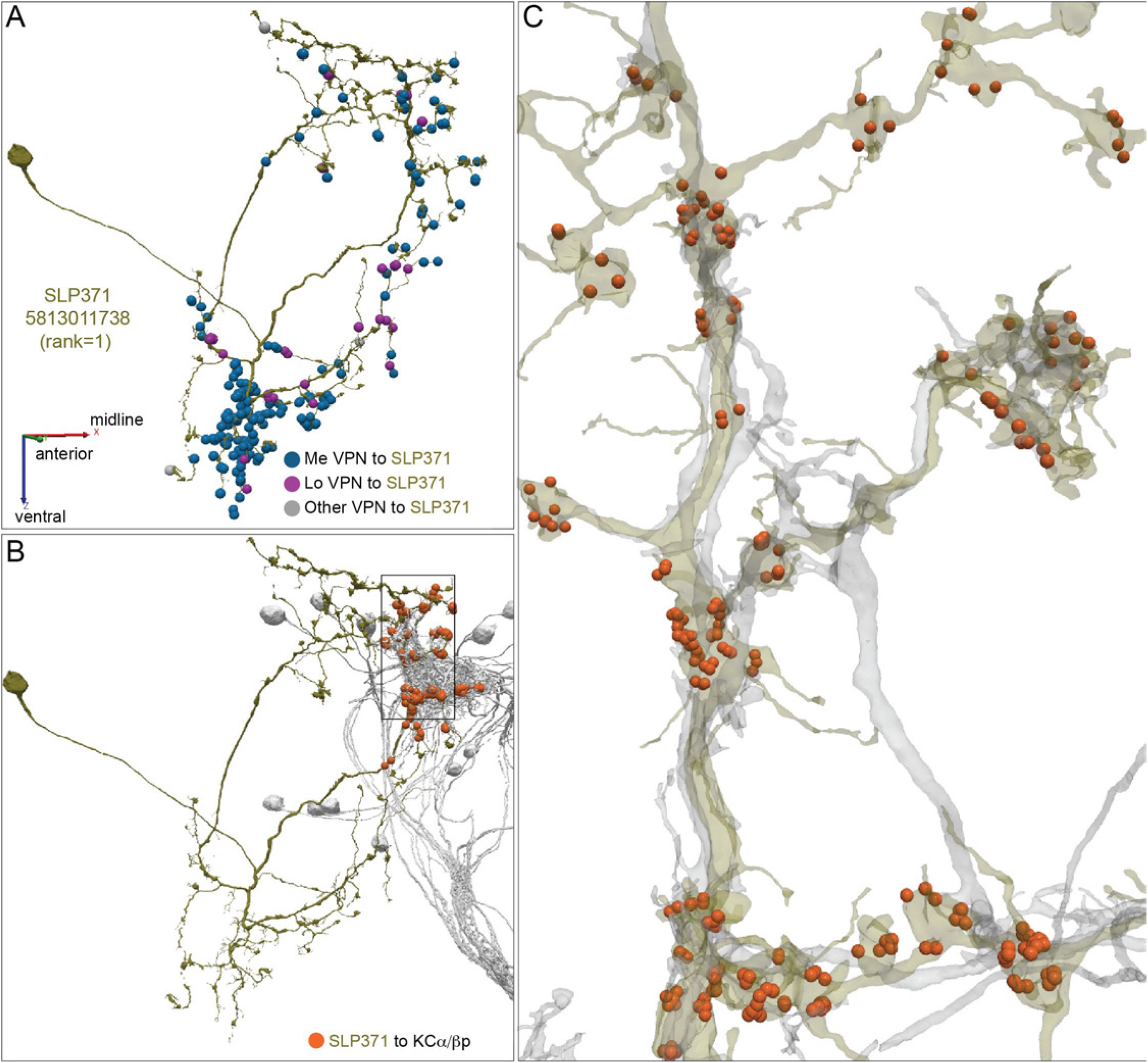
Detailed morphology of a dACA LVIN. (A) One of the two SLP371 neurons (5813011738; green) is shown with the positions of its synaptic input from three color-coded classes of VPNs. (B) Synapses from this LVIN (green) onto the subset of α/βp KCs (grey) that it contacts are shown as orange dots. (C) Enlarged and rotated view of the box in (B). Note that connections from the LVIN to α/βp KCs lack the claw-like structure typical of the synapses between olfactory projection neurons and KCs in the CA.

**Figure 11-figure supplement 4.**
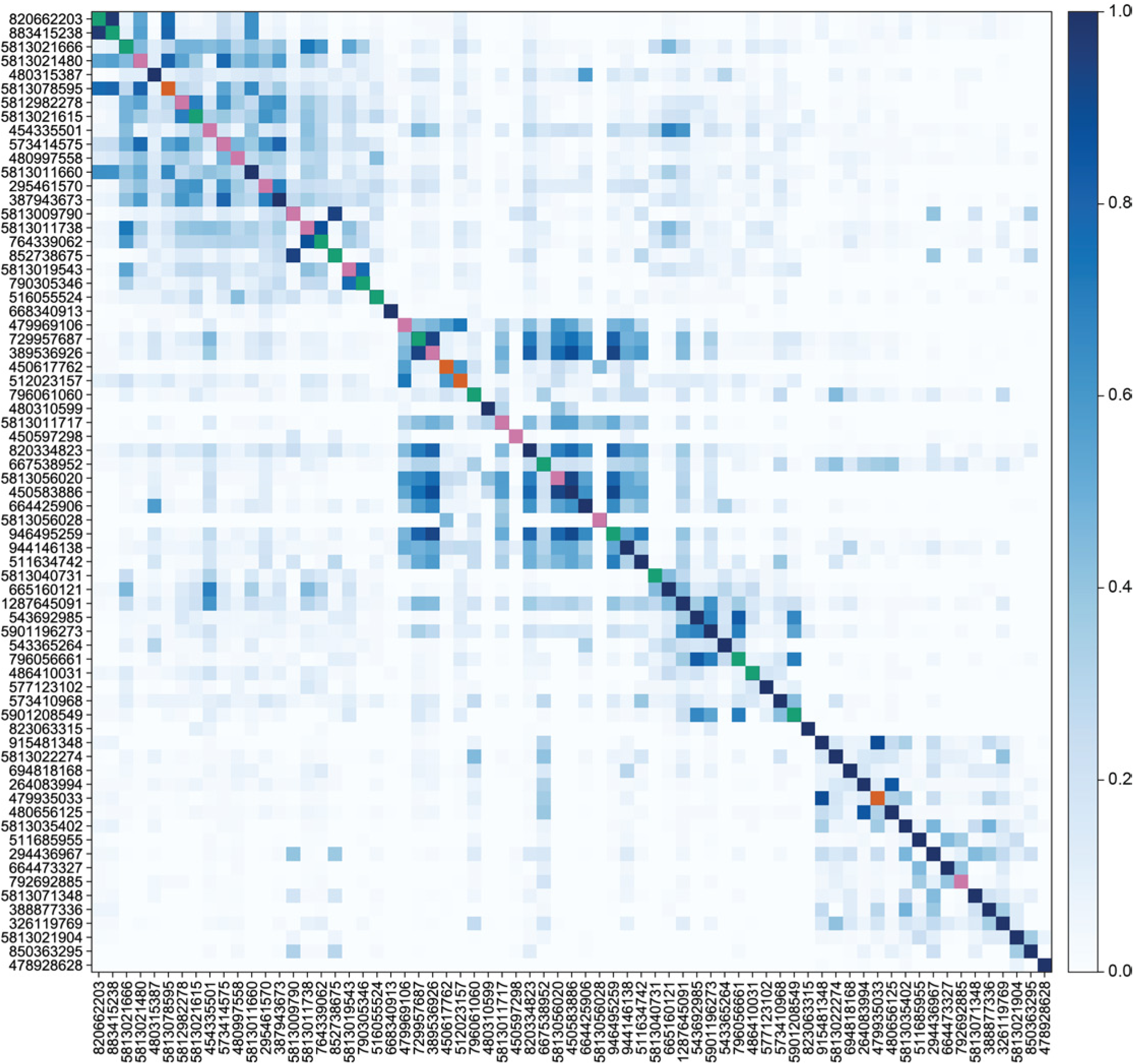
Similarity of VPN inputs to individual LVINs that innervate the vACA or the dACA. A heatmap showing the similarity of LVIN inputs from VPNs. It reveals the degree of diversity of the VPN inputs received by LVINs, as well as the existence of clusters of LVINs that receive similar inputs. Each square in the heat map represents a cosine similarity of the VPN inputs to a pair of LVINs. The ordering of the LVINs shown was determined by clustering based on their inputs. Colors on the diagonals indicate whether the given LVIN is among the top 20 inputs to the vACA (green), to the dACA (pink), to both the vACA and dACA (orange), or neither (dark blue).

**Figure 11-figure supplement 5.**
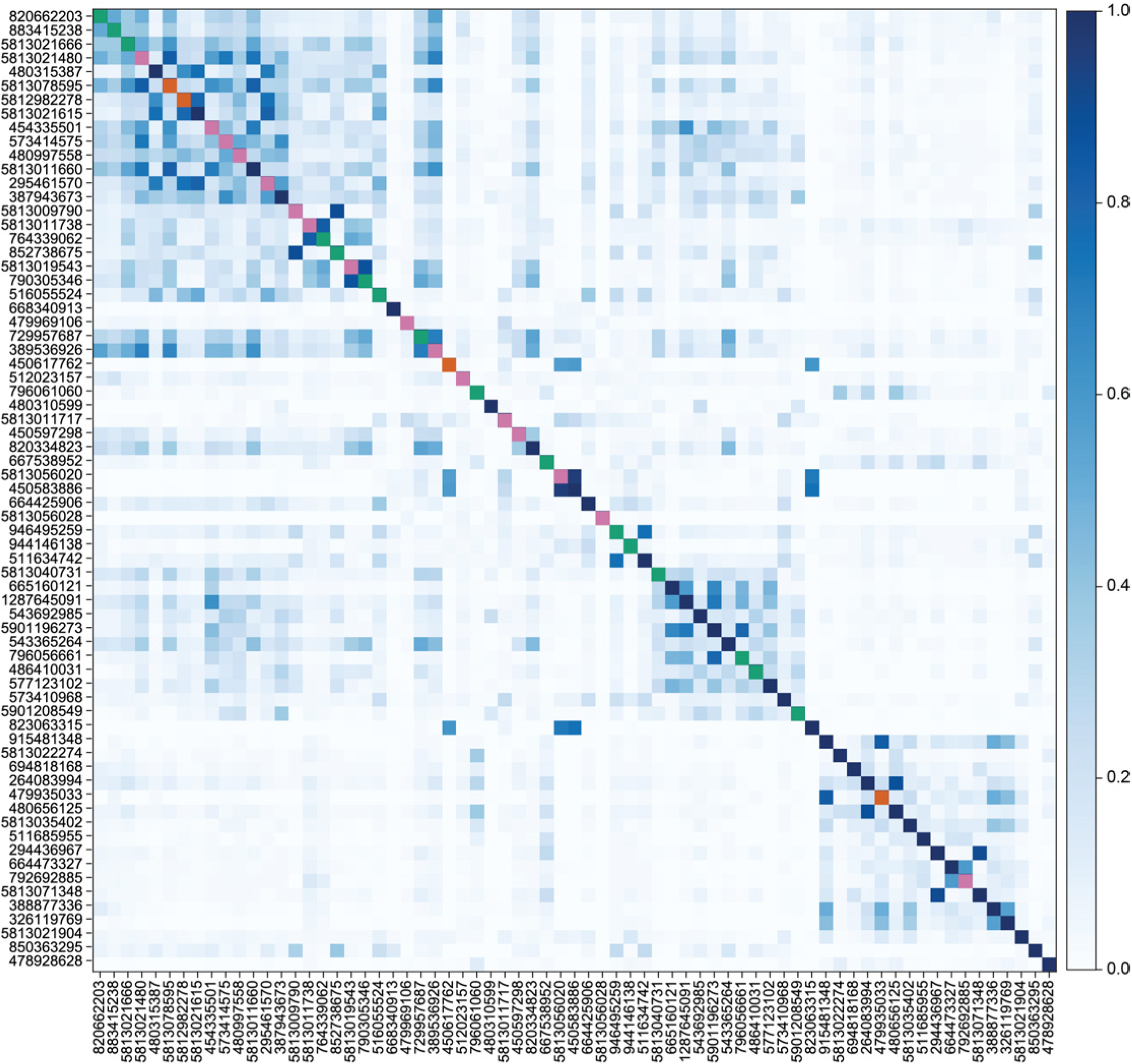
Similarity of non-visual inputs to individual LVINs. A heatmap in the same format as Figure 11-figure supplement 4 but showing the similarity of LVIN inputs from non-visual (i.e. non-VPN, non-LVIN) neurons. LVINs are shown in the same order as in Figure 11-figure supplement 4. The fact that substantial structure is preserved between these plots indicates that clusters of LVINs with similar visual inputs also tend to receive similar inputs from neurons that do not (directly) relay visual information. Colors on the diagonals indicate whether the given LVIN is among the top 20 inputs to the vACA (green), to the dACA (pink), to both the vACA and dACA (orange), or neither (dark blue).

**Figure 12-figure supplement 1.**
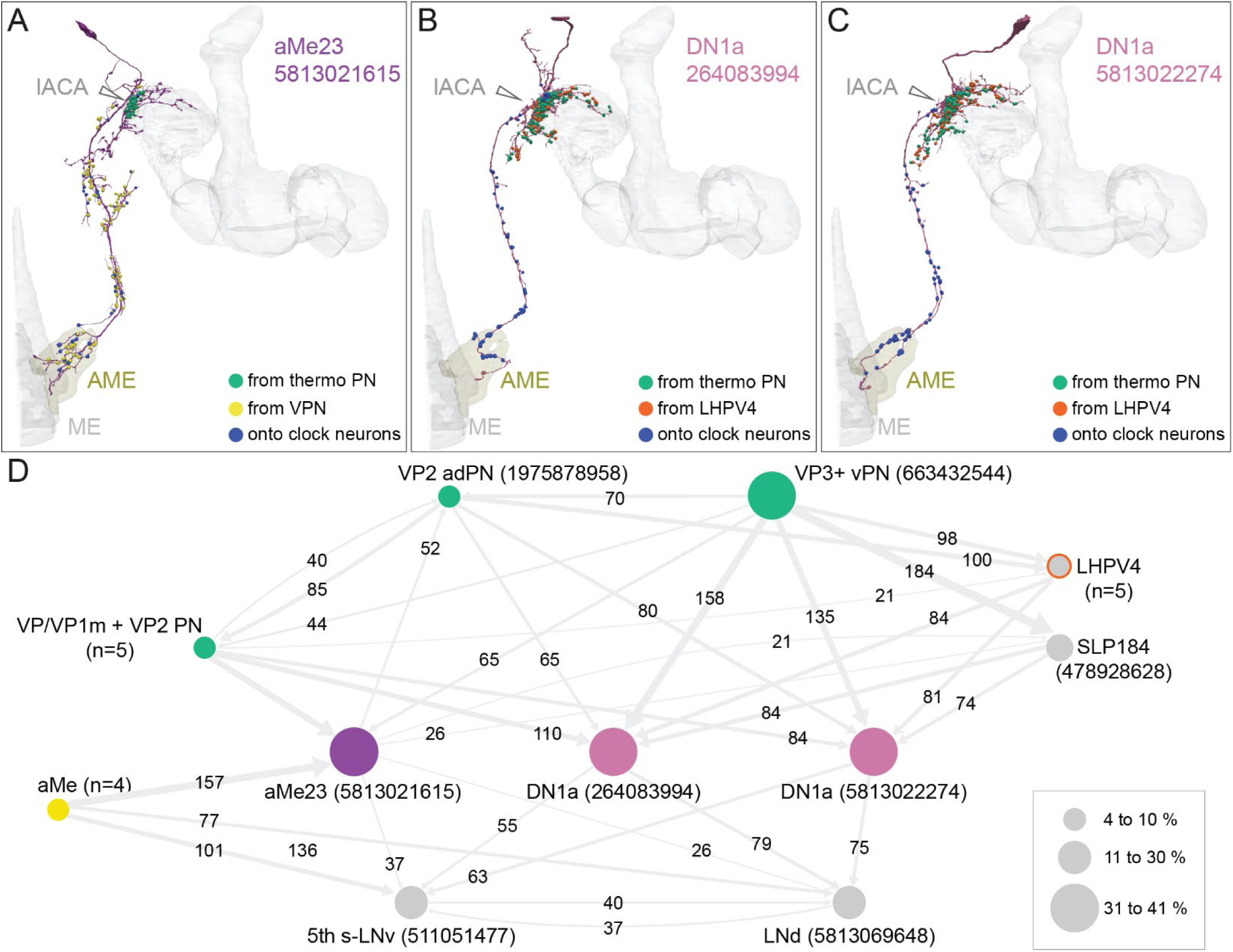
DN1a neurons relay temperature cues from the lACA to the circadian clock. (A) Morphology of a DNa1-like neuron (<u>aMe23</u>) with the positions where it receives thermosensory input in the lACA (green) and visual inputs (yellow) in the accessory medulla (AME) and the PLP. The position of its synapses onto two clock neurons, the 5th s-LNv and one LNd, are shown in blue. (B-C) Morphologies of <u>two DN1a neurons</u> with inputs from thermosensory PNs and the <u>LHPV4</u> interneuron, as well as outputs to the same two clock neurons 5th s-LNv and LNd, shown color-coded. (D). Simplified circuit diagram. The <u>aMe23</u> and DN1a neurons show different patterns of input from visual projection neurons (<u>aMe</u>; yellow), thermosensory and hygrosensory projection neurons (<u>VP/VP1m + VP2 PN</u>, <u>VP3+ vPN</u>, <u>VP2 adPN</u>; green), and interneurons (LHPV4; grey with red outline) (Marin et al., 2020). Temperature entrainment of the circadian clock requires input from the aristae (Yadlapalli et al., 2018; Alpert et al., 2020), which house the VP3 and VP2 sensory neurons (Gallio et al., 2011); the connections we observed provide a potential anatomical substrate for arista-dependent temperature entrainment of the clock. DN1a (Helfrich-Förster, 2004; Shafer et al., 2006) and aMe23 neurons provide input to the circadian clock neurons 5th <u>s-LNv</u> and <u>LNd</u>. Connections of fewer than 20 synapses have been omitted. The sizes of the three green circles representing PN types indicate the percentage of those neurons’ total output (based on synapse number) going directly to other neurons in the diagram; the sizes of the circles representing all other neurons indicate what percent of that neuron’s total input comes directly from other neurons in the diagram.

**Figure 12-figure supplement 2.**
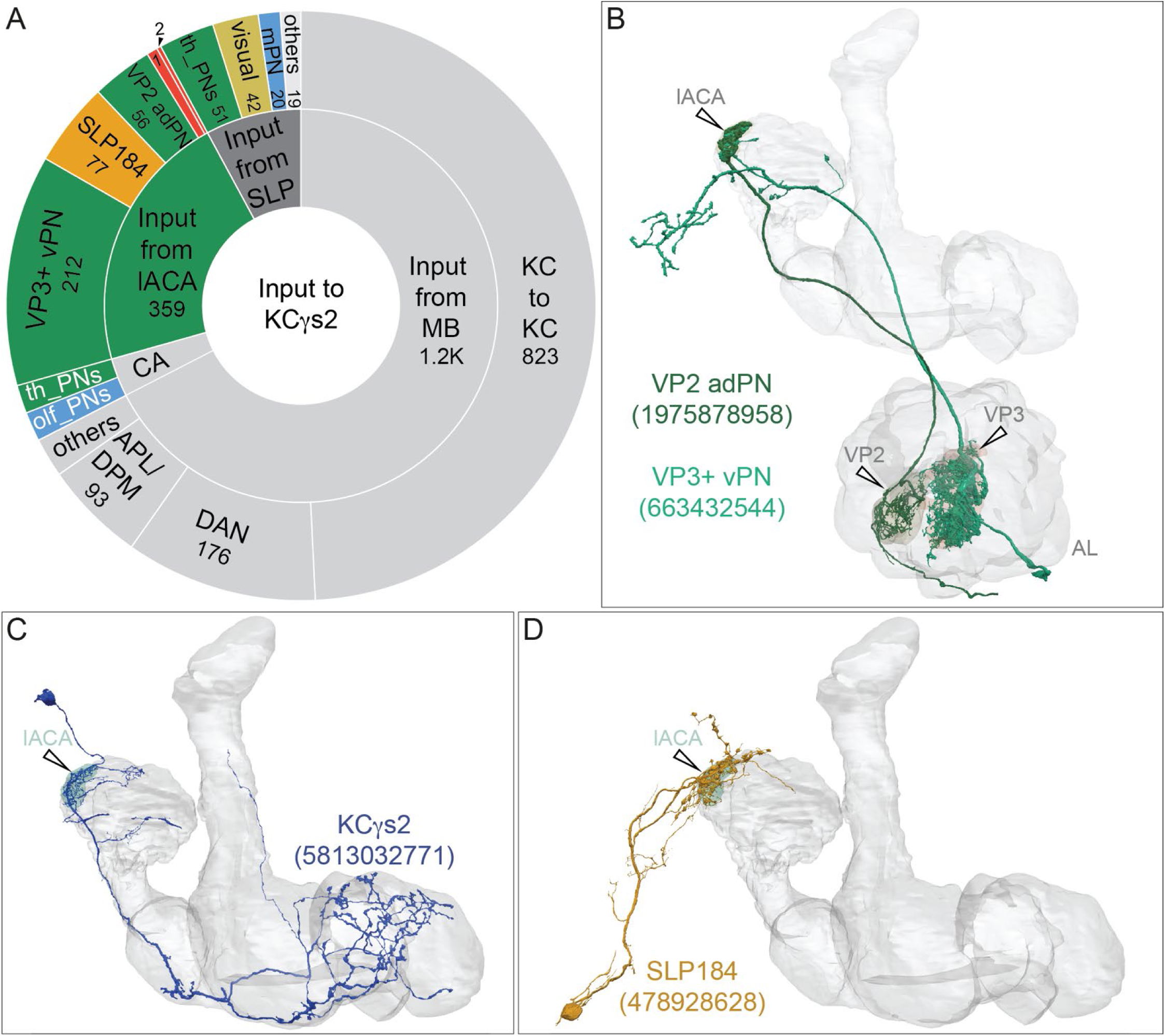
KCγs2 and its inputs. (A) Distribution of inputs to <u>KCgs2</u>. KCγs2 receives the vast majority of its sensory input in the lACA from thermo/hygrosensory PNs, but also receives a small amount of visual and olfactory information. Note that the pie chart reflects all of KCγs2’s inputs, not just those that occur in the lACA. KCγs2 receives nearly half its input from putative KC-to-KC synapses in the γ lobe. The γs2 KC receives the majority of its sensory input in the lACA from two thermal PNs (VP3+ vPN and VP2 adPN); other lACA inputs include the interneuron <u>SLP184</u> (77, 4.5%) and neurons represented by the numbered sectors: 1, aMe23 (10, 0.7%) and 2, VP1m+VP2_lvPN2 (8, 0.5%). The γs2 KC also receives 132 synapses (7.7% of its input) from its arbor that lies in the SLP, outside main CA and lACA, indicated in the grey wedge, as well as a small amount of input from other thermal and olfactory PNs in the CA. (B) Morphologies of thermosensory PNs VP2 adPN and VP3+ vPN, which are the major inputs to the lACA, are shown. (C) Morphology of KCγs2. (D) Morphology of the interneuron SLP184.

**Figure 13-figure supplement 1.**
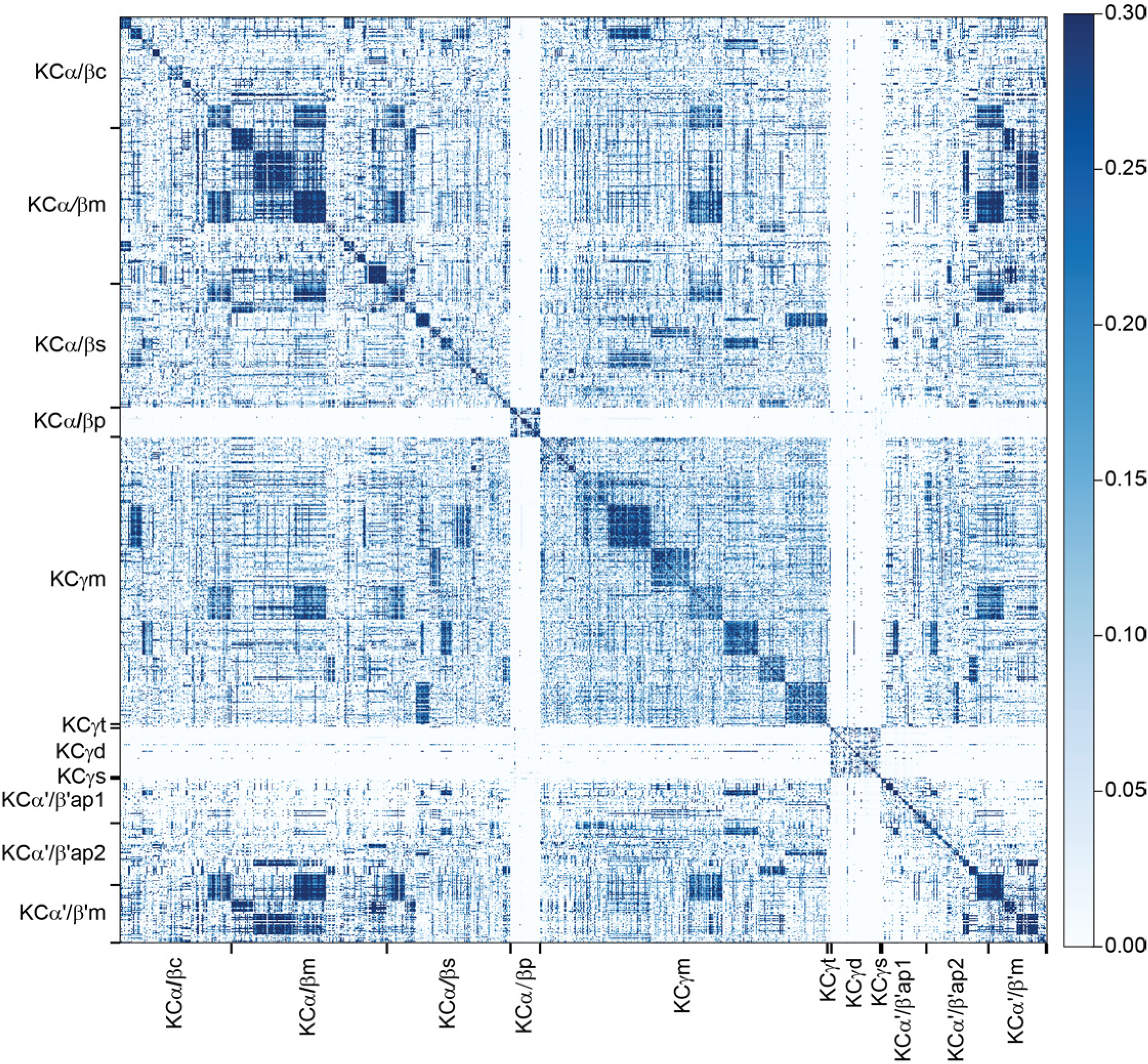
Clustering KCs based on the similarity of their inputs from PNs. Cosine similarity of PN inputs to KCs. This metric measures the degree of overlap in the PN inputs, weighted by synapse count, to each pair of KCs. Each PN is treated individually, rather than being grouped by cell type. KCs are grouped by subtype, and then the ordering within each subtype is determined by spectral clustering. The optimal number of clusters within each type is determined using the silhouette score and capped at 10. The potential computational significance of groups of KCs responsible for distinct PN inputs is modeled in Figure 37.

**Figure 13-figure supplement 2.**
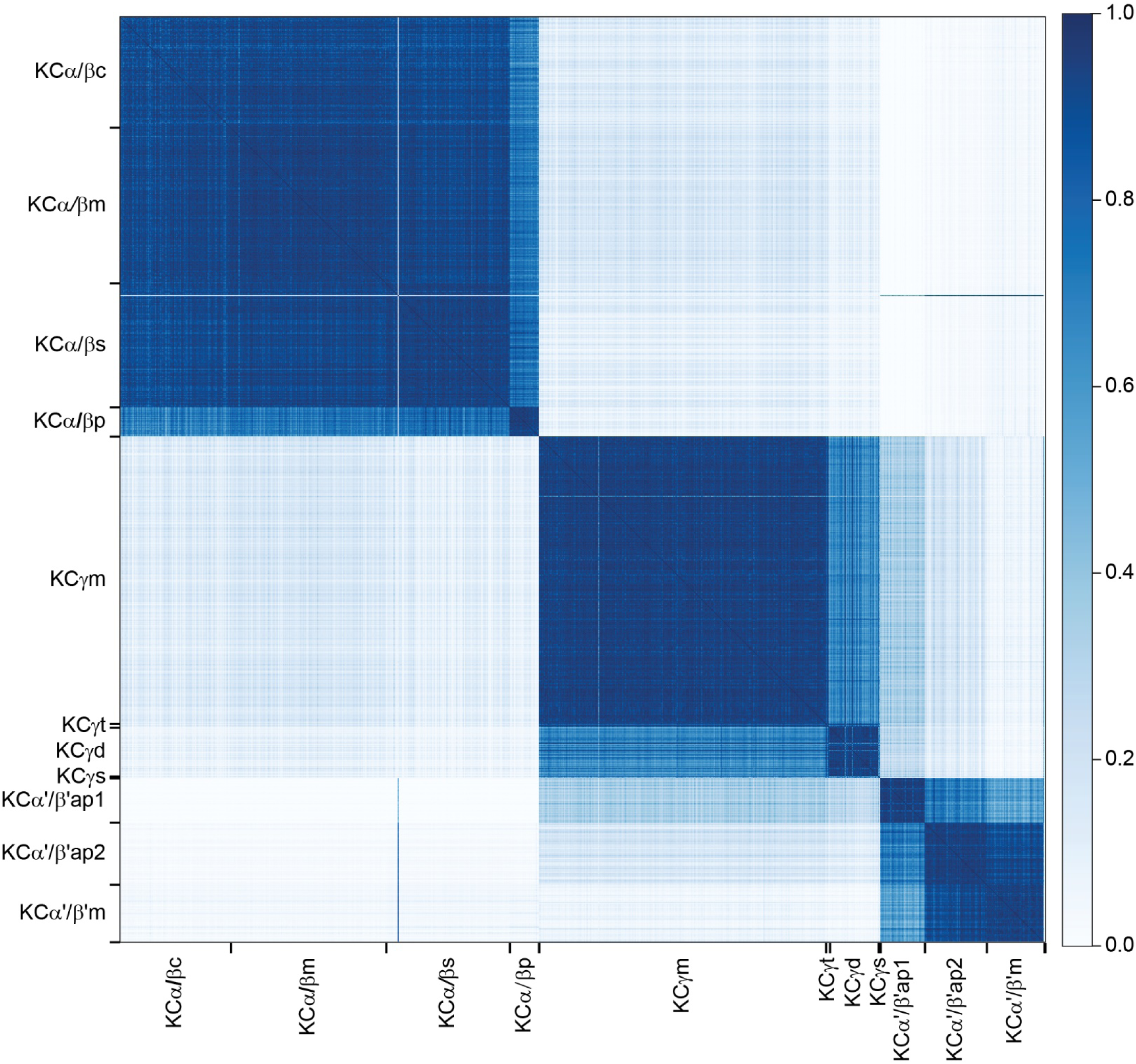
Similarity of KC outputs to MBONs. Cosine similarity of KCs outputs to MBONs. KCs are grouped by subtype, and then indexed in the same order as in Figure 13.

**Figure 15-figure supplement 1.**
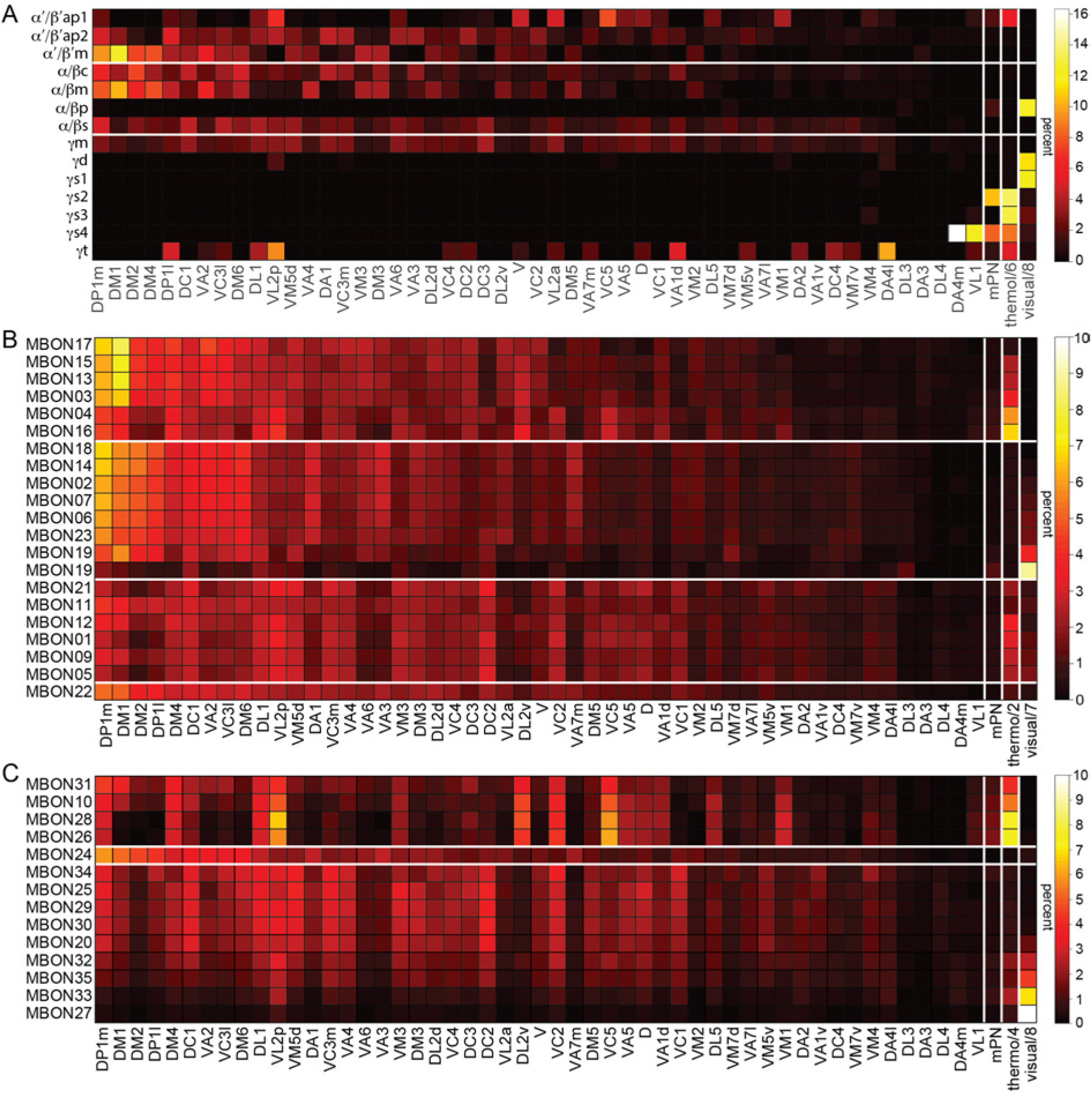
PN connections to KC types and effective PN to MBON connectivity. Detailed views of the PN inputs to KCs and MBONs. The uPN types are listed individually. Inputs from multiglomerular PNs (mPN), thermo/hygrosensory (thermo) and visual pathways have been pooled over all input neurons of each type. Percentages indicate the amount of effective input due to a particular PN type divided by the total sensory input to each target cell or population; the thermo-hygrosensory and visual results have been divided by the factors indicated in each column, to keep the color scale within reasonable bounds. (A) Inputs from uPNs, mPNs, thermo/hygrosensory and visual pathways to KCs of different subtypes. (B) Inputs from uPNs, mPNs, thermo/hygrosensory and visual pathways to each typical MBON. (C) The same plot as in (B), but for the atypical MBONs. Note that this plot reflects only input that atypical MBONs receive from KCs and ignores other input that they receive from their dendrites that extend outside the MB lobes. Approximately 80 and 55 percent of the inputs to MBON27 (γ5d) and MBON33 (γ2γ3), respectively, convey visual information (see Figure 15B). Note that input from VC5 and VL2p is highly represented by the same MBONs that receive strong input from thermo-hygrosensory PNs, supporting the idea that these glomeruli may also convey thermo-hygrosensory information (Hamada et al., 2008; Frank et al., 2015; Marin et al., 2020).

**Figure 15-figure supplement 2.**
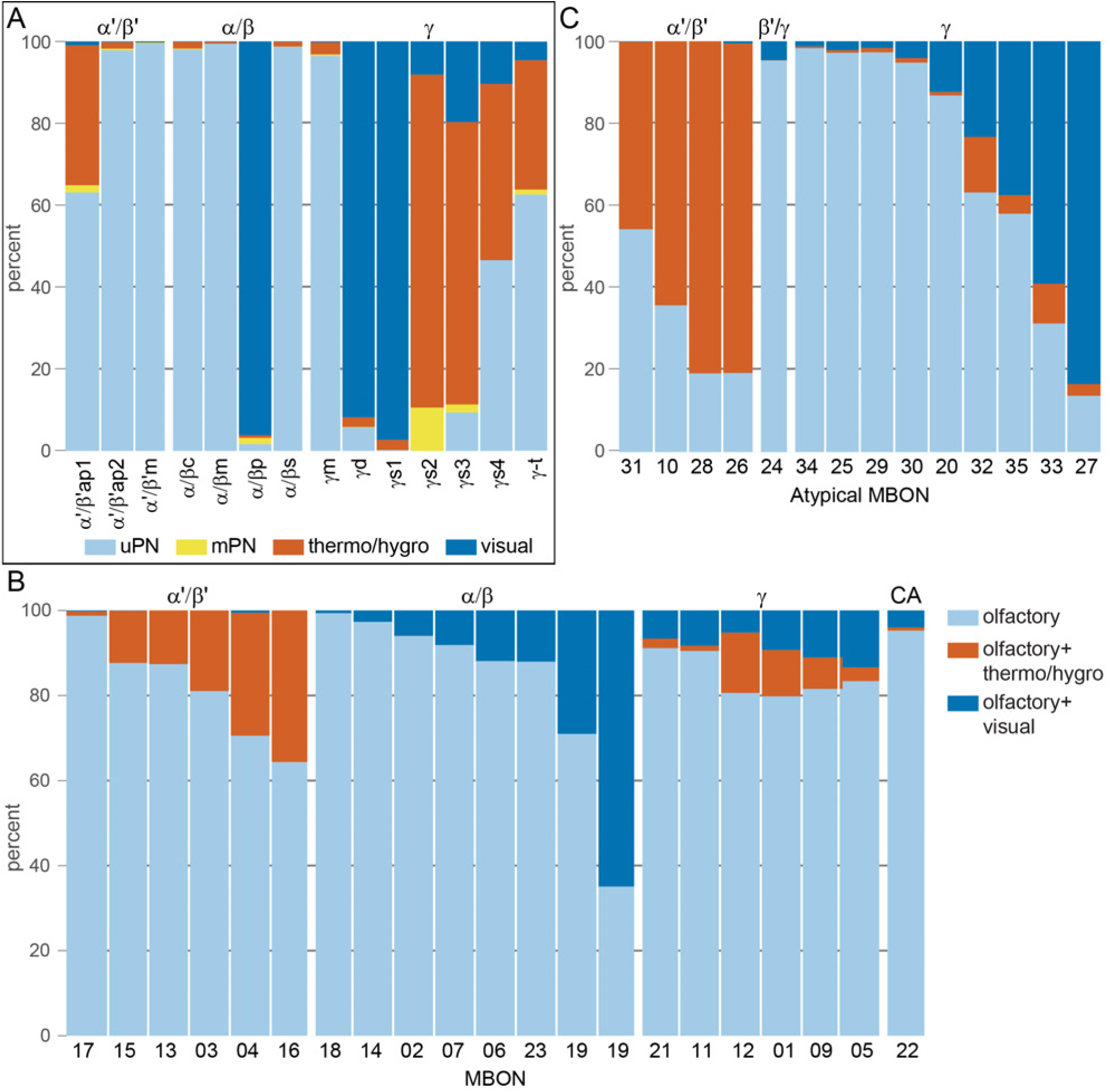
PN to KC and KC to MBON connectivity by sensory modality. (A) Olfactory input from uniglomerular PNs is carried by most of the KC types (α′/β′m, α′/β′ap2, α/βc, α/βm, α/βs, γm), but thermo-hygrosensory and visual inputs are conveyed by specific KC subtypes. In particular α/βp, γd and γs1 KCs carry substantial fractions of visual input, and α′/β′ap1, γs2-4 and γt KCs convey thermo-hygrosensory information. (B) Within the α′/β′, α/β and γ lobes, the amount of non-olfactory input to different MBONs varies across MBON types due to differential KC innervation. MBONs innervating the α′/β′ lobe received variable amounts of thermo-hygrosensory input due to different levels of α′/β′ap1 innervation. Visual input varies across the α/β MBONs due to different amounts of α/βp KC input. The γ lobe, in contrast, has a more uniform distribution of different sensory modalities across its MBONs. (C) Same as (B), but for atypical MBONs. Percentages indicate the number of synapses of a given type divided by the total number of sensory input synapses to each KC type (panel A) or KC synapses to each MBON (panels B and C). Panels B and C reflect KC input to MBONs with the KCs divided into classes according to panel A. In contrast, Figures 15A and 15B show the full input-to-output effective connectivity.

**Figure 15-figure supplement 3.**
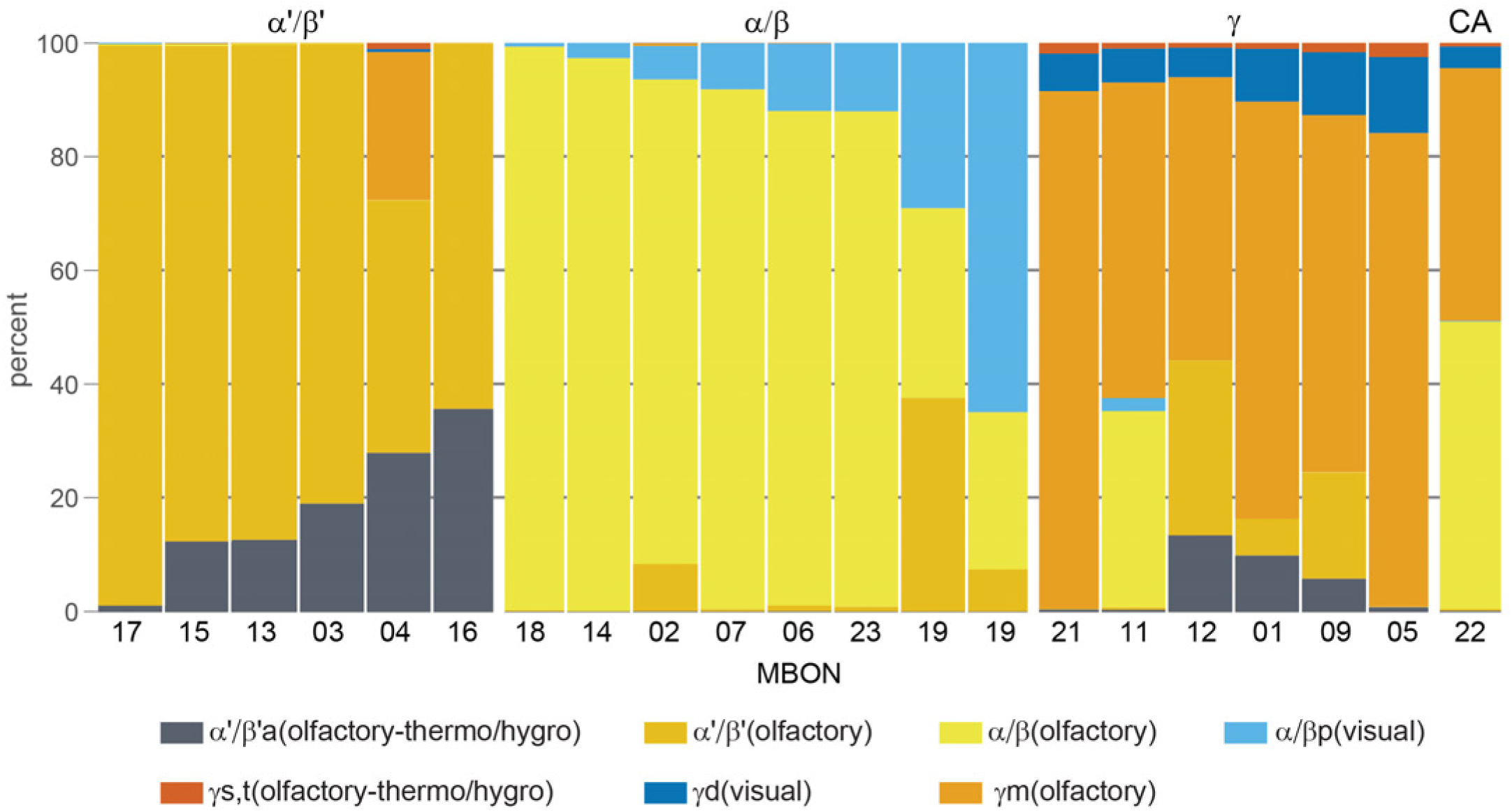
MBON connectivity by KC type. Percentage of input to typical MBONs by KC type. Similar to Figure 15-figure supplement 2C, except that the KCs have been divided by lobes as well as by sensory modality. MBONs exhibit different biases toward KC types, which underlies differences (as shown in Figure 15-figure supplement 2) in the proportion of effective input they receive from each sensory modality. We also observe a few instances of MBONs receiving substantial input from KCs in unexpected (relative to prior findings) lobes, including the large percentage of γ KC input to MBON04 (β′2mp_bilateral), and the large percentages of α′/β′ input to one of the two MBON19s (α2p3p). It is unknown if these unexpected connections are stereotyped.

**Figure 16-figure supplement 1.**
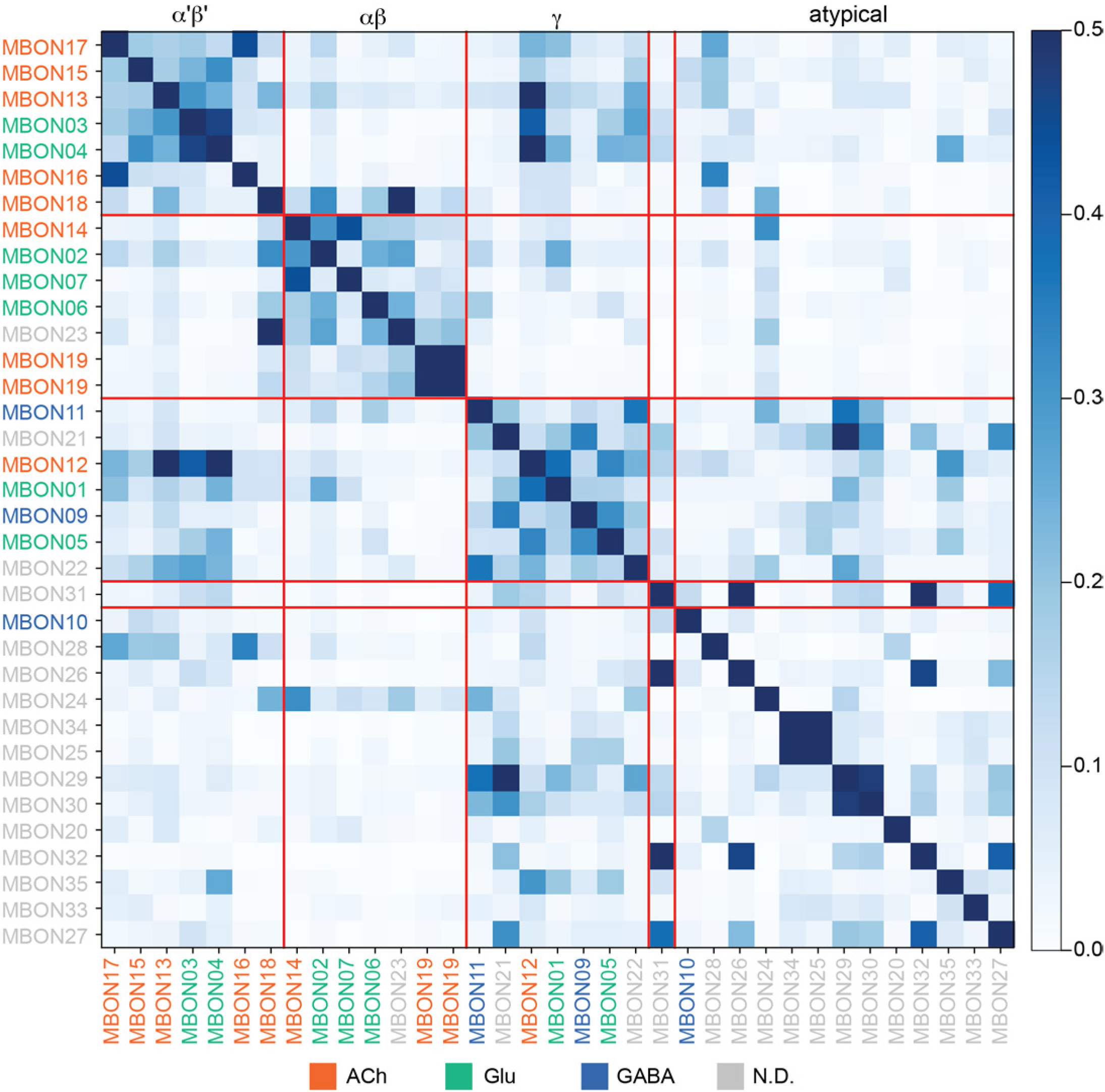
MBON output similarity by lobe. Cosine similarity of MBONs based on the similarity of their outputs to all neurons (unnamed neuronal fragments have been excluded). Each cell of the heat map indicates the output similarity of the indicated pair of MBONs. This is the same data as plotted in Figure 16, except that the MBONs are organized by lobe, shown in the same order as in Figure 15. The preservation of lobe-wise sensory input structure in the MBONs suggests that MBONs may mediate multiple parallel pathways that transform sensory information into behavioral output. The MBON names are color-coded to indicate their neurotransmitter, as indicated.

**Figure 16-figure supplement 2.**
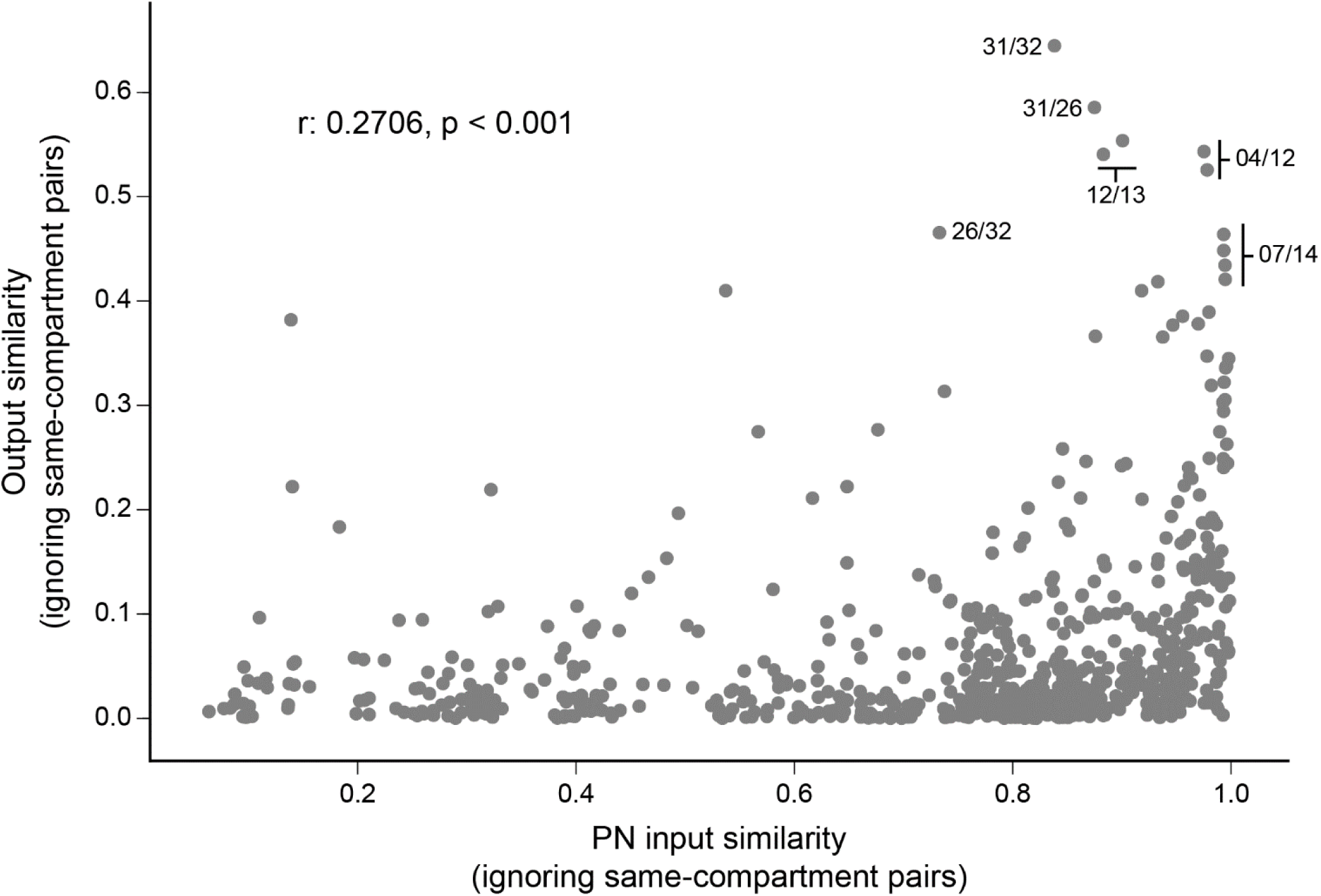
Correlation of MBON output and PN input similarity. Scatter plot of MBON output similarity vs. similarity of their inputs from PNs, quantifying the relationship between the structure of MBON outputs and their sensory inputs. Each point represents a pair of MBONs from different compartments. Output similarity is computed using the cosine similarity of outputs, as in Figure 16. PN input similarity is computed using cosine similarity of effective PN inputs, as in Figure 16. The highest ten points on the y-axis correspond to the following pairs (fewer than ten listed due to multiple MBONs of the same type): MBON31(α′1) and MBON32 (γ2). MBON26 (β′2d) and MBON31 (α′1), MBON12 (γ2α’1) MBON13 (α′2). MBON04 (β′2mp_bilateral) and MBON12 (γ2α′1), MBON26 (β′2d) and MBON32 (γ2), MBON07 (α1) and MBON14 (α3).

**Figure 16-figure supplement 3.**
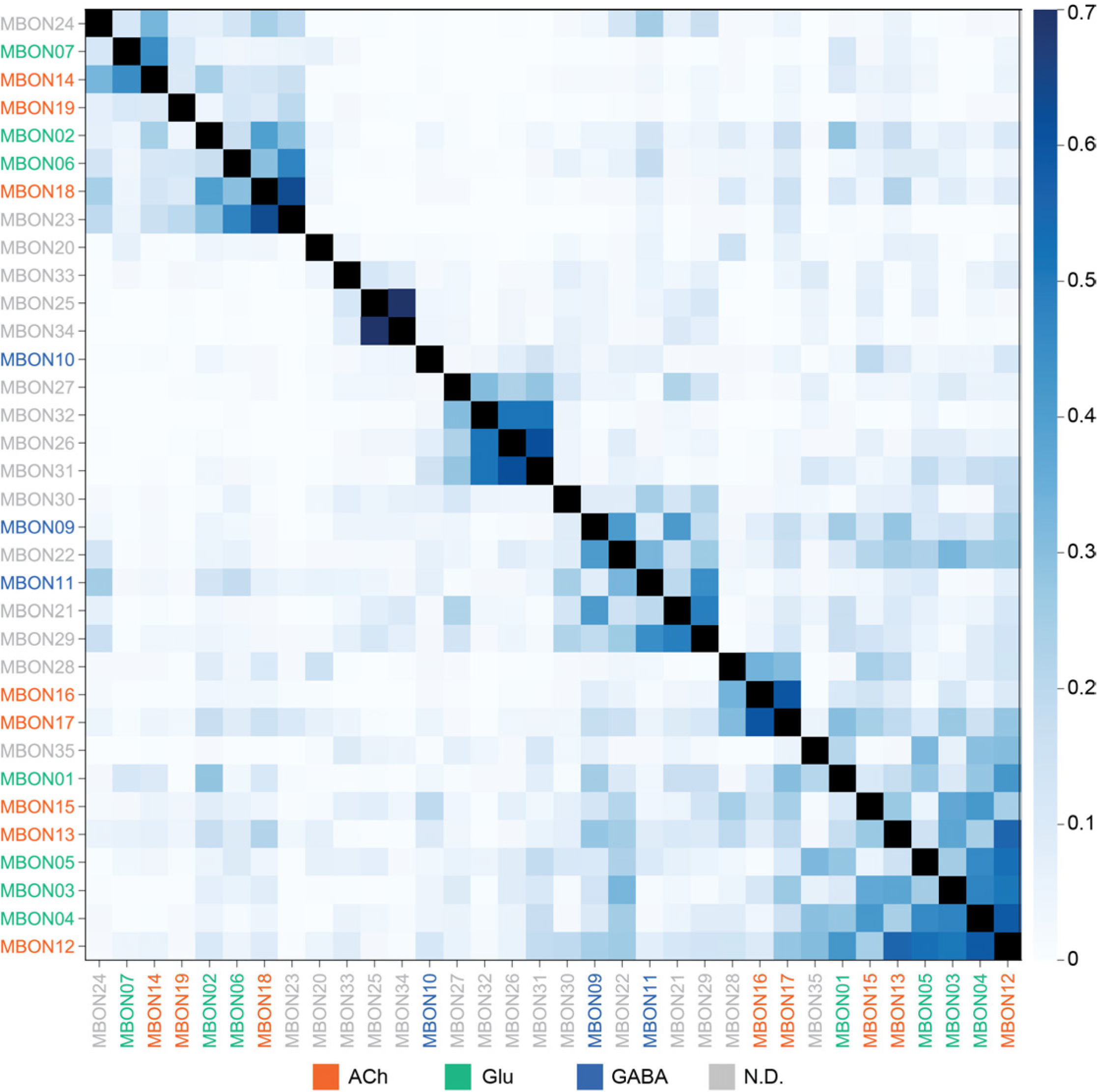
MBONs clustered by output similarity. Cosine similarity between each MBON type and their target population. The ordering of MBONs was determined using average clustering. Unlike Figure 16 or Figure 16-figure supplement 1, the ordering of the MBONs in this figure does not take into account input structure or anatomical structure, allowing for unbiased grouping of MBONs based on their outputs. This reveals some clusters of MBONs with convergent outputs that span multiple lobes. Only named neurons were considered in the target population. MBON names are color-coded to indicate their neurotransmitter.

**Figure 16-figure supplement 4.**
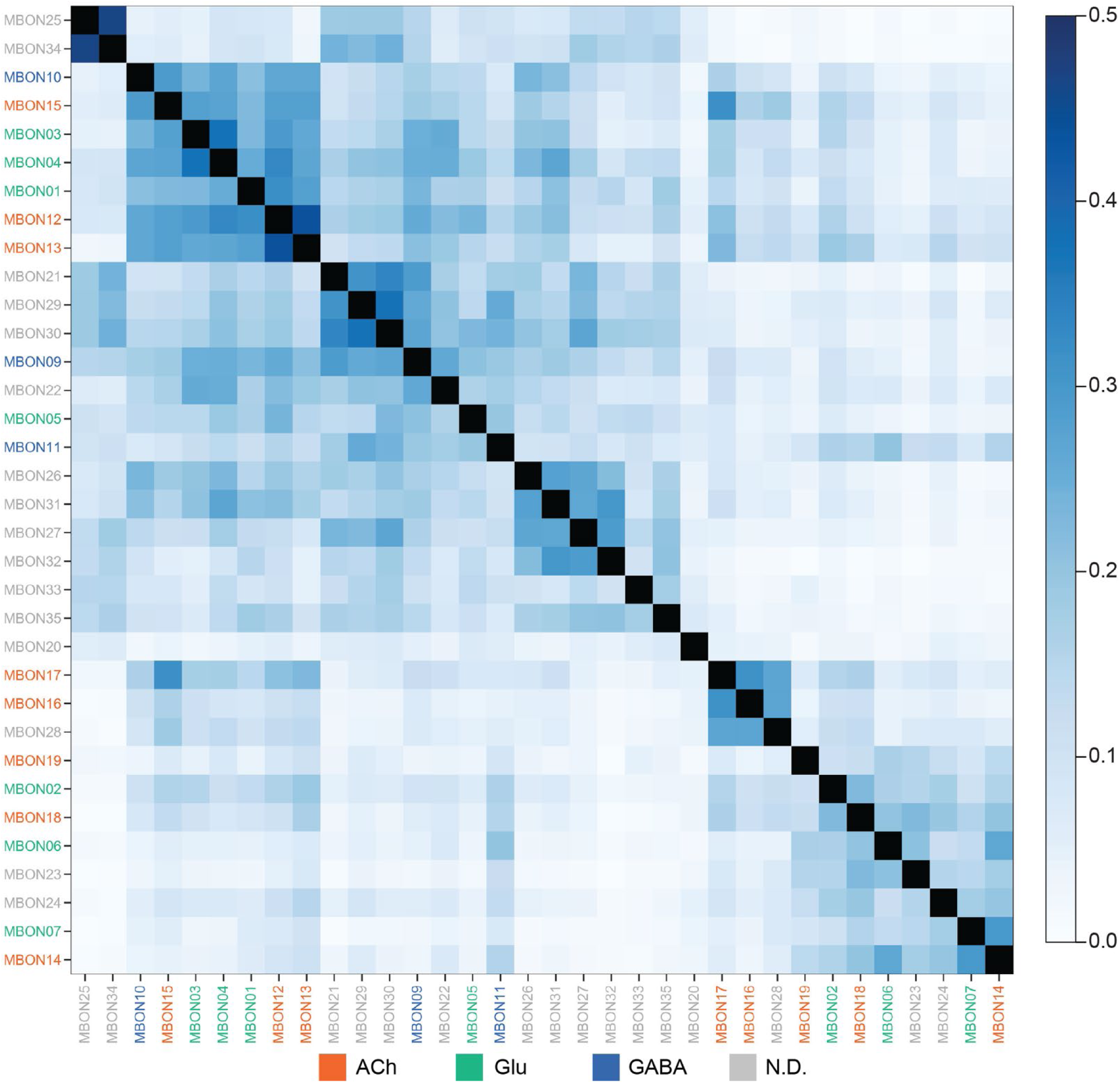
MBON morphologies grouped by output similarity. Radial dendrogram shows that MBONs with similar downstream connectivity innervate different compartments. A dendrogram based on the MBON cosine similarity matrix (Figure 16-figure supplement 3) was extracted and the morphologies of the MBON in each group were plotted. Colored dots match the colors of the MBONs in the morphology images. MBONs can have similar downstream connectivity regardless of whether they innervate the same (MBONs 16, 17 and 28) or different (MBONs 23, 18, 06 and 02) compartments. The asterisk indicates MBONs (MBON10, 13, 15, 20 and 33) that could not be grouped with another partner; they are singletons where their downstream connectivity is different from all other MBONs.

**Figure 17-figure supplement 1.**
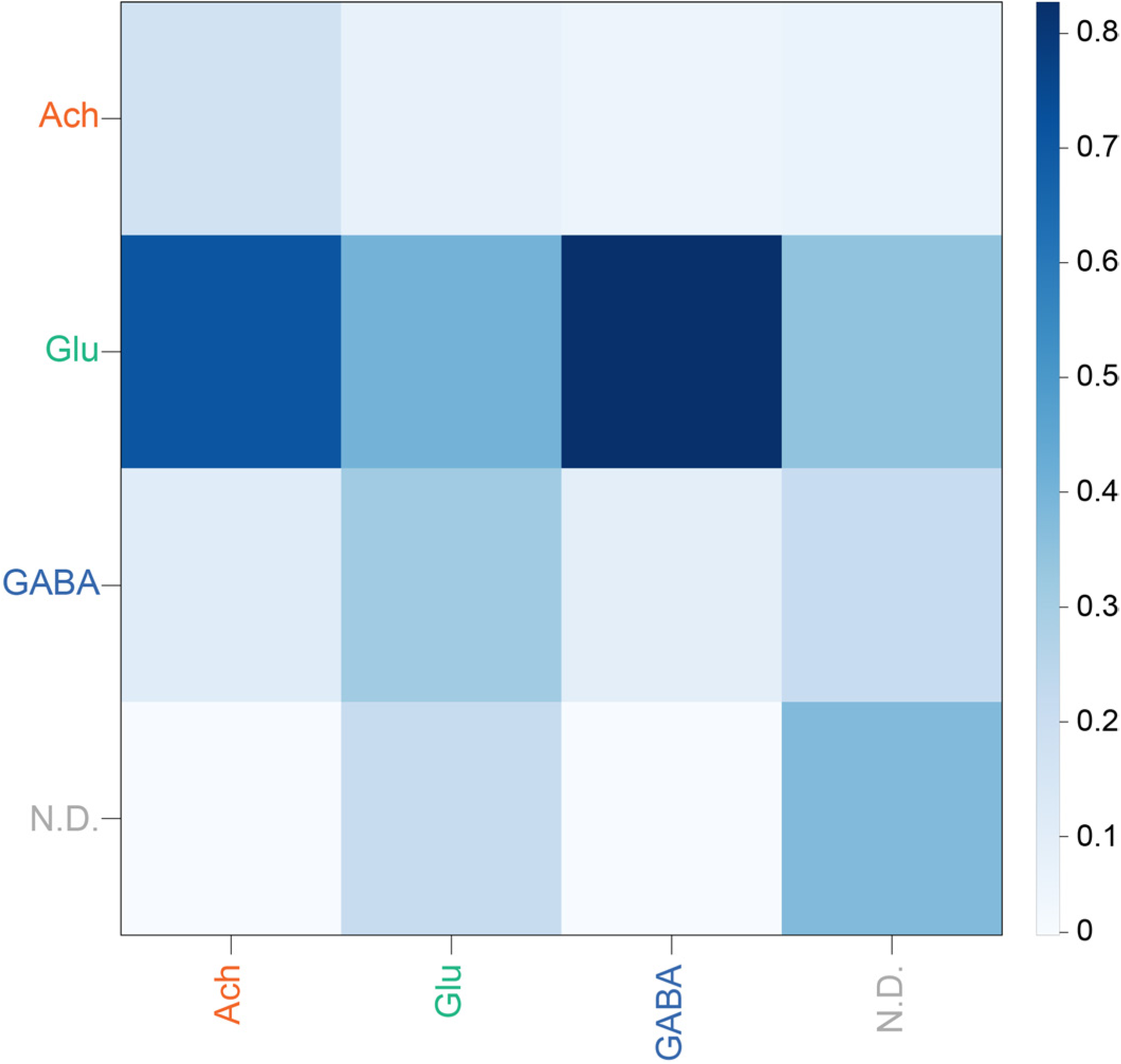
MBON output similarity pooled by neurotransmitter. Average cosine similarity of pairs of MBONs with the indicated neurotransmitters, computed by collapsing the blocks in Figure 17.

**Figure 18-figure supplement 1.**
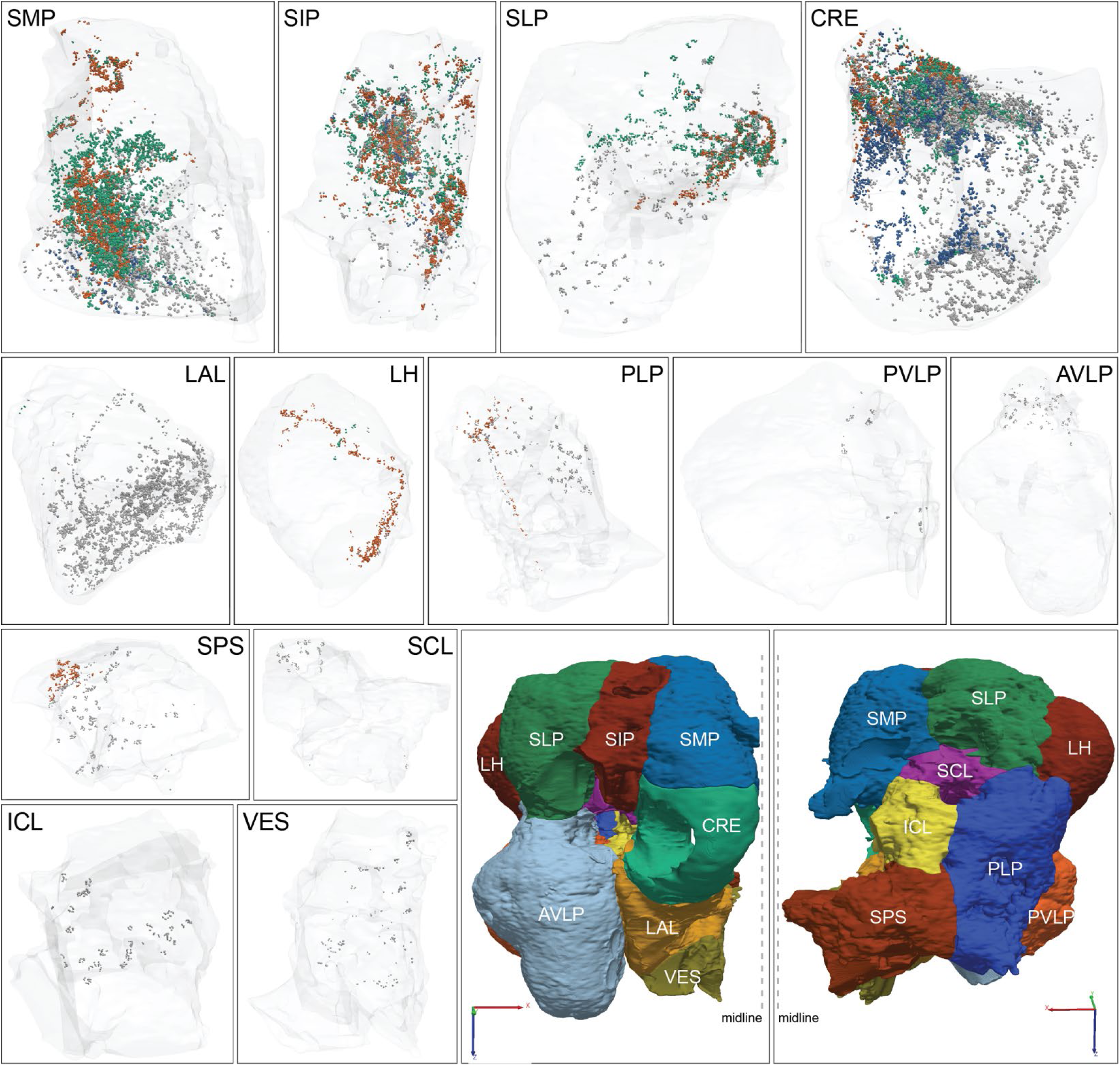
MBON output distribution by neurotransmitter presented on separated brain areas. Visualization of the spatial position of MBON output synapses within individual brain areas, as indicated.

**Figure 18-figure supplement 2.**
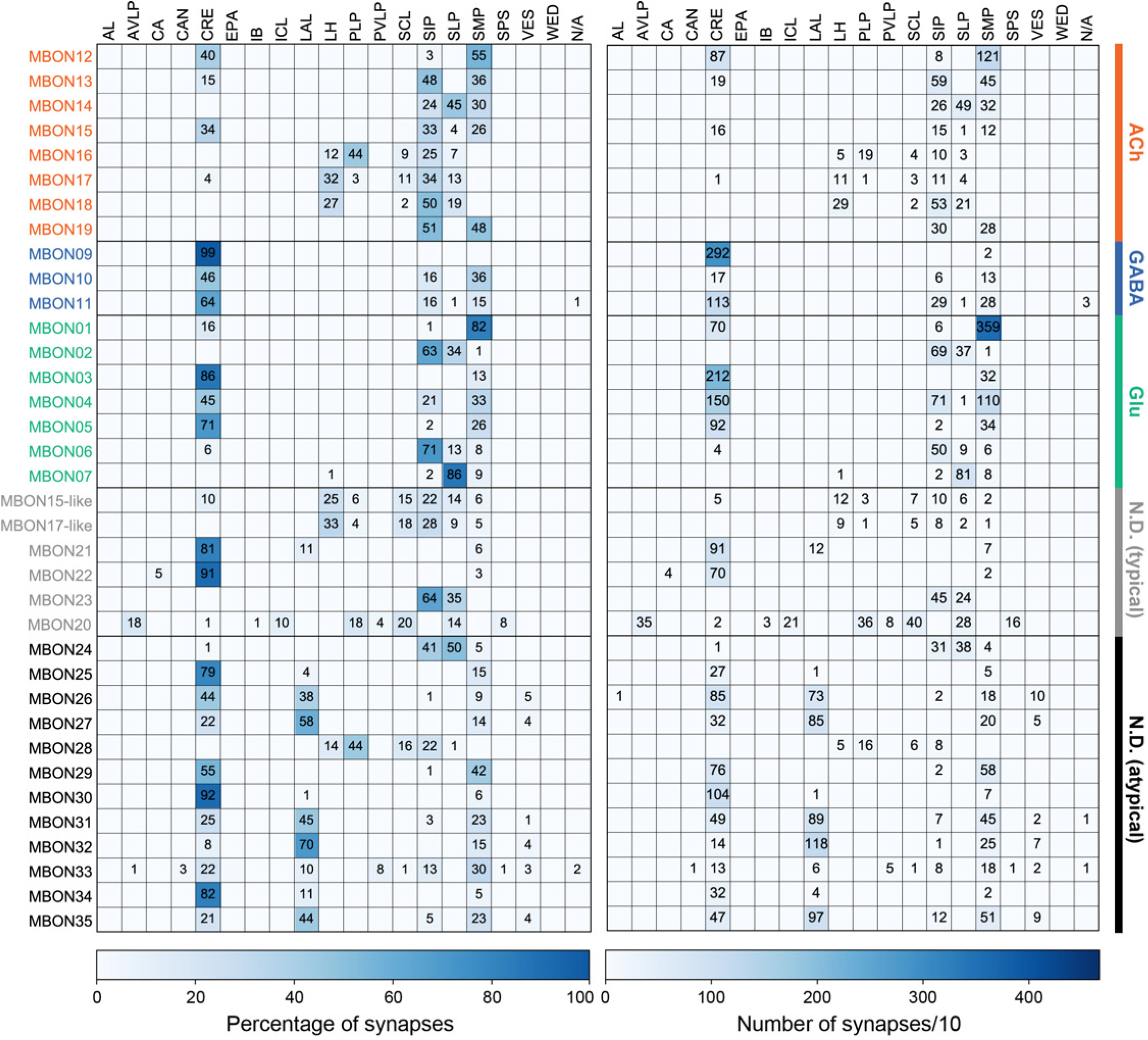
Distribution of individual MBON outputs by brain region. MBON outputs to brain areas, grouped by neurotransmitter; typical and atypical MBONs with undetermined transmitter were grouped separately. (Left) The value of each cell indicates the percentage of the given MBON’s output synapses that reside in the given brain area. Blank cells indicate values of less than 1%. (Right) The value of each cell indicates the number of the given MBON’s output synapses that reside in the given brain region divided by 10 and rounded to an integer. Blank cells indicate brain regions that receive fewer than 10 synapses from the corresponding MBON.

**Figure 19-figure supplement 1.**
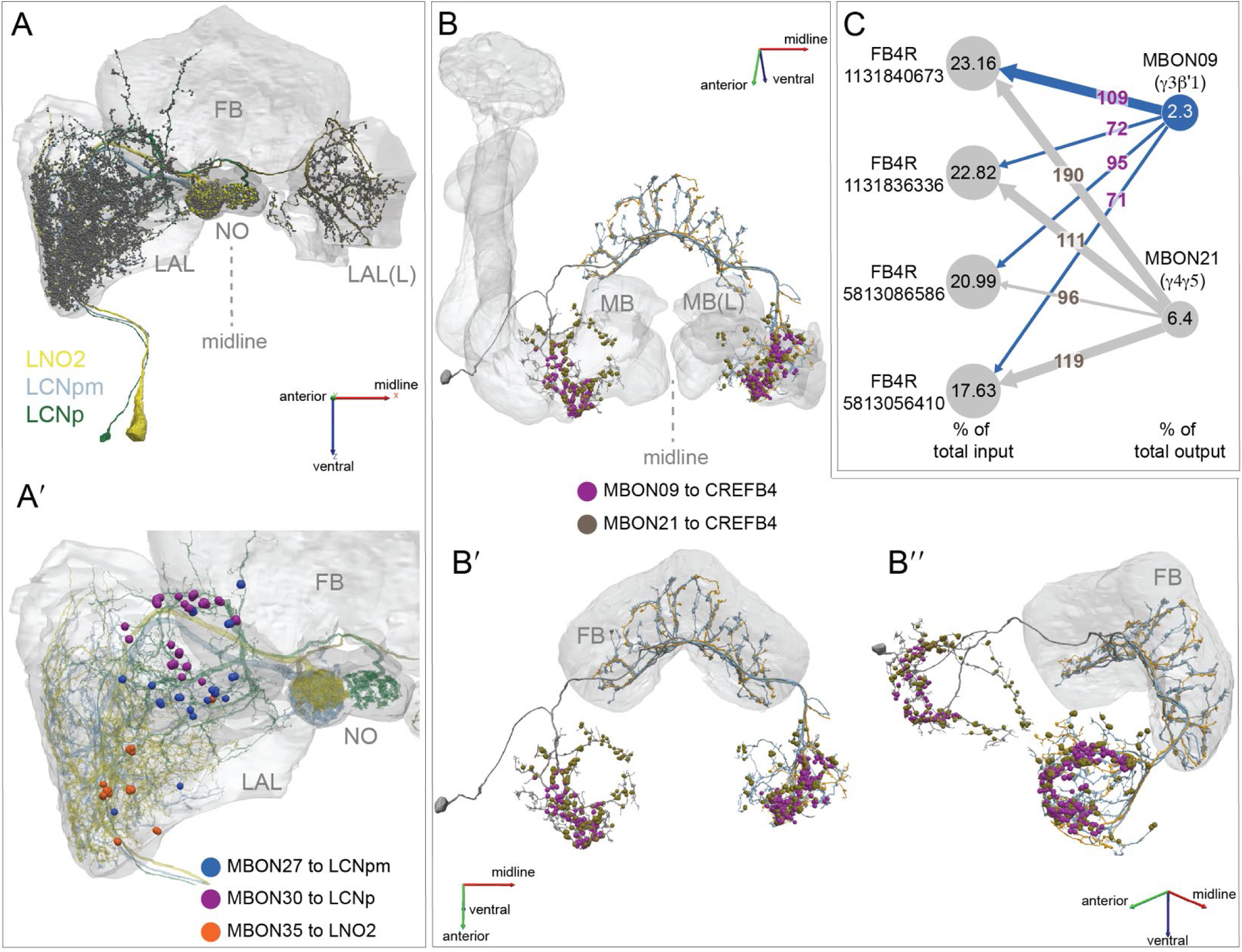
The morphology and connectivity of nodulus and CREFB4 neurons. (A) Morphologies of the LCNp, LCNpm, LNO2 nodulus neurons that are downstream of MBONs. The FB, NO and LAL are shown in grey. Presynaptic sites are shown as yellow dots and postsynaptic sites as grey dots. (A′) Higher magnification view, showing the locations of MBON inputs onto these neurons. (B) <u>CREFB4</u> (FB4R) neurons are shown with each of the four neurons of this type displayed in a different color. The locations of the synapses CREFB4 (FB4R) cells receive from MBON09 (γ3β′1) and MBON21 are shown, color-coded. (B′, B′′) Alternative views of image shown in (B). (C) Diagram of synaptic connectivity. The numbers in the circles that represent the cells indicate, for CREFB4 (FB4R) cells, the percentage of their input that comes from these two MBONs and, for the MBONs, the percent of their output that goes to CREFB4 (FB4R). The numbers in the arrows are the number of input synapses. The neurotransmitters used by MBON21 and CREFB4 have not been determined; MBON09 is GABAergic.

**Figure 20-figure supplement 1.**
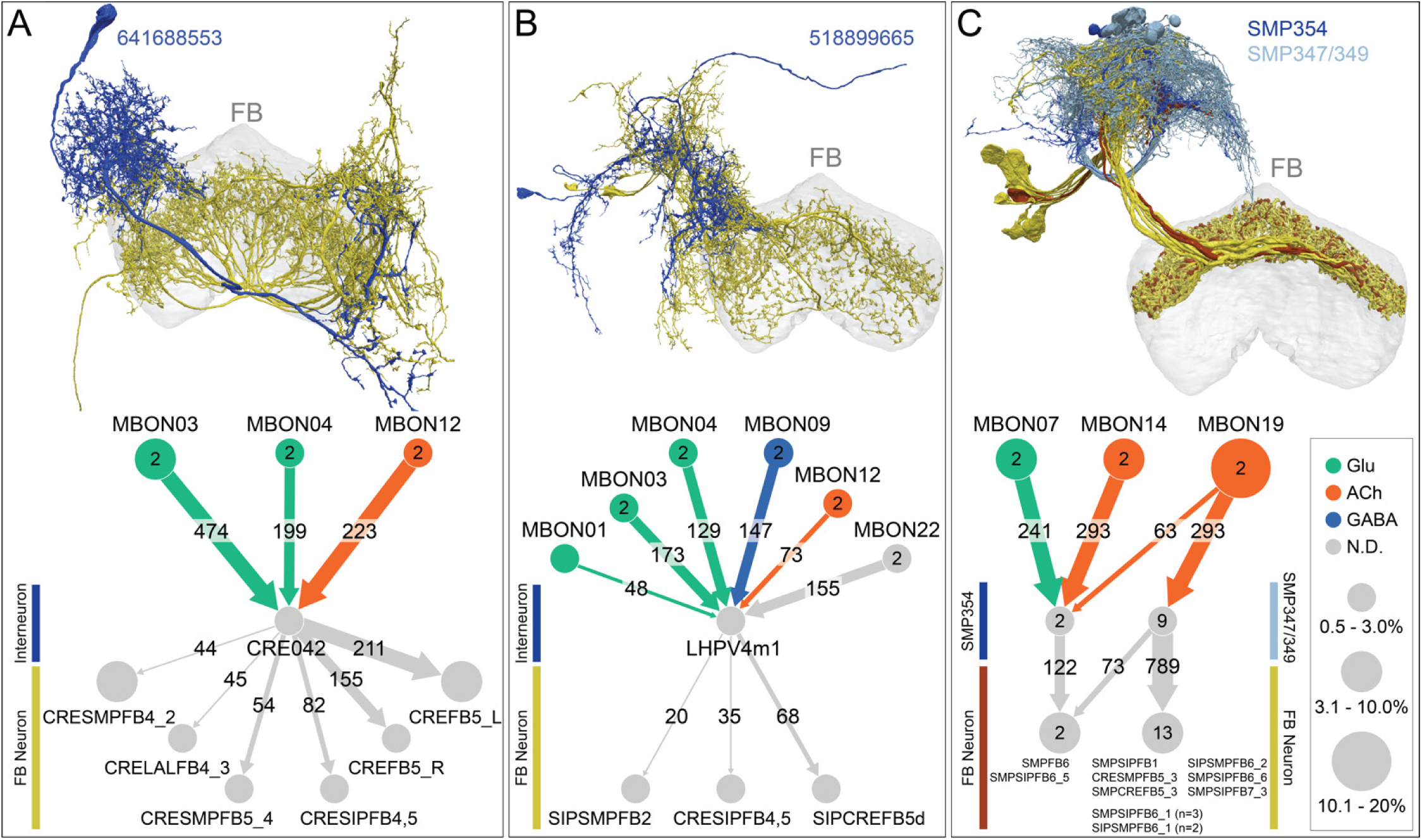
Examples of MBON to FB connectivity involving an interneuron. (A-C) Examples of connectivity patterns observed for connection of MBONs to the FB through an interneuron. At the top of each panel the morphologies of the interneuron (blue) and FB neurons (yellow/brown) are shown. At the bottom of each panel a connectivity diagram is shown, with neurotransmitters color-coded and the size of the circles for MBONs representing the percentage of their outputs that go to the shown interneurons. For FB neurons, the size of the circle represents the percentage of their inputs that come from those interneurons. Note that interneurons in this pathway are often targets of multiple MBONs with neurotransmitters of opposite sign, a so-called “push-pull” arrangement. (A) An interneuron, <u>CRE042</u>, that is strongly connected to MBONs of different valence inputs onto FB neurons of layers 4 and 5: <u>CRESMPFB4_2 (FB4D)</u>, <u>CRELALFB4_4 (FB4H)</u>, <u>CRESIPFB4,5 (FB4X)</u>, <u>CRESMPFB5 (FB5K)</u>, and <u>CRESMPFB5_4</u> <u>(FB5L)</u> (see Video 27). (B) An interneuron, <u>LHPV4m1</u>, that receives input from six MBON types and connects to three FB neurons of layers 2, 4 and 5, and 5: <u>SIPSMPFB2 (FB2F_a)</u>, <u>CRESIPFB4,5 (FB4X)</u>, and <u>SIPCREFB5d (FB5AB)</u>. No other interneuron connecting to the FB gets input from more different MBONs (see Video 28). (C) This interneuron can be divided into subtypes based on connectivity to α lobe MBONs: <u>SMP347</u>, <u>SMP349</u>, and <u>SMP354</u>, all of which are downstream targets of MBON19 (α2p3p), which conveys visual information (Figure 15A), but SMP354 consisting of two cells also gets strong input from MBON07 (α1) and MBON14 (α3) in a “push-pull” arrangement. All these interneurons convey information mostly to FB layer 6: <u>SMPSIPFB1,3 (FB1A)</u>, <u>CRESMPFB5_3 (FB5H)</u>, <u>SMPCREFB5_3 (FB5I)</u>, <u>SMPSIPFB6_1 (FB6A)</u>, <u>SIPSMPFB6_1 (FB6C_b)</u>, <u>SMPFB6 (FB6D)</u>, <u>SIPSMPFB6_2 (FB6E)</u>, <u>SMPSIPFB6_5 (FB6I)</u>, <u>SMPSIPFB6_6 (FB6K)</u>, and <u>SMPSIPFB7_3 (FB7F)</u> (see Video 29). Note these FB neurons contain 7 of the 9 cells in cluster 31 of Figure 34A.

**Figure 21-figure supplement 1.**
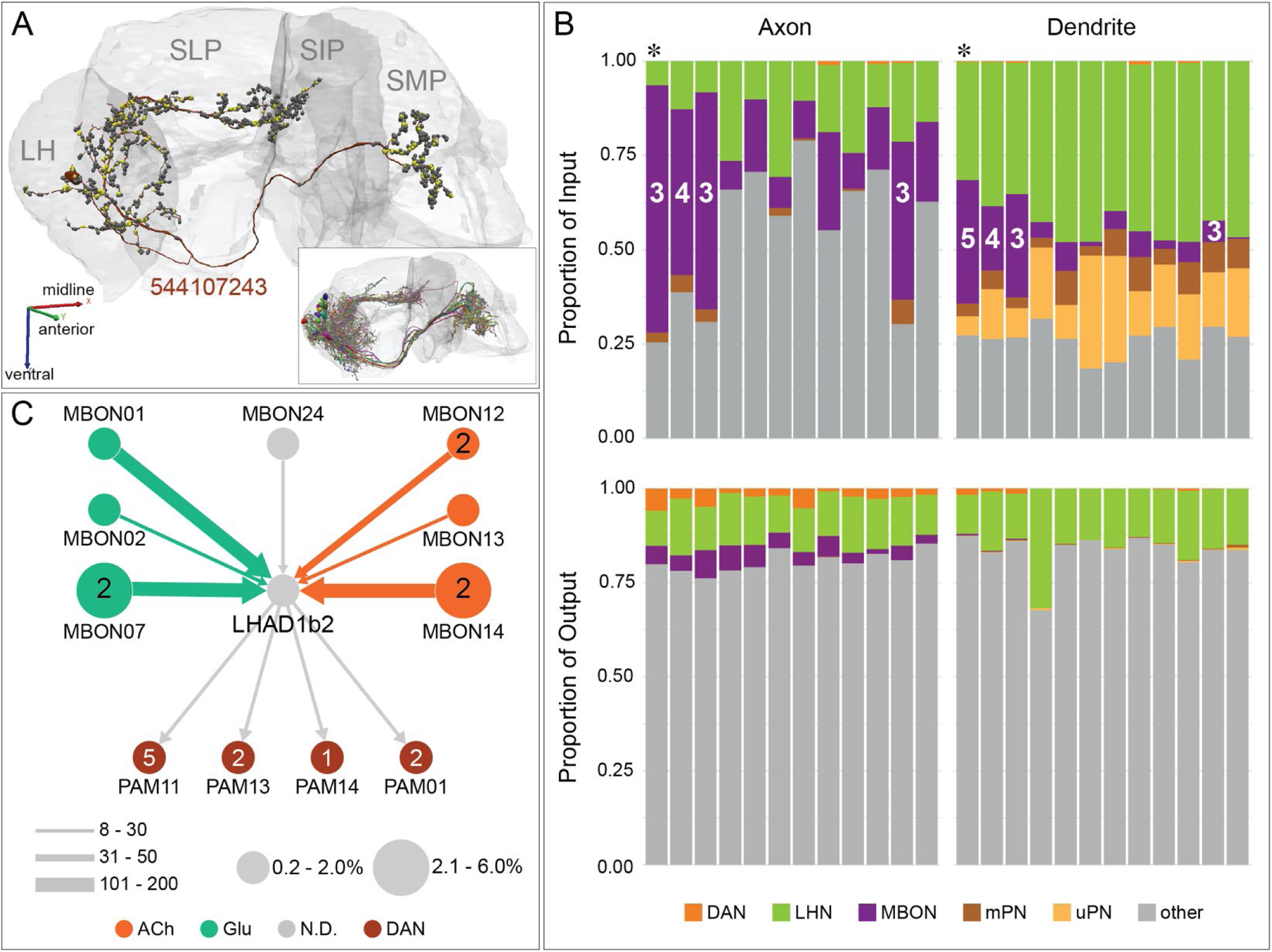
A LH neuron downstream of seven MBON types. The previously reported LH output neuronal type (LHAD1b2; Dolan et al., 2019) acts as a site of integration of innate and learned information. (A) The morphology of a <u>LHAD1b2_d</u> neuron (544107243) is shown in maroon, with its presynaptic and postsynaptic sites represented by yellow and grey dots, respectively. The inset shows the morphologies of all twelve neurons of this cell type: <u>LHAD1b2_a</u>: 673426956 and 544107335; <u>LHAD1b2_b:</u> 5813022459, 573683438 and 483017681; <u>LHAD1b2_c:</u> 5813052205, 543321179 and 574040939; <u>LHAD1b2_d:</u> 544107243 and 730562988; <u>LHAD1c1:</u> 729867599; and <u>LHAD1c2:</u> 673426956. (B) Histograms of the proportion of inputs to (top) and outputs from (bottom) each of the twelve LHAD1 neurons from various classes of cells are shown, color coded. These connections are divided into two groups based on the positions on the neurons’ arbors where they occur, labeled as Axon and Dendrite, based on the distribution of pre- and postsynaptic sites although all its arbors are mixed; the neuron order is the same in all histograms. The number of different MBON types innervating individual LHAD1 neurons varies; the number of MBON types providing input is indicated for those receiving input from three or more types. The neuron (544107243) shown in (A) and (C) is marked with an asterisk. (C) A connectivity diagram of the <u>LHAD1b2_d</u> neuron displayed in (A) is shown. This neuron receives input from seven MBON types and its downstream targets include four DAN cell types. The transmitters used by the MBONs are color-coded, the thickness of the arrows reflects synapse number, and the sizes of the circles that represent the MBONs reflects the percentage of their output directed to this LHAD1b2_d cell. The size of the circle representing LHAD1b2_d indicates the proportion of its total input that is derived from MBONs and the size of the circles representing the DANs indicate the fraction of their input provided by the LHAD1b2_d cell.

**Figure 21-figure supplement 2.**
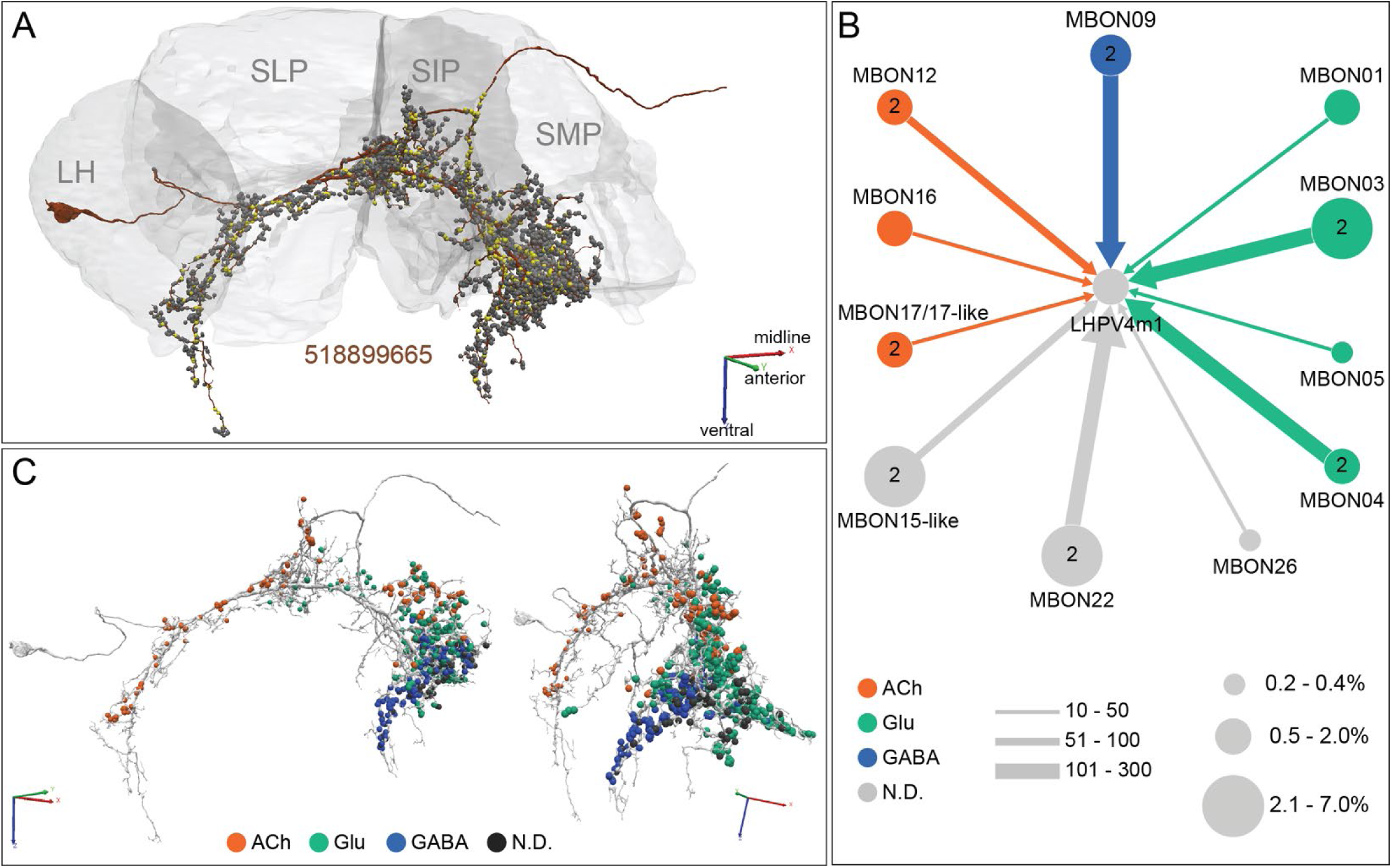
A neuron downstream of 11 MBON types. This neuron <u>LHPV4m1</u> (518899665) provides an extreme example of a neuron that integrates input from different MBON cell types. (A) The morphology of the neuron is shown in maroon, with its presynaptic and postsynaptic sites represented by yellow and grey dots, respectively. (B) Diagram showing the strengths of input from eleven different MBONs. The transmitters used by the MBONs are color-coded, the thickness of the arrows reflects synapse number, and the sizes of the circles that represent the MBONs reflect the percentage of their output directed to this neuron. (C) Distribution of sites of MBON input to the neuron, color-coded by MBON neurotransmitter type. Note that connections from MBONs with different neurotransmitters are segregated along the neuron’s different arbors; for example, cholinergic inputs are concentrated in its posterior branches.

**Figure 22-figure supplement 1.**
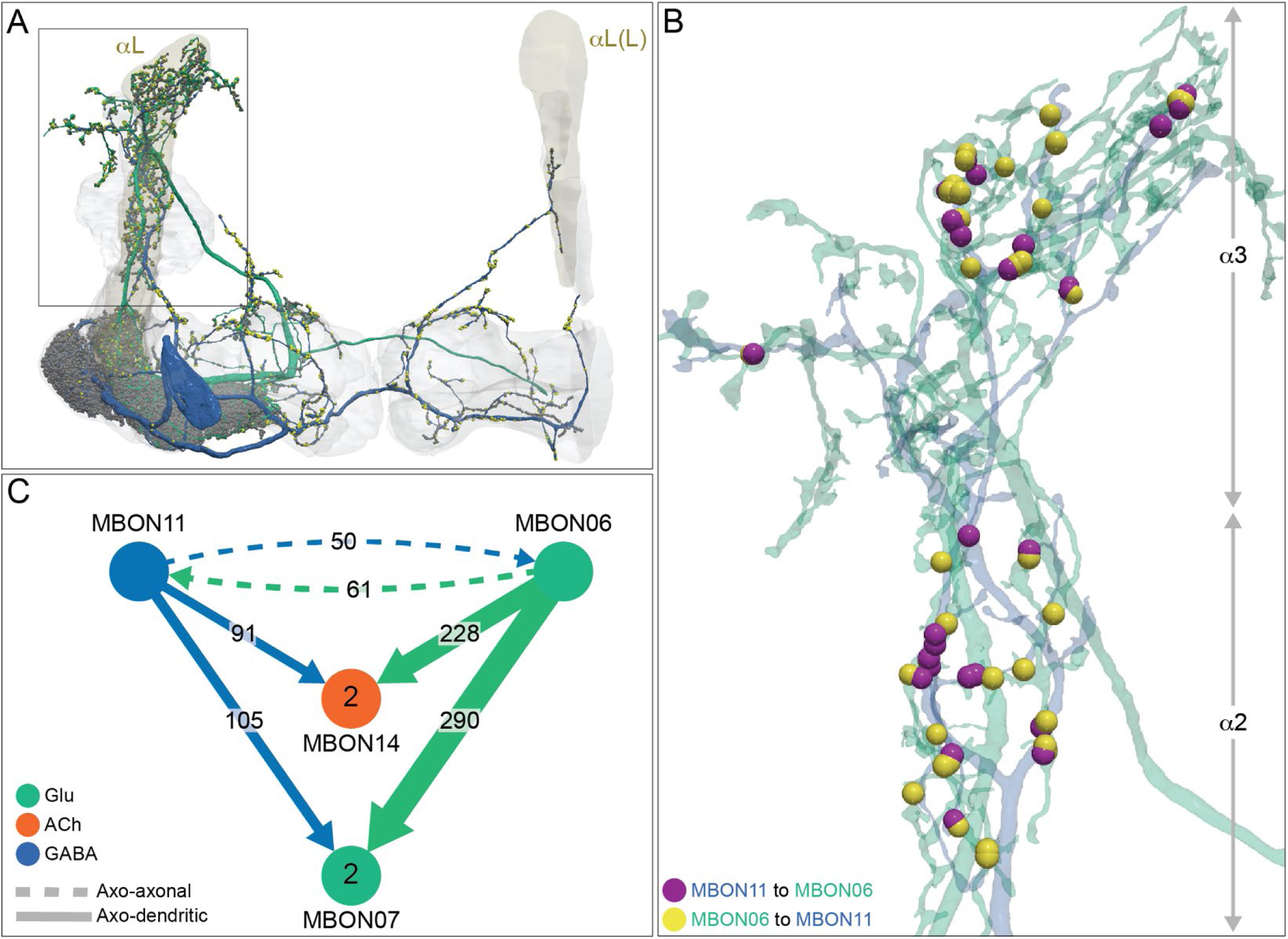
MBON06 and MBON11 make reciprocal axo-axonal connections in the α lobe. (A) MBON06 (β1>α; green) and MBON11 (γ1pedc>α/β; blue) send axonal processes to the α2, α3 and, to a lesser extent, α1 compartments. (B) MBON06 and MBON11 make reciprocal axo-axonal connections, mostly in the α2 and α3 compartments. The positions of these synapses are shown, color coded. (C) Diagram of a subset of connections involving MBON06 and MBON11, which both also make axo-dendritic connections onto MBON07 (α1) and MBON14 (α3) inside the MB lobes.

**Figure 23-figure supplement 1.**
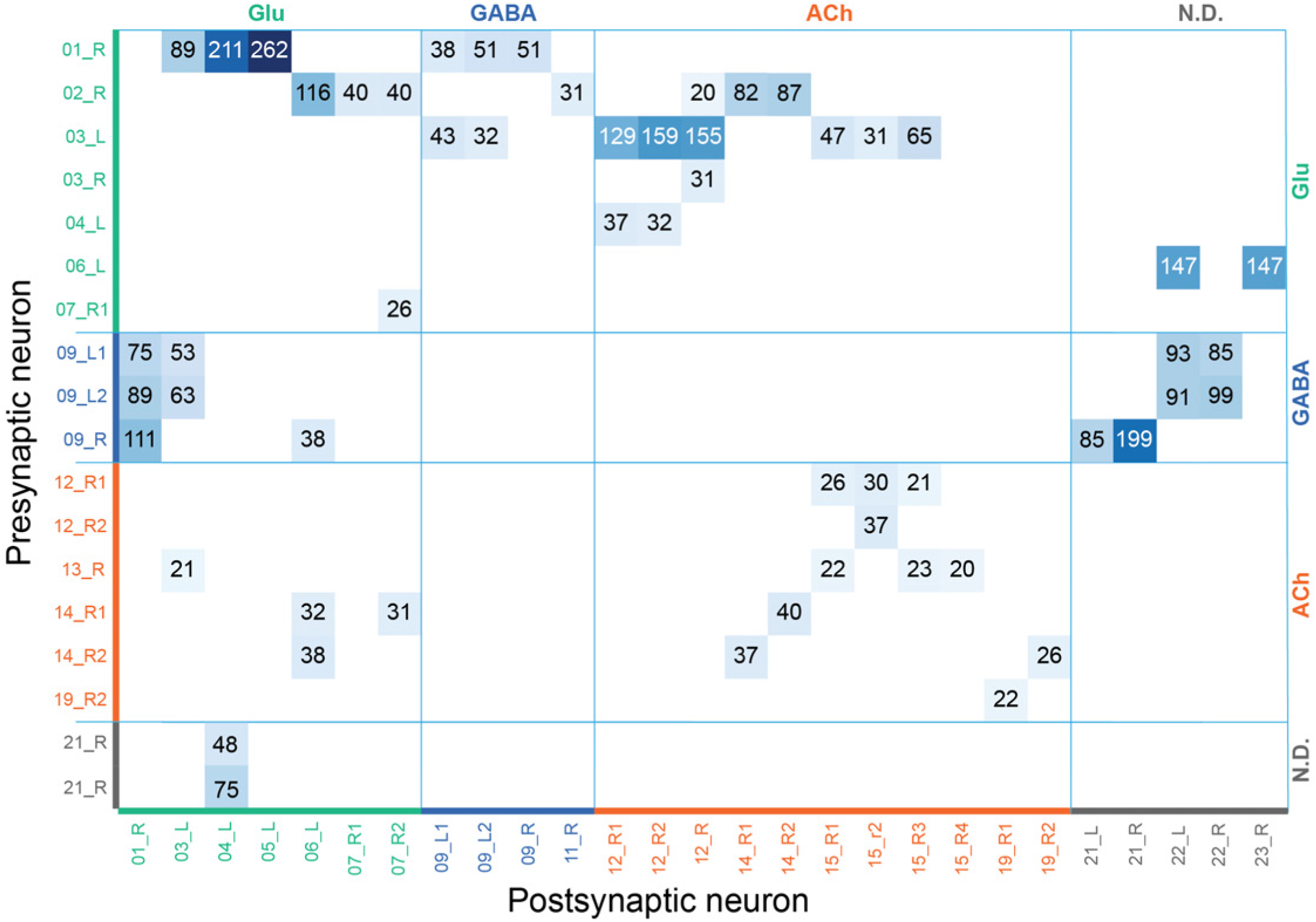
Axo-axonal connections among typical MBONs outside the MB presented as a connection matrix. This matrix shows all axo-axonal connections between pairs of typical MBONs that involve 20 or more synapses. The MBONs on the vertical axis are presynaptic to those of the horizontal axis. MBON types are sorted by neurotransmitter. Blank rows or columns are not shown. Connections are seen more frequently and are stronger between glutamatergic MBONs onto cholinergic MBONs or other glutamatergic MBONs. Likewise, GABAergic MBONs frequently synapse onto the axons of glutamatergic MBONs and MBONs with undetermined transmitter. In contrast, no axo-axonal synapses between GABAergic MBONs and cholinergic MBONs were seen in either direction. In several cases, axo-axonal synapses were highly localized and concentrated in a single axonal branch (see Figure 23-figure supplement 2).

**Figure 23-figure supplement 2.**
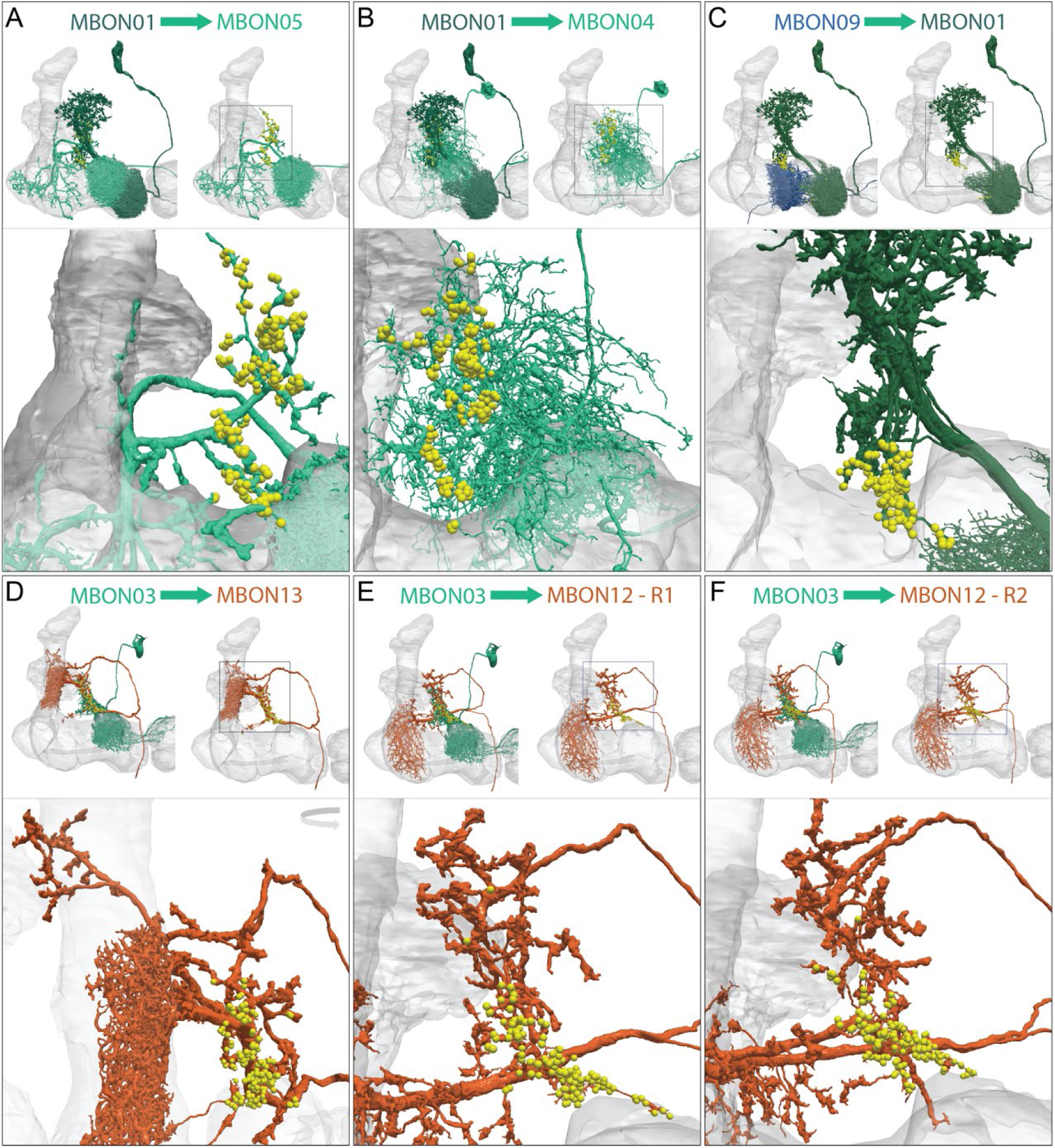
Axo-axonal synapses between MBONs can be highly localized. (A-F) Each panel shows three images: a low magnification view (upper left) of two neurons that make axo-axonal connections, a low magnification view (upper right) of just the postsynaptic neuron, and a higher resolution view of the relevant portion of its axonal arbor (lower). Neurons are color coded by neurotransmitter type. Synapses are depicted as yellow dots. (A-B) MBON01 (γ5β’2a; dark green) forms axo-axonal connections onto MBON04 (β′2mp_bilateral) and MBON05 (γ4>γ1γ2) with similar synapse counts, 211 and 262 respectively. (A) Synapses from MBON01 to MBON05 are widely distributed throughout MBON05’s axonal terminal. (B) Synapses from MBON01 to MBON04 are highly biased towards the lateral and anterior of MBON04’s axonal terminal. (C) MBON01 receives over 100 synapses from MBON09 (γ3β′1)_R onto its axon. Most of these connections are localized in a smaller branch that is ventral and isolated from its main axonal arbor. (D) Localized axo-axonal connections from MBON03 (β′2mp) to MBON13 (α′2). Note that the enlarged image has been rotated to improve visibility of the relevant arbor. (E-F) Localized axo-axonal connections between MBON03 and each of two MBON12 (γ2α′1) cells occur in the same location on the axonal arbors of <u>MBON12-R1</u> (E) and <u>MBON12-R2</u> (F).

**Figure 23-figure supplement 3.**
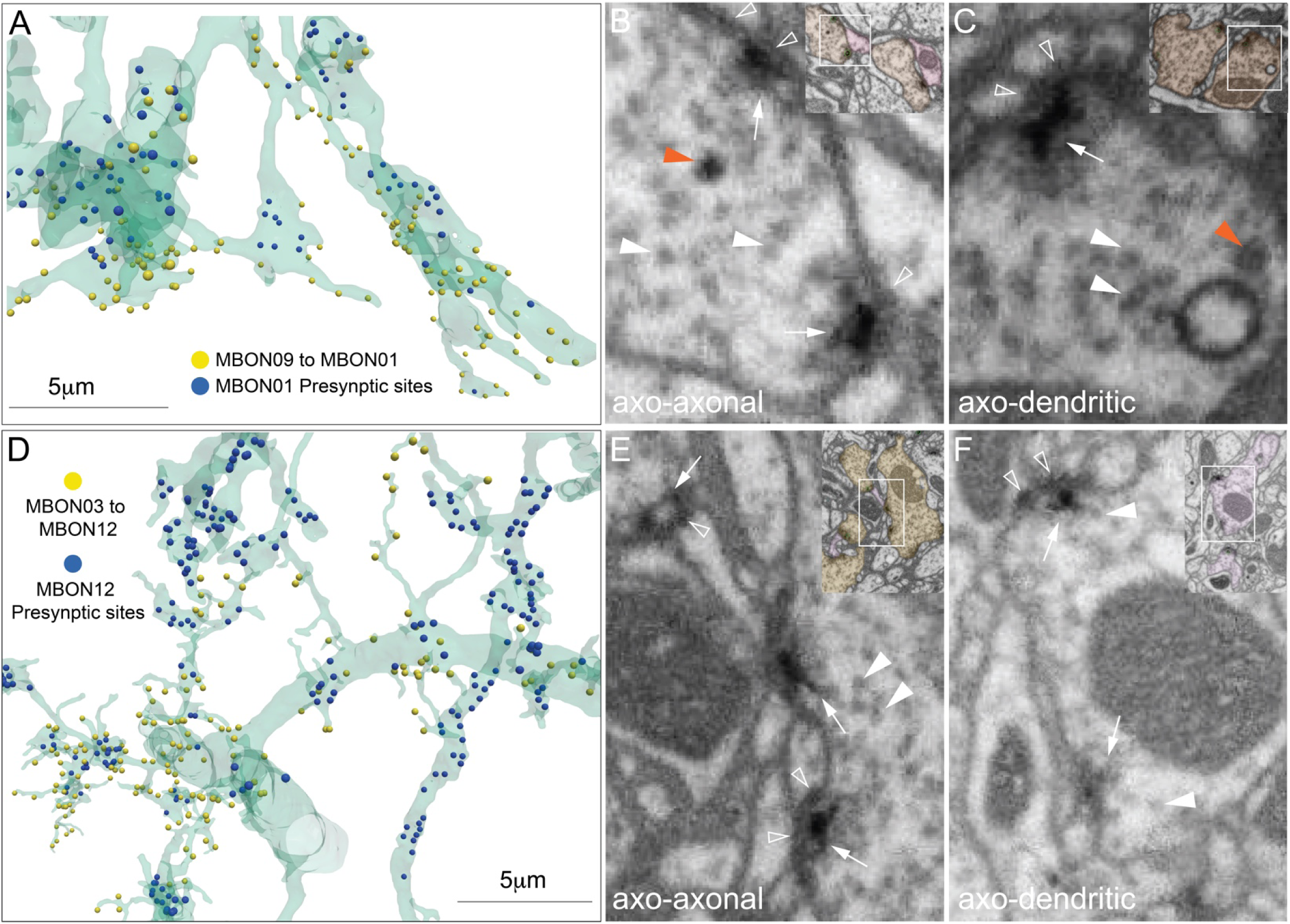
Morphology of axo-axonal synapses. (A) A portion of the MBON01 (γ5β’2a) arbor from within the enlarged area shown in Figure 23-figure supplement 2C is shown in transparent green. The relative locations of axo-axonal synapses from <u>MBON09</u> (γ3β′1) onto MBON01 (yellow dots) and MBON01’s output synapses to its downstream targets (blue dots) are indicated. (B) Electron micrograph from the hemibrain data set showing two MBON09 to MBON01 axo-axonal synapses (white arrows). Two types of vesicles are indicated by the arrowhead (small clear vesicle, white arrowhead; larger dense core vesicle, orange arrowhead). Postsynaptic densities are indicated by the hollow white arrowheads. The inset in the upper right shows a lower magnification view; MBON09 is colored brown, MBON01 is pink and annotated presynaptic sites are marked by the small green circles. (C) An axo-dendritic synapse from MBON09 to one of its downstream targets is shown for comparison. The morphologies of MBON’s axo-axonal and axo-dendritic synapses are similar. (D) A portion of the MBON12 (γ2α′1) arbor from within the enlarged area shown in Figure 23-figure supplement 2F is shown in transparent green. The relative locations of axo-axonal synapses from MBON03 (β′2mp) onto <u>MBON12</u> (yellow dots) and MBON12’s output synapses to its downstream targets (blue dots) are indicated. (E) Electron micrograph from the hemibrain data set showing three MBON03 to MBON012 axo-axonal synapses (white arrows). Small clear vesicles are indicated by the white arrowheads. Postsynaptic densities are indicated by the hollow white arrowheads. The inset in the upper right shows a lower magnification view; MBON03 is shaded brown, MBON12 is pink. (F) An axo-dendritic synapse from MBON12 to one of its downstream targets is shown for comparison. The inset in the upper right shows a lower magnification view; MBON12 is shaded pink. The morphologies of MBON’s axo-axonal and axo-dendritic synapses are similar.

**Figure 24-figure supplement 1.**
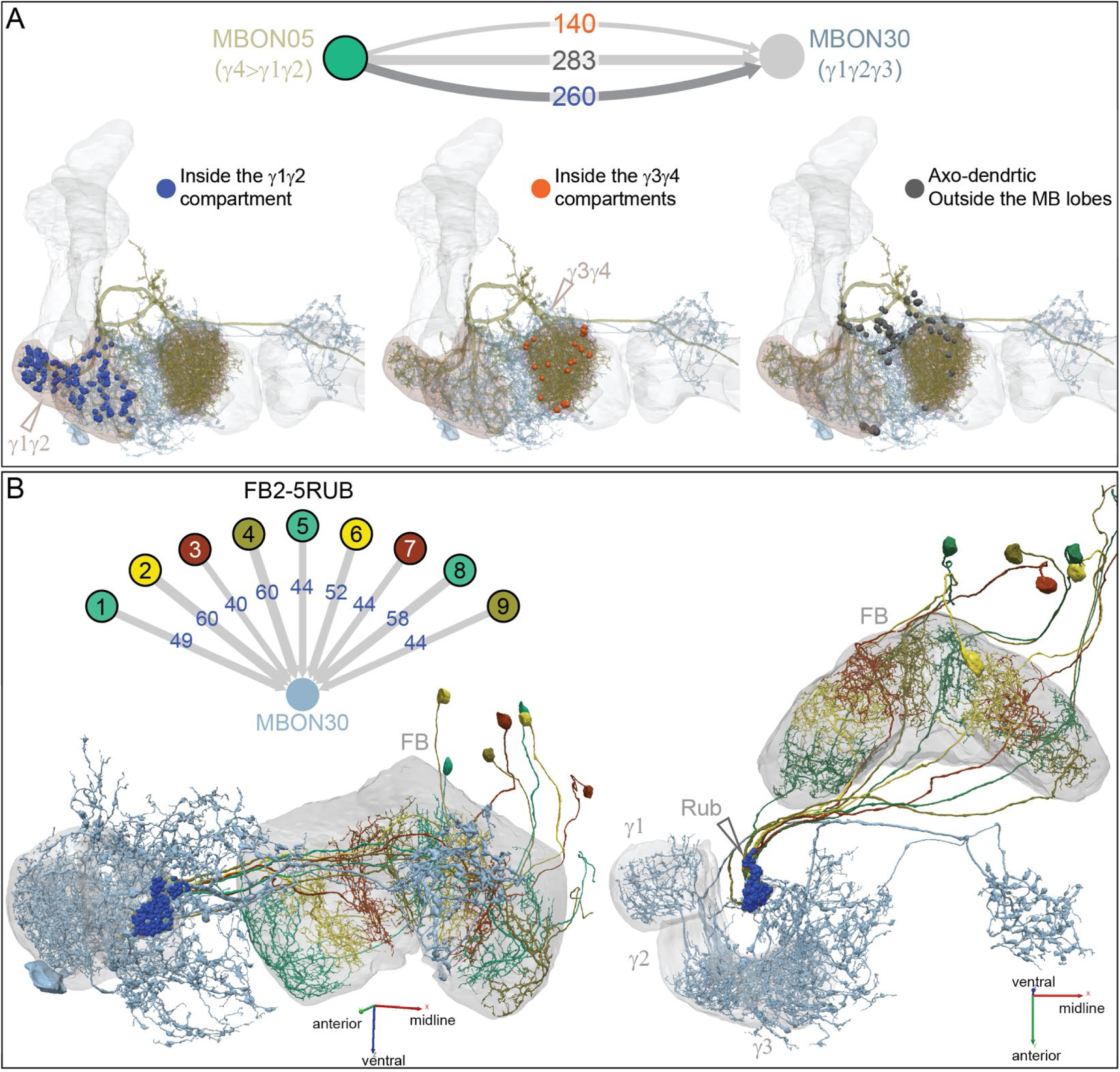
MBON30 displays an unusual pattern of innervation from other MBONs and FB neurons. (A) MBON30 receives strong input from MBON05, a feedforward glutamatergic MBON, through three distinct axo-dendritic paths: inside the γ1 and γ2 compartments (260 synapses); inside the γ3 and γ4 compartments (140 synapses); and outside the MB lobes (283 synapses). The locations of synapses used in each of these three connections are shown. Note that in the connectivity diagram in Figure 24, the arrow represents all 400 synapses that occur in either the γ1 and γ2 compartments (260) or in the γ3 and γ4 compartments (140). (B) MBON30 receives strong input from the FB mediated by the FB2-5RUB cell type which has nine columnar cells. The connection occurs inside the RUB. MBON30 is the only MBON that has input from an FB cell type among its top 10 inputs.

**Figure 25-figure supplement 1.**
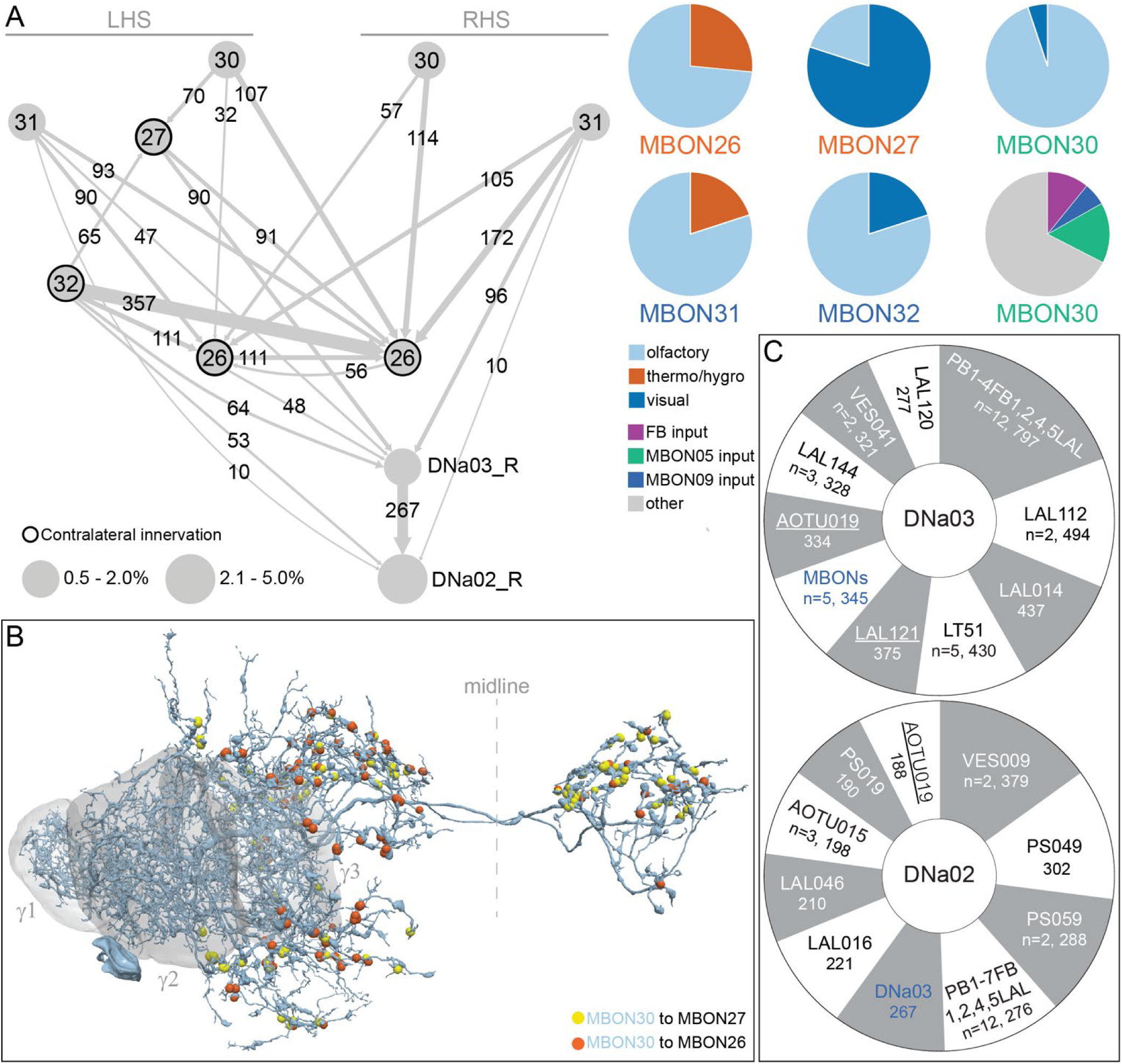
Top inputs to DNa02 and DNa03 (A) Left: Expanded connectivity diagram now including MBON30. MBON30 provides strong input to MBON26 and MBON27 (Figure 24) and conveys information from the FB (Figure 24-figure supplement 1; Video 15). Right: Pie charts describing inputs to MBON26, MBON27, MBON30, MBON31 and MBON32. The first five pie charts show the modalities of sensory information conveyed to these MBONs by KCs (data taken from Figure 15-supplement 2). Note MBON26 and MBON31 receive substantial thermo/hygrosensory input and MBON 27 and MBON32 receive substantial visual input. MBON30 receives strong input from MBON05 and MBON09, as well as from a single FB columnar neuron cell type (FB2-5RUB); these three cell types provide 36 percent of MBON30’s input, when input from KCs, DANs, APL and DPM is excluded. (B) The presynaptic sites on MBON30 that connect to MBON26 and MBON27 are shown, color coded. (C) Sunburst charts showing the ten cell types providing the largest amount of input, based on synapse number, to DNa03 (top) and DNa02 (bottom). The number of synapses and the number of cells in the cell type if more than one (n=) are shown. Neuron names with underscore indicate they innervate the contralateral side.

**Figure 25-figure supplement 2.**
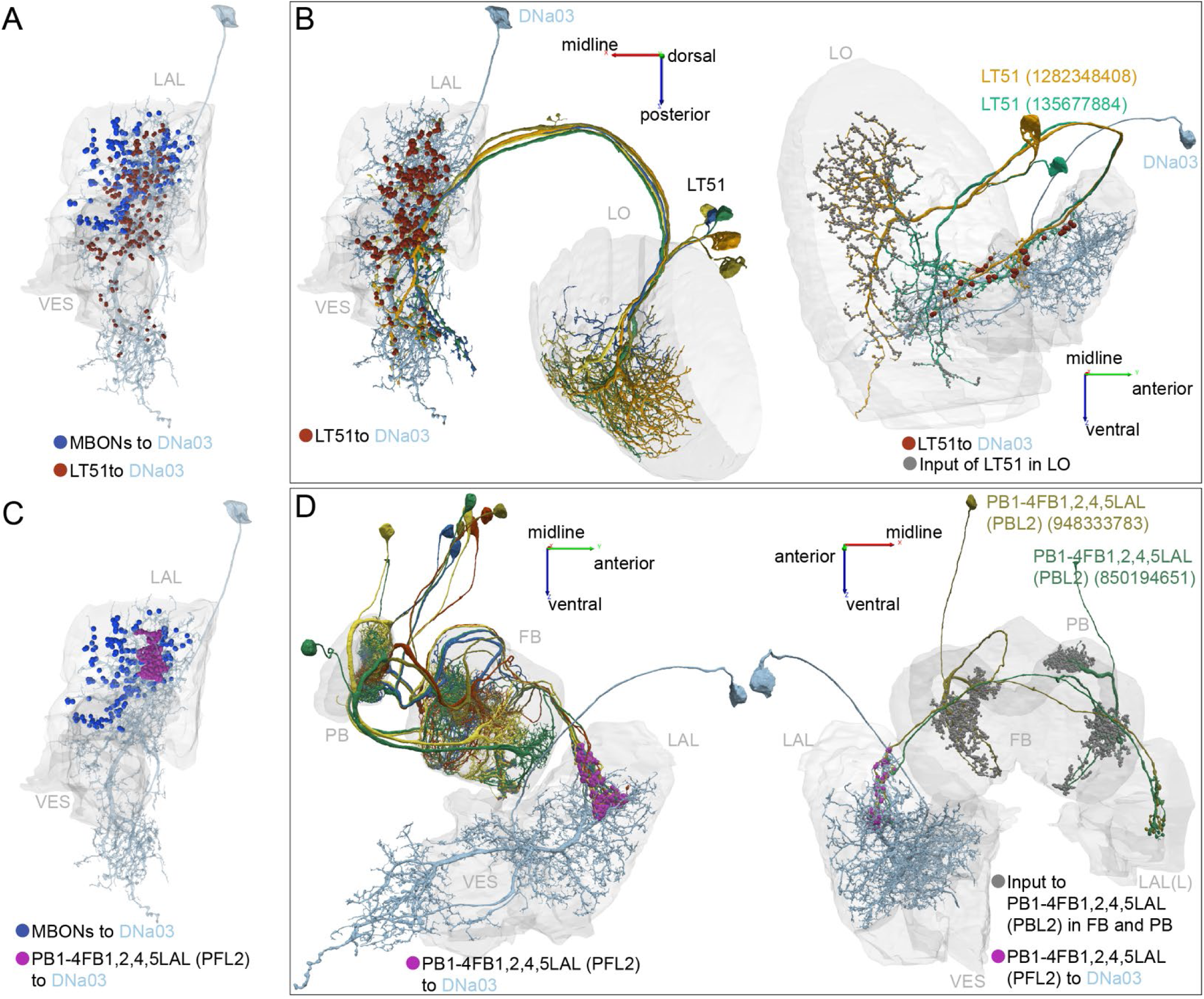
Central complex and LO inputs to DNa03. (A) DNa03 receives strong visual input from lobula tangential (LT) neurons, LT51 (also see Figure 25-figure supplement 1 C). The <u>five LT51</u> neurons make 430 synapses onto DNa03, shown as reddish brown dots. These connections are located at the core of the DNa03 dendrite while the MBONs synapse onto DNa03 around the periphery (shown as blue dots). The five LT51s that synapse onto DNa03 represent only a subset of the twelve LT51 identified in the right optic lobe and the two of these cells account for 362 of the 430 synapses from LT51 cells to DNa03. Interestingly, these five connected LT51 cells have their dendrites in the same half of the lobula, corresponding to the frontal part of the visual field. (B) Left: a dorsal view of LT51 connecting to DNa03. Right: two LT51 neurons connecting to DNa03 are shown with their postsynaptic sites in the LO indicated by the grey dots and presynaptic connections to DNa03 as reddish brown dots. (C) The strongest input cell type to DNa03 is a cell type with dendrites in the PB and the FB, <u>PB1-4FB1,2,4,5LAL(PFL2),</u> with 12 such cells making a total of ∼800 synapses (also see Figure 25-figure supplement 1 C). These synaptic connections are highly concentrated on a small dendritic area of DNa03, shown in purple with MBON input on the periphery (blue). (D) Left: Side view of the connections between the PFL2 neurons and DNa03. Right: Two of the PFL2 neurons are shown with their output to DNa03 (purple) and their input sites in FB and PB (grey). Note that PFL2 neurons are highly polarized.

**Figure 26-figure supplement 1.**
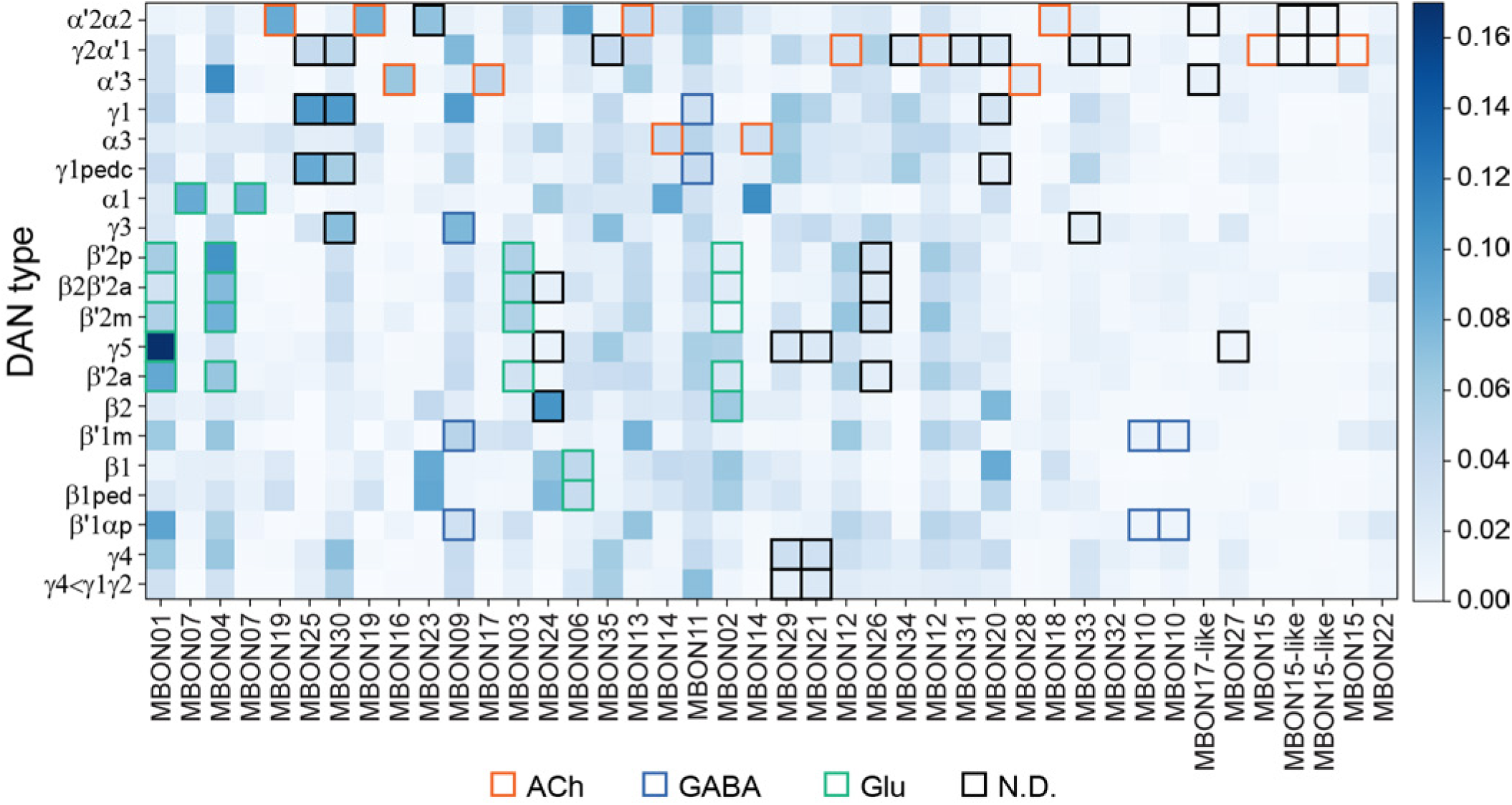
MBON to DAN feedback mediated by an interneuron. MBONs also make indirect connections to DANs in which the MBON connects to an interneuron which then connects to a DAN. This matrix shows the effective connection strength of such MBON-to-DAN feedback loops and is meant to complement the data presented in Figure 26, which describes direct connections between MBONs and DANs. Effective connection strength is a measure that takes into account both the strength of the MBON’s connection to the interneuron and that of the interneuron to the DAN. Effective connection strength is computed by multiplying the matrix of DAN inputs with the matrix of MBON outputs (both matrices are normalized so that the inputs to each neuron sum to 1). This connection strength is averaged across all DANs of the indicated type. Highlighted boxes indicate interactions between MBONs and DANs that share a compartment, and the box color indicates the neurotransmitter of the MBON. MBONs are ordered by their proportion of self-feedback, as defined and shown in Figure 26-figure supplement 2.

**Figure 26-figure supplement 2.**
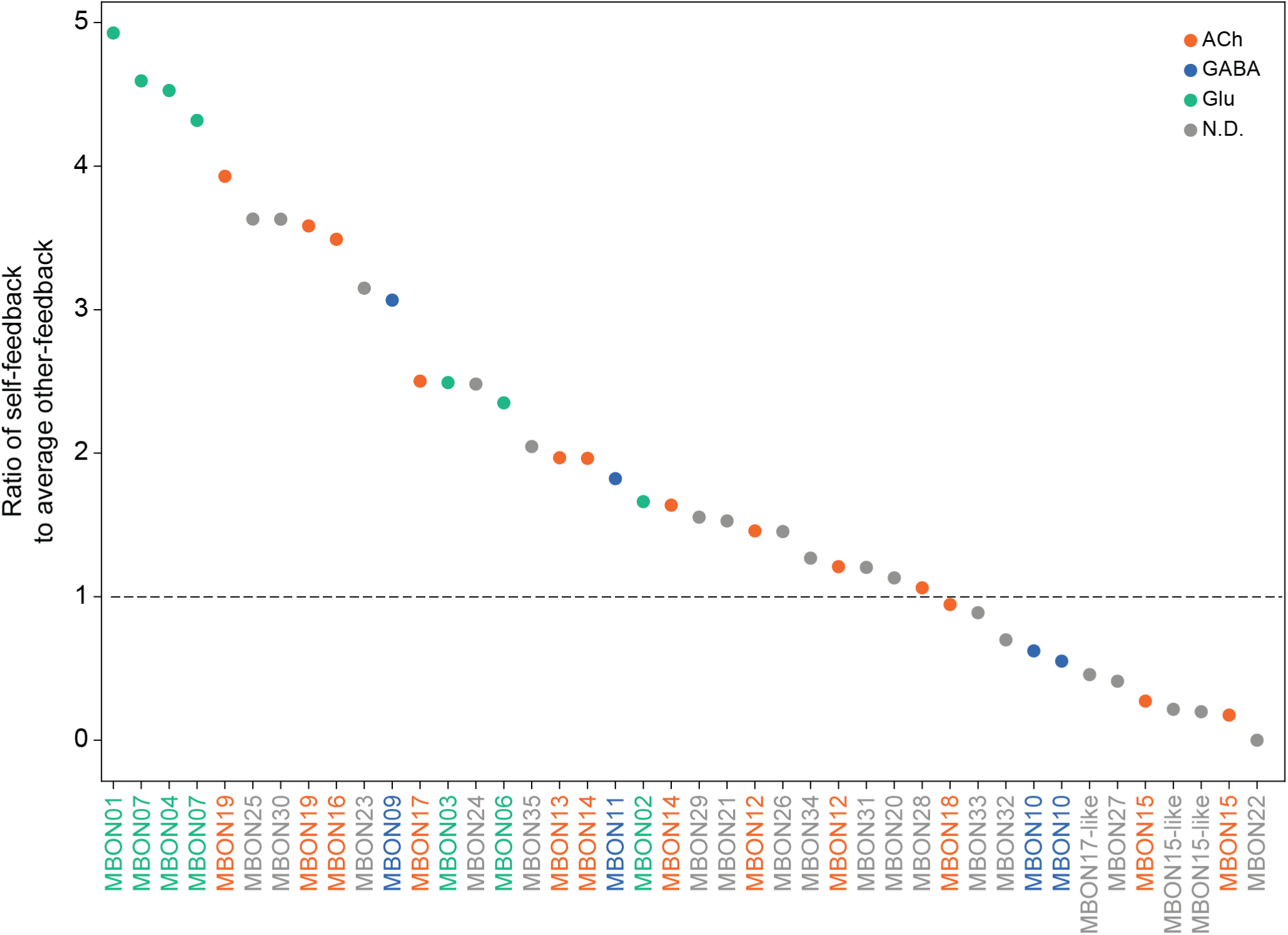
Distribution of DAN feedback mediated by an interneuron. This plot shows the ratio of feedback to the same compartment (self-feedback) to feedback to other compartments (other-feedback) for each MBON type; color-coding indicates the neurotransmitter used by the MBON. The plot reveals a robust pattern of glutamatergic (and to a lesser extent, cholinergic) MBONs indirectly modulating their associated DANs. The plot is meant to complement the data presented in Figure 26, which describes direct connections between MBONs and DANs. The strength of indirect self-feedback is defined as the average effective connection strength mediated by one interneuron between a given MBON and the DANs that innervate the same compartment. The strength of average other-feedback is defined as the average effective connection strength between the MBON and all other DANs, again mediated by one interneuron. The value plotted on the y-axis is the ratio of these two quantities. That is, MBONs with higher ratios have a larger fraction of the interneuron-mediated feedback they initiate directed back to their own compartment, rather than to other compartments.

**Figure 27-figure supplement 1.**
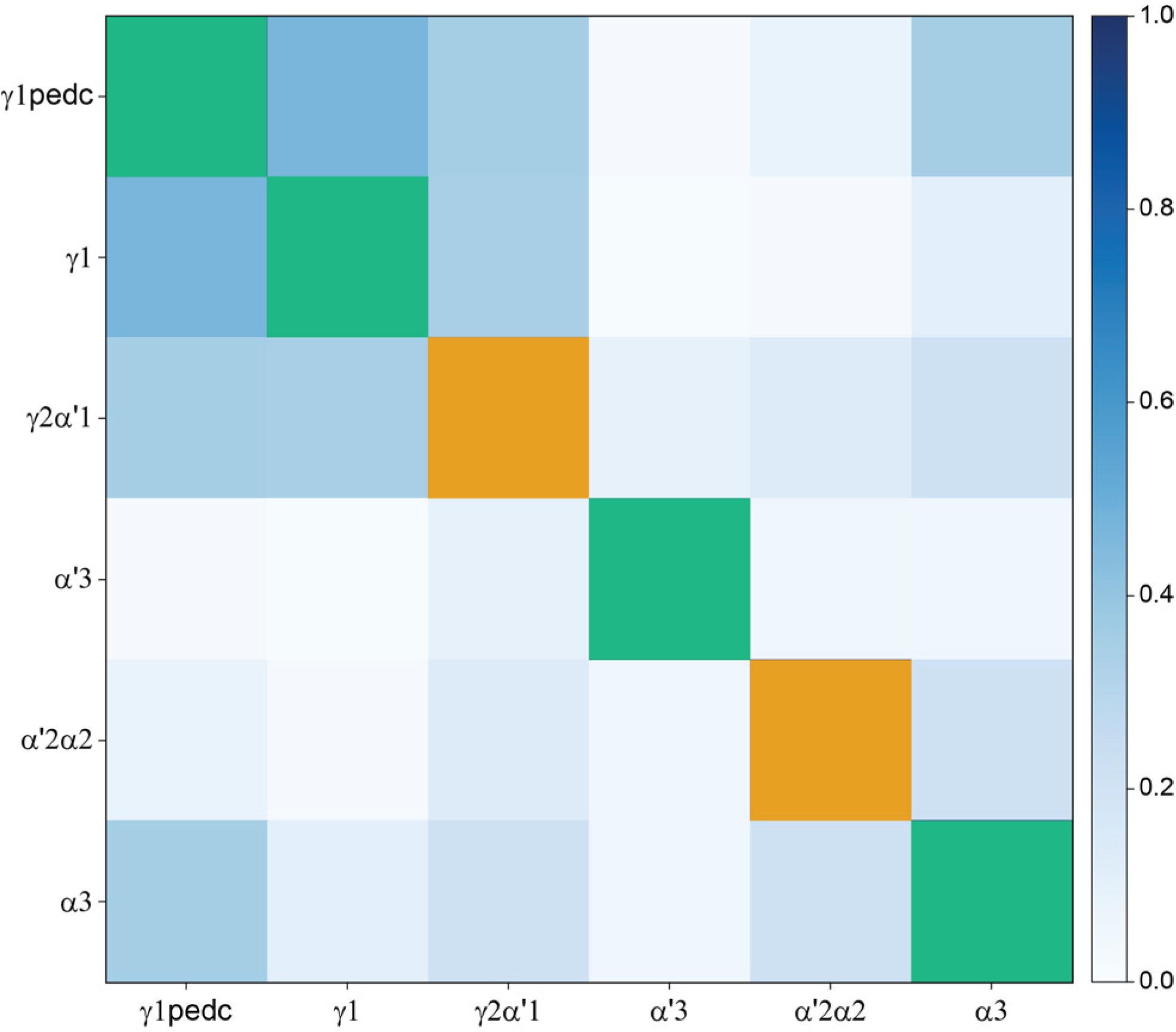
Similarity of inputs to PPL1 DANs. Expanded view of Figure 27 for PPL1 DANs (red square in Figure 27). Note that each PPL1 DAN receives direct feedback from MBONs, including the MBON from its own compartment. Colors on the diagonal indicate whether the given DAN receives feedback from MBONs in the same compartment (yellow) or from both the same and different compartments (green).

**Figure 27-figure supplement 2.**
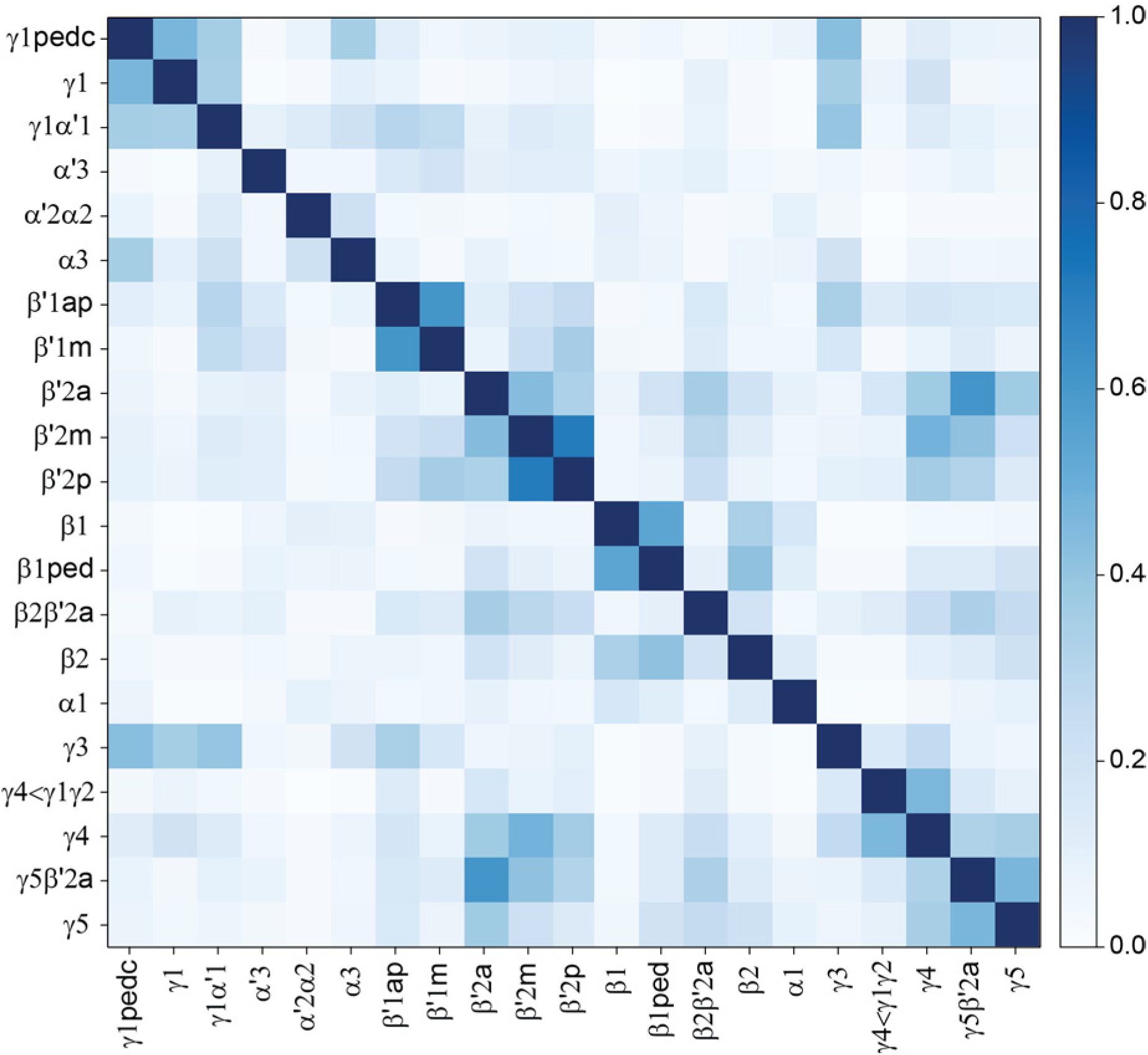
Similarity of inputs to DAN cell types. Average input similarity between DAN cell types computed after pooling the data for all cells of a given type, as compared to Figure 27 where each individual DAN is plotted separately.

**Figure 27-figure supplement 3.**
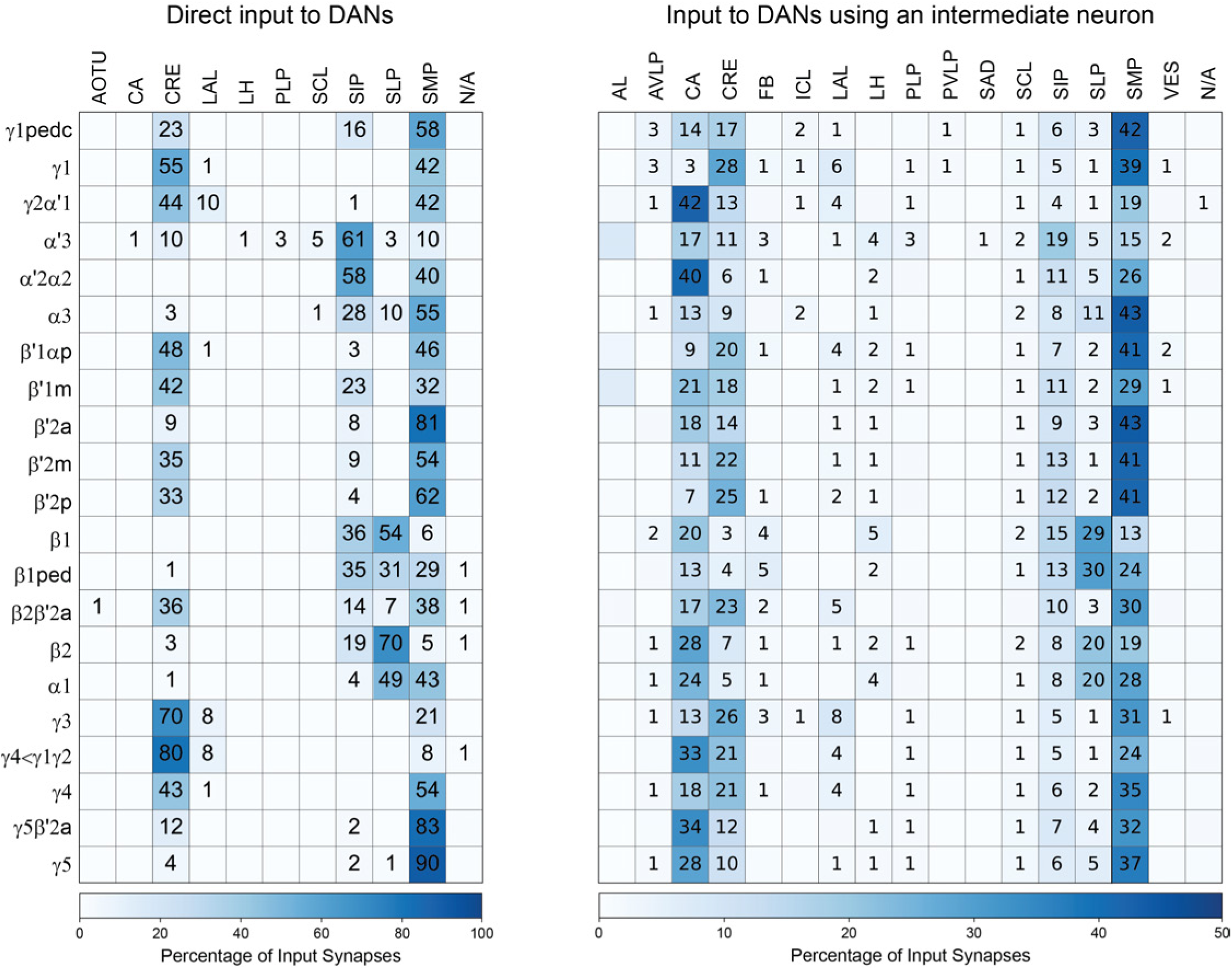
DAN input distribution by brain region. (Left) The value in each box indicates the percentage of the given DAN type’s input synapses that lie in the given brain region. Blank boxes indicate values of less than 1%. (Right) Distribution of brain regions providing input to DAN types using an intermediate interneuron. The value in each box indicates the effective input to the given DAN from the given brain region, mediated by one interneuron. Effective input is a measure that takes into account both the strength of the connection of each of the neurons that provide input to the DAN and the connection strength of other neurons to each of those input interneurons in each brain region. Effective input is computed by matrix-multiplying the inputs to the DANs and the inputs to those DAN-presynaptic neurons (normalizing both matrices so that inputs to all neurons sum to 1). Blank boxes indicate values of less than 1%. The two distributions are generally similar, but there are some clear differences. For example, direct input from CA is minimal but very prominent in input mediated by an interneuron; for example, the γ2α’1 PPL103 DAN receives over 40% of its indirect input from the CA.

**Figure 27-figure supplement 4.**
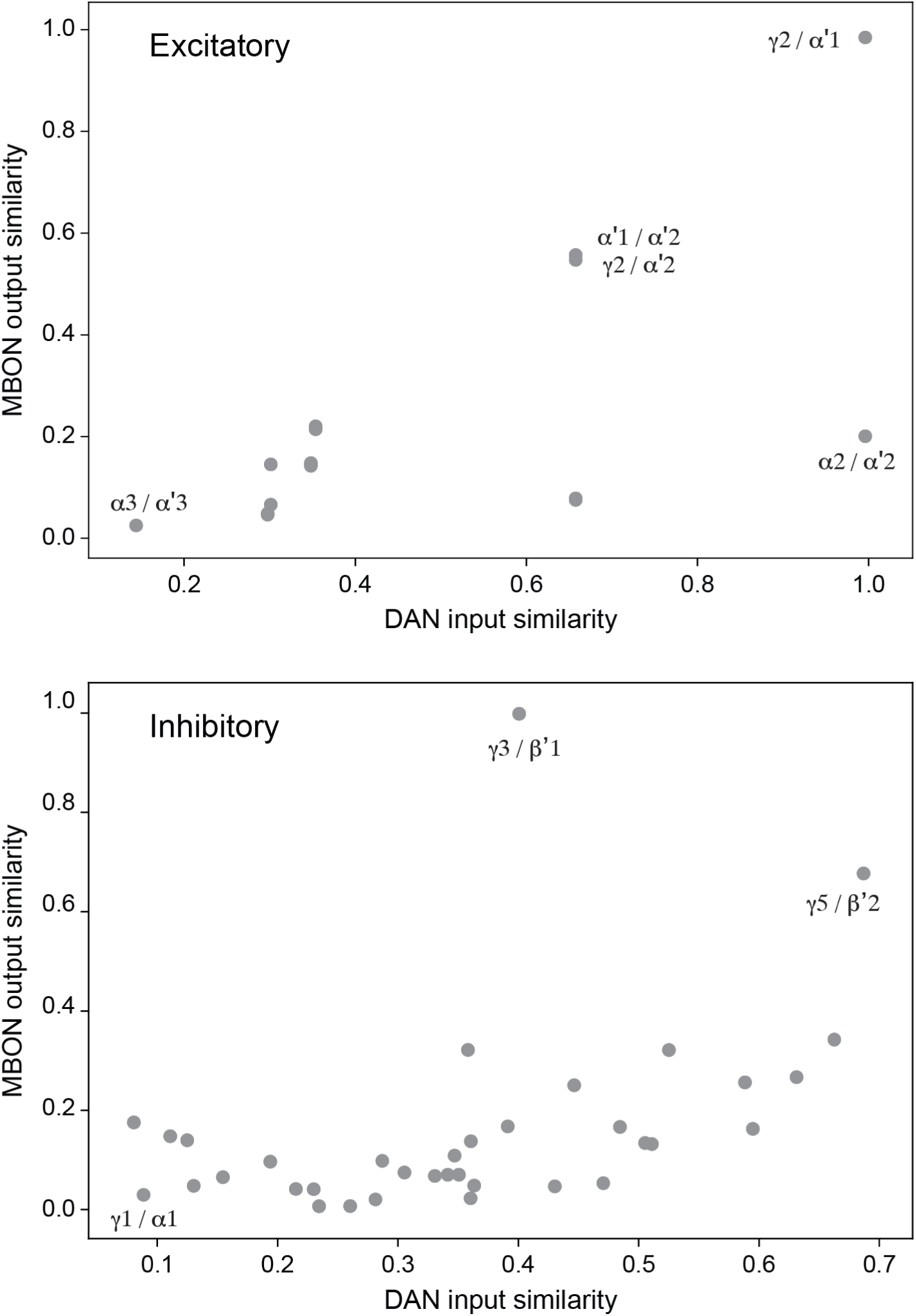
Comparing DAN inputs and MBON outputs reveals credit assignment by DANs. Scatter plot of DAN input similarity (data from Figure 27) and MBON output similarity (data from Figure 16, collapsed by compartment). Each point represents a pair of compartments; notable pairs are labeled in the figure. DAN input similarity is defined as the cosine similarity of the inputs to the DANs of two compartments. MBON output similarly measures the cosine similarity of the outputs from the MBONs in those compartments. The relationship between the two similarity measures indicates that compartments whose MBONs project to similar downstream targets have DANs that receive similar inputs. This structure suggests a form of “credit assignment” in which compartments whose MBONs control similar behaviors also receive similar reinforcement signals. Results are shown for compartments whose MBONs express excitatory (top, ACh) or putatively inhibitory (bottom, GABA and Glu) neurotransmitters.

**Figure 28-figure supplement 1.**
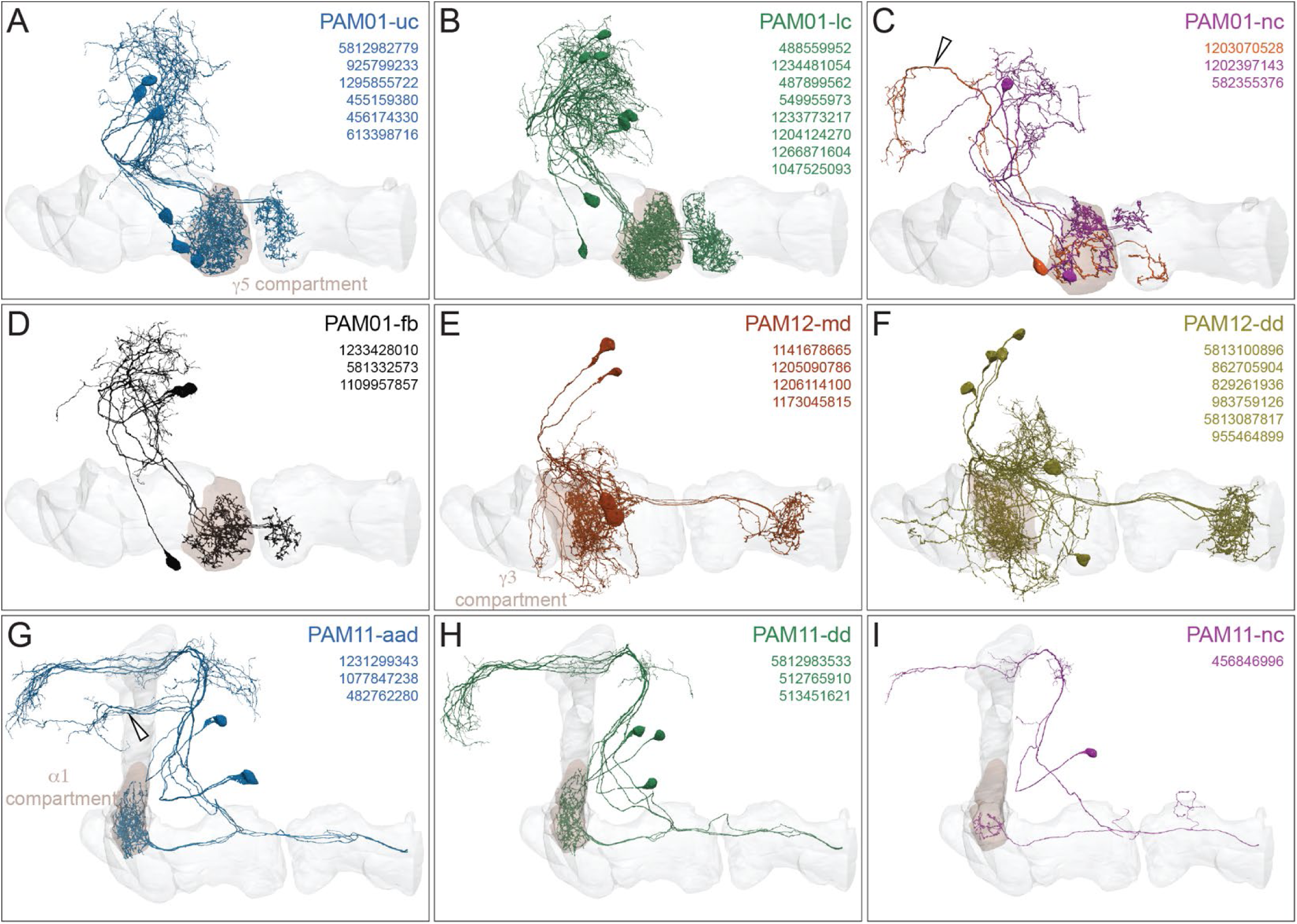
Anatomy of PAM01, PAM12, and PAM11 DAN subtypes. Morphologies of DAN subtypes discussed in Figure 28. Innervated MB lobes are shown in grey and the relevant compartment in brown. (A) Six PAM01-uc DANs (blue) project across the upper commissure (uc). (B) Eight PAM01-lc DANs (green) project across the lower commissure (lc). The dendritic field of PAM01-lc also extends slightly more anterior and ventral compared to PAM01-uc DANs (A). (C) Three non-canonical (nc) PAM01-nc DANs (magenta and orange) shown together. A single <u>PAM01-nc</u> DAN (orange, crossing the lc) has an additional dendritic branch in the SLP (indicated by arrowhead) and two other <u>PAM01-nc</u> DANs (1202397143, 582355376, crossing the uc) have axons confined to the dorsal or lateral portion of the γ5 compartment. (D) Three <u>PAM01-fb</u> DANs receive direct feedback (fb) from MBON01 (γ5β’2a) and their presynaptic fields are confined to the ventral portion of the γ5 compartment, in contrast to other PAM01 DANs. They also project through the lower commissure. Subtypes match those in Otto et al. 2020. (E-F) PAM12 DANs divide into (E) four medial dendrite (md) <u>PAM12-md</u> DANs (maroon) and (F) six dorsal dendrite (dd) <u>PAM01-dd</u> DANs (ochre) (also see Figure 33). (G-I) The three morphological groups of PAM11 DANs. These DANs are also shown in Video 32. (G) Three <u>PAM11-aad</u> DANs (blue) have an additional anterior dendrite (aad) anterior to the α lobe (arrowhead). (H) Three dorsal dendrite (dd) <u>PAM11-dd</u> DANs (green). (I) One non-canonical (nc) <u>PAM11-nc</u> DAN (magenta), which unlike the other PAM11 DANs, has an axonal field confined to the middle of the α1 compartment.

**Figure 30-figure supplement 1.**
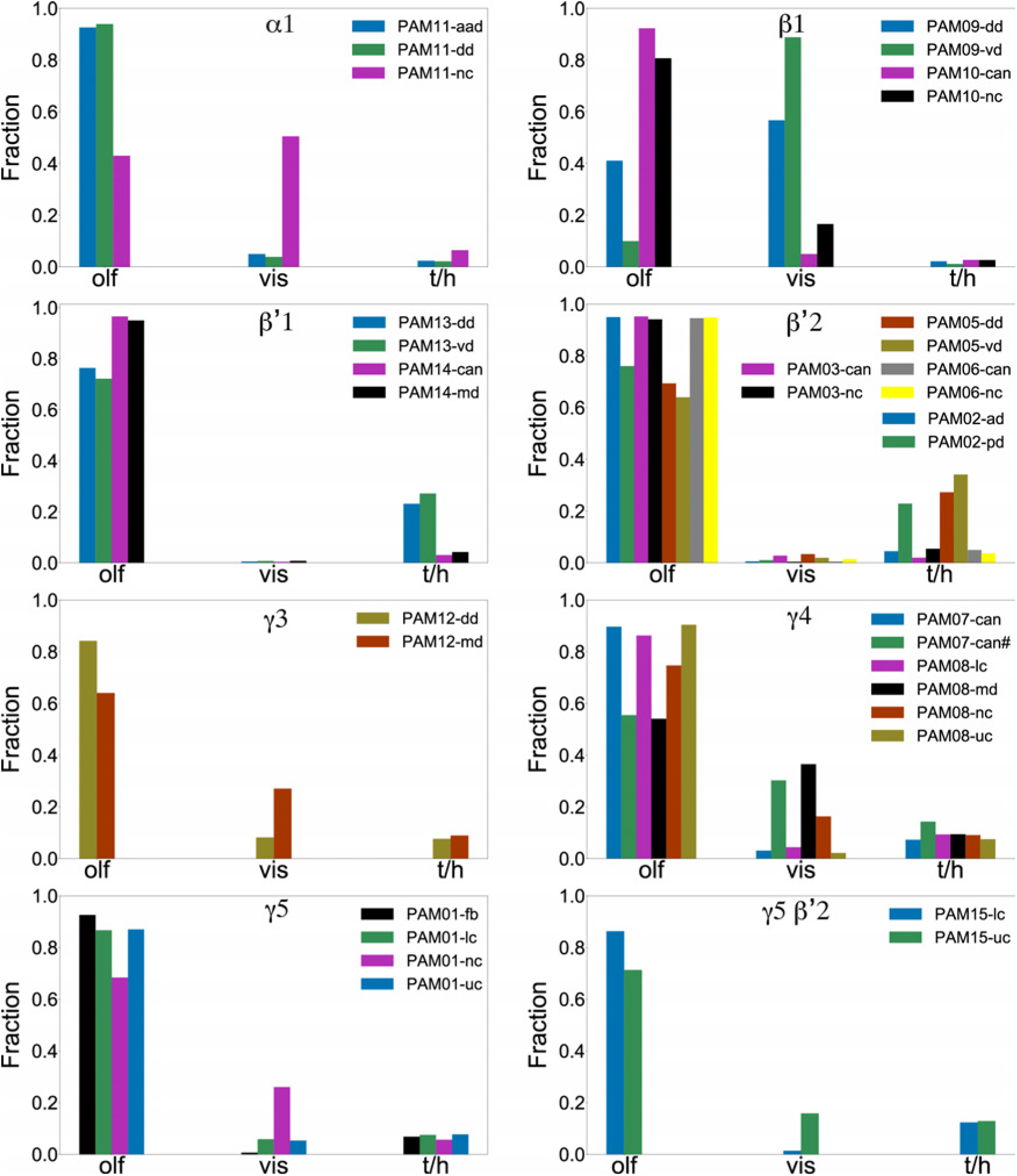
DAN subtypes within a compartment can be biased toward KC types representing different sensory modalities. These plots reveal that in many cases, selective innervation of KC subclasses by particular DAN subtypes within a compartment is organized according to the sensory modality represented by those KCs. Sensory modalities of KCs are assigned according to their PN inputs (derived from the same data as Figure 15-figure supplement 2A). In the indicated compartments, the DANs of a given type are divided into subtypes following criteria from Figure 28. For the PAM07-can (γ4<γ1γ2) subtype, one outlier, indicated by the # sign, which was not apparent with clustering either by morphology or input connectivity, was identified by spectral clustering of DAN-KC connectivity (Figure 29). The y-axis values indicate the fraction of each of the indicated DAN cluster’s output it provides to KCs representing three different sensory modalities. The contributions of each KC to this value are weighted by the number of input synapses they receive from DANs of the given type. Olf - olfactory, vis - visual, and t/h - thermo/hygrosensory.

**Figure 30-figure supplement 2.**
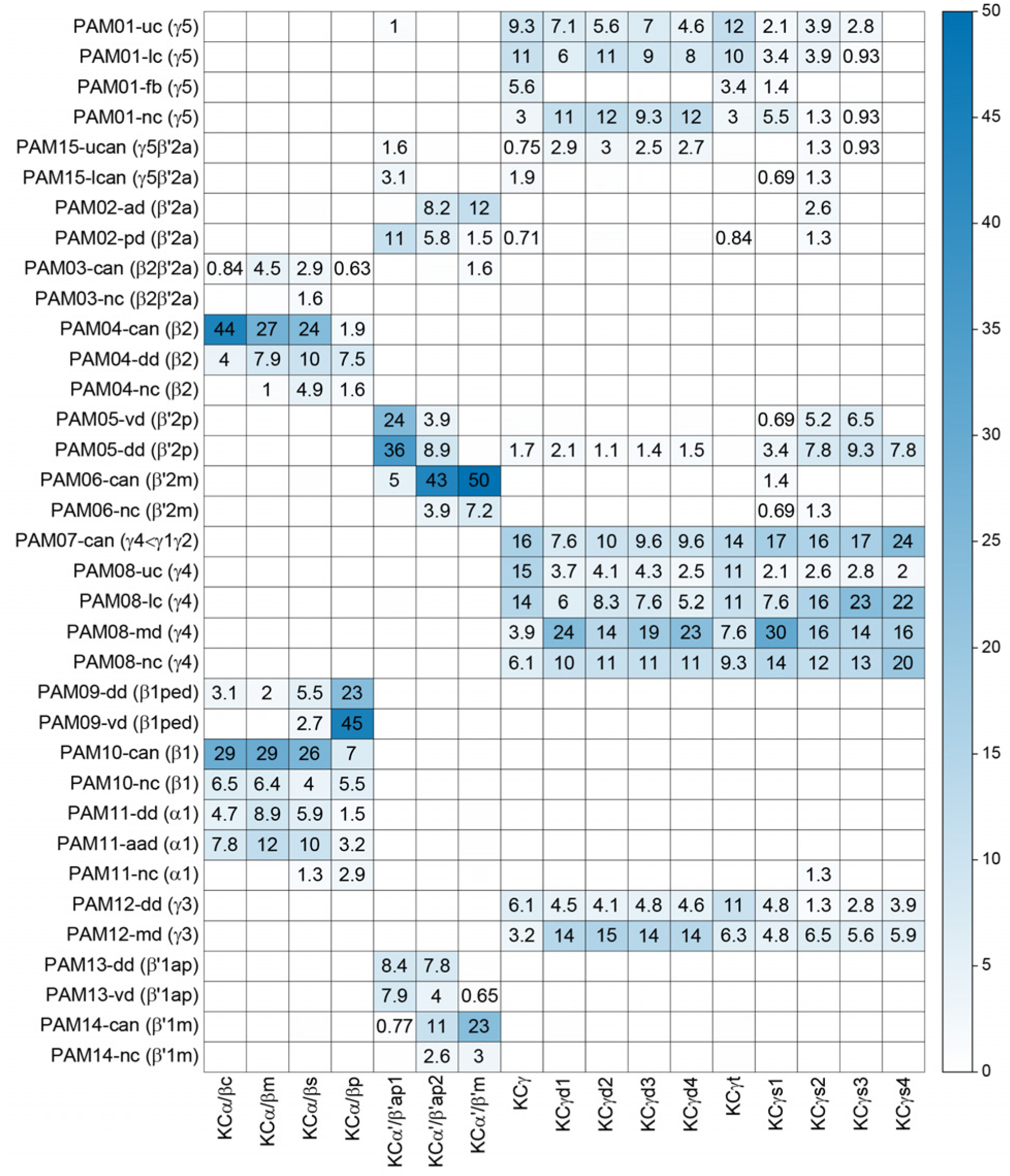
PAM DAN subtypes synapse onto specific KC classes within a compartment. DAN to KC connectivity matrix underlying Figure 30. DAN subtypes, defined by morphological and upstream connectivity clustering (from Figure 28), synapse onto specific types of KCs within a compartment. Each cell of the matrix shows the percentage of inputs provided by a particular DAN subtype, normalized by all DAN inputs to that KC type. DAN subtypes that have axons confined to portions of the same compartment often differ in their KC connectivity. Only connections constituting greater than 0.5% of the KC’s total DAN input in the compartment are shown.

**Figure 30-figure supplement 3.**
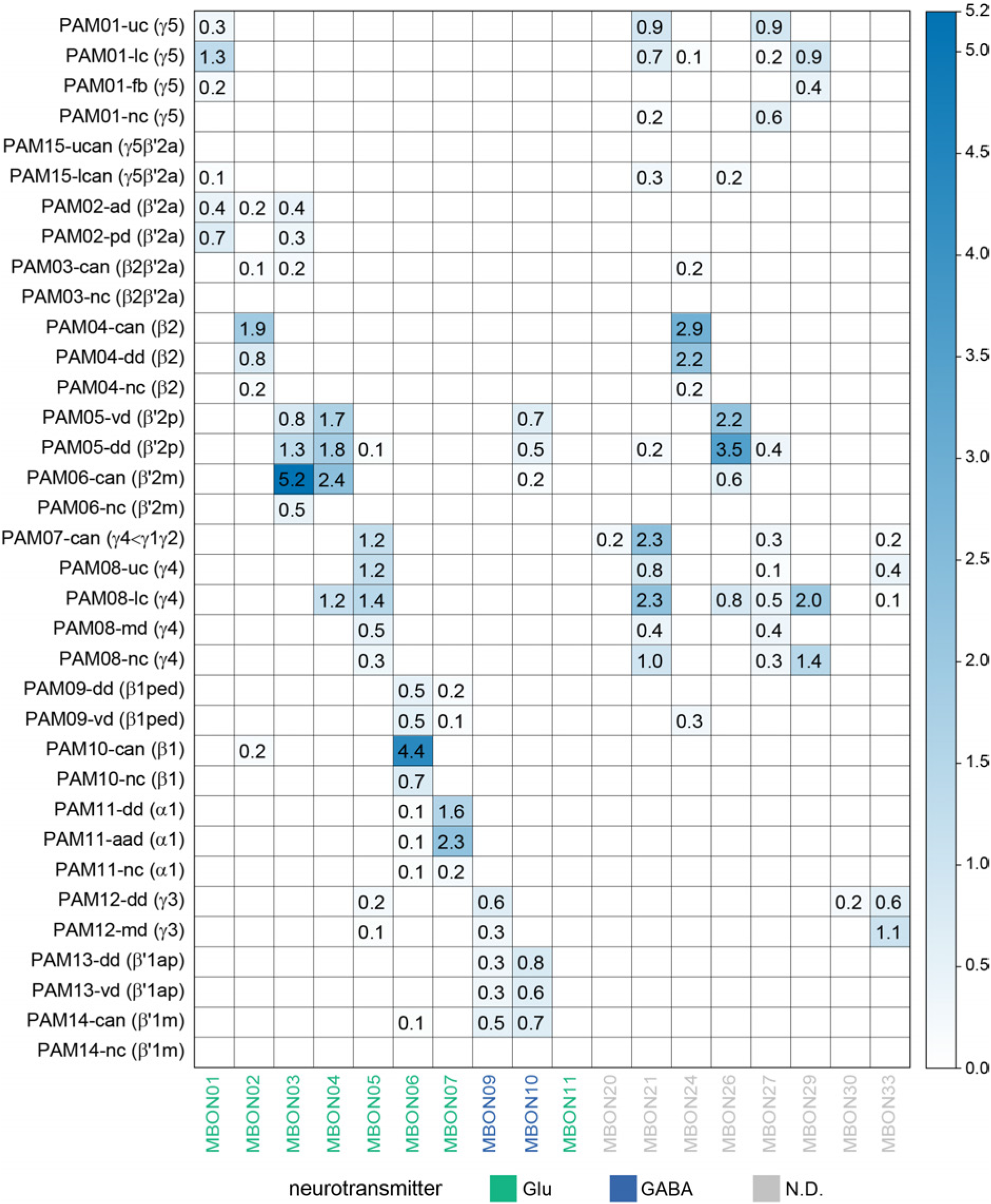
PAM DAN subtypes synapse selectively onto particular MBONs within a compartment. DAN to MBON connectivity matrix underlying Figure 30. DAN subtypes as defined by morphological and upstream connectivity clustering (from Figure 28) connect to specific MBONs within the respective compartment. Each cell of the matrix shows the percentage of input provided by the indicated DAN subtype, normalized by the MBON’s synaptic input. DAN subtypes that have axons confined to a specific portion of the same compartment often differ in their connectivity to MBONs. Only connections constituting greater than 0.1% of the synaptic input to each MBON are shown.

**Figure 32-figure supplement 1.**
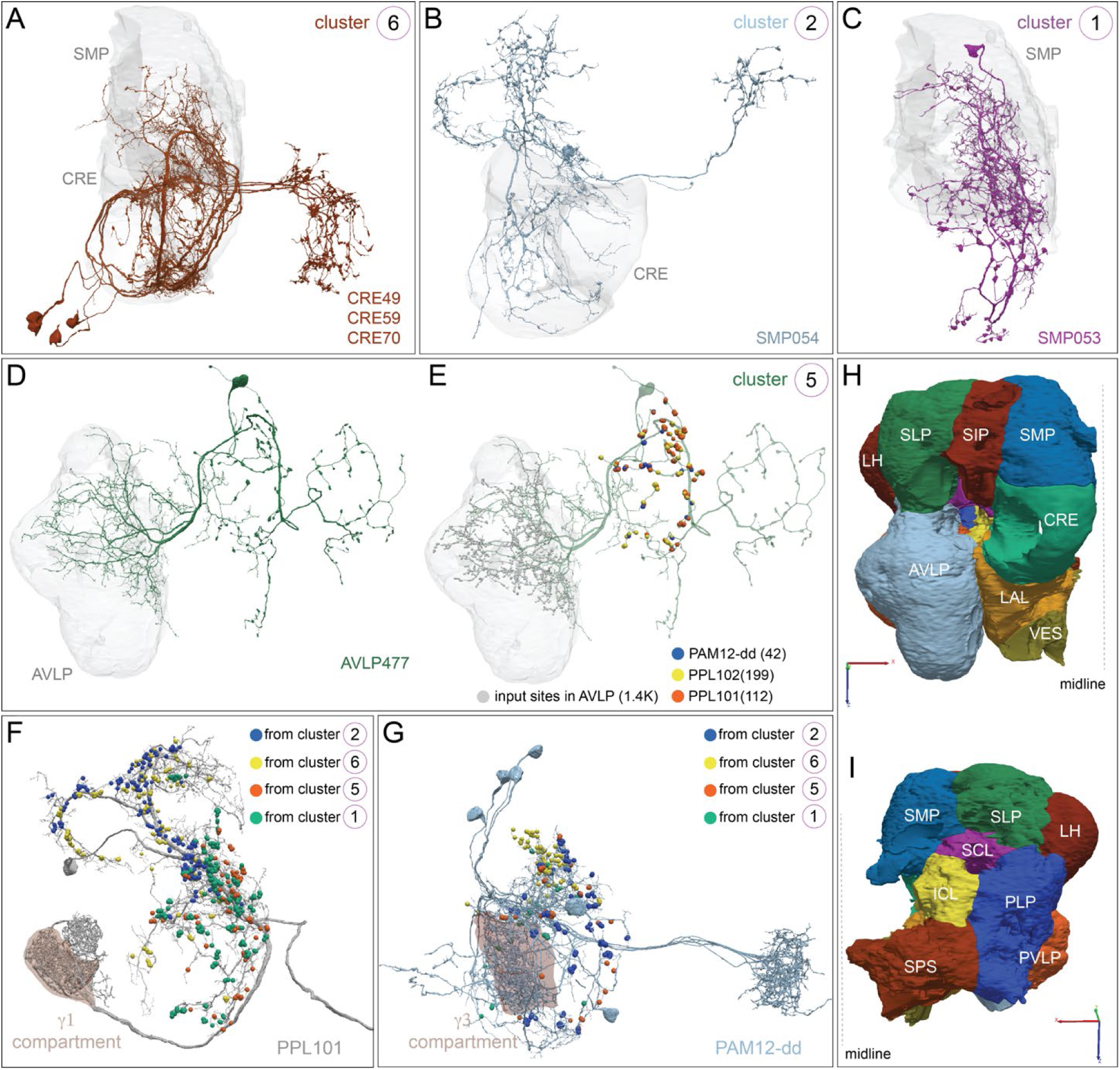
Strong inputs to PAM12-dd and PPL101 localize to different dendritic arbors. More detailed representations of connectivity between input clusters 6, 2, 1 and 5 (Figure 31) and PPL101 and PAM12-dd DANs. Neuropils in which DAN input neurons receive most of their inputs are shown in grey. (A-C) Neurons in cluster 6 (cell types <u>CRE49</u>, <u>CRE59</u>, <u>CRE70</u>), cluster 2 (cell type <u>SMP054</u>) and cluster 1 (cell type <u>SMP053</u>), shown in Figure 32 D-E are shown here without their output synapses marked to allow better visibility of their morphologies. (D and E) Cluster 5 (cell type <u>AVLP477</u>) shown without synapses (D) and with synapses marked (E) also synapses onto PAM12-dd (blue dots), PPL101 (γ1pedc; orange dots) and PPL102 (γ1; yellow dots) DANs; synapse numbers are given in parentheses. (F) Detail of inputs to PPL101, whose outputs are in the γ1 compartment (brown) and the core of the pedunculus. Inputs from valence-specific clusters 2, 6, 5 and 1 (synaptic input color-coded as indicated) localize to specific branches of PPL101. (G) Detail of inputs to PAM12-dd, whose outputs are in the γ3 compartment (brown) and which receives input from clusters 2, 6, 5 and 1 (synapses color-coded as indicated). While clusters 5 and 2 inputs are evenly distributed, cluster 6 and 1 localize to the dorsal portion of the PAM12-dd’s dendrites. (H and I) Two views of the input neuropils discussed in Figures 32-36; see Figure 2-figure supplement 1 for a key to abbreviations. Connections contributing less than 0.5% of a neuron’s total dendritic input have been excluded.

**Figure 33-figure supplement 1.**
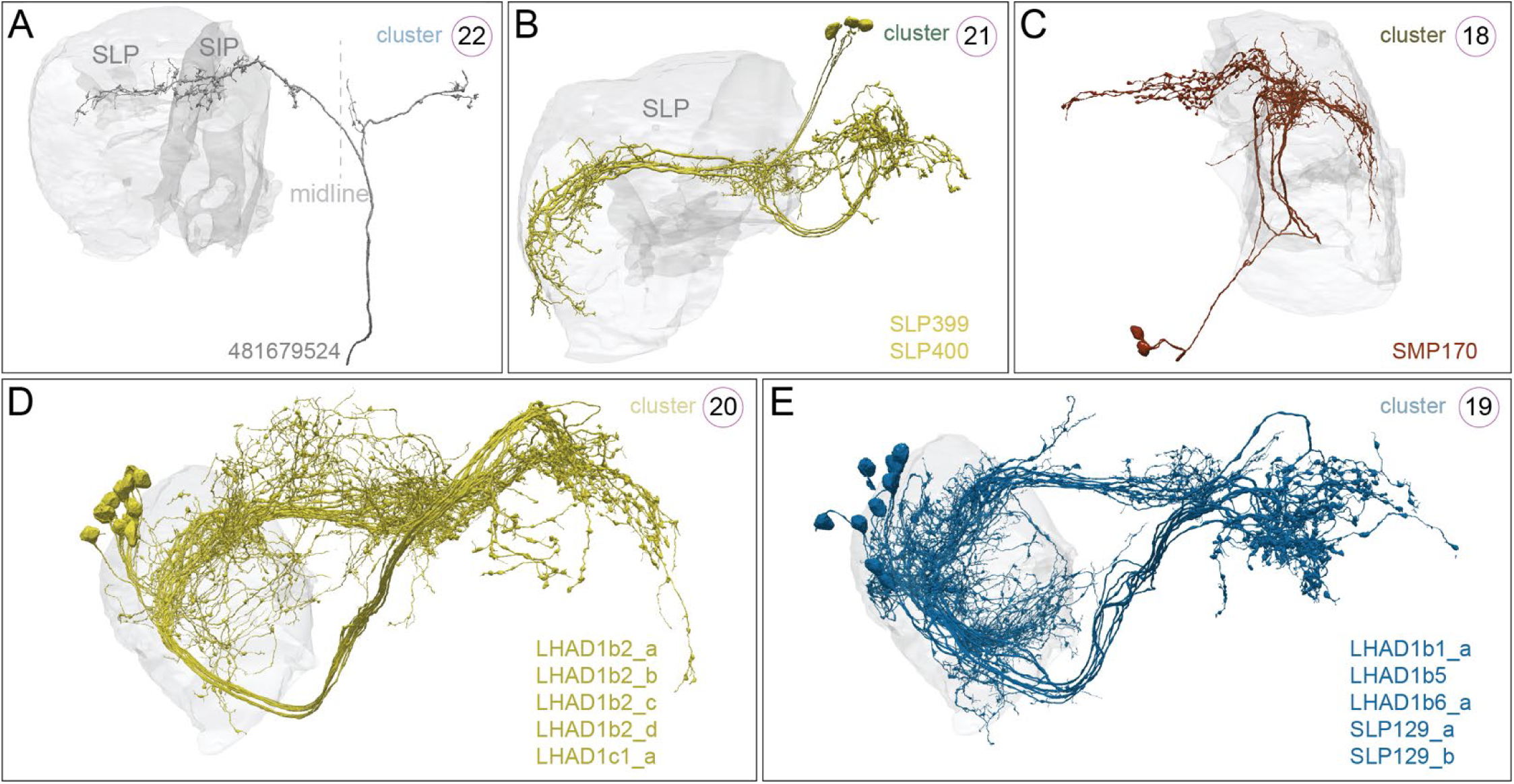
More information on the cell types shown in Figure 32. Panels show the neurons shown in panels (B-F) of Figure 33 with their corresponding neuPrint IDs or cell types. Links to neuPrint are as follows: (A) <u>481679524</u>; (B) <u>SLP399</u>, <u>SLP400</u>; (C) <u>SMP170</u> (D) <u>LHAD1b2_a</u>; <u>LHAD1b2_b</u>; <u>LHAD1b2_c</u>; <u>LHAD1b2_d</u>; <u>LHAD1c1_a</u> (E) <u>LHAD1b1_a</u>; <u>LHAD1b5</u>; <u>LHAD1b6_a</u>; <u>SLP129_a</u>; <u>SLP129_b</u>.

**Figure 35-figure supplement 1.**
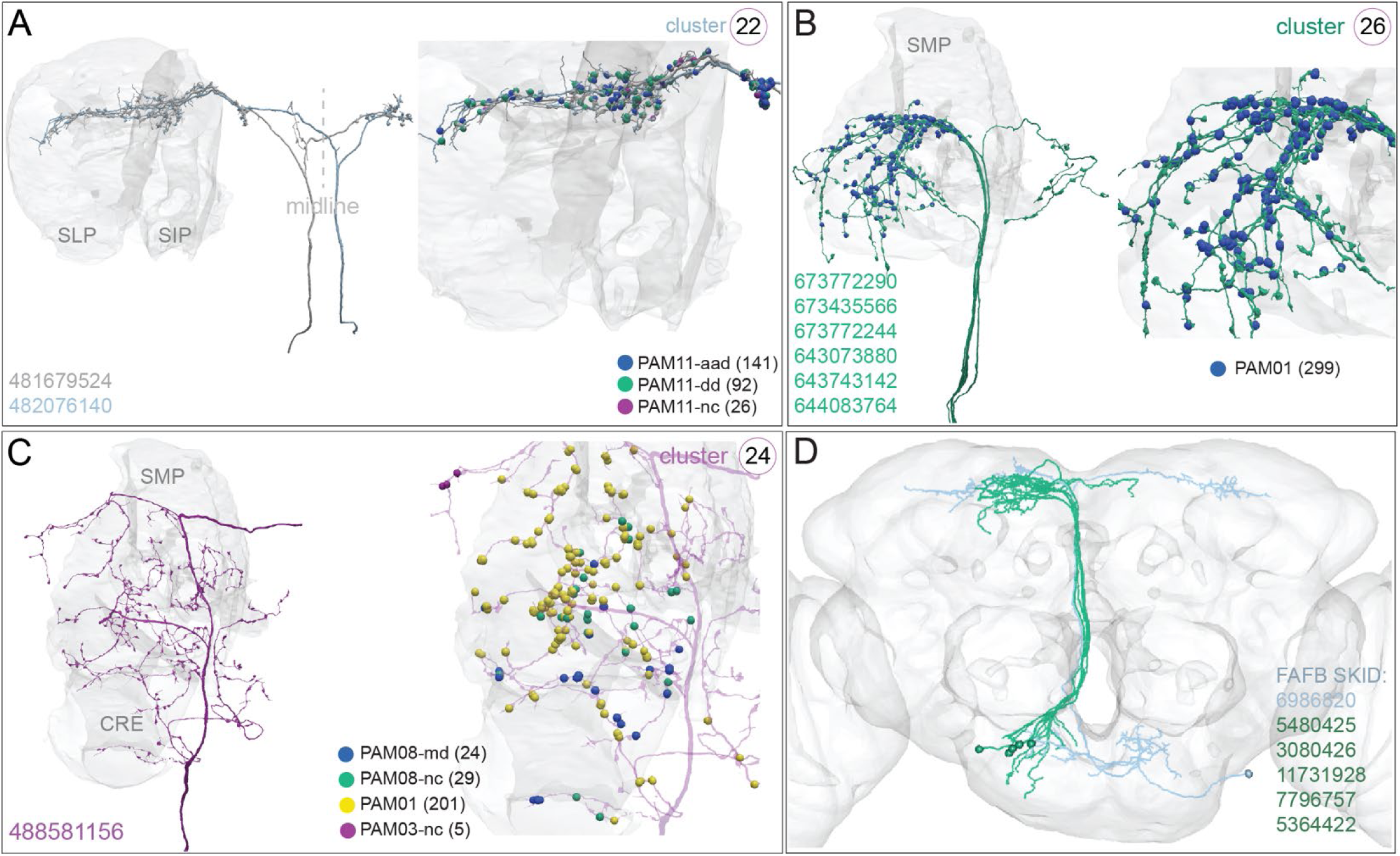
Most strongly connected SEZON clusters exclusively synapse onto positive valence PAM DANs. (A-C) Further examples of clusters consisting of SEZONs that connect to specific DAN subtypes. In each panel the neuronal morphology is shown on the left with the brain regions where they contact DANs shown in grey, and on the right a higher magnification view of their output synapses onto specific DAN types are shown, color coded, with synapse numbers given in parentheses. The threshold for connectivity is 0.5% of the DANs total inputs (Figure 31). (A) The neurons of SEZON <u>cluster</u> 22 (481679524) and its contralateral partner (482076140) branch in the SMP and SLP and synapse onto all 3 PAM11 subtypes (compare to Figure 33). (B) The six neurons in SEZON <u>cluster 26</u> branch in the SMP and connect to positive valence γ5 DAN subtypes (blue) and weakly to PAM15 (γ5β′2a) and PAM02 (β′2a) subtypes. (C) The single neuron in SEZON <u>cluster 24</u> branches in the SMP and connects specifically to PAM08-md (γ4; blue) and PAM08-nc subtypes (green), all PAM01 (γ5) subtypes (yellow), as well as the PAM03-nc (β2β′2a) subtype (magenta). (D) The axons of SEZONs in (A and B) can be matched to those in the FAFB EM volume (Zheng et al. 2018). This registration permits the retrieval of their dendritic fields, which reside in tissue that is missing from the hemibrain dataset. FAFB matches to hemibrain SEZONs are shown and are colored to match their hemibrain counterparts in panels (A) and (B). The dendritic fields differ, suggesting these SEZONs convey distinct information. Connections contributing less than 0.5% of a neuron’s total dendritic input have been excluded.

**Figure 36-figure supplement 1.**
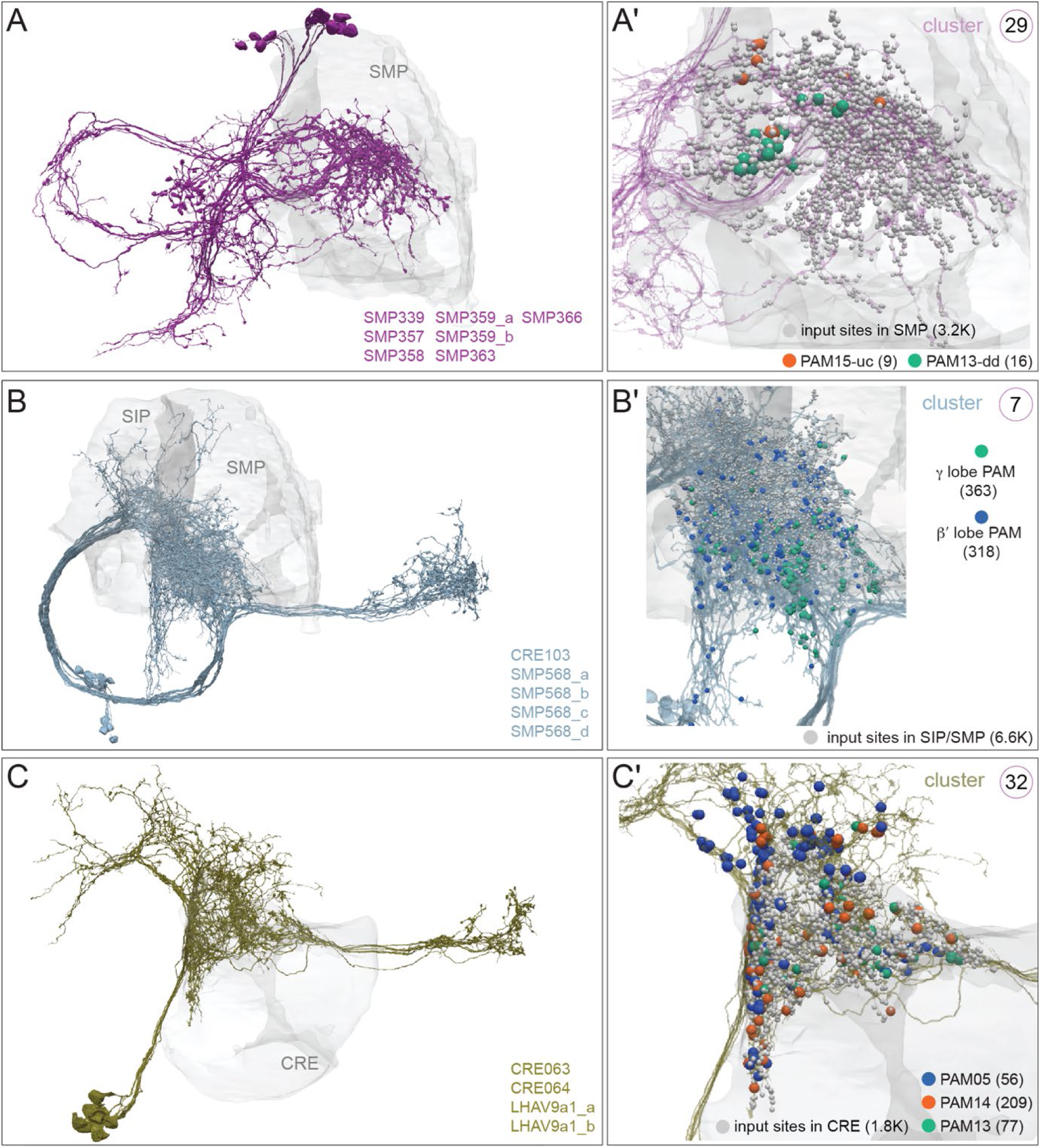
Inputs specific to PAM DANs. Neurons providing strong input to DANs of the PAM cluster are shown. Neuropil areas indicated (grey) and synapses to DANs are color coded; synapse numbers are given in parentheses. (A, A′) The nine neurons of <u>cluster 29</u> fall into seven cell types and receive inputs in the SMP and connect to PAM15-uc (γ5β′2a; red) and the PAM13-dd (β′1ap; green). (B, B′) The 13 neurons of <u>cluster 7</u> fall into five cell types and receive inputs in the SMP and SIP and connect strongly to PAM DANs that are restricted to the β′ (blue synapses) and the γ lobes (green synapses): PAM13 (β′1ap), PAM06 (β′1m), PAM02 (β′2a), PAM06 (β′2m), PAM05 (β′2p), PAM03 (β2β′2a) and PAM12-md (γ3), PAM08 (γ4), PAM01 (γ5), PAM(γ5β′2a). (C, C′) <u>Cluster 32</u> has nine neurons of four cell types that receive some inputs in the CRE. Cluster 32 provides strong input to PAM14 (β′1) subtypes (red synapses) and is also upstream of the PAM05-vd (β′2p) subtype (blue synapses) and all PAM13 (β′1ap) subtypes (green synapse). The threshold for connectivity is 0.5% of the DANs total inputs (Figure 31).

**Figure 36-figure supplement 2.**
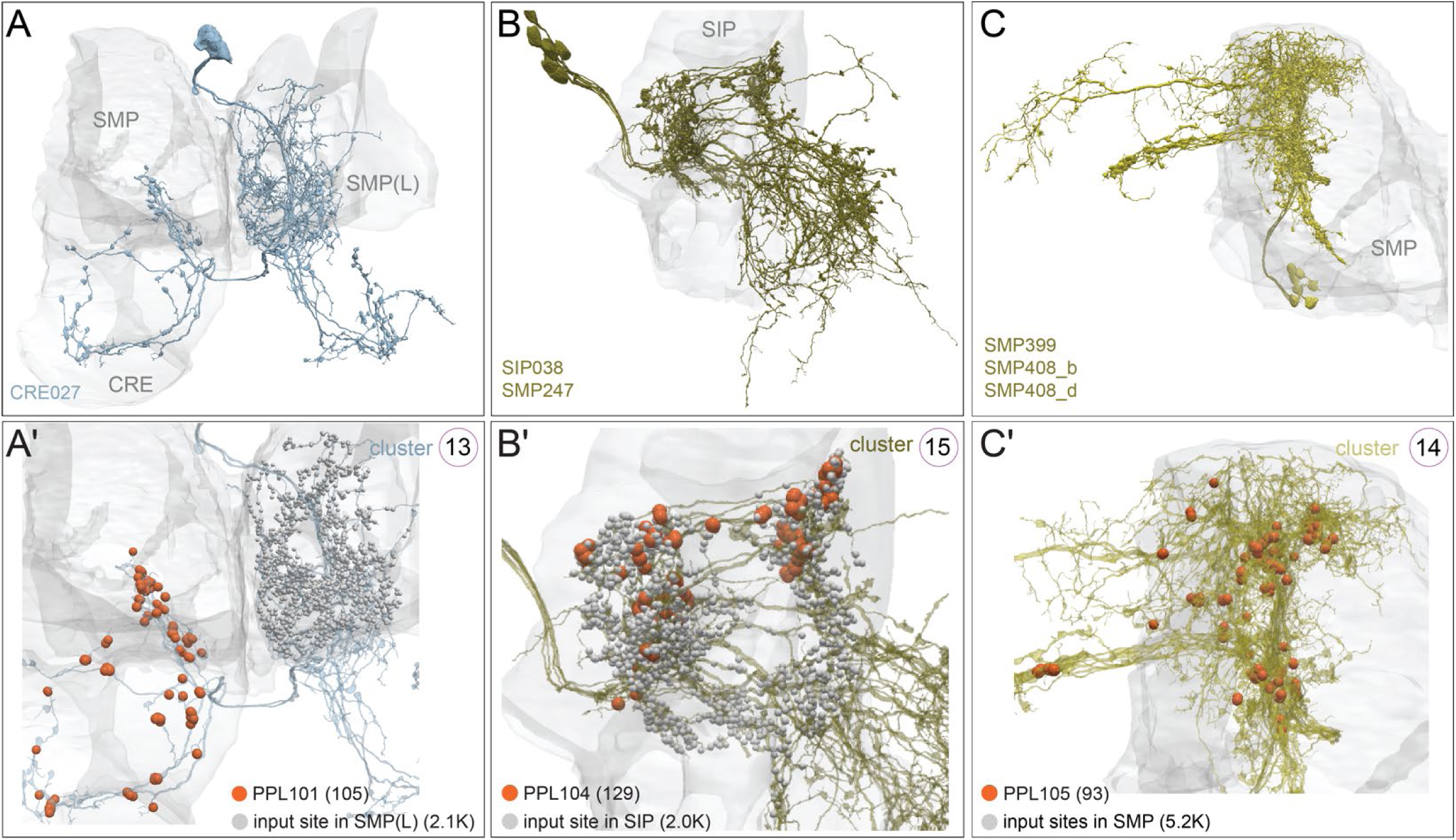
Inputs specific to PPL1 DANs. Neuropil areas indicated (grey) and synapses to DANs are color coded; synapse numbers are given in parentheses. (A, A′) The two neurons (light blue) of <u>cluster 13</u> receive inputs in the contralateral SMPs and connect to the negative valence PPL101(γ1pedc; red) DANs. (B, B′) <u>Cluster 15</u> has six neurons receiving inputs in the SIP and provide strong input to PPL104 (α′3). (C, C′) The seven neurons of <u>cluster 14</u> receive inputs in the SMP and are strongly connected to the PPL105 (α′2α2). The threshold for connectivity is 0.5% of the DAN’s total inputs (Figure 31).

**Figure 37-figure supplement 1.**
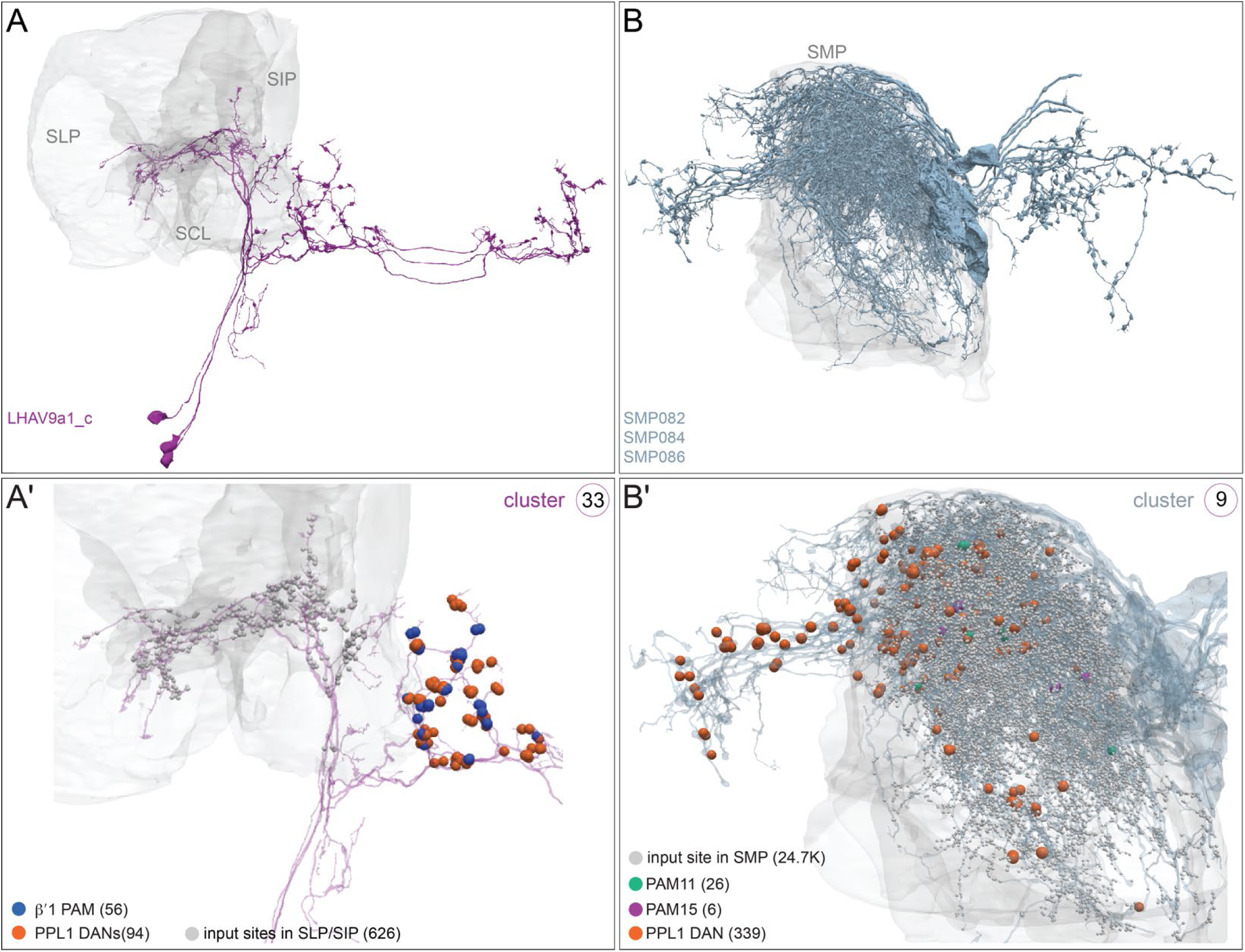
Additional examples of neurons providing common input to both PAM and PPL1 DANs. Neuropil areas where neurons of clusters 33 and cluster 9 receive input are shown in grey and their synapses onto DANs are color coded; synapse numbers are given in parentheses. (A, A′) The three neurons of <u>cluster 33</u> receive inputs in the SIP, SLP, and SCL and connect to both PAM13 (β′1ap) subtypes (blue synapses) and to a collection of PPL1 DANs (PPL101, PPL102, PPL103 and PPL104; red synapses). (B, B′) <u>Cluster 9</u> has six neurons that receive inputs in the SMP and connect to the negative valence PPL1 DANs (PPL101, PPL102, PPL105 and PPL106; red synapses). Cluster 9 is also upstream, but more weakly, of positive valence PAM DANs (dark blue synapses) including all PAM11 subtypes (green synapses), the PAM15 subtypes (magenta synapses) and more weakly to PAM01-lc and PAM01-fb subtypes.

## Video Thumbnails

While Videos are referred to in the text, this version of the manuscript does not include the Videos themselves.

On the following four pages, we provide thumbnails of the first frame of each Video to give the reader a glimpse of their content.

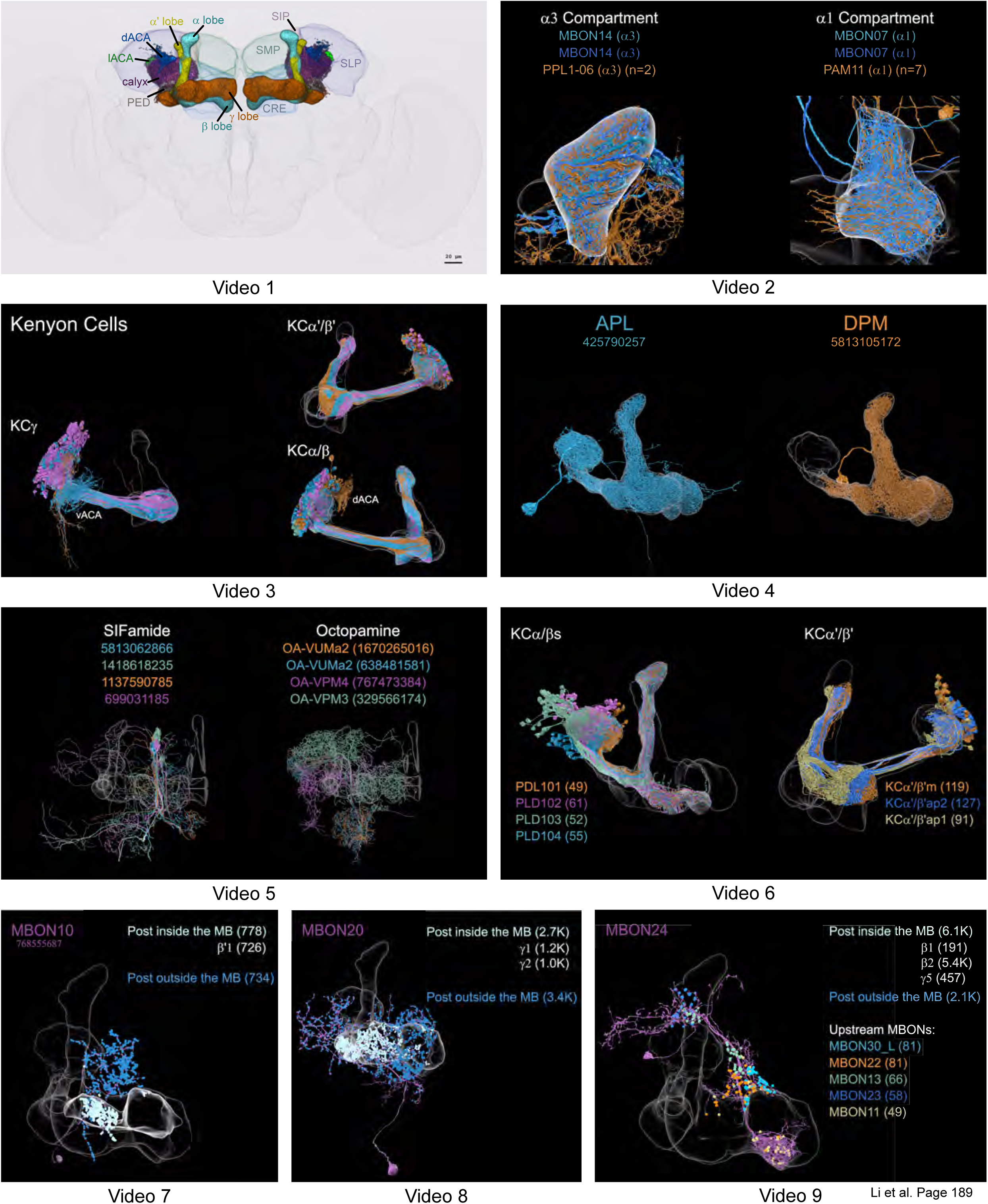
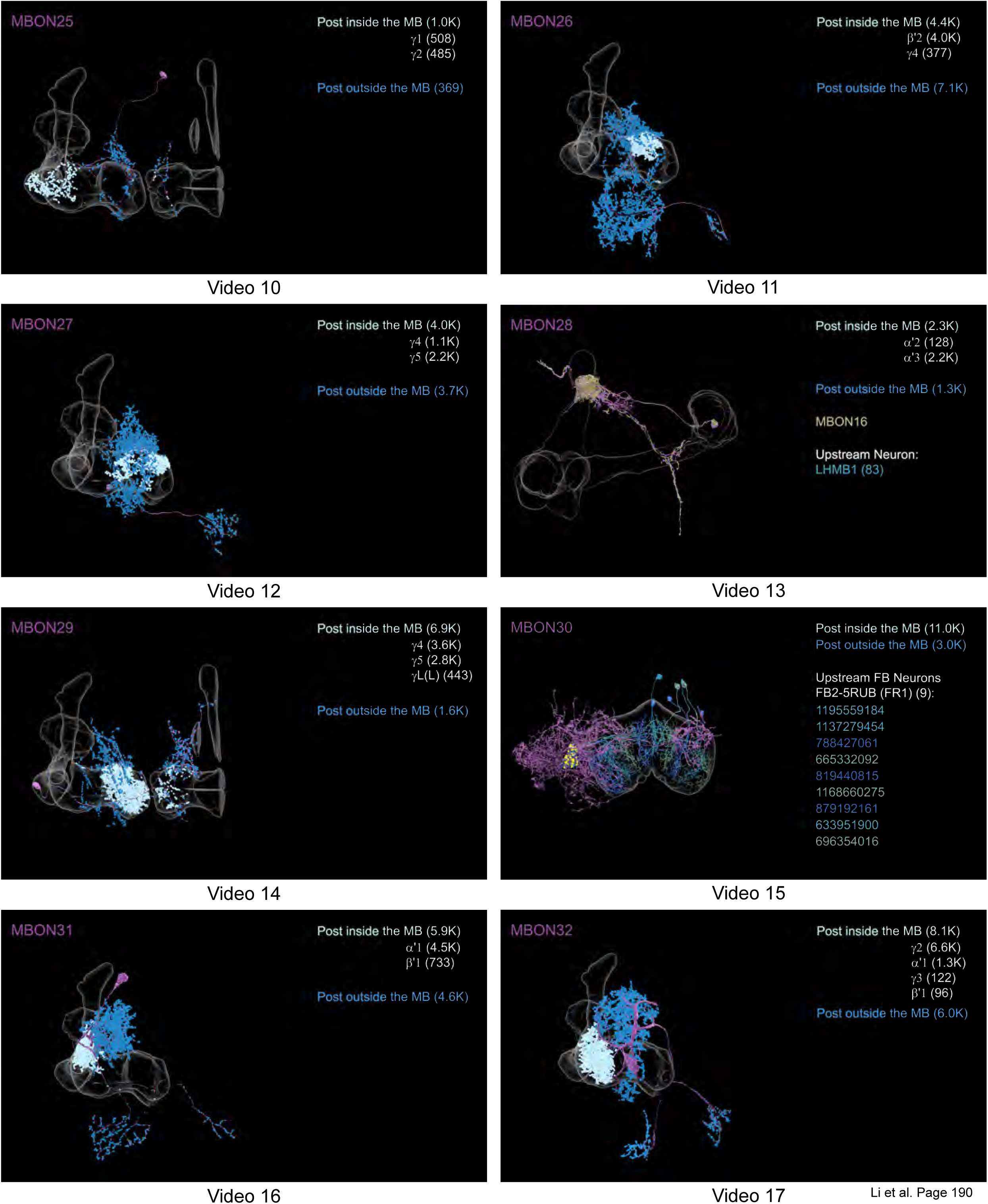
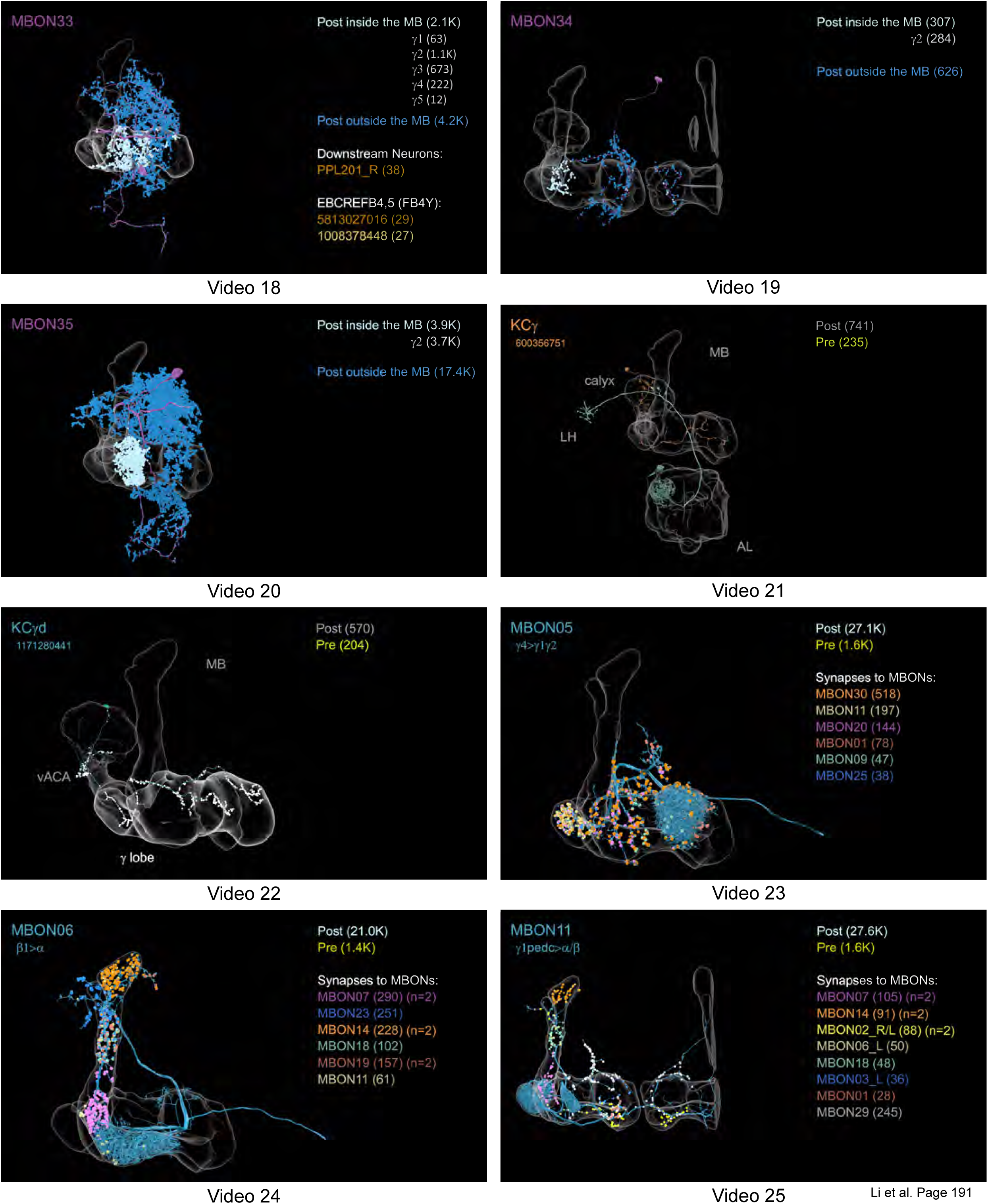
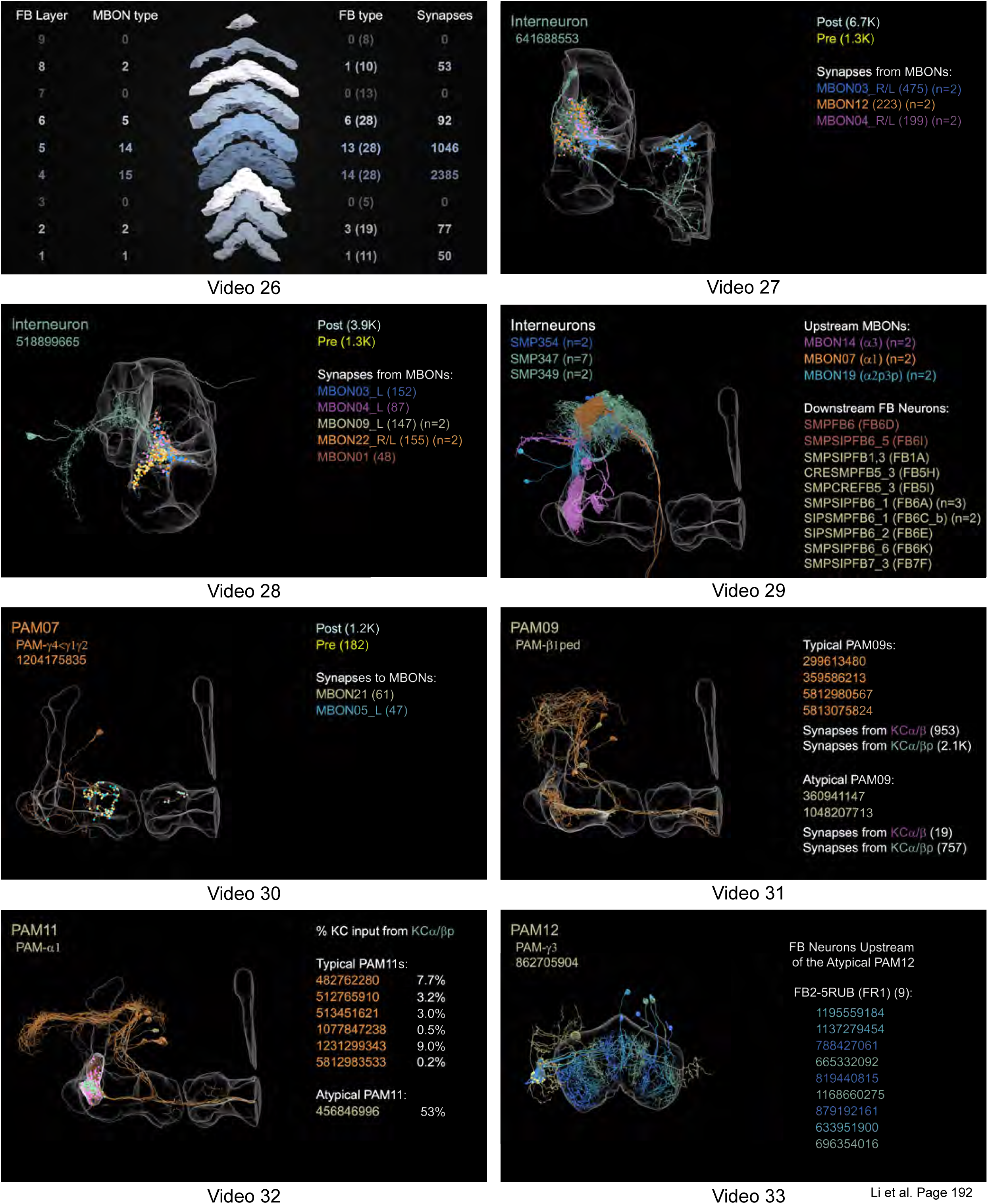

